# Autophagy flux during human aging is sex- and cell type-specific, and is associated with physical fitness

**DOI:** 10.64898/2026.05.15.725565

**Authors:** Tatiana M. Moreno, Stephanie R. Heimler, Ryan J. Moran, Hava Shoshana Barkai, Lina Scandalis, Larissa Traxler, Ashley Neil, Stephen Dozier, Jaclyn Bergstrom, Sanjeev S. Ranade, Anne G. Bang, Jerome Mertens, David Wing, Anthony J. Molina, Caroline Kumsta

## Abstract

Autophagy is widely proposed to decline with age; however, direct evidence for this across cell and tissue types in humans remains limited. Furthermore, it remains unknown whether interventions that improve physiological health during aging can modify autophagic activity in humans. Here, we performed transcriptomic and functional autophagy analyses across subject-matched human cell types from a healthy aging cohort spanning the adult lifespan. RNA-seq of primary dermal fibroblasts and induced neurons (iNs) revealed increased transcription of many autophagy-related genes with age, most markedly in fibroblasts. The impact of age on autophagic activity, measured using autophagy flux assays, was cell type- and sex-dependent, and uncoupled from autophagy-gene transcription. Autophagy flux decreased with age in male fibroblasts, was unchanged in female fibroblasts, and increased in female iNs. In freshly isolated peripheral blood mononuclear cells (PBMCs), autophagy flux became more heterogeneous with age and trended higher in older individuals, independent of sex. Although autophagy flux levels did not match across different cell types, higher autophagy flux in all cell types was associated with reduced physical function in older adults (≥70 years). Importantly, autophagy flux decreased following 12 weeks of mild exercise in parallel with improved physical function. These findings indicate that autophagy is regulated in a cell type-, sex-and physiological function-dependent manner during human aging, and highlight PBMC autophagy flux as a potentially modifiable, blood-accessible readout of physiological state in older adults.

## INTRODUCTION

Aging is characterized by the progressive loss of cellular and organismal health, driven by the accumulation of molecular damage, chronic stress and impaired repair mechanisms^1^. Among the cellular processes that maintain homeostasis during aging, macroautophagy, hereafter ‘autophagy’, is a conserved lysosomal degradative pathway that supports proteostasis, organelle quality control, nutrient recycling and cellular stress adaptation^2–4^. Consistent with this central role, disabled autophagy has been identified as a primary hallmark of aging^1^, and evidence from model organisms and post-mortem human tissues have suggested that autophagy declines with age^5–7^. Reduced autophagy has been linked to the pathogenesis of age-associated diseases including neurodegenerative diseases^8^, and interventions that enhance autophagy extend lifespan and healthspan in multiple model systems^9–13^.

However, autophagy is not uniformly reduced during aging. Autophagic activity can increase in specific cellular states, such as senescence and cancer^14,15^, and transcriptional and static readouts suggest that autophagy dynamics can vary across tissues, cell types and between sexes during aging^16,17^. Consistent with this, recent work demonstrates cell type- and sex-specific differences in chaperone-mediated autophagy across murine tissues, although evidencing an overall trend toward decline with age^18^. However, whether cell type- and sex-specific changes occur across human tissues remains unclear, in part due to the difficulty of directly measuring autophagic activity in humans. Autophagy is a multi-step process, and commonly used ‘static’ readouts, including autophagosome abundance, levels of the autophagosome-associated protein LC3/Atg8-II, selective autophagy receptors, and autophagy-gene expression cannot distinguish between increased autophagosome formation and impaired degradation^19^. Direct measurement of autophagy flux, achieved by pharmacological inhibition of lysosomal degradation, is not feasible *in vivo* in humans^20^. As a result, how autophagic activity is regulated across human tissues and physiological states during aging remains unclear.

To address these limitations, accessible human cell models that retain age-associated features provide an opportunity to examine autophagy across tissues within the same individuals. Primary dermal fibroblasts preserve transcriptional signatures of aging^21,22^, while induced neurons (iNs) derived from these cells enable investigation of neuronal aging in a subject-matched context^19,23–26^. In parallel, peripheral blood mononuclear cells (PBMCs) provide a minimally invasive system to assess autophagy in physiologically relevant conditions, and offer potential for biomarker development^27–31^. However, these approaches have not been integrated to assess functional autophagy across multiple human cell types within the same individuals.

Consequently, the relationship between autophagy flux and physiological health in human aging remains poorly defined. It is unknown whether flux is coordinated across cell types within the same individual, whether variation in flux across individuals reflects differences in physiological state or fitness, and whether it can be modified by behavioral intervention. These questions have direct relevance to whether autophagy flux could serve as a meaningful marker of aging trajectories.

Here, we performed integrated transcriptional analyses of autophagy in primary dermal fibroblasts and iNs across a healthy human cohort spanning the adult lifespan. We uncovered age-associated changes in autophagy-gene transcription across cell types. Moreover, we found that transcriptional and functional autophagic activity were uncoupled, and that autophagy flux was altered sex- and cell type-specifically with age. To establish the physiological relevance of autophagy in human aging, we examined relationships between autophagy flux and physiological health measures, finding that increased autophagy flux in PBMCs, fibroblasts and iNs is associated with reduced physical function in adults over 70 years old. Finally, we performed a pilot intervention to test whether autophagy flux in PBMCs is modifiable by mild exercise in older adults, and found that exercise reduced elevated autophagy flux to more youthful levels. Together, these studies provide a comprehensive assessment of autophagy across human tissues, link autophagic activity to physiological health and fitness, and demonstrate its modulation in response to behavioral intervention during aging.

## RESULTS

### Autophagy-gene transcription is upregulated differently with age in human primary dermal fibroblasts and induced neurons (iNs)

Aging is associated with robust transcriptomic changes in human primary dermal fibroblasts^21,22,32^. To define age-associated changes in autophagy-gene transcription at a global level, we performed bulk RNA-seq on low passage (<10) primary dermal fibroblasts from young and older healthy adult donors (young: 23-33 years, N = 4; older: 64-72 years, N = 5) from the San Diego Nathan Shock Center Clinical Cohort (SHOCK; **Tables 1**, **S1**), a well-characterized cohort of healthy individuals across the age-span who self-reported to be free of chronic disease^33^. Principal component analysis (PCA) of global fibroblast transcriptomes separated samples by age (**Fig. 1A**), indicating retention of robust age-associated transcriptional signatures^21,34^. Differential expression analysis identified 94 significantly up- and 188 downregulated genes (FC>1.0; p_adj_<0.05) in older versus younger samples (**Fig. S1A**). Gene set enrichment analysis (GSEA) and gene ontology (GO) analysis revealed that downregulated genes were enriched for cell cycle-related biological processes including nuclear chromosome segregation, nuclear division and mitotic cell cycle checkpoint signaling (**Fig. 1B**; **Tables S2**, **S3**), consistent with reduced proliferation capacity in fibroblasts from older donors^21,35–37^. Consistent with senescence-like phenotypes, MYC and EF2 target genes were negatively enriched, whereas TNFα, IL6-JAK-STAT3 and P53 signaling were positively enriched (**Fig. 1B**)^37–40^. Upregulated genes were enriched for lipid-associated processes including lipid localization, transport and storage (**Table S2**), as well as muscle- and endothelial-associated terms, suggesting partial loss of cell identity with age^21,41,42^. Together, these findings validate that low-passage SHOCK fibroblasts retain robust age-associated transcriptomic signatures.

**Figure 1.**
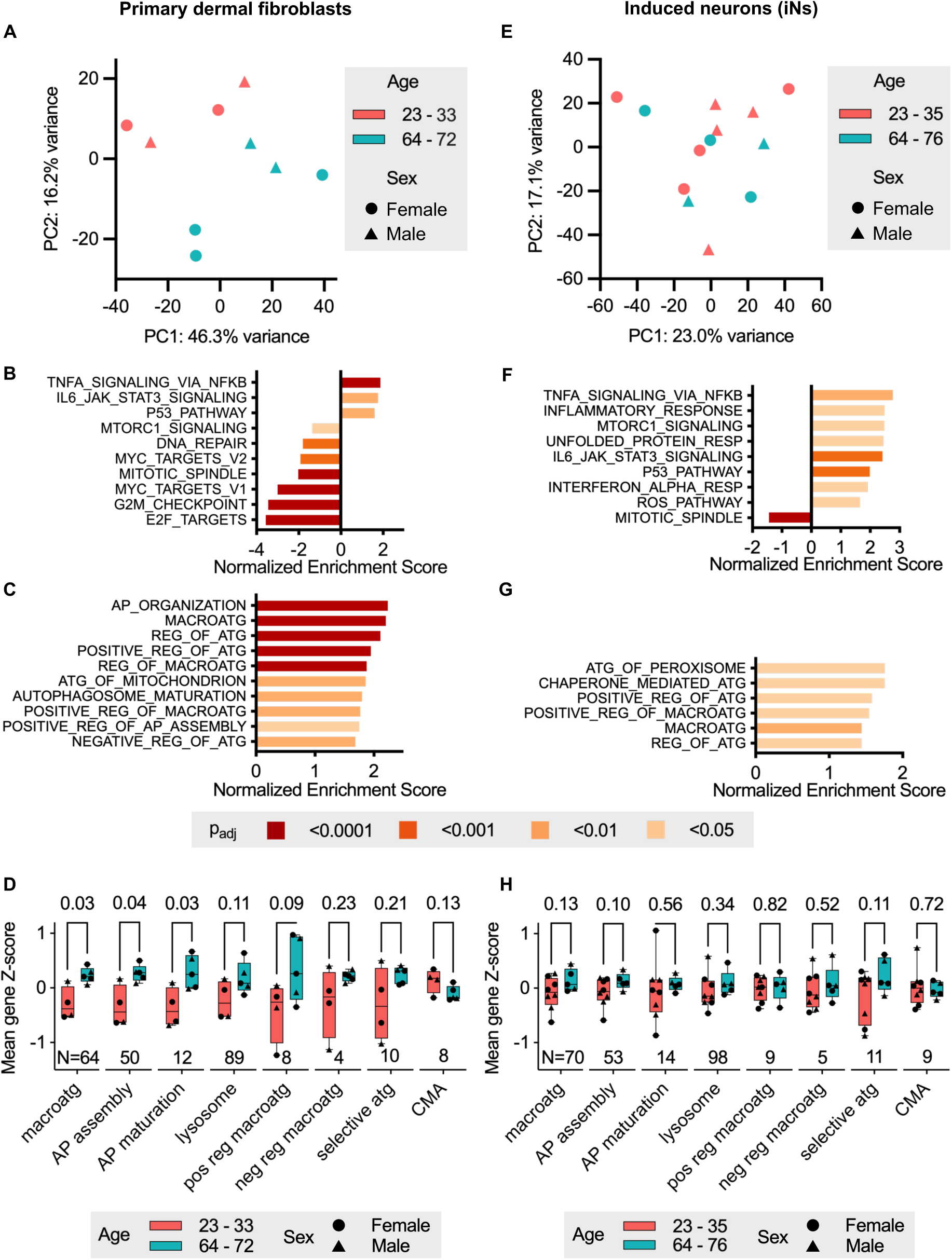
Aging is associated with upregulation of autophagy-related transcriptional programs in primary dermal fibroblasts and iNs. (**A**) PCA of primary dermal fibroblast samples labeled by age group (young: 23-33 years; older: 64-72 years) and sex. (**B-C**) Significantly enriched (**B**) age-related GSEA Hallmark and (**C**) autophagy-related GO Biological Process terms in primary dermal fibroblasts from older donors. (**D**) Mean gene Z-scores of autophagy-related GO Biological Process gene lists in primary dermal fibroblasts from young and older donors. P-values by independent t-tests. (**E**) PCA of iN samples labeled by age group (young: 23-35 years; older: 64-76 years) and sex. (**F-G**) Significantly enriched (**F**) age-related GSEA Hallmark and (**G**) autophagy-related GO Biological Process terms in iNs from older donors. (**H**) Mean gene Z-scores of autophagy-related GO Biological Process gene lists in iNs from young and older donors. P-values by independent t-tests. For **B-C** and **F-G**: bars are colored by adjusted p-value (p_adj_) as indicated in the key below panels **C** and **G**. All significantly enriched terms (p_adj_<0.05) are listed in **Tables S3** and **S6**. For **D** and **H**, gene modules are listed in **Table S6** and statistical comparisons are provided in **Table S5**. AP, autophagosome; macroatg, macroautophagy; reg, regulation; atg, autophagy; resp, response; ROS, reactive oxygen species; pos, positive; neg, negative; CMA, chaperone-mediated autophagy. See also **Figure S1**.

In primary dermal fibroblasts from older donors, autophagy-related GO Biological Process terms were positively enriched (**Fig. 1C**). To further resolve age-associated changes in autophagy gene transcription, we analyzed curated gene modules (**Tables S4**, **S5**). In fibroblasts, gene modules associated with macroautophagy, autophagosome components, and autophagosome assembly and maturation were significantly upregulated with age (**Fig. 1D**; **Table S4**). No clear sex-associated patterns were detected, although this may be due to limited sample size. In contrast, other autophagy-related modules, including selective autophagy receptors and chaperone-mediated autophagy (CMA), were not significantly altered with age (**Fig. 1D**). In addition, positive regulators of macroautophagy showed a trend toward upregulation (p = 0.09), whereas negative regulators were not significantly changed (p = 0.23), consistent with a regulated transcriptional response rather than stochastic age-associated dysregulation (**Fig. 1D**; **Table S4**). At the individual gene level, age-associated changes were heterogenous: while some macroautophagy genes (e.g., *ATG5*, *ATG7*, *ATG4C*) decreased with age, consistent with prior observations^43–46^, the majority of macroautophagy genes increased with age or peaked in mid-to-late adulthood (∼30-65 years; **Fig. S1B**). Together, these findings indicate that aging in human dermal fibroblasts is associated with coordinated but gene-specific remodeling of autophagy-gene transcription resulting in overall upregulation rather than a global decline.

We next sought to determine whether age-related changes in autophagy-gene transcription occur in a coordinated manner across human tissues, or have cell type-specific phenotypes. As long-lived, post-mitotic cells, neurons rely heavily on autophagy to maintain proteostasis^47,48^. However, age-associated changes in autophagy have not been systematically characterized in induced neurons (iNs) from healthy individuals. We therefore assessed autophagy-gene transcription in iNs as a model of human neuronal aging since they retain age-associated transcriptomic signatures^23^. We generated iNs via direct conversion^49^ of additional SHOCK donor fibroblasts, including five subject-matched lines, and performed bulk RNA-seq. Following conversion, iNs clustered separately from fibroblasts by PCA, confirming successful lineage conversion (**Fig. S1C**). Although iNs showed more modest global age-associated transcriptomic separation (**Fig. 1E**) and fewer differentially expressed genes (**Fig. S1D**) compared to fibroblasts, they exhibited positive enrichment of established age-associated pathways including inflammation, loss of proteostasis, oxidative stress and senescence (**Fig 1F**; **Table S6**)^50–54^. GSEA analysis further revealed enrichment of multiple autophagy-related pathways, including macroautophagy, and its positive regulation in aged iNs (**Fig. 1G**; **Table S6**). However, in contrast to fibroblasts, autophagy gene modules were not significantly changed with age in iNs (**Fig. 1H**), and even more heterogenous at the individual gene level (**Fig. S1E**). This divergence between pathway-level enrichment and module-level stability, suggests that age-associated changes in autophagy-gene transcription in iNs are variable across genes and individuals, rather than reflecting a uniform, coordinated response (**Fig. S1E**). Together, these data indicate that aging is associated with upregulation of autophagy-related transcriptional programs in both fibroblasts and iNs, but with stronger and more coordinated upregulation in fibroblasts and more variable responses in iNs.

### Age-related changes in autophagy flux are cell type- and sex-specific

Given the observed increase in autophagy-related gene transcription with age, we next asked whether these changes translated into increased autophagic activity. To address this, we measured autophagosome abundance, assessed as LC3B-positive structures (quantified as foci area per cell area), and autophagy flux using chloroquine (CQ)-mediated inhibition of lysosomal degradation in primary dermal fibroblasts (N = 44) and subject-matched iNs (N = 17) from SHOCK donors. We found that static measures of LC3B-positive foci did not reflect autophagic activity, as LC3B-positive foci area per cell area did not correlate with autophagy flux (**Fig. S2A**, **B**), demonstrating that autophagosome abundance alone is not a reliable measure of autophagic activity.

In primary dermal fibroblasts, autophagy flux exhibited sex-specific and age-dependent differences. Autophagy flux in fibroblasts from young male donors trended higher than those from young female donors (**Fig. 2A-C**), although sample size was insufficient to test sub-group differences in autophagy flux. With increasing age, autophagy flux declined in males but remained stable in females, resulting in comparable levels between sexes in older donors (**Fig. 2B**, **C**). The age-associated decrease in autophagy flux in male was independent of total fibroblast area (**Fig. S3A-D**). In a multiple linear regression model including an age-by-sex interaction, age was significantly associated with decreased autophagy flux in males (p = 0.021), whereas no significant age-associated change was observed in females. The age-by-sex interaction showed a trend toward significance (p = 0.075), supporting a sex-dependent effect of aging on autophagy flux in fibroblasts.

**Figure 2.**
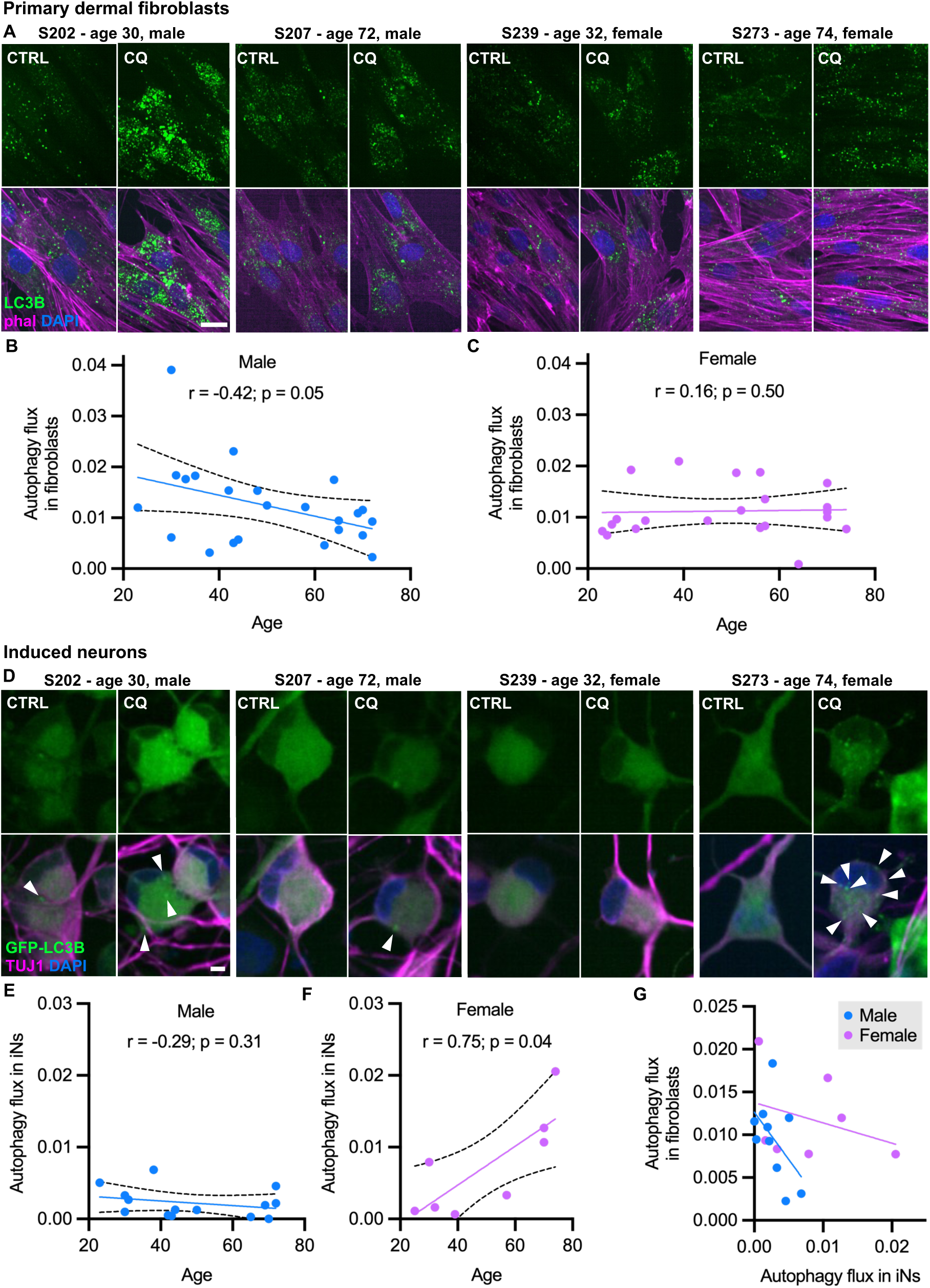
Age-related changes in autophagy flux are cell type- and sex-specific. (**A**) Representative images of LC3B-positive foci (green) in primary dermal fibroblasts from young and older male (left) and female (right) donors treated ± CQ for 2 hours. Actin cytoskeleton in magenta (phal – phalloidin) and nuclei in blue (DAPI). 40x magnification. Scale bar: 20 µm. (**B**) Autophagy flux in primary dermal fibroblasts from male SHOCK cohort donors spanning the adult lifespan, with each point representing one donor (N = 23 donors) (R^2^ = 0.17; p = 0.05); Spearman r and p-values reported. (**C**) Autophagy flux in primary dermal fibroblasts from female SHOCK cohort donors spanning the adult lifespan (N = 21 donors). Linear regression ± 95% confidence interval shown with each point representing one donor (R^2^ = 0.001; p = 0.87); Spearman r and p-values reported. (**D**) Representative images of GFP-LC3B-positive foci (green) in iNs from young and older male (left) and female (right) donors treated ± CQ for 2 hours. Neuronal marker TUJ1 in magenta and nuclei in blue (DAPI). 40x magnification. Scale bar: 50 µm. (**E**) Autophagy flux in iNs from male SHOCK cohort donors spanning the adult lifespan (N = 14 donors). Linear regression ± 95% confidence interval shown (R^2^ = 0.08; p_slope_ = 0.33); Spearman r and p-values reported. (**F**) Autophagy flux in iNs from female SHOCK cohort donors spanning the adult lifespan (N = 8 donors). Linear regression ± 95% confidence interval shown (R^2^ = 0.61; p = 0.02); Spearman r and p-values reported. (**G**) Autophagy flux in subject-matched fibroblasts and iNs labeled by donor sex. Linear regression for males (R^2^ = 0.26; p_slope_ = 0.14) and females (R^2^ = 0.11; p_slope_ = 0.47) shown. For **B-C** and **E-G**, autophagy flux reflects the absolute change in LC3B or GFP-LC3B-positive foci area per cell area ± CQ treatment. See also **Figures S2** and **S3**.

In iNs we found that static measures of LC3B-positive foci did not reflect autophagic activity, as in primary dermal fibroblasts, as GFP-LC3B-positive autophagosome area per cell area did not correlate with autophagy flux (**Fig. S2C**, **D**), demonstrating that flux-based assays are required to reliably measure autophagic activity in this cell type.

Autophagy flux in iNs also exhibited an age- and sex-dependent pattern, but distinct from fibroblasts. Autophagy flux was similar between male and female cells at younger ages, remaining relatively stable with advancing age in males but increasing with age in females (**Fig. 2D-F**). In female iNs, CQ treatment resulted in increased accumulation of GFP-LC3B-positive foci with increased age, consistent with increased autophagy flux (**Fig. S3E-H**). In a multiple linear regression model, sex was significantly associated with autophagy flux (p = 0.016), indicating a statistically significant difference in autophagy flux between male and female samples. Inclusion of an age-by-sex interaction term revealed a significant interaction (p = 0.001), indicating that the relationship between age and autophagy flux differs by sex, with flux increasing with age in females but not in males. Together, these findings indicate that age-related changes in autophagy flux are both cell type- and sex-dependent, with opposite trends observed in fibroblasts and iNs.

We next asked whether autophagy flux was coordinated across cell types within individuals. Autophagy flux in fibroblasts was not associated with autophagy flux in iNs across the 10 subject-matched male samples (r = −0.41; p = 0.25) or 7 subject-matched female samples (r = −0.46; p = 0.30) by Spearman’s analysis (**Fig. 2G**). Multiple linear regression analysis revealed a significant sex-dependent effect on the relationship between fibroblast and iN autophagy flux (p = 0.02), with a stronger negative association observed in males and a weaker relationship in females. This indicates that the relationship between autophagy flux across cell types differs by sex and is not uniformly coordinated across cell types within individuals.

Together, these findings demonstrate that autophagy flux is regulated in a cell type- and sex-dependent manner during human aging and does not mirror transcriptional changes in autophagy-related genes. This uncoupling highlights the importance of directly measuring autophagy flux to accurately assess autophagic activity.

### Autophagy flux becomes more heterogeneous with age in freshly isolated human PBMCs

Given the tissue- and sex-specific regulation of autophagy flux in fibroblasts and iNs, we next assessed age-associated changes in autophagy flux in freshly isolated PBMCs from SHOCK cohort participants (**Fig. 3A**). As circulating cells directly exposed to plasma, PBMCs provide a readout of systemic influences on autophagic activity^55–57^. Whole-blood and isolated-cell Coulter counts showed that our isolation method strongly enriched for lymphocytes (**Fig. S4A**), indicating that measured autophagy flux primarily reflected activity in T-, B- and NK cells. Technical variability across CQ treatment conditions was low for lymphocytes (CV < 20%) and increased only in scarce cell types (**Fig. S4B**), indicating consistent PBMC composition across samples.

**Figure 3.**
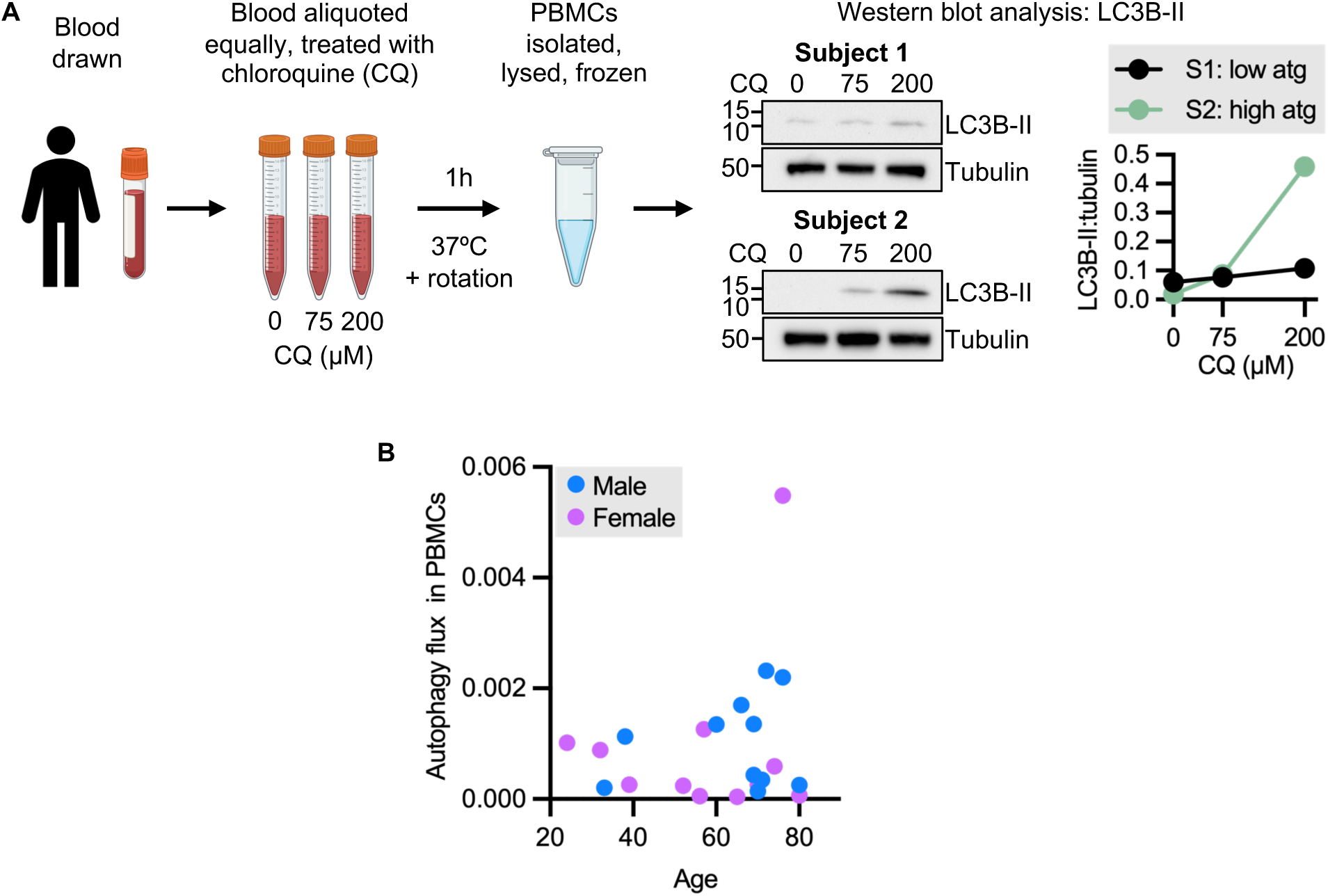
Autophagy flux becomes more heterogeneous with age in freshly isolated human PBMCs. (**A**) Experimental schematic of the autophagy flux assay in freshly drawn whole blood, followed by PBMC isolation and Western blot analysis of autophagosome marker LC3B-II. S1, subject 1; S2, subject 2; atg – autophagy. (**B**) Autophagy flux in PBMCs from male (N = 11) and female (N = 12) SHOCK cohort donors spanning the adult lifespan, with each point representing one donor. See also **Figures S4-S6**.

As in fibroblasts and iNs, basal LC3B-II levels in untreated samples did not correlate with autophagic flux (**Fig. 3A**; **S5A**), whereas LC3B-II levels following CQ treatment correlated strongly with measured flux (**Fig. 3A**; **S5B**), confirming the requirement for flux-based assays in PBMCs.

Autophagy flux in PBMCs from 23 SHOCK cohort participants was not associated with age (p = 0.44); however, autophagy flux became more heterogeneous with increasing age and showed a trend toward higher flux in individuals age ∼70 and above (**Fig. 3B**). This pattern is consistent with previous reports of increased autophagy flux with age in freshly isolated PBMCs^27,28^. No sex-associated differences in PBMC autophagy flux were observed (multiple linear regression main effects p = 0.86; age:sex interaction p = 0.91), indicating that PBMC autophagy flux is not significantly influenced by sex in this cohort. We further confirmed that these findings were not explained by age-related differences in PBMC composition, either in whole blood or after isolation (**Fig. S6A**, **B**).

Importantly, these data further support a model where autophagic activity is not coordinately regulated across the three assayed cell types within the limits of this cohort, as age- and sex-associated changes differed by cell type.

### Increased autophagy flux in human cells correlates with reduced physical fitness in older adults

We next asked whether autophagy flux in primary dermal fibroblasts, iNs and PBMCs could be indicative of physical health and fitness. To assess organismal fitness, SHOCK cohort participants underwent standardized clinical and functional assessments, performed at rest and during maximal exertion during progressive treadmill exercise (VO_2max_)^33^. Measures of exercise capacity, metabolic function and physical performance declined with age in the SHOCK healthy aging cohort, including resting VO_2_, (**Fig. S7A**), maximum heart rate (**Fig. S7B**), VO_2max_ (**Fig. S7C**), minute ventilation (**Fig. S7D**), grip strength (**Fig. S7E**), time spent in the very vigorous activity zone as measured by actigraphy (age-associated decline strongly driven by extremely high values in two young participants) (**Fig. S7F**), and walking speed (**Fig. S7G**), consistent with well-established age-related reductions in aerobic capacity, muscle function and high-intensity physical activity^58–61^. In contrast, measures of cardiovascular load and decreased physical function increased with age, including systolic and diastolic blood pressure (**Fig. S7H**, **I**), and chair stand time (**Fig. S7J**), reflecting known age-associated increases in vascular stiffness and cardiovascular risk^62,63^, and decreased mobility^64,65^. Notably, a subset of measures showed little or no association with age, including body-mass-index (BMI) (**Fig. S7K**), resting heart rate (**Fig. S7L**), Short Physical Performance Battery (SPPB) score (a composite measure of lower extremity function including balance, gait speed, and chair stand tests) (**Fig. S7M**), and accelerometer-derived sedentary time (**Fig. S7N**), indicating preservation of baseline functional status in this cohort.

We next examined how these physiological measures relate to autophagy flux across cell types. Across SHOCK cohort participants autophagy flux showed distinct associations with physiological outcomes that differed by cell type. In fibroblasts and PBMCs, higher autophagy flux correlated with measures indicative of more youthful cardiovascular function, including higher maximum heart rate (fibroblasts: Spearman r = 0.27; p = 0.075)^66^ and lower resting heart rate (PBMCs: Spearman r = −0.61; p = 0.002)^67,68^ (**Fig. 4A**). In contrast, in iNs higher autophagy flux correlated with reduced maximal grip strength (Spearman r = −0.51; p = 0.016) and walking speed (Spearman r = −0.52; p = 0.01) (**Fig. 4A**), suggesting that increased autophagy flux in iNs may be associated with reduced physical function^59^. Together, these findings indicate that autophagy flux is not simply a readout of chronological aging, but instead reflects aspects of physiological health that vary across individual and cell type.

**Figure 4.**
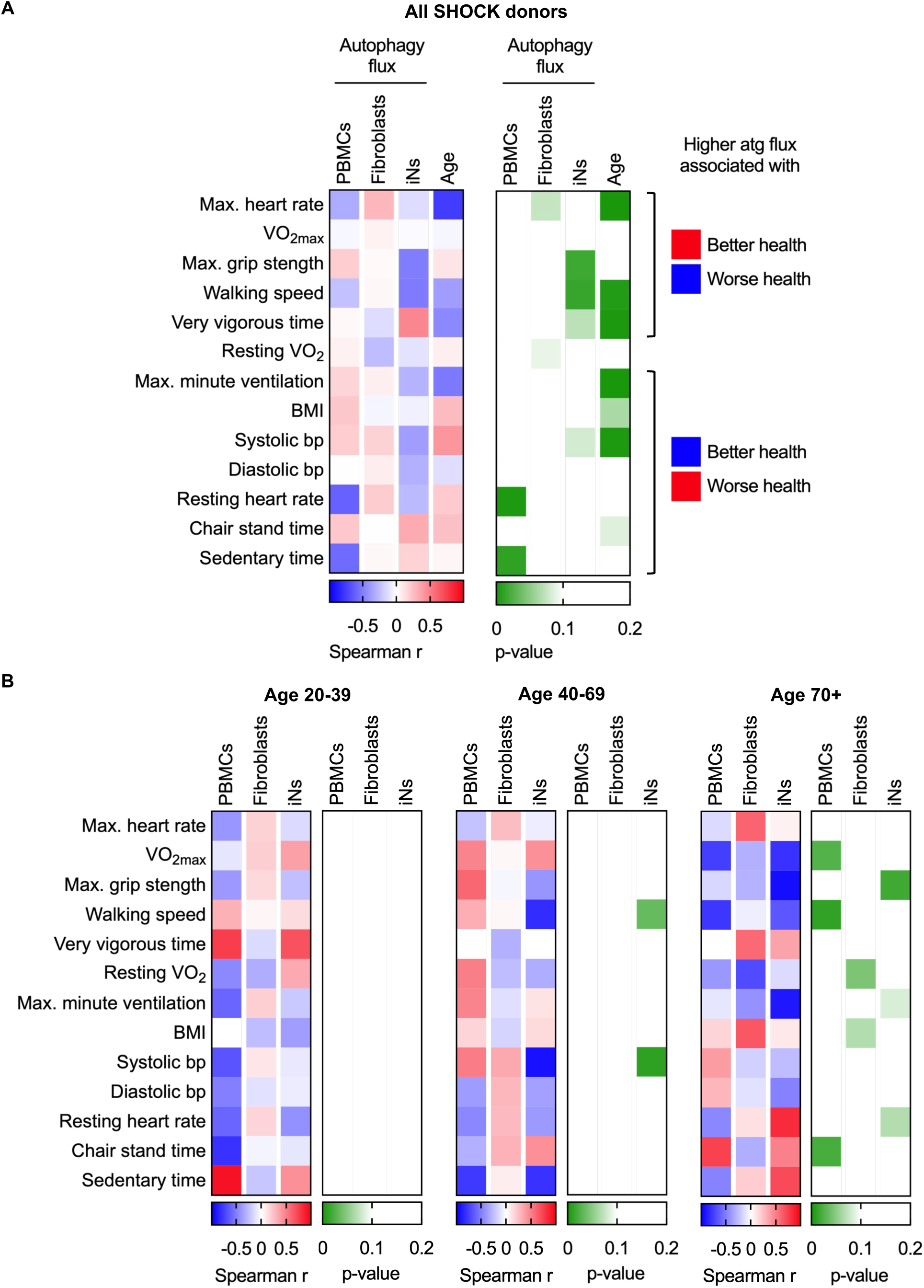
Increased autophagy flux in human cells correlates with reduced physical fitness in older adults. Heatmaps showing Spearman correlation coefficients (r) and corresponding p-values between autophagy flux in PBMCs, primary dermal fibroblasts or iNs, as well as age and physiological health readouts. Data are shown in (**A**) all SHOCK cohort participants, and (**B**) SHOCK cohort participants stratified by age group. In (**A**), the annotation on the right indicates whether higher values of each physiological measure are associated with better or worse health. Accordingly, positive correlations between autophagy flux and measures associated with better health indicate improved fitness, whereas positive correlations with measures associated with worse health indicate reduced fitness, and vice versa for negative correlations. Spearman correlations were calculated separately for each cell type and visualized as heatmaps. For all SHOCK cohort participants shown in (**A**), PBMCs, N = 23; fibroblasts, N = 44; iNs, N = 22; Age, N = 54. For SHOCK cohort participants in (**B**), Age 20-39: PBMCs, N = 5; fibroblasts, N = 15; iNs, N = 9; age 40-69: PBMCs, N = 8; fibroblasts, N = 20; iNs, N = 7; age 70+: PBMCs, N = 10; fibroblasts, N = 9; iNs, N = 6. See also **Figure S7**.

Since autophagy flux became more heterogeneous in PBMCs in adults age approximately 70 and older, and maximum autophagy flux in these subjects was relatively higher than in younger subjects (**Fig. 3B**), we next asked whether these relationships differed at advanced ages. Indeed, in SHOCK participants aged 70 years and older, higher autophagy flux across cell types was more consistently associated with reduced physical fitness. Specifically, higher PBMC autophagy flux was associated with lower VO_2max_ (Spearman r = −0.75; p = 0.026), slower walking speed (Spearman r = −0.78; p = 0.01) and longer time to complete the chair stand test (Spearman r = 0.73; p = 0.02); higher fibroblast autophagy flux correlated with increased BMI (Spearman r = 0.65; p = 0.067) and reduced resting VO_2_ (Spearman r = −0.70; p = 0.043); and higher iN autophagy flux was associated with reduced maximal grip strength (Spearman r = −0.94; p = 0.017), and increased resting heart rate (Spearman r = 0.82; p = 0.067) (**Fig. 4B**), although these relationships should be interpreted cautiously given the small sample size for these outcomes. With the exception of resting VO_2_, which is not indicative of health status on its own^69,70^, these measures are broadly consistent with reduced physical fitness in individuals with higher autophagy flux.

Together, these results indicate that the relationship between autophagy flux and physiological health differs with age. While higher autophagy flux shows variable associations with fitness across the adult lifespan depending on cell type, in older adults higher autophagy flux is consistently associated with reduced physical fitness.

### Mild exercise reverses elevated autophagy flux in fresh PBMCs

To test whether PBMC autophagy flux is modifiable in older adults with reduced physical function, we measured autophagy flux in participants from the Strong Foundations 2.5 (STRONG) cohort, a digitally delivered fall-prevention exercise program for older adults^71^ (**Tables 2**, **S7**). STRONG participants completed a 12-week mild exercise program consisting of weekly 1-hour sessions focused on improving balance, muscular strength and posture. Health and fitness measures were collected before and after the 12-week intervention program (**Fig. S8**), and included measures that overlapped with those assessed in the SHOCK healthy aging cohort, such as BMI, SPPB, maximal grip strength, resting heart rate, and resting systolic and diastolic blood pressure (**Fig. S9**). Maximal exertion testing was not performed in STRONG participants. This pilot study included STRONG participants aged 77-88 (**Fig. S9A**; **Table S7**). SPPB scores in the STRONG cohort were significantly lower than in the SHOCK cohort, indicating significantly lower functional capacity in STRONG participants (**Fig. S9C**; **Table S8**).

Baseline PBMC autophagy flux measurement in STRONG cohort participants revealed elevated flux relative to the SHOCK cohort mean, with 4 of 5 STRONG participants exceeding one standard deviation above the SHOCK cohort average across all ages (**Fig. 5A**). These findings strengthen the observation that higher PBMC autophagy flux in older adults is associated with reduced physical function.

**Figure 5.**
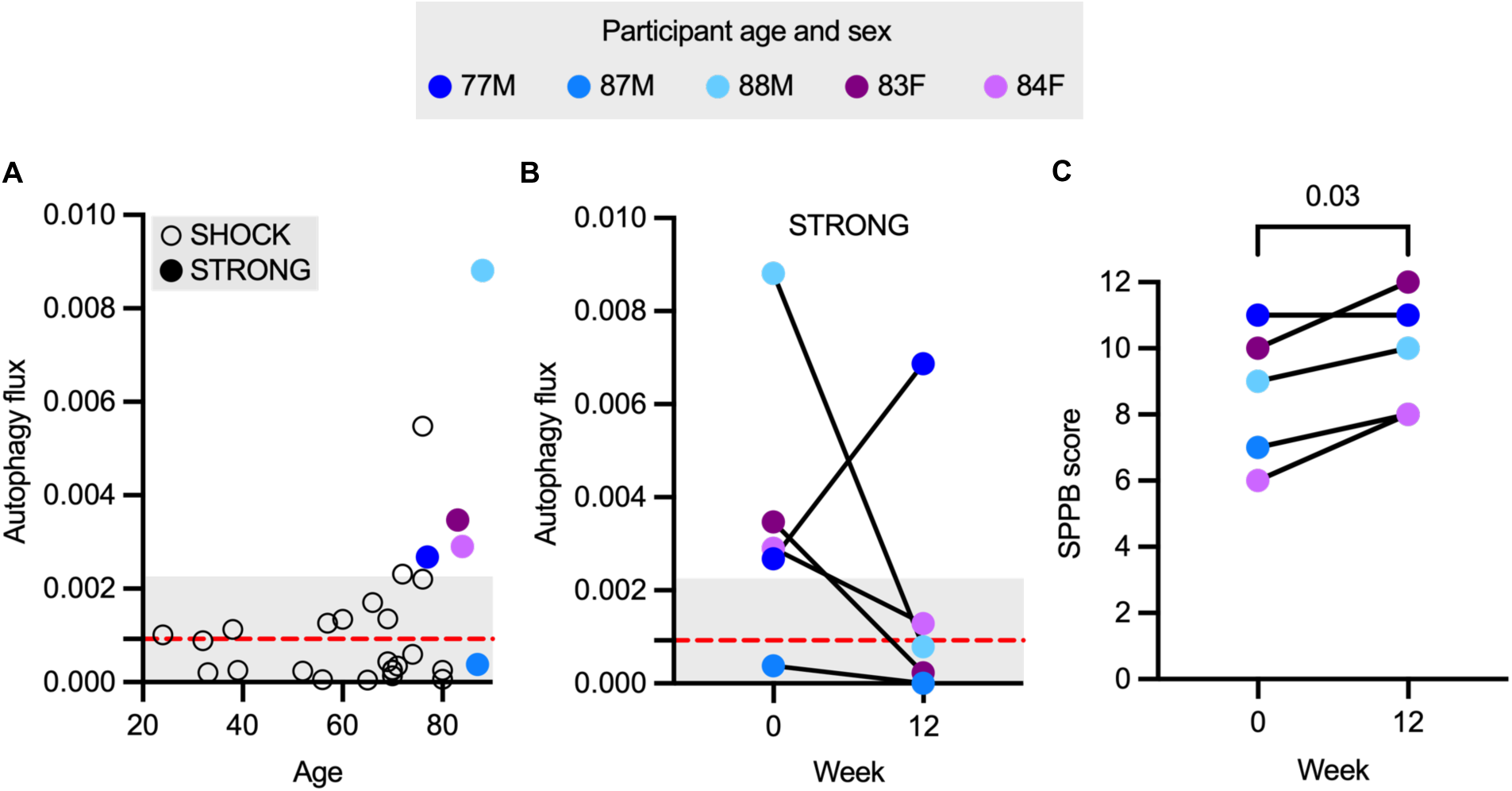
Mild exercise is associated with reduced PBMC autophagy flux and improved physical performance in older adults. Data points for STRONG cohort participant are colored by age (years) and sex (M – male; F – female). (**A**) Association between age and autophagy flux in PBMCs from SHOCK cohort participants (open circles; N = 23) and STRONG cohort participants at baseline (colored circles; N = 5). (**B**) Autophagy flux in STRONG cohort participants before and after a 12-week mild exercise program (N = 5). Each line represents one participant. P-value by paired t-test. Red dotted line and grey shading in (**A**) and (**B**) indicate mean SHOCK cohort autophagy flux ± standard deviation. (**C**) SPPB score in STRONG cohort participants before and after a 12-week mild exercise program. P-value by paired t-test. See also **Figures S8** and **S9**.

Following the 12-week mild exercise program, PBMC autophagy flux decreased in 4 out of 5 participants and fell within one standard deviation of the SHOCK Healthy Aging cohort mean (**Fig. 5B**). In parallel, SPPB scores improved in STRONG subjects (**Fig. 5C**), while BMI (**Fig. S8A**) and heart rate (**Fig. S8B**) were mildly increased, and maximal grip strength (**Fig. S8C**), posture (**Fig. S8D**, **E**), blood pressure (**Fig. S8F**, **G**), and chair stands (**Fig. S8H**; **Table S7**, **S8**) were unchanged.

These pilot data suggest that mild exercise may reduce PBMC autophagic activity while improving physical performance in older adults, supporting the potential for autophagy flux in PBMCs to be used as a biomarker of changes in physical function.

Interestingly, the single participant in whom autophagy flux increased following the intervention was the youngest (age 77) and the most physically functional at baseline (SPPB score = 11), whereas all other participants were age 83 or older. Although limited by sample size, this observation is consistent with the possibility that the relationship between PBMC autophagy flux and physiological fitness state may differ at more advanced ages. Overall, these preliminary findings indicate that autophagy flux in circulating immune cells is modifiable in humans and support its potential as a biomarker of physiological state in older adults.

## DISCUSSION

In this study, we identified cell type- and sex-specific differences in how autophagy flux changes with age in humans across PBMCs and subject-matched cultured primary dermal fibroblasts and iNs, and show that these functional changes are uncoupled from autophagy-related gene transcription. We further show that autophagy flux is associated with measures of physical function and fitness, and may be modulated by mild exercise in older adults. These findings challenge the prevailing view that autophagy uniformly declines with age and instead support a model of context-dependent regulation across tissues, cell types and physiological states.

In primary dermal fibroblasts, we find that autophagy-related gene expression is broadly upregulated with age, with enrichment of pathways associated with macroautophagy and lysosomal function, while modules associated with selective autophagy receptors and chaperone-mediated autophagy were not significantly altered. This suggests macroautophagy-specific transcriptional remodeling with heterogeneous changes on the individual gene level^45,46^. This transcriptional upregulation may reflect a compensatory response to age-associated cellular stress and accumulation of autophagic substrates, potentially driven by activation of stress-responsive regulators of autophagy such as transcription factor EB (TFEB) or forkhead box O (FOXO)^72,73^. Despite this overall transcriptional upregulation, these changes did not translate into uniformly increased functional autophagic activity in human dermal fibroblasts. Instead, our data indicate that age-associated changes in autophagy flux in fibroblasts are sex-dependent, with a decline observed in males and relative stability in females. Whether this response is protective or reflects insufficient maintenance of autophagic capacity remains unclear and will require direct assessment of autophagosome quality and lysosomal function. Prior studies assessing autophagy flux in human dermal fibroblasts have yielded conflicting results^74,75^, likely due to differences in cohort composition and experimental design. Our findings extend this literature by demonstrating that age-associated changes in fibroblast autophagy flux are sex-dependent, underscoring the importance of considering biological sex when evaluating autophagic function in human aging.

In iNs, autophagy flux exhibited distinct age- and sex-dependent patterns that differed from those observed in fibroblasts, including an increase in flux with age in females and relative stability in males. These findings indicate that neuronal autophagy is regulated in a cell type- and sex-specific manner and, when compared with subject-matched fibroblasts, further suggest that autophagy is not coordinately regulated across tissues within individuals. At the transcriptional level, aging in iNs was associated with enrichment of autophagy-related pathways, including macroautophagy and its positive regulation; however, in contrast to fibroblasts, autophagy gene modules were not significantly altered with age, indicating a less coordinated transcriptional response. Future studies comparing autophagy gene transcription and flux in subject-matched directly converted and iPSC-derived neurons will additionally help clarify how conversion strategy influences autophagy regulation and strengthen the generalizability of these findings across neuronal models. Moreover, given that autophagy is altered at the proteomic and functional levels in iNs derived from patients with neurodegenerative diseases^76–78^, it will be important to map these changes longitudinally to better understand how autophagy evolves during the transition from healthy aging to disease states.

Taken together, these observations reveal a disconnect between transcriptional and functional measures of autophagy, highlighting that similar transcriptional upregulation across cell types can be associated with divergent functional outcomes. This underscores the importance of directly measuring autophagy flux rather than relying on transcriptional proxies, and points to post-transcriptional and post-translational mechanisms as regulators of autophagic activity during human aging, a possibility that warrants direct investigation in future work.

Recent studies have begun to measure autophagy flux in freshly isolated PBMCs, demonstrating that autophagy flux is heterogeneous and can increase with age^27,28^. Consistent with these observations, our data show that autophagy flux in PBMCs becomes increasingly heterogeneous with age and trends higher in older individuals. In line with prior reports showing that increased PBMC autophagy flux is associated with adverse cardiometabolic parameters, i.e. increased systolic blood pressure in pre-diabetic individuals^27^, we find that elevated autophagy flux in older adults is associated with reduced physical function and fitness. While dietary interventions including caloric restriction, intermittent fasting and acute protein intake^30,31^ do not significantly modulate of autophagy flux in freshly isolated PBMCs, our pilot intervention of mild exercise over a 12-week period induced measurable changes in autophagy flux in older adults, reducing autophagy flux toward levels observed in younger individuals. These data support a role for exercise as a distinct modulator of autophagic activity in circulating immune cells, unlikely to be confounded by dietary variability. Notably, the reduction of elevated PBMC autophagy flux after exercise suggests that high autophagy flux in older adults may represent a compensatory response to increased cellular stress rather than enhanced autophagy capacity. One potential explanation is that exercise reduces systemic inflammatory signals, including in older adults^79–81^, thereby lowering the demand for compensatory autophagy. Determining whether autophagosome quality and lysosomal function are preserved in aged PBMCs and iNs will be important to distinguish adaptive from dysfunctional autophagy.

Age-associated changes in autophagy were not coordinated across tissues but instead reflected tissue-specific regulation. Autophagy flux in PBMCs did not correlate with flux in fibroblasts or iNs, and fibroblast and neuronal autophagy showed sex-specific opposing patterns, suggesting that autophagy is shaped by cell-intrinsic properties and tissue-specific environmental context rather than a unified systemic program. The relationship between autophagy and health appears to shift with age: while associations between autophagy flux and fitness are variable across the adult lifespan, elevated autophagy in older individuals is more consistently associated with reduced physical performance, raising the possibility that autophagy becomes increasingly stress-responsive or dysregulated in late life.

These findings have important implications for autophagy-targeting interventions. Efforts to globally enhance autophagy may not be uniformly beneficial and could exacerbate dysfunction if autophagic processes are already impaired or inefficient. Instead, a more nuanced understanding of tissue-specific autophagy regulation and functional capacity will be required. In this context, PBMC autophagy flux represents a promising, accessible readout of physiological state that is sensitive to intervention and may serve as an early biomarker of changes in organismal function, but importantly, likely does not reflect autophagy flux in skin fibroblasts and neurons. Future studies will be required to define the molecular mechanisms linking systemic signals to autophagy regulation, assess autophagosome quality and lysosomal function across tissues, and extend these analyses to additional human cell types.

### Limitations of the study

This study has several limitations. First, the sample size for both the SHOCK and STRONG cohorts was modest, limiting statistical power to detect subtle age- or intervention-associated effects. In the STRONG cohort, limited sampling timepoints further restrict the ability to resolve the temporal dynamics of autophagy flux during mild exercise. Second, participant ethnic diversity was limited in both cohorts, which may affect the generalizability of these findings across populations. Third, analyses of fresh tissue were restricted to PBMCs, which may not capture tissue-specific autophagy regulation in organs such as brain or skin. Although primary dermal fibroblasts and directly reprogrammed neurons retain aging signatures, they remain *in vitro* models and may not fully reflect the *in vivo* cellular environment, including exposure to circulating factors and tissue-specific architecture. Finally, autophagic activity was assessed using LC3B-based flux assays, which do not assess cargo selectivity, autophagosome structure or lysosomal function, limiting insight into the qualitative nature of age-related changes in autophagy. These limitations highlight the need for larger, longitudinal studies and multimodal assessments of autophagy to further define its role in human aging and disease.

## RESOURCE AVAILABILITY

### Lead contact

Requests for further information and resources should be directed to and will be fulfilled by the lead contact, Caroline Kumsta.

### Materials availability

All unique/stable reagents generated in this study are available from the lead contact with a completed materials transfer agreement.

### Data and code availability

- All data reported in this paper will be shared by the lead contact upon request.
- All original code used in this paper will be shared by the lead contact upon request.
- Any additional information required to reanalyze the data reported in this paper is available from the lead contact upon request.

## EXPERIMENTAL MODEL AND STUDY PARTICIPANT DETAILS

### SHOCK cohort for primary dermal fibroblast, induced neuron and peripheral blood mononuclear cell studies

The San Diego Nathan Shock Center Clinical Cohort (SHOCK) has previously been described^33^, including standardized assessments of physical and cognitive performance. All participants provided informed consent, and all procedures were approved by University of California, San Diego (UC San Diego) IRB 201141. For the present study, a subset of SHOCK cohort participants was selected based on sample availability for analysis of autophagy-gene transcription and autophagy flux in primary dermal fibroblasts and iNs, and autophagy flux in freshly isolated peripheral blood mononuclear cells (PBMCs) (**Table 1**). Physical health measures for all SHOCK participants were collected, as previously described^33^. Autophagy flux measurements in PBMCs were correlated to participant age, sex and physiological fitness measures obtained under resting conditions and following exercise to volitional exhaustion. Additional experimental, demographic and health data for each participant are available in **Table S1**.

**Table 1.** Summary of San Diego Nathan Shock Center Healthy Aging (“SHOCK”) cohort demographics by experiment.

| Assay | RNA-seq |  | Autophagy flux |  |  |
| --- | --- | --- | --- | --- | --- |
| Cell type | Fibroblast<br>[mean<br>(range)] | iN<br>[mean<br>(range)] | Fibroblast<br>[mean<br>(range)] | iN<br>[mean<br>(range)] | PBMC<br>[mean<br>(range)] |
| **N** | 9 | 13 | 44 | 17 | 23 |
| **Age**<br>**(years)** | 49.3<br>(23 – 72) | 42.6<br>(23 – 76) | 49.5<br>(23 – 74) | 52.5<br>(23 – 74) | 60.8<br>(24 – 80) |
| **BMI**<br>**(kg/m<sup>2</sup>)** | 23.30<br>(20.3 – 29.7) | 24.63<br>(20.3 – 30.0) | 24.64<br>(20.0 – 30.1) | 24.51<br>(20.3 – 30.0) | 24.51<br>(19.5 – 30.1) |
| **Sex**<br>**(% male)** | 44.4 | 46.2 | 52.3 | 58.8 | 47.8 |
| **Race**<br>**(% White)** | 88.9 | 92.3 | 86.4 | 82.4 | 95.7 |
| **SPPB**<br>**score** | 12.0<br>(12 – 12) | 11.8<br>(10 – 12) | 12.0<br>(11 – 12) | 11.9<br>(11 – 12) | 11.8<br>(8 – 12) |

BMI, body mass index; PBMC, peripheral blood mononuclear cell; iN, induced neuron; SPPB, Short Physical Performance Battery, a composite measure of lower extremity function including balance, gait speed, and chair stand tests (score range 0-12, with higher scores indicating better physical function).

**Table 2.** Summary of STRONG participant demographics, and fitness measures before and after a 12-week mild exercise intervention.

|  | Baseline, week 0<br>[mean (range)] | 12-weeks mild<br>exercise<br>[mean (range)] |
| --- | --- | --- |
| **N** | 5 |  |
| **Age (years)** | 83.8 (77 – 88) |  |
| **Sex (% male)** | 60.0 |  |
| **Race (% White)** | 100.0 |  |
| **BMI (kg/m<sup>2</sup>)** | 27.5<br>(23.9 – 31.6) | 29.0<br>(24.6 – 36.7) |
| **SPPB score** | 8.6 (6 – 11) | 9.8 (8 – 12) |

BMI, body mass index; PBMC, peripheral blood mononuclear cell; SPPB, Short Physical Performance Battery, a composite measure of lower extremity function including balance, gait speed, and chair stand tests (score range 0-12, with higher scores indicating better physical function). All measurements were obtained from the same participants at baseline (week 0) and after 12 weeks of exercise.

### STRONG cohort for PBMC studies before and after 12 weeks of mild exercise

The Strong Foundations 2.5 (STRONG) cohort sponsored by UCSD’s Exercise and Physical Activity Resource Center underwent similar intervention to a Strong Foundations cohort that has previously been described^71^, including standardized assessments of physical and cognitive performance. All participants provided informed consent, and all procedures were approved by UC San Diego’s IRB 809304. Participants were residents of senior living facilities in San Diego, CA, identified as low- or moderate-risk for falls, who volunteered to participate in a modified, partially digitally delivered exercise program designed to improve strength and prevent falls. For the present study, a subset of STRONG cohort participants was selected based on sample availability and data completeness for blood draws to assess autophagy flux in PBMCs before and after 12 weeks of participation in the exercise program. Although more limited in scope than within the foundational study, the methodology behind the physical health and performance metrics for all STRONG participants collected at baseline (week 0) and post-intervention (week 12) timepoints have been previously described^71^ (**Table S7**). Changes in autophagy flux in PBMCs and corresponding changes in physical fitness between readouts from week 0 and week 12 were analyzed (**Table S8**).

## METHOD DETAILS

### Cell culture

Primary dermal fibroblasts from donors aged 23-76 years were obtained from the San Diego Nathan Shock Center (SD-NSC). Cells were cultured at 37°C, 5% CO_2_ and ambient O_2_ (∼21%) in DMEM (Gibco #11965092) supplemented with 15% fetal bovine serum (FBS, Corning), 1% non-essential amino acids (NEAA, Gibco #11140050), and 1% GlutaMAX (Gibco #35050061), referred to as ‘fibroblast media’. All primary dermal fibroblast lines were at or below passage 12 for all experiments (**Table S1**).

### Lentivirus production and titration

Production of UNA lentivirus for fibroblast to iN conversion was performed as previously described^49^. Briefly, HEK293T cells were transfected with pLVXUbC-rtTA-Ngn2:2A:Ascl1 (Addgene: #127289), and the packaging plasmids psPAX2 (Addgene #12260) and pMD2.G (Addgene #12259) using PEI. Viral titers were quantified using Lenti-X GoStix Plus (Takara) and concentrated by ultracentrifugation after 72 hours of incubation. Concentrated virus was resuspended in DPBS and stored at −80°C.

For production of EGFP-LC3B lentivirus, large-scale lentivirus production and concentration were performed by the Sanford Burnham Prebys Functional Genomics core facility. The lentivirus production procedure and conditions were optimized using Biomek i7 automation at high-throughput scale and subsequently scaled up to 150 cm² culture dishes for larger-scale production^82^. Briefly, pCDH-EF1a-mCherry-EGFP-LC3B lentivector DNA (Addgene #170446) was co-transfected with psPAX2 (Addgene #12260) and pMD2.G (Addgene #12259) plasmids into HEK293T cells in nine 150 cm^2^ dishes. Viral supernatant was collected every 24 hours from day 2 to day 4 post-transfection. Crude virus was filtered through a 0.22 µm filter and concentrated by ultracentrifugation at 21,000 rpm for 2 hours at 4°C using a 20% sucrose gradient. Concentrated viral particles were resuspended in DPBS, aliquoted and stored at - 80°C. Viral titers were determined by measuring transduction efficiency in target cells using a LSRFortessa (BD) flow cytometer at 4 days post-transduction.

### Generation of directly-reprogrammed neurons from primary dermal fibroblasts

Induced neurons (iNs) were generated from SHOCK cohort primary dermal fibroblasts as previously described^49^. Briefly, primary dermal fibroblasts were transduced with lentivirus pLVXUbC-rtTA-Ngn2:2A:Ascl1 (Addgene #127289) carrying puromycin resistance and doxycycline-inducible neuronal transcription factors NGN2 and ASCL1 (UNA cassette). Transduced cells were expanded under puromycin selection (1 µg/mL) in TFM-P media^49^ for at least five passages. Primary dermal fibroblasts carrying the UNA cassette are referred to as ‘UNA+ fibroblasts’. UNA+ fibroblasts were generated either in-house or obtained from the SD-NSC. All UNA+ fibroblast lines were maintained below passage 18.12, meaning 18 passages from fibroblast isolation and 12 passages after transfection, at the time of conversion (**Table S1**). For iN conversion, UNA+ fibroblasts were pooled to 300% confluency in TFM-P, and after 24 hours, changed to iN conversion media (referred to as ‘NK media’), containing doxycycline and a defined combination of neuronal conversion factors^49^. NK media was replaced every 2 days for the first week, followed by three times per week thereafter. iNs were harvested on day 21 post-induction for RNA-seq analysis, or purified on day 23 post-induction for autophagy flux experiments.

### RNA-seq analysis of primary dermal fibroblasts and iNs

Primary human dermal fibroblasts from 4 young (23-33 years) and 5 older (64-72 years) SHOCK cohort donors were analyzed (see **Tables 1**, **S1**). Cells were cultured in fibroblast media. For RNA-seq, 200,000 cells per culture were seeded in single wells of 6-well plates and grown under the previously described conditions for 48 hours. RNA was extracted following the RNeasy RNA purification kit protocol (Qiagen #74104). Cells were washed with Dulbecco’s phosphate-buffered saline (DPBS, Gibco # 14190094) and lysed in RNeasy lysis buffer containing β-mercaptoethanol, followed by homogenization using QIAshredder columns (Qiagen #79656), and purification with RNeasy Mini spin columns, including on-column DNase digestion. PolyA library prep was performed by the Salk Institute for Biological Studies Next Generation Sequencing and Genomics Core using the Illumina NovaSeq 6000 SP to generate paired-end reads (50 bp) at an average depth of ∼50 million reads per sample.

iNs directly converted from UNA+ fibroblasts from 8 young (23-35 years) and 5 older (64-76 years) SHOCK cohort donors were analyzed. After three weeks of conversion, iNs were detached using TrypLE and purified by magnetic-associated cell sorting (MACS) using PSA-NCAM antibody-associated beads (Miltenyi cat# 130-092-966) over MACS columns (Miltenyi cat# 130-042-201). 200,000 – 300,000 purified iNs per sample were pelleted and snap-frozen. RNA was extracted using the TRIzol LS protocol, and genomic DNA was removed using TURBO DNase (Thermo Fisher Scientific). Directional mRNA libraries were generated using poly(A) enrichment and sequenced on the NovaSeq X Plus platform (Illumina), generating paired-end 150 bp reads. Library preparation and sequencing were performed by Novogene USA.

Paired-end libraries from primary dermal fibroblasts and induced neurons (iNs) were processed independently with the nf-core/rnaseq pipeline executed on Amazon Web Services EC2 under Nextflow v25.10.4, using Docker containerization for software reproducibility^83^. Fibroblast libraries were processed with nf-core/rnaseq v3.23.0 against the iGenomes NCBI build of the human reference genome GRCh38, including the pre-built STAR index and matching NCBI GTF annotation^84^. iN libraries were processed with nf-core/rnaseq v3.20.0 against the Ensembl GRCh38 primary assembly with the Ensembl release 113 GTF. For both datasets, reads were adapter- and quality-trimmed with Trim Galore (v0.6.10), aligned to the genome with STAR (v2.6.1d for fibroblasts and v2.7.11b for iNs), and transcript-level abundances were quantified from the genome alignments with Salmon (v1.10.3) in alignment-based mode. Gene-level count matrices were generated by the pipeline’s built-in tximeta/tximport aggregation step (salmon.merged.gene_counts.tsv). Per-sample sequencing and alignment quality were assessed with FastQC (v0.12.1), Picard MarkDuplicates, RSeQC, Qualimap, and dupRadar, and aggregated into MultiQC reports; all samples met quality thresholds (min_trimmed_reads = 10,000; min_mapped_reads = 5) prior to downstream statistical analysis.

All downstream analyses were performed in R. Gene-level counts were imported into DESeq2, and genes with fewer than 10 total counts across all samples were removed prior to analysis. A multi-factor differential expression model estimating age and sex effects simultaneously (design: ∼ sex + age) was fit. DESeq2 applied a negative binomial generalized linear model with empirical Bayes dispersion shrinkage and Wald test statistics. Log₂ fold changes were further regularized using adaptive shrinkage (apeglm). Genes were considered significantly differentially expressed at an adjusted p-value (Benjamini–Hochberg, BH) < 0.05 and |log₂ fold change| > 1. For principal component analyses, variance-stabilizing transformation (VST, blind = TRUE) was applied to the filtered count matrix and PCA was computed using prcomp(). Gene Ontology (GO) Biological Process enrichment was performed with enrichGO from clusterProfiler, using all tested genes as the background universe and converting HGNC gene symbols to Entrez IDs via org.Hs.eg.db; significance was defined at p < 0.05 and q < 0.05 (BH). Up- and downregulated gene sets were tested separately to preserve directional specificity. For gene set enrichment analysis (GSEA), all tested genes were pre-ranked and analyzed using fgsea() with the c1.Hallmark, c2.KEGG:Medicus and c5.GO:BP gene sets. For hypothesis-driven pathway analysis, curated gene sets were organized by PANTHER Biological Process and Cellular Component annotations, and selective autophagy receptor and CMA lists^18^. VST expression values (blind = FALSE, using the ∼ sex + age design) were computed from the multi-factor DESeq2 object and collapsed to the gene-symbol level by averaging across multiple Ensembl IDs mapping to the same symbol. Gene expression was z-scored row-wise across samples, and a module score was computed per sample as the mean z-score across all module genes. Module scores were compared between Young and Older groups using Welch’s two-sample t-test (var.equal = FALSE). P-values were corrected for multiple comparisons across all modules using the Benjamini–Hochberg method. Modules were considered nominally significant at p < 0.05; no modules reached FDR < 0.05, consistent with limited statistical power at N = 9.

### Autophagy flux assay in primary dermal fibroblasts

SHOCK primary dermal fibroblasts were seeded in fibroblast media in four 384-well plates over a period of 2 days. On day 1, 22 fibroblast cultures were seeded at a density of 150 cells per well, and on day 2, the remaining 22 fibroblast cultures were seeded at a density of 250 cells per well. A 70% total volume media change was performed on all wells on day 4. On day 6, cells were subjected to an autophagy flux assay. For the autophagy flux assay, 50% of fibroblast media was removed from each well and replaced with either plain fibroblast media or fibroblast media containing 100 µM chloroquine (CQ; Sigma #C6628) for a final concentration of 50 µM CQ per well. Cells were incubated for 2 hours at 37°C, 5% CO_2_. Media was then replaced with 4% paraformaldehyde (PFA) in DPBS for 20 minutes. Cells were then rinsed 3 times for 5 minutes with DPBS and processed for immunofluorescent staining by incubating in 1:100 anti-LC3B (MBL Life Science #M152-3) in 0.2% saponin (Acros Organics #419231000)/10% FBS in DPBS at 4°C for 1h30, followed by 1:1,000 Alexa Fluor 488-labeled secondary antibody (Invitrogen # A11001) and 1X Alexa Fluor 647-labeled phalloidin (Thermo Scientific #A22287) in 0.2% saponin/10% FBS in DPBS at room temperature for 1 hour, and finally 1X DAPI (Sigma #D9542) in DPBS for 15 minutes.

### Autophagy flux assay in iNs

On day 23 post-induction, iNs were separated by MACS as described above. Purified iNs were seeded in 12 Geltrex (Gibco #A1413201)-coated wells of a 384-well plate per cell line at 6,000 cells per well in NK media. 24 hours after seeding, iNs were transduced with pCDH-EF1a-mCherry-EGFP-LC3B (Addgene #170446) at MOI 20 in NK media containing 8 µg/mL polybrene. After 12 hours transduction, wells were cycled with fresh NK media 5 times to remove the lentivirus. 72 hours after lentivirus removal, an autophagy flux assay was performed as described above. Briefly, iNs were incubated in NK media at a final concentration of 0 or 50 µM CQ for 2 hours at 37°C, 5% CO_2_. After treatment, media was replaced with ice cold PFA at a final concentration of 4% in DPBS for 20 minutes. Cells were then rinsed 3 times for 5 minutes with DPBS. iNs were processed for immunofluorescent staining by incubating in 1:500 anti-TUJ1 (neuronal marker, BioLegend #801202) in 0.2% saponin/10% FBS in DPBS at 4°C overnight, followed by 1:1,000 Alexa Fluor 647-labeled secondary antibody (Invitrogen #A32728) in 0.2% saponin/10% FBS in DPBS at room temperature for 4 hours, and finally, 1X DAPI in DPBS for 15 minutes.

### Image Acquisition and quantification of LC3B-positive structures

Cells were imaged on an Opera Phenix High-Content Screening system in confocal mode with a 40X objective. In primary dermal fibroblasts, total cell and LC3B-positive foci area was quantified using a custom CellProfiler v4.2.8^85^ pipeline in which LC3B-positive foci were identified based on adaptive intensity, and masked by phalloidin staining to assign to the correct cell. Foci were quantified in at least 81 fields of view per condition. Autophagy flux was determined by subtracting the average total LC3B-positive foci area per cell area in CTRL-treated cells from that in CQ-treated cells. In iNs, total cell body and GFP-LC3B-positive foci area in TUJ1-positive iNs was quantified by manually outlining and measuring the area of regions and of interest using ImageJ2 v2.16.0/1.54p. Foci were quantified in at least 100 cells per condition. Autophagy flux was determined by subtracting the average total GFP-LC3B-positive foci area per cell body area in CTRL-treated cells from that in CQ-treated cells.

### Autophagy flux assay and isolation of fresh human PBMCs

Autophagy flux assays in PBMCs were performed similarly to a previously described method^86^. Blood was drawn by a licensed phlebotomist into a 9 mL VACUETTE lithium heparin collection tube (Greiner #455084). Samples were processed within 45 minutes for SHOCK cohort participants, and within 2.5 hours for STRONG cohort participants. After an initial Coulter count (Beckman Coulter DxH 520 hematology analyzer), blood was split evenly into three 9 mL VACUETTE lithium heparin tubes. Prostaglandin E (50 µL in DPBS; 18 µM stock) and CQ (final concentrations 0, 75 or 200 µM) were added to each sample. Samples were incubated at 37°C on a rotator (10 rpm) for 1 hour prior to PBMC isolation. For PBMC isolation, samples were centrifuged at 500 x g for 15 minutes to separate the plasma from hematocrit. The hematocrit containing PBMCs was diluted in 12 mL warm Roswell Park Memorial Institute Medium (RPMI, Gibco #11-835-030) containing 50 µL prostaglandin E, layered over 3 mL Histopaque 1077 (Sigma #10771), and centrifuged at 700 x g for 30 minutes. The buffy coat was collected, diluted in 20 mL RPMI + 50 µL prostaglandin E, and centrifuged at 700 x g for 15 minutes. The PBMC pellet was resuspended in 950 µL PRMI + 50 µL prostaglandin E for a final Coulter count, then pelleted at 500 x g for 5 minutes and lysed in ice-cold lysis buffer (10 mM Tris/HCl, pH 7.0, 1 mM EDTA, 0.5 mM EGTA, 1% Triton X-100, 0.1% sodium deoxycholate, 0.1% SDS, 140 mM NaCl, 2.5 mM sodium pyrophosphate, 1 mM sodium orthovanadate, 1 mM 2-glycerophosphate, 1X cOmplete protease inhibitor, 1X PhosSTOP phosphatase inhibitor in water). Lysates were incubated at 4°C on a rotator for 45-60 minutes, centrifuged at 16,000 x g at 4°C for 10 minutes, and the supernatant was aliquoted and flash-frozen in liquid N_2_, and stored at −80°C. At a later timepoint, total protein concentration was determined using Pierce 660 Protein Assay Reagent (Thermo #22660) with 1X Ionic Detergent Compatibility Reagent (Thermo #22663).

### Western blots

For Western blotting, 1.5 µg of protein was mixed with 6X Laemmli buffer (Thermo #J61337.AD) and brought to a final volume of 36 µL with lysis buffer. Protein was denatured at 95°C for 10 minutes, after which samples were loaded onto a 10% Bis-Tris SDS-PAGE gel (NuPAGE #NP0315) in MOPS running buffer (NuPAGE, #NP0001), and separated at 110 V. Proteins were transferred to a PVDF membrane (Bio-Rad Laboratories, #1704156) using the Trans-Blot Turbo system (Bio-Rad Laboratories) at 1.3 A for 7 minutes. The membrane was cut horizontally at ∼30 kDa. The bottom portion was probed with rabbit anti-LC3B (1:1,000 in 1% milk in TBS-T; Cell Signaling #3868) and secondary anti-rabbit IgG HRP (1:10,000 in 1% milk in TBS-T; Cell Signaling #7074), and the top portion was probed with mouse anti-tubulin (1:1,000 in 1% milk in TBS-T; Cell Signaling #3873) and secondary anti-mouse IgG HRP (1:10,000 in 1% milk in TBS-T; Cell Signaling #7076). Blots were developed using SuperSignal chemiluminescence reagent (Thermo Scientific, #34577 or #34095) and imaged on a ChemiDoc imager (Bio-Rad Laboratories, Inc.). Protein expression was quantified using ImageJ2 (v2.16.0/1.54p) by measuring mean grey values within equal-sized rectangular regions of interest. Background signal was calculated from three evenly spaced regions and subtracted from each measurement. LC3B-II signal was normalized to tubulin, and autophagy flux was calculated as the difference between tubulin-normalized LC3B-II signal in CQ-treated and CTRL-treated samples.

## QUANTIFICATION AND STATISTICAL ANALYSIS

Statistics for all RNA-seq analyses were calculated using R as described above. All other statistics including Spearman correlation analyses, simple and multiple linear regression models, t-tests, and graph generation were performed using Prism v10.6.0 (GraphPad). Statistical tests and significance are specified in corresponding figure legends.

## KEY RESOURCES

| Reagent/Resource | Source | Identifier/Catalog # | RRID |
| --- | --- | --- | --- |
| **Cell cultures** |  |  |  |
| Primary dermal fibroblasts | San Diego Nathan Shock Center | See **Table S1** | N/A |
| UNA+ primary dermal fibroblasts | San Diego Nathan Shock Center; this study | See **Table S1** | N/A |
| HEK293T | Cell Biolabs | LTV-100 | CVCL_JZ09 |
| **Antibodies** |  |  |  |
| LC3B mouse mAb | MBL Life Science | M152-3 | AB_1279144 |
| LC3B rabbit mAb | Cell Signaling | 3868 | AB_2137707 |
| TUJ1 mouse mAb | BioLegend | 801202 | AB_2313773 |
| Tubulin mouse mAb | Cell Signaling | 3873 | AB_1904178 |
| Alexa Fluor 488-labeled goat-anti-mouse IgG | Invitrogen | A11001 | AB_2534069 |
| Anti-rabbit IgG HRP | Cell Signaling | 7074 | AB_2099233 |
| Anti-mouse IgG HRP | Cell Signaling | 7076 | AB_330924 |
| Alexa Fluor 647-labeled goat-anti-mouse IgG | Invitrogen | A32728 | AB_2633277 |
| **Software tools** |  |  |  |
| Nf-core/rnaseq | N/A | N/A | SCR_026973 |
| RStudio | N/A | N/A | SCR_000432 |
| CellProfiler v4.2.8 | Broad Institute | N/A | SCR_007358 |
| ImageJ2 v2.16.0/1.54p | N/A | N/A | SCR_002285 |
| Prism | GraphPad | N/A | SCR_002798 |
| **Plasmids** |  |  |  |
| pLVXUbC-rtTA-Ngn2:2A:Ascl1 | Addgene | 127289 | Addgene_127289 |
| psPAX2 | Addgene | 12260 | Addgene_12260 |
| pMD2.G | Addgene | 12259 | Addgene_12259 |
| pCDH-EF1a-mCherry-EGFP-LC3B | Addgene | 170446 | Addgene_170446 |

## ACKNOWLEDGMENTS

This work was supported by a Conrad Prebys Foundation Predoctoral Fellowship (TMM), San Diego Nathan Shock Center P30-AG068635 (pilot grant to CK), NIH-NIA R01-AG083373 (CK), NIH-NIA R01-AG085634 (JM), NIH-NIA Longevity Consortium U19-AG023122 (JM), the American Federation for Aging

Research AFAR-30376715 (JM). Funding for core facilities: NCI Cancer Center Support Grant P30 CA030199 and Shared Instrumentation Grant S10 OD036254, The Razavi Newman Integrative Genomics and Bioinformatics Core Facility of the Salk Institute (RRID:SCR_014842 and SCR_014846) with funding from NIH-NCI CCSG P30 CA014195, NIH-NIA San Diego Nathan Shock Center P30 AG068635, the NIH-NIA Liver Cancer P01 AG073084-04, the Howard and Maryam Newman Family Foundation and the Helmsley Trust.

Figure 3A was generated with help from BioRender.com.

## AUTHOR CONTRIBUTIONS

TMM: conceptualization, data curation, formal analysis, funding acquisition, investigation, methodology, resources, software, supervision, validation, visualization, writing – original draft, writing – review and editing.

SRH: data curation, investigation, methodology, resources, validation.

RJM: conceptualization, funding acquisition, project supervision, writing – review. HSB: data curation, investigation, writing – review.

LS: data curation, project administration, resources.

LT: data curation, investigation, methodology, resources, writing – review and editing. AN: data curation, investigation, resources, writing – review and editing.

SD: methodology, resources.

JB: formal analysis, writing – review and editing.

SSR: data curation, formal analysis, resources, software, writing – review and editing. AB: conceptualization, resources.

JM: conceptualization, resources.

DW: conceptualization, investigation, methodology, project supervision, writing – review. AJM: conceptualization, resources.

CK: conceptualization, funding acquisition, project administration, resources, supervision, writing – review and editing.

## DECLARATION OF GENERATIVE AI AND AI-ASSISTED TECHNOLOGIES

During the preparation of this work, the authors used ChatGPT v5.2 to improve grammar and clarity. After using this tool, the authors reviewed and edited the content as needed and take full responsibility for the content of the publication.

## SUPPLEMENTAL INFORMATION

Document S1. Figures S1-S9, Tables S1-S8.

