## Supplementary material for "Autophagy flux during human aging is sex- and cell type-specific, and is associated with physical fitness": DocumentS1

**Supplemental document S1.**

**Contents**

**Figure S1.** Global and autophagy-related differential gene expression in young and older primary dermal fibroblasts and iNs.

**Figure S2.** Chloroquine treatment reveals autophagy flux in primary dermal fibroblasts and iNs.

**Figure S3.** LC3B-positive puncta area is not associated with cell size following chloroquine treatment in primary dermal fibroblasts and iNs.

**Figure S4.** PBMC isolation enriches for lymphocytes and yields consistent cell composition across samples.

**Figure S5.** Chloroquine treatment of freshly drawn whole blood reveals autophagy flux in human PBMCs.

**Figure S6.** PBMC composition in whole blood and isolated PBMCs does not correlate with age or autophagy flux.

**Figure S7.** Changes in SHOCK cohort health and fitness with age.

**Figure S8.** Comparison of health and fitness readouts in STRONG participants between week 0 and week 12 of mild exercise.

**Figure S9.** Comparison of health and fitness readouts in SHOCK PBMC donors and STRONG participants.

**Table S1.** Experimental details, demographics and health measures of participants of the San Diego Nathan Shock Center Healthy Aging ("SHOCK") cohort.

**Table S2.** Table S2. PANTHER Slim-GO Biological Process module scores in SHOCK fibroblasts from older versus younger donors.

**Table S3.** Significantly up- and downregulated GSEA terms with age in SHOCK primary dermal fibroblasts.

**Table S4.** Significantly up- and downregulated GSEA terms with age in SHOCK induced neurons.

**Table S5.** Summary of autophagy-related gene modules defined by the PANTHER Classification System.

**Table S6.** Assignment of autophagy-related genes to GO biological Process modules and regulatory categories.

**Table S7.** Demographics and health measures of the STRONG cohort before and after a 12-week exercise intervention.

**Table S8.** Demographics and health measures of the SHOCK PBMC cohort and STRONG cohort before and after a 12-week exercise intervention.

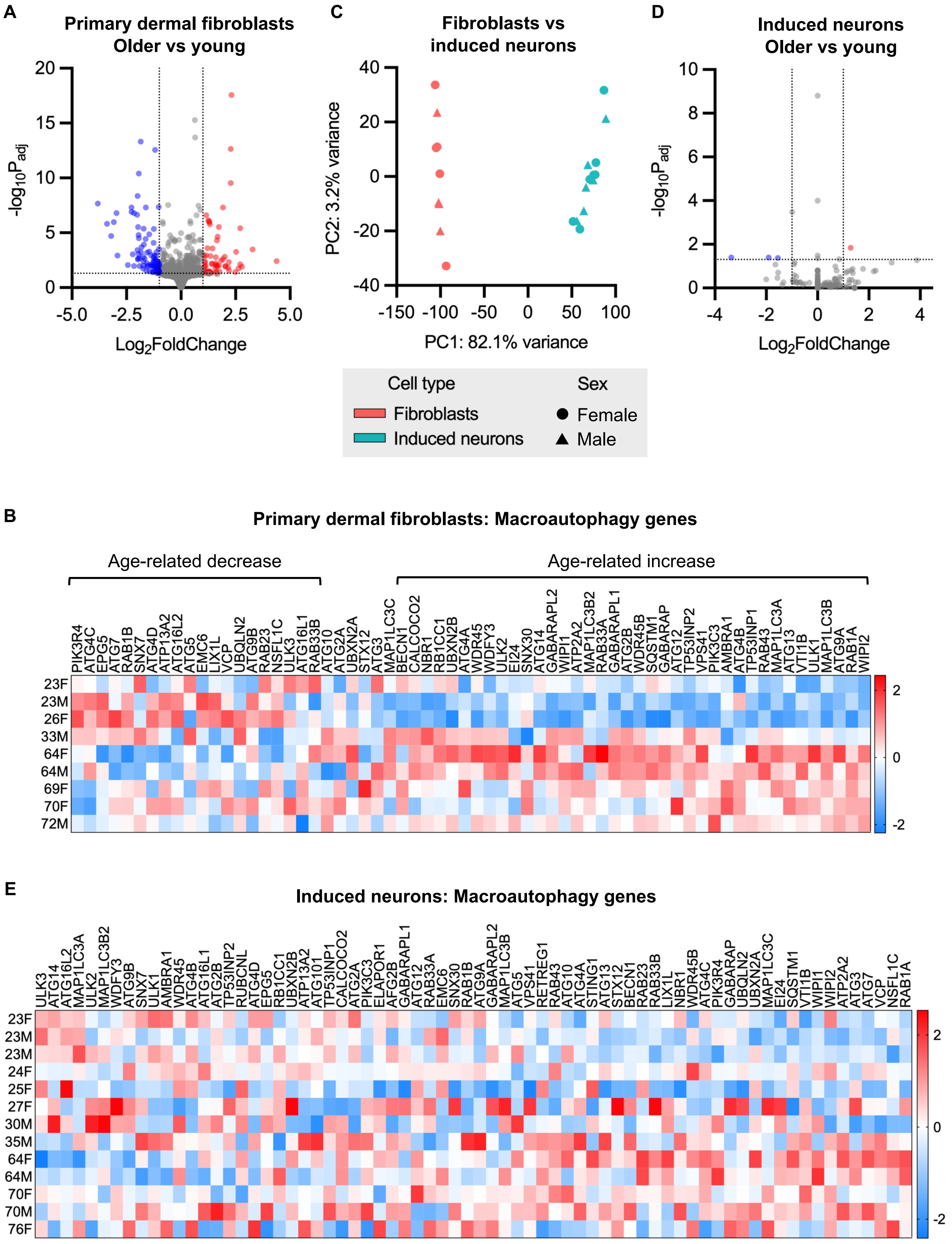
A
Primary dermal fibroblasts
Older vs young
-log10Padj
Log2FoldChange
20
15
10
5
0
-5.0
-2.5
0.0
2.5
5.0
C
Fibroblasts vs
induced neurons
PC2: 3.2% variance
PC1: 82.1% variance
40
20
0
-20
-40
-150
-100
-50
0
50
100
D
Induced neurons
Older vs young
-log10Padj
Log2FoldChange
10
8
6
4
2
0
-4
-2
0
2
4
Cell type
Fibroblasts
Induced neurons
Sex
Female
Male
B
Primary dermal fibroblasts: Macroautophagy genes
Age-related decrease
Age-related increase
PIK3R4 ATG4C EPG5 ATG7 RAB1B SNX7 ATG4D ATP13A2 ATG16L2 ATG5 EMC6 LIX1L VCP UBQLN2 ATG9B RAB23 NSFL1C ULK3 ATG16L1 RAB33B ATG10 ATG2A UBXN2A STX12 ATG3 MAP1LC3C BECN1 CALCOCO2 NBR1 RB1CC1 UBXN2B ATG4A WDR45 WDFY3 ULK2 EI24 SNX30 ATG14 GABARAPL2 WIPI1 ATP2A2 MAP1LC3B2 RAB33A GABARAPL1 ATG2B WDR45B SQSTM1 GABARAP ATG12 TP53INP2 VPS41 PIK3C3 AMBRA1 ATG4B TP53INP1 RAB43 MAP1LC3A ATG13 VTI1B ULK1 MAP1LC3B ATG9A RAB1A WIPI2
23F
23M
26F
33M
64F
64M
69F
70F
72M
2
0
-2
E
Induced neurons: Macroautophagy genes
ULK3 ATG14 ATG16L2 MAP1LC3A ULK2 MAP1LC3B2 WDFY3 ATG9B SNX7 ULK1 AMBRA1 WDR45 ATG4B ATG16L1 ATG2B TP53INP2 RUBCNL ATG4D EPG5 RB1CC1 UBXN2B ATP13A2 ATG101 TP53INP1 CALCOCO2 ATG2A PIK3C3 ELAPOR1 AFG2B GABARAPL1 ATG12 RAB33A EMC6 SNX30 RAB1B ATG9A GABARAPL2 MAP1LC3B ATG5 VPS41 RETREG1 RAB43 ATG10 ATG4A STING1 ATG13 STX12 BECN1 RAB23 RAB33B LIX1L NBR1 WDR45B ATG4C PIK3R4 GABARAP UBQLN2 UBXN2A MAP1LC3C EI24 SQSTM1 VTI1B WIPI1 WIPI2 ATP2A2 ATG3 ATG7 VCP NSFL1C RAB1A
23F
23M
23M
24F
25F
27F
30M
35M
64F
64M
70F
70M
76F
2
0
-2

**Figure S1. Global and autophagy-related differential gene expression in young and older primary dermal fibroblasts and iNs.**

(**A**) Volcano plot of differentially-expressed genes in older vs young primary dermal fibroblasts (|$\log_2$ fold change cutoff| ≥ 1.0; $p_{adj} < 0.05$). (**B**) Z-scored expression of GO Biological Process macroautophagy genes in primary dermal fibroblasts ordered by ascending age. (**C**) PCA of primary dermal fibroblast and iN samples labeled by cell type and donor sex. (**D**) Volcano plot of differentially-expressed genes in older vs young iNs (|$\log_2$ fold change cutoff| ≥ 1.0; $p_{adj} < 0.05$). (**E**) Z-scored expression of GO Biological Process macroautophagy genes in iNs ordered by ascending age. Statistical analysis: Differential expression in (**A**) and (**D**) was determined using DESeq2 with Benjamini-Hochberg adjustment ($p_{adj} < 0.05$). Each sample represents one biological replicate (independent donor; fibroblasts: N = 4 young, N = 5 older; iNs: N = 8 young, N = 5 older).

**Primary dermal fibroblasts**

**Induced neurons**

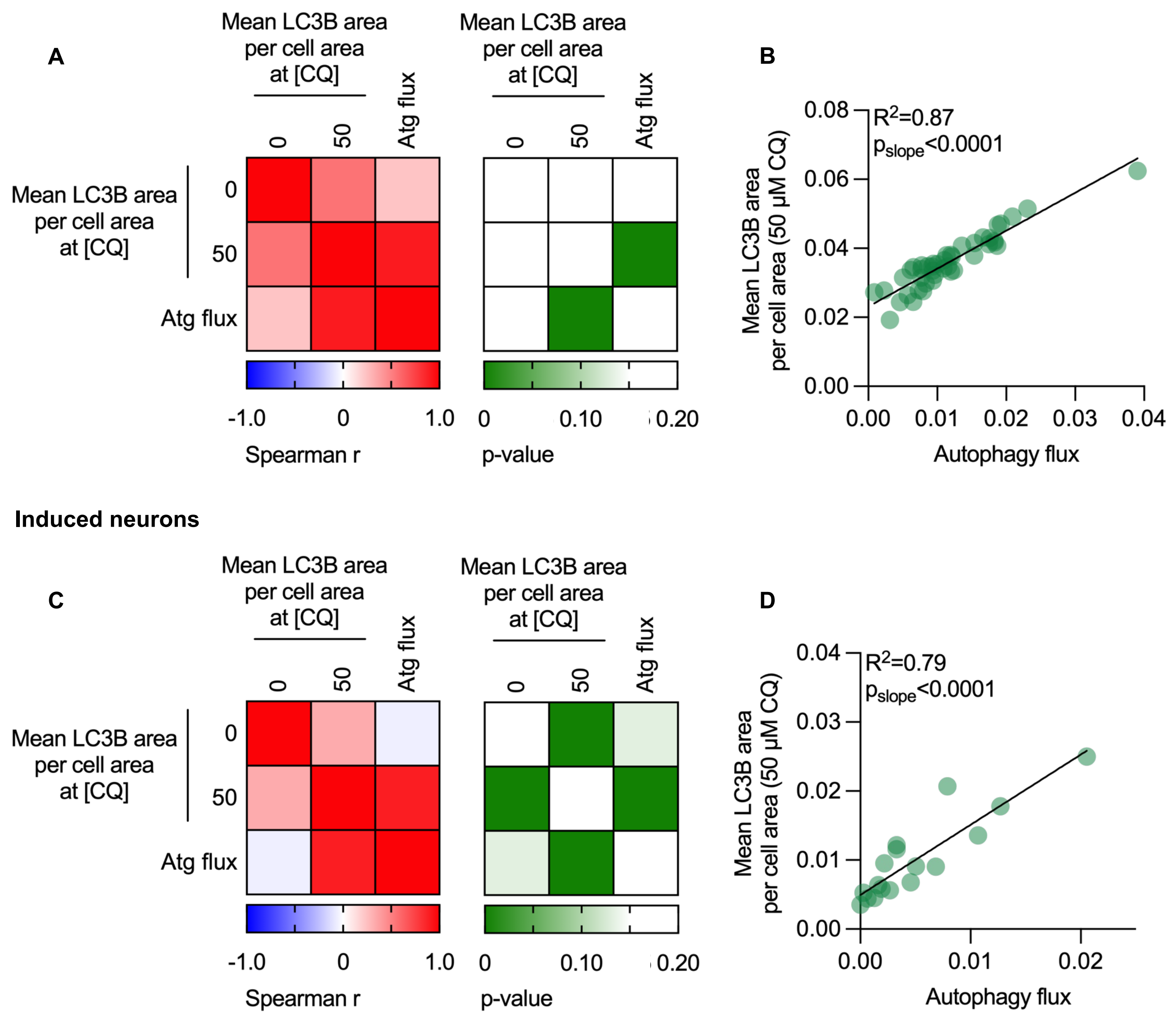

**Figure S2. Chloroquine treatment reveals autophagy flux in primary dermal fibroblasts and iNs.**
(**A**) Spearman correlation coefficients (left) and corresponding p-values (right) of autophagy flux assay measurements from assays performed in primary dermal fibroblasts from SHOCK cohort donors (N = 44). (**B**) Correlation between autophagy flux and mean LC3B-positive puncta area per cell area following 50 µM CQ treatment in primary dermal fibroblasts, with each point representing one donor. Linear regression shown ($R^2$ and p-values from linear regression model). (**C**) Spearman correlation coefficients (left) and corresponding p-values (right) of autophagy flux assay measurements from assays performed in iNs from SHOCK cohort donors (N = 17). (**D**) Correlation between autophagy flux and mean LC3B-positive puncta area per cell area at 50 µM CQ treatment in iNs, with each point representing one donor. Linear regression shown ($R^2$ and p-values from linear regression model). Autophagy flux is defined as the CQ-induced change in LC3B or GFP-LC3B-positive puncta area per cell (CQ-CTRL).

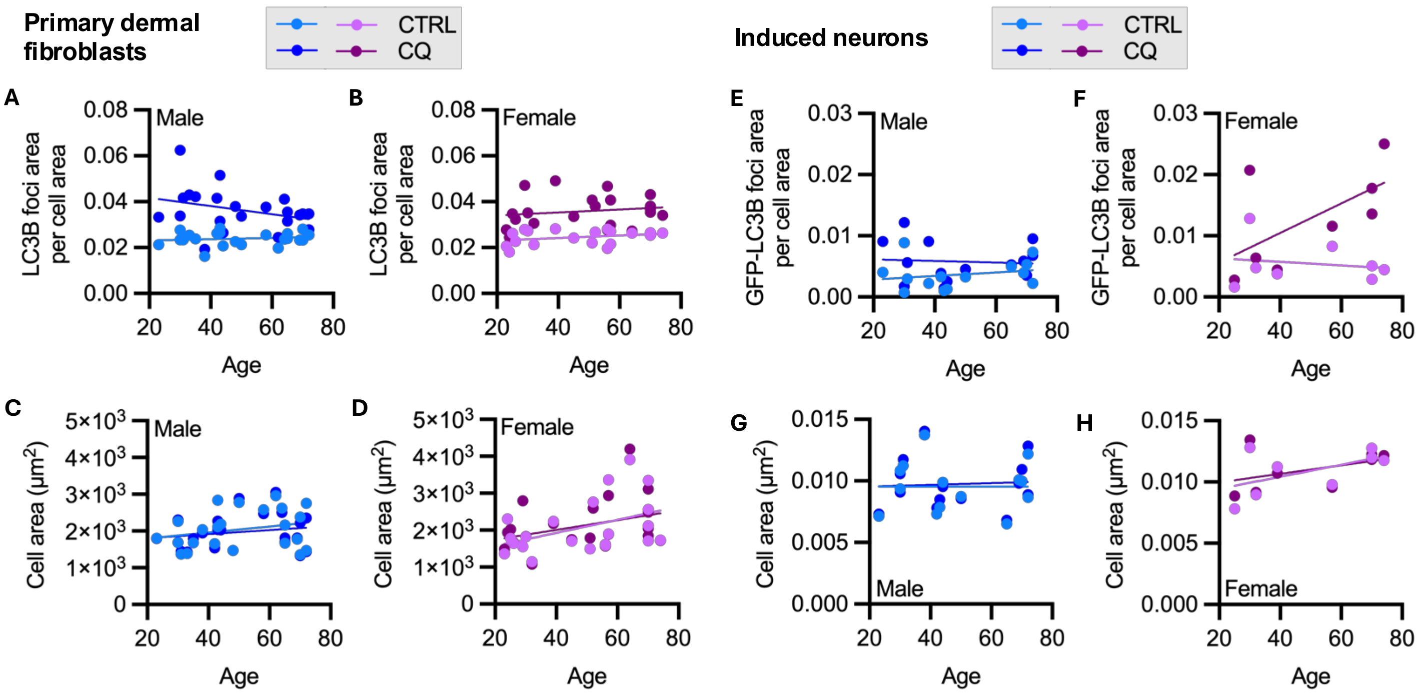

**Figure S3. LC3B-positive foci area is not associated with cell size following chloroquine treatment in primary dermal fibroblasts and iNs.**

(**A**) Mean LC3B-positive foci area per cell area in primary dermal fibroblasts from male SHOCK cohort donors treated ± CQ (N = 23). Linear regression per treatment by age ($R^2_{CTRL}$= 0.03; $R^2_{CQ}$ = 0.10); comparison of CTRL vs CQ slopes, p = 0.09; comparison of CTRL vs CQ intercepts, p < 0.0001. (**B**) Mean LC3B-positive foci area per cell area in primary dermal fibroblasts from female SHOCK cohort donors treated ± CQ (N = 21). Linear regression per treatment by age ($R^2_{CTRL}$ = 0.10; $R^2_{CQ}$ = 0.03); comparison of CTRL vs CQ slopes, p = 0.91; comparison of CTRL vs CQ intercepts, p < 0.0001. (**C**) Mean cell area of primary dermal fibroblasts from male SHOCK cohort donors treated ± CQ (N = 23). Comparison of CTRL vs CQ, p = 0.13 by Wilcoxon matched-pairs signed rank test. (**D**) Mean cell area of primary dermal fibroblasts from female SHOCK cohort donors treated ± CQ (N = 21). Comparison of CTRL vs CQ, p > 0.99 by Wilcoxon matched-pairs signed rank test. (**E**) Mean GFP-LC3B-positive foci area per cell body area in iNs from male SHOCK cohort donors treated ± CQ (N = 14). Linear regression per treatment by age ($R^2_{CTRL}$= 0.05; $R^2_{CQ}$ = 0.01); comparison of CTRL vs CQ slopes, p = 0.52; comparison of CTRL vs CQ intercepts, p = 0.06. (**F**) Mean GFP-LC3B-positive foci area per cell body area in iNs from female SHOCK cohort donors treated ± CQ (N = 8). Linear regression per treatment by age ($R^2_{CTRL}$ = 0.03; $R^2_{CQ}$ = 0.38); comparison of CTRL vs CQ slopes, p = 0.09; comparison of CTRL vs CQ intercepts, p = 0.03. (**G**) Mean cell body area of iNs from male SHOCK cohort donors treated ± CQ (N = 14). Comparison of CTRL vs CQ, p = 0.09 by Wilcoxon matched-pairs signed rank test. (**H**) Mean cell body area of iNs from female SHOCK cohort donors treated ± CQ (N = 8). Comparison of CTRL vs CQ, p > 0.74 by Wilcoxon matched-pairs signed rank test.

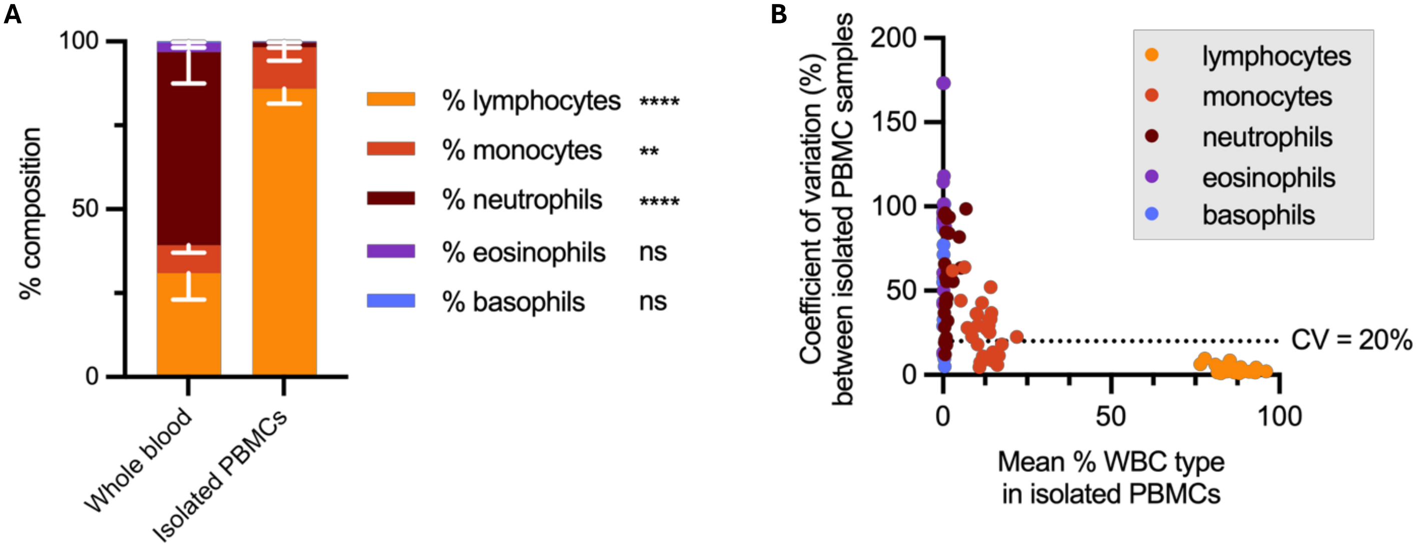

**Figure S4. PBMC isolation enriches for lymphocytes and yields consistent cell composition across samples.** (**A**) PBMC composition in whole blood and isolated PBMC samples from SHOCK cohort participants, measured by Coulter counter analysis (N = 23). Mean ± standard deviation. Statistical comparison of cell type abundance before and after PBMC isolation were performed by two-way ANOVA; ns – p>0.05; *p<0.05; **p<0.01; ***p<0.001, ****p<0.0001. (**B**) Coefficient of variation (CV) of PBMC cell type abundance (% of total cells) across 0, 75 and 200 μM CQ-treated samples from SHOCK participants. Each data point represents the CV calculated from 3 treatment conditions for one participant. Dotted line indicates CV threshold of 20%.

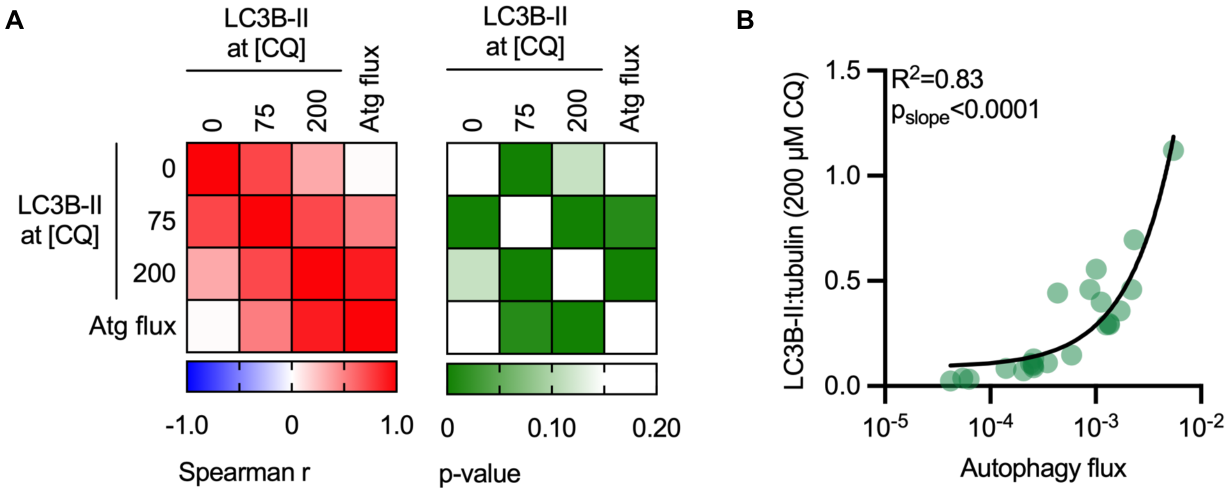

**Figure S5. Chloroquine treatment of freshly drawn whole blood reveals autophagy flux in human PBMCs.**
(**A**) Spearman correlation coefficients (left) and corresponding p-values (right) of autophagy flux assay measurements in freshly isolated PBMCs from SHOCK cohort donors (N = 23). (**B**) Correlation between autophagy flux and LC3B-II levels following 200 µM CQ treatment, with $R^2$ and p-values from linear regression analysis. The $log_{10}$-scaled x-axis results in the curved appearance of the linear trend line.

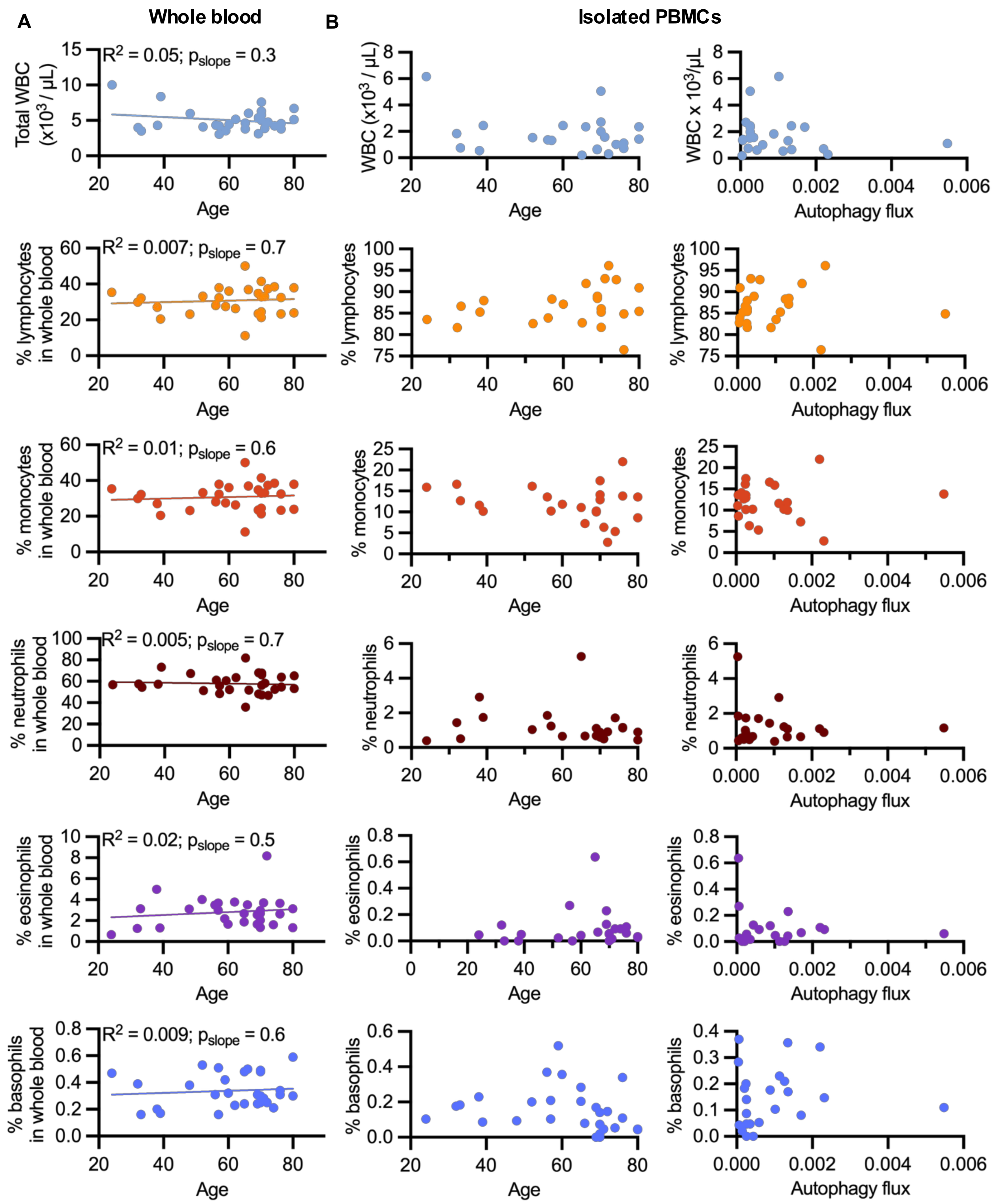

**Figure S6. PBMC composition in whole blood and isolated PBMCs does not correlate with age or autophagy flux.** (**A**) Relationship between age and total white blood cell counts (WBC; top), and PBMC cell type composition (bottom) in whole blood from SHOCK cohort participants. Each data point represents one participant. $R^2$ values and $p_{slope}$ derived from linear regression. (**B**) Relationship between total white blood cell counts (WBC; top), and PBMC cell type composition (bottom) with age (left) and autophagy flux (right) in SHOCK cohort participant isolated PBMC samples. Each data point represents the mean abundance of the indicated cell type across 0, 75 and 200 µM CQ-treated samples for each individual.

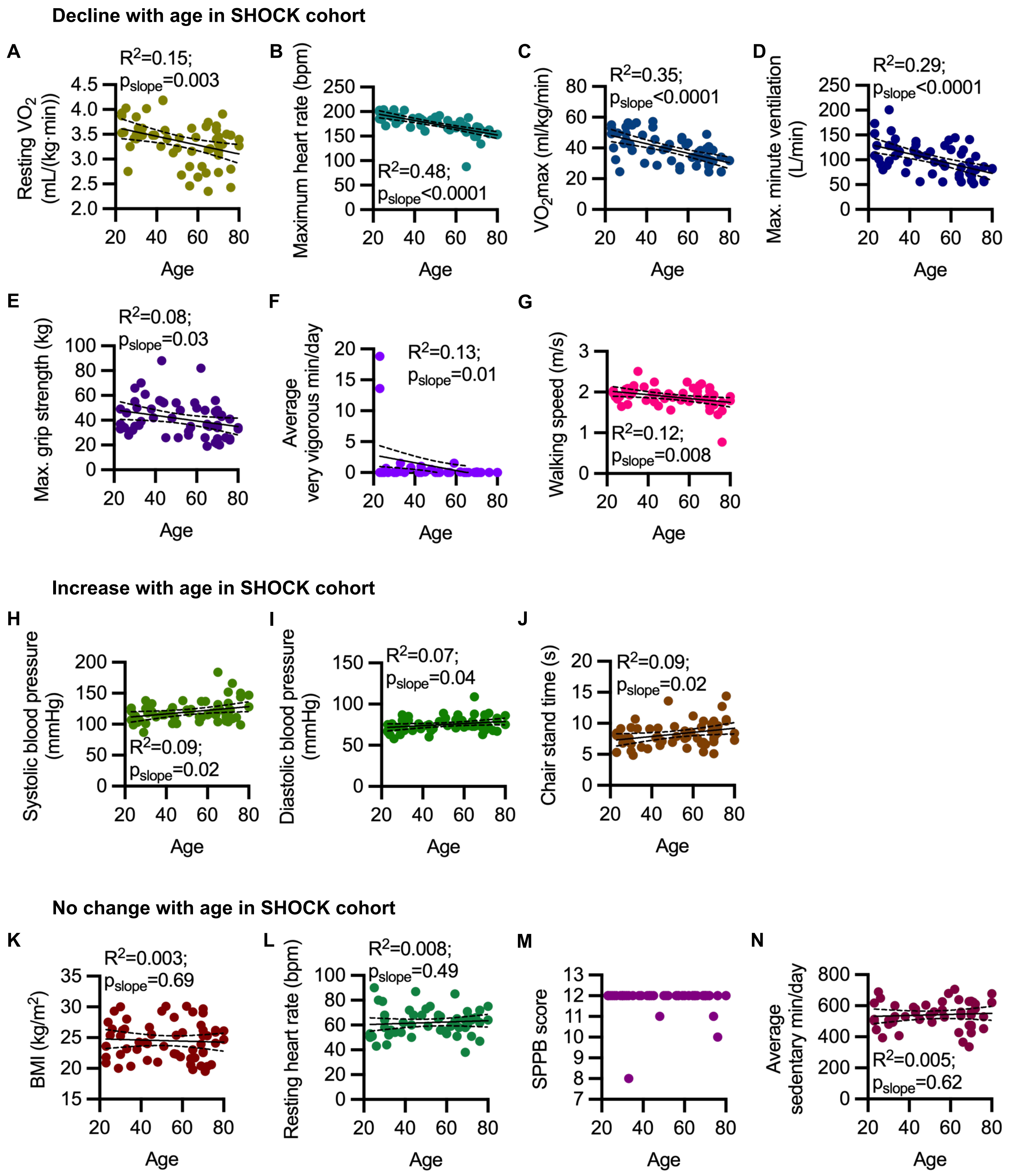

**Figure S7. Changes in SHOCK cohort health and fitness with age.**
Health and fitness readout for all SHOCK participants (N = 49-58) as a function of age. (**A**) resting $VO_2$; (**B**) maximum heart rate; (**C**) $VO_{2max}$; (**D**) maximum minute ventilation (VE); (**E**) maximal grip strength of strongest hand; (**F**) average very vigorous minutes per day; (**G**) walking speed in a 6-minute walking test; (**H**) systolic blood pressure; (**I**) diastolic blood pressure; (**J**) SPPB test chair stand time; (**K**) body mass index (BMI); (**L**) resting heart rate; (**M**) short physical performance battery (SPPB) score out of 12; (**N**) average sedentary minutes per day.

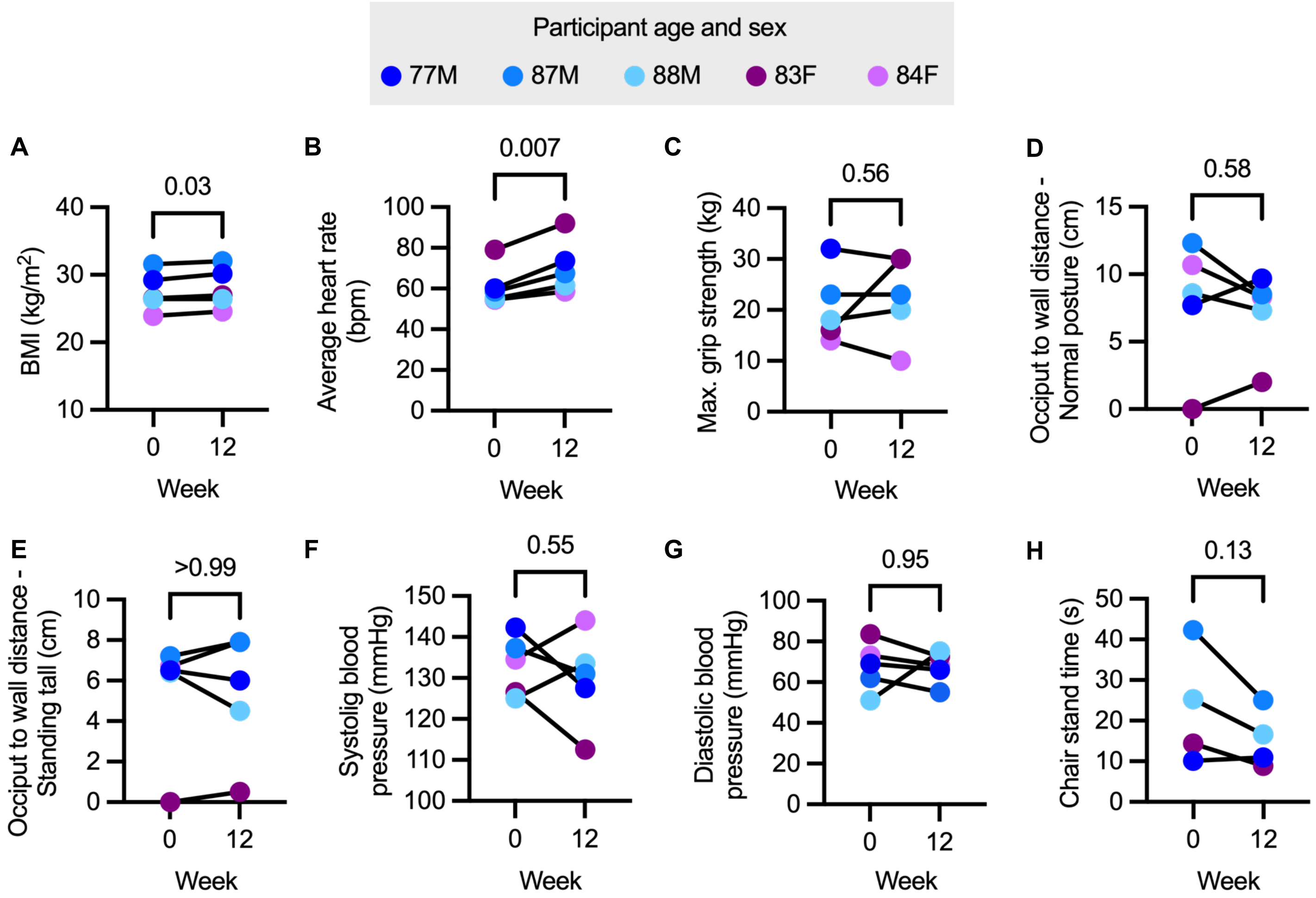

**Figure S8. Comparison of health and fitness readouts in STRONG participants between week 0 and week 12 of mild exercise.**

Week 0 and week 12 STRONG participant (N = 5) (**A**) body mass index (BMI); (**B**) average heart rate; (**C**) maximal grip strength of strongest hand; (**D**) occiput to wall distance standing with a normal posture; (**E**) occiput to wall distance standing as tall as possible; (**F**) systolic blood pressure; (**G**) diastolic blood pressure; (**H**) SPPB test chair stand time. P-values by paired t-tests.

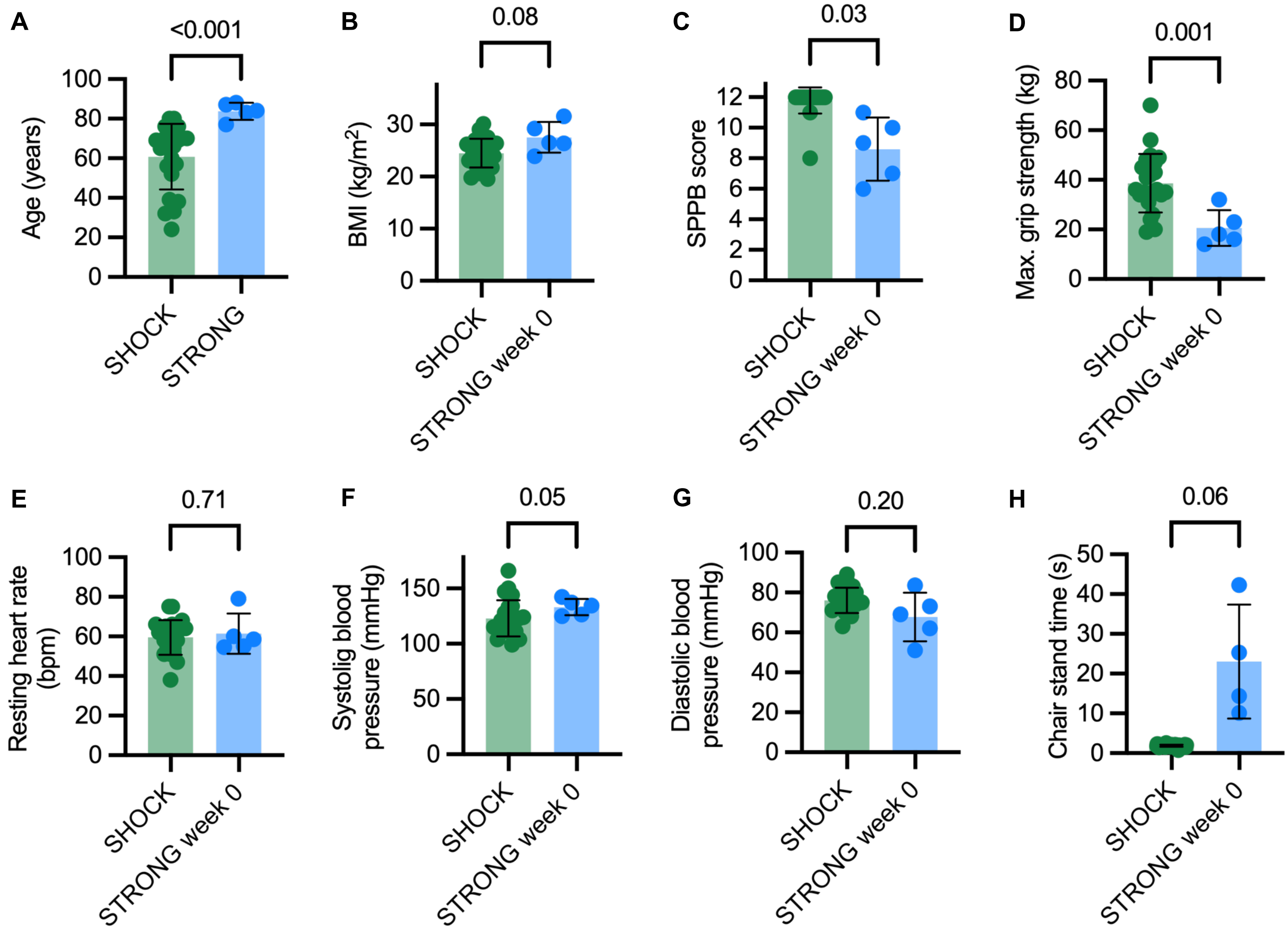

**Figure S9. Comparison of health and fitness readouts in SHOCK PBMC donors and STRONG participants.**
(**A**) SHOCK and STRONG participant age. SHOCK PBMC donor (N = 23), and week 0 STRONG participant (N = 5) (**B**) body mass index (BMI); (**C**) short physical performance battery (SPPB) score; (**D**) maximal grip strength of strongest hand; (**E**) resting heart rate; (**F**) systolic blood pressure; (**G**) diastolic blood pressure; (**H**) SPPB test chair stand time. P-values by Welch's t-test.

Table S1. Experimental details, demographics and health measures of participants of the San Diego Nathan Shock Center Healthy Aging ("SHOCK") cohort.

| | Assay / Cell type / Passage # | | | | | | | | | Demographics | | | | Health and fitness readouts | | | | | | | | | | | | | |
|---|---|---|---|---|---|---|---|---|---|---|---|---|---|---|---|---|---|---|---|---|---|---|---|---|---|---|---|
| | RNA-seq | | | | Autophagy flux | | | | | | | | | | | | | | | | | | | | | | |
| | Fibroblast | | iN | | Fibroblast | | iN | | PBMC | | | | | | | | | | | | | | | | | | |
| SHOCK donor ID | Assayed? | Passage # | Assayed? | Passage # | Assayed? | Passage # | Assayed? | Passage # | Assayed? | Age (years) | Sex | Race | Hispanic/Latino? | BMI (kg/m$^2$) | Resting $VO_2$ (mL/(kg·min)) | Systolic blood pressure (mmHg) | Diastolic blood pressure (mmHg) | Resting heart rate (bpm) | Maximum heart rate (bpm) | $VO_2$max (mL/(kg·min)) | Maximum minute ventilation (VE; L/min) | Maximal grip strength (kg) | Walking speed (m/s) | Chair stand time (s) | SPPB score | Average Freedson sedentary minutes per day | Average Freedson very vigorous minutes per day |
| 201 | X | P9 | X | P15.6 | X | P6 | | | | 23 | F | White | X | 20.9 | 3.87 | 107 | 68 | 53 | 185 | 49.31 | 108.55 | 37 | 8.43 | 1.99 | 12 | 502.3 | 13.6 |
| 202 | | | | | X | P11 | X | | | 30 | M | White | | 25.29 | 3.41 | 125 | 80 | 68 | 195 | 53.67 | 200.92 | 66 | 6.69 | 2.09 | 12 | 480.9 | 0.0 |
| 203 | | | X | P15.7 | X | P9 | | | | 35 | M | White | | 20.34 | 4.02 | 112 | 68 | 54 | 203 | 56.47 | 116.84 | 61 | 6.16 | 2.51 | 12 | 407.5 | 0.0 |
| 204 | X | P7 | | | X | P4 | | | | 26 | F | White | | 26.4 | 2.75 | 103 | 58 | 43 | 171 | 49.71 | 90.58 | 46 | 6.80 | 1.88 | 12 | 648.5 | 0.7 |
| 205 | | | | | X | P4 | | | | 51 | F | White | | 23.6 | 3.43 | 119 | 83 | 59 | 178 | 42.43 | 71.45 | 26 | 9.47 | 1.56 | 12 | 494.5 | 0.0 |
| 207 | X | P8 | | | X | P5 | X | P17.11 | | 72 | M | White | | 20.3 | 3.91 | 109 | 72 | 64 | 163 | 38.66 | 80.14 | 44 | 9.62 | 1.93 | 12 | 611.6 | 0.0 |
| 208 | | | | | X | P8 | X | P16.10 | | 50 | M | Asian | | 29.3 | 3.22 | 123 | 89 | 73 | 189 | 39.85 | 116.48 | 54 | 6.94 | 1.96 | 12 | 539.3 | 0.0 |
| 209 | | | | | X | P9 | X | | | 43 | M | White | | 21.3 | 4.18 | 127 | 75 | 60 | 185 | 57.25 | 137.40 | 55 | 7.77 | 2.24 | 12 | 555.5 | 0.0 |
| 210 | | | | | X | P9 | X | | | 42 | M | White | | 24.2 | 3.26 | 120 | 74 | 72 | 188 | 38.60 | 119.41 | 56 | 7.66 | 1.97 | 12 | 591.0 | 0.0 |
| 212 | | | | | X | P8 | | | | 45 | F | Asian | | 25.5 | 3.41 | 101 | 70 | 87 | 160 | 28.69 | 55.99 | 26 | 7.12 | 1.66 | 12 | 512.1 | 0.1 |
| 213 | | | | | X | P12 | X | P15.9 | X | 38 | M | White | X | 23.1 | 3.48 | 115 | 76 | 54 | 168 | 48.43 | 74.37 | 49 | 5.90 | 2.13 | 12 | 530.6 | 0.8 |
| 215 | X | P8 | X | P11.5 | X | P5 | | | | 70 | F | White | | 29.7 | 3.17 | 119 | 76 | 60 | 149 | 24.41 | 58.57 | 21 | 9.69 | 1.55 | 12 | 474.2 | 0.0 |
| 216 | | | | | X | P10 | X | | | 44 | M | White | | 26.7 | 3.58 | 111 | 68 | 68 | 182 | 45.62 | 110.65 | 54 | 6.45 | 1.98 | 12 | - | - |
| 219 | X | P7 | | | | | | | | 69 | F | White | | 20.4 | 3.67 | 105 | 63 | 64 | 160 | 39.80 | 72.99 | 22 | 8.72 | 1.82 | 12 | 627.5 | 0.0 |
| 220 | X | P9 | X | P13.7 | X | P6 | X | P17.9 | | 23 | M | White | | 21.8 | 3.53 | 126 | 75 | 50 | 199 | 52.87 | 152.87 | 33 | 8.07 | 2.03 | 12 | 616.9 | 18.8 |
| 221 | X | P8 | X | P13.5 | X | P7 | | | | 64 | F | White | | 20.6 | 3.66 | 115 | 73 | 63 | 164 | 34.63 | 70.09 | 26 | 6.75 | 2.02 | 12 | 427.3 | 0.0 |
| 223 | X | P6 | | | X | P9 | | | | 33 | M | Asian | | 28 | 3.48 | 120 | 80 | 56 | 189 | 38.36 | 108.68 | 39 | 9.16 | 2.16 | 12 | 600.6 | 1.5 |
| 224 | | | | | X | P10 | X | P14.8 | | 65 | M | White | | 23.04 | 3.74 | 184 | 109 | 65 | 88 | 36.19 | 85.11 | 36 | 5.40 | 2.16 | 12 | - | - |
| 225 | | | | | X | P8 | X | P12.8 | | 31 | M | White | | 26.2 | 3.63 | 118 | 68 | 44 | 176 | 56.41 | 158.43 | 50 | 4.84 | 1.93 | 12 | 498.6 | 0.0 |
| 227 | X | P5 | X | P12.6 | X | P10 | | | | 64 | M | White | | 21.6 | 2.72 | 120 | 79 | 85 | 171 | 28.21 | 64.45 | 48 | 9.38 | 1.91 | 12 | 638.5 | 0.0 |
| 231 | | | X | P10.5 | X | P8 | | | | 70 | M | White | | 25.3 | 3.31 | 152 | 87 | 62 | 158 | 29.47 | 72.94 | 48 | 7.03 | 2.11 | 12 | 425.8 | 0.0 |
| 233 | | | X | P12.5 | X | P10 | X | | | 25 | F | White | | 25.4 | 4.03 | 113 | 76 | 90 | 204 | 32.34 | 101.43 | 34 | 9.16 | 1.83 | 12 | 689.8 | 0.0 |
| 234 | | | X | P11.5 | | | | | | 23 | M | White | | 23.6 | 3.93 | 121 | 66 | 51 | 203 | 55.11 | 173.36 | 49 | 5.31 | 1.97 | 12 | 513.1 | 0.0 |
| 235 | | | X | P13.6 | X | P11 | X | P15.10 | | 30 | M | Asian | | 30 | 3.49 | 138 | 87 | 64 | 204 | 45.21 | 128.86 | 55 | 5.50 | 1.99 | 12 | 507.4 | 0.0 |
| 236 | | | X | P11.6 | X | P5 | | | X | 24 | F | White | | 27.5 | 3.54 | 99 | 63 | 51 | 184 | 40.00 | 128.80 | 35 | 8.09 | 1.89 | 12 | 445.9 | 0.0 |
| 237 | | | | | X | P10 | | | | 43 | M | White | | 24 | 4.19 | 101 | 60 | 50 | 172 | 53.54 | 130.33 | 88 | 9.22 | 1.76 | 12 | 578.8 | 1.0 |
| 238 | | | X | P11.5 | | | | | | 27 | F | White | X | 29.3 | 3.25 | 106 | 72 | 80 | 194 | 24.60 | 79.70 | 28 | 7.47 | 1.64 | 12 | 394.8 | 0.0 |
| 239 | | | | | X | P5 | X | P12.9 | X | 32 | F | White | | 26.4 | 3.50 | 123 | 78 | 52 | 183 | 42.30 | 127.82 | 36 | 10.70 | 1.97 | 12 | 494.0 | 0.4 |
| 241 | | | | | X | P5 | | | | 57 | F | White | | 25.4 | 3.16 | 137 | 87 | 67 | 163 | 45.21 | 128.86 | 34 | 5.97 | 1.97 | 12 | 526.9 | 0.0 |
| 242 | | | | | X | P6 | X | P13.9 | X | 57 | F | White | | 22.3 | 3.62 | 111 | 75 | 56 | 169 | 46.90 | 87.34 | 35 | 7.75 | 1.86 | 12 | - | - |
| 244 | | | | | X | P8 | | | | 56 | F | White | | 28.3 | 2.46 | 104 | 71 | 65 | 166 | 25.67 | 55.64 | 28 | 7.56 | 1.81 | 12 | 571.9 | 0.0 |
| 245 | | | | | X | P5 | | | | 62 | M | White | | 30.1 | 2.85 | 118 | 74 | 59 | 180 | 33.84 | 144.59 | 82 | 7.60 | 2.10 | 12 | 705.5 | 0.1 |
| 248 | | | | | X | P6 | | | X | 56 | F | White | X | 22.4 | 2.85 | 112 | 81 | 68 | 182 | 27.84 | 92.99 | 31 | 8.78 | 1.81 | 12 | 625.4 | 0.0 |
| 249 | | | | | X | P5 | | | | 65 | M | White | X | 29.2 | 3.19 | 132 | 88 | 51 | 158 | 37.26 | 111.48 | 36.8 | 6.94 | - | 12 | 536.4 | 0.0 |
| 251 | | | | | X | P5 | | | | 59 | M | White | | 21.9 | 3.44 | 137 | 80 | 55 | 172 | 42.03 | 139.63 | 54 | 8.59 | 2.25 | 12 | 679.0 | 1.5 |
| 253 | | | | | X | P6 | | | X | 52 | F | White | | 30.1 | 2.68 | 124 | 74 | 75 | 192 | 31.72 | 95.74 | 48 | 7.88 | 1.87 | 12 | 560.3 | 0.0 |
| 254 | | | X | P11.8 | | | | | | 76 | F | White | X | 24.1 | 3.48 | 99 | 67 | 72 | 162 | 34.13 | 58.58 | 25 | 14.40 | 1.75 | 10 | - | - |
| 255 | | | | | | | | | X | 33 | M | White | | 23.2 | 3.62 | 133 | 85 | 55 | 183 | 54.86 | 141.85 | 70 | - | 2.06 | 8 | 530.1 | 0.2 |
| 257 | | | | | X | P5 | | | | 70 | F | White | | 22.7 | 3.08 | 105 | 68 | 71 | 159 | 28.29 | 58.48 | 25 | 10.37 | 1.65 | 12 | 465.6 | 0.0 |
| 258 | | | | | X | P6 | | | | 48 | M | Asian | | 21 | 3.13 | 138 | 79 | 68 | 183 | 38.92 | 108.79 | 41.8 | 13.59 | 1.84 | 11 | 604.9 | 0.5 |
| 261 | | | | | | | | | X | 60 | M | Asian | | 25.8 | 2.64 | 115 | 72 | 53 | 167 | 43.21 | 120.80 | 49.8 | 11.19 | 1.77 | 12 | - | - |
| 262 | | | | | | | | | X | 76 | M | White | | 25.9 | 2.80 | 150 | 80 | 47 | 155 | 24.60 | 86.40 | 41 | 10.34 | 0.77 | 12 | - | - |
| 263 | | | | | | | | | X | 80 | F | White | | 24.7 | 3.40 | 128 | 75 | 64 | 153 | 31.86 | 82.37 | 33 | 8.41 | 1.79 | 12 | 620.4 | 0.0 |
| 265 | | | | | | | | | X | 65 | F | White | | 19.8 | 2.35 | 104 | 73 | 65 | 169 | 33.30 | 55.53 | 19 | 10.34 | 1.82 | 12 | 559.8 | 0.0 |
| 266 | | | | | | | | | X | 69 | M | White | | 27 | 3.59 | 115 | 71 | 58 | 149 | 33.83 | 95.16 | 56 | 9.72 | 1.86 | 12 | 336.5 | 0.0 |
| 267 | | | | | | | | | X | 76 | F | White, Asian | | 26.2 | 2.43 | 144 | 83 | 62 | 163 | 24.36 | 55.88 | 24 | 10.60 | 1.54 | 12 | 452.0 | 0.0 |
| 268 | | | | | | | | | X | 66 | M | White | | 21.74 | 3.21 | 123 | 74 | 58 | 188 | 38.39 | 120.24 | 35 | 6.62 | 2.20 | 12 | 365.8 | 0.0 |
| 269 | | | | | X | P7 | X | P11.6 | X | 70 | F | White | | 26.19 | 2.73 | 134 | 81 | 65 | 168 | 29.65 | 57.57 | 34 | 7.53 | 1.74 | 12 | 629.3 | 0.0 |
| 270 | | | | | | | | | X | 71 | M | White | | 19.51 | 3.33 | 123 | 75 | 66 | 168 | 31.85 | 51.67 | 20 | 8.22 | 1.78 | 12 | 621.7 | 0.0 |
| 271 | | | | | X | P9 | X | P12.8 | X | 69 | M | White | | 23.93 | 3.40 | 111 | 75 | 38 | 152 | 48.99 | 141.11 | 47 | 9.91 | 1.81 | 12 | 473.8 | 0.0 |
| 272 | | | | | X | P7 | X | P13.9 | X | 72 | M | White | | 25.31 | 3.30 | 166 | 89 | 51 | 134 | 31.37 | 107.11 | 43 | 8.53 | 1.69 | 12 | 580.0 | 0.0 |
| 273 | | | | | X | P9 | X | P13.9 | X | 74 | F | White | X | 20.59 | 3.50 | 111 | 68 | 66 | - | - | - | 26 | 12.72 | 1.45 | 11 | - | - |
| 274 | | | | | X | P9 | X | P12.8 | X | 70 | M | White | | 22.57 | 3.45 | 121 | 79 | 64 | 169 | 40.85 | 80.66 | 45 | 5.10 | 2.03 | 12 | 540.7 | 0.0 |
| 275 | | | | | | | | | X | 80 | M | White, Native Hawaiian/Pacific Islander | X | 26.12 | 3.27 | 147 | 86 | 75 | 154 | 31.79 | 81.43 | 34 | 7.25 | 1.89 | 12 | 677.1 | 0.0 |
| 276 | | | | | X | P9 | X | P12.8 | X | 70 | F | White | | 29.03 | 3.15 | 104 | 68 | 64 | 169 | 28.27 | 68.26 | 36 | 8.19 | 1.80 | 12 | 555.2 | 0.1 |
| 278 | | | | | X | P10 | X | P12.8 | X | 39 | F | White | X | 24.29 | 3.48 | 115 | 70 | 62 | 186 | 30.07 | 95.43 | 42 | 9.09 | 1.79 | 12 | - | - |
| 279 | | | | | X | P8 | | | | 29 | F | White | X | 20 | 3.42 | 87 | 63 | 79 | 189 | 42.97 | 88.68 | 32 | 7.66 | 2.06 | 12 | - | - |
| 280 | | | | | X | P10 | X | P12.8 | | 30 | F | Asian | | 22.29 | 3.85 | 101 | 74 | 58 | 188 | 49.38 | 97.56 | 35 | 9.31 | 1.70 | 12 | - | - |

Fibroblast, primary dermal fibroblast; iN, induced neuron; PBMC, peripheral blood mononuclear cell; M, male; F, female; BMI, body mass index; walking speed - in 6-minute walk test; chair stand time - in SPPB test; SPPB, Short Physical Performance Battery

Table S2. PANTHER Slim-GO Biological Process module scores in SHOCK fibroblasts from older versus younger donors.

| GO terms positively enriched with age | | | |
|---|---|---|---|
| GO ID | GO description | Gene count (input = 79) | Adjusted p-value |
| GO:0044057 | regulation of system process | 16 | 1.33E-09 |
| GO:0008015 | blood circulation | 13 | 3.61E-07 |
| GO:0003012 | muscle system process | 12 | 6.63E-07 |
| GO:1903522 | regulation of blood circulation | 9 | 1.77E-06 |
| GO:0003013 | circulatory system process | 13 | 2.63E-06 |
| GO:0010876 | lipid localization | 11 | 1.08E-05 |
| GO:0050731 | positive regulation of peptidyl-tyrosine phosphorylation | 7 | 1.22E-05 |
| GO:0003018 | vascular process in circulatory system | 8 | 3.28E-05 |
| GO:0015850 | organic hydroxy compound transport | 8 | 3.41E-05 |
| GO:0008016 | regulation of heart contraction | 7 | 3.74E-05 |
| GO:0014911 | positive regulation of smooth muscle cell migration | 4 | 6.15E-05 |
| GO:0044331 | cell-cell adhesion mediated by cadherin | 4 | 6.15E-05 |
| GO:0120254 | olefinic compound metabolic process | 6 | 6.68E-05 |
| GO:0090257 | regulation of muscle system process | 7 | 7.88E-05 |
| GO:0034308 | primary alcohol metabolic process | 5 | 8.71E-05 |
| GO:0072109 | glomerular mesangium development | 3 | 1.08E-04 |
| GO:0050730 | regulation of peptidyl-tyrosine phosphorylation | 7 | 1.10E-04 |
| GO:0060047 | heart contraction | 7 | 1.19E-04 |
| GO:0006869 | lipid transport | 9 | 1.41E-04 |
| GO:0006936 | muscle contraction | 8 | 1.54E-04 |
| GO:0001934 | positive regulation of protein phosphorylation | 11 | 1.58E-04 |
| GO:0070374 | positive regulation of ERK1 and ERK2 cascade | 6 | 1.63E-04 |
| GO:0003015 | heart process | 7 | 1.74E-04 |
| GO:1902430 | negative regulation of amyloid-beta formation | 3 | 1.90E-04 |
| GO:0036120 | cellular response to platelet-derived growth factor stimulus | 3 | 2.25E-04 |
| GO:0015844 | monoamine transport | 4 | 2.41E-04 |
| GO:0034332 | adherens junction organization | 4 | 2.41E-04 |
| GO:1902992 | negative regulation of amyloid precursor protein catabolic process | 3 | 2.63E-04 |
| GO:0006813 | potassium ion transport | 6 | 2.65E-04 |
| GO:0070372 | regulation of ERK1 and ERK2 cascade | 7 | 2.72E-04 |
| GO:0007507 | heart development | 11 | 2.82E-04 |
| GO:0098742 | cell-cell adhesion via plasma-membrane adhesion molecules | 6 | 2.97E-04 |
| GO:0036119 | response to platelet-derived growth factor | 3 | 3.06E-04 |
| GO:0099560 | synaptic membrane adhesion | 3 | 3.06E-04 |
| GO:0006809 | nitric oxide biosynthetic process | 4 | 3.29E-04 |
| GO:1990573 | potassium ion import across plasma membrane | 3 | 3.53E-04 |
| GO:0048144 | fibroblast proliferation | 5 | 3.61E-04 |
| GO:0018108 | peptidyl-tyrosine phosphorylation | 7 | 3.76E-04 |
| GO:0018212 | peptidyl-tyrosine modification | 7 | 3.99E-04 |
| GO:0043269 | regulation of monoatomic ion transport | 9 | 4.23E-04 |
| GO:0046209 | nitric oxide metabolic process | 4 | 4.38E-04 |
| GO:0016339 | calcium-dependent cell-cell adhesion via plasma membrane cell adhesion molecules | 3 | 4.59E-04 |
| GO:2000352 | negative regulation of endothelial cell apoptotic process | 3 | 4.59E-04 |
| GO:0032835 | glomerulus development | 4 | 4.69E-04 |
| GO:2001057 | reactive nitrogen species metabolic process | 4 | 4.69E-04 |
| GO:0070371 | ERK1 and ERK2 cascade | 7 | 4.84E-04 |
| GO:0014910 | regulation of smooth muscle cell migration | 4 | 5.01E-04 |
| GO:0017158 | regulation of calcium ion-dependent exocytosis | 3 | 5.85E-04 |
| GO:0072012 | glomerulus vasculature development | 3 | 5.85E-04 |
| GO:0062013 | positive regulation of small molecule metabolic process | 5 | 5.95E-04 |
| GO:0045860 | positive regulation of protein kinase activity | 7 | 6.65E-04 |
| GO:0061448 | connective tissue development | 7 | 6.83E-04 |
| GO:0007200 | phospholipase C-activating G protein-coupled receptor signaling pathway | 4 | 6.87E-04 |
| GO:0014909 | smooth muscle cell migration | 4 | 7.29E-04 |
| GO:0034381 | plasma lipoprotein particle clearance | 3 | 7.30E-04 |
| GO:0061437 | renal system vasculature development | 3 | 7.30E-04 |
| GO:0061440 | kidney vasculature development | 3 | 7.30E-04 |
| GO:0043502 | regulation of muscle adaptation | 4 | 7.73E-04 |
| GO:0045913 | positive regulation of carbohydrate metabolic process | 4 | 7.73E-04 |
| GO:0048660 | regulation of smooth muscle cell proliferation | 5 | 7.94E-04 |
| GO:0050433 | regulation of catecholamine secretion | 3 | 8.10E-04 |
| GO:0098869 | cellular oxidant detoxification | 4 | 8.67E-04 |
| GO:0048659 | smooth muscle cell proliferation | 5 | 8.93E-04 |
| GO:0048661 | positive regulation of smooth muscle cell proliferation | 4 | 9.69E-04 |
| GO:0043408 | regulation of MAPK cascade | 10 | 9.86E-04 |
| GO:1903530 | regulation of secretion by cell | 9 | 1.03E-03 |
| GO:0019216 | regulation of lipid metabolic process | 7 | 1.06E-03 |
| GO:0019915 | lipid storage | 4 | 1.08E-03 |
| GO:0050432 | catecholamine secretion | 3 | 1.08E-03 |
| GO:1902003 | regulation of amyloid-beta formation | 3 | 1.08E-03 |
| GO:0032147 | activation of protein kinase activity | 4 | 1.14E-03 |
| GO:0042311 | vasodilation | 3 | 1.19E-03 |
| GO:0042572 | retinol metabolic process | 3 | 1.19E-03 |
| GO:0045859 | regulation of protein kinase activity | 9 | 1.28E-03 |
| GO:0043410 | positive regulation of MAPK cascade | 8 | 1.30E-03 |
| GO:0045766 | positive regulation of angiogenesis | 5 | 1.38E-03 |
| GO:1904018 | positive regulation of vasculature development | 5 | 1.38E-03 |
| GO:0050708 | regulation of protein secretion | 6 | 1.40E-03 |
| GO:0023061 | signal release | 8 | 1.42E-03 |
| GO:0001822 | kidney development | 7 | 1.45E-03 |
| GO:0034329 | cell junction assembly | 8 | 1.45E-03 |
| GO:0007156 | homophilic cell adhesion via plasma membrane adhesion molecules | 4 | 1.46E-03 |
| GO:0070252 | actin-mediated cell contraction | 4 | 1.46E-03 |
| GO:0001938 | positive regulation of endothelial cell proliferation | 4 | 1.53E-03 |
| GO:0014812 | muscle cell migration | 4 | 1.53E-03 |
| GO:0014706 | striated muscle tissue development | 6 | 1.59E-03 |
| GO:0009306 | protein secretion | 7 | 1.61E-03 |
| GO:0035592 | establishment of protein localization to extracellular region | 7 | 1.61E-03 |
| GO:0002285 | lymphocyte activation involved in immune response | 5 | 1.64E-03 |
| GO:0030335 | positive regulation of cell migration | 9 | 1.64E-03 |
| GO:0051046 | regulation of secretion | 9 | 1.75E-03 |
| GO:0043691 | reverse cholesterol transport | 2 | 1.75E-03 |
| GO:0072110 | glomerular mesangial cell proliferation | 2 | 1.75E-03 |
| GO:1904238 | pericyte cell differentiation | 2 | 1.75E-03 |
| GO:0072001 | renal system development | 7 | 1.79E-03 |
| GO:0043114 | regulation of vascular permeability | 3 | 1.80E-03 |
| GO:0046460 | neutral lipid biosynthetic process | 3 | 1.80E-03 |
| GO:0046463 | acylglycerol biosynthetic process | 3 | 1.80E-03 |
| GO:1902991 | regulation of amyloid precursor protein catabolic process | 3 | 1.80E-03 |
| GO:0033674 | positive regulation of kinase activity | 7 | 1.83E-03 |
| GO:0071692 | protein localization to extracellular region | 7 | 1.83E-03 |
| GO:0043255 | regulation of carbohydrate biosynthetic process | 4 | 1.84E-03 |
| GO:1990748 | cellular detoxification | 4 | 1.92E-03 |
| GO:2000351 | regulation of endothelial cell apoptotic process | 3 | 1.94E-03 |
| GO:2000147 | positive regulation of cell motility | 9 | 2.01E-03 |
| GO:0043500 | muscle adaptation | 4 | 2.01E-03 |
| GO:0010817 | regulation of hormone levels | 8 | 2.08E-03 |
| GO:0034205 | amyloid-beta formation | 3 | 2.08E-03 |
| GO:0046890 | regulation of lipid biosynthetic process | 5 | 2.11E-03 |
| GO:0030277 | maintenance of gastrointestinal epithelium | 2 | 2.13E-03 |
| GO:0045428 | regulation of nitric oxide biosynthetic process | 3 | 2.24E-03 |
| GO:0045017 | glycerolipid biosynthetic process | 6 | 2.31E-03 |
| GO:0040017 | positive regulation of locomotion | 9 | 2.34E-03 |
| GO:0045765 | regulation of angiogenesis | 6 | 2.36E-03 |
| GO:0097237 | cellular response to toxic substance | 4 | 2.38E-03 |
| GO:0080164 | regulation of nitric oxide metabolic process | 3 | 2.40E-03 |
| GO:0006816 | calcium ion transport | 7 | 2.47E-03 |
| GO:1901342 | regulation of vasculature development | 6 | 2.48E-03 |
| GO:0032352 | positive regulation of hormone metabolic process | 2 | 2.54E-03 |
| GO:1901722 | regulation of cell proliferation involved in kidney development | 2 | 2.54E-03 |
| GO:0051937 | catecholamine transport | 3 | 2.57E-03 |
| GO:0072577 | endothelial cell apoptotic process | 3 | 2.57E-03 |
| GO:0006639 | acylglycerol metabolic process | 4 | 2.59E-03 |
| GO:0006109 | regulation of carbohydrate metabolic process | 5 | 2.60E-03 |
| GO:0006638 | neutral lipid metabolic process | 4 | 2.69E-03 |
| GO:1904036 | negative regulation of epithelial cell apoptotic process | 3 | 2.74E-03 |
| GO:0045216 | cell-cell junction organization | 5 | 2.92E-03 |
| GO:0046486 | glycerolipid metabolic process | 7 | 2.98E-03 |
| GO:0050746 | regulation of lipoprotein metabolic process | 2 | 3.00E-03 |
| GO:0061050 | regulation of cell growth involved in cardiac muscle cell development | 2 | 3.00E-03 |
| GO:0071071 | regulation of phospholipid biosynthetic process | 2 | 3.00E-03 |

| GO terms negatively enriched with age | | | |
|---|---|---|---|
| GO ID | GO description | Gene count (input = 160) | Adjusted p-value |
| GO:0098813 | nuclear chromosome segregation | 23 | 1.08E-09 |
| GO:0000280 | nuclear division | 26 | 1.08E-09 |
| GO:0048285 | organelle fission | 27 | 1.46E-09 |
| GO:0000819 | sister chromatid segregation | 19 | 1.55E-08 |
| GO:0007059 | chromosome segregation | 24 | 2.71E-08 |
| GO:0140014 | mitotic nuclear division | 19 | 2.47E-07 |
| GO:0000070 | mitotic sister chromatid segregation | 16 | 2.47E-07 |
| GO:0051783 | regulation of nuclear division | 14 | 2.47E-07 |
| GO:0007088 | regulation of mitotic nuclear division | 13 | 2.70E-07 |
| GO:0007093 | mitotic cell cycle checkpoint signaling | 14 | 2.70E-07 |
| GO:0033045 | regulation of sister chromatid segregation | 12 | 8.80E-07 |
| GO:0000075 | cell cycle checkpoint signaling | 15 | 1.77E-06 |
| GO:0045930 | negative regulation of mitotic cell cycle | 16 | 1.79E-06 |
| GO:0030071 | regulation of mitotic metaphase/anaphase transition | 11 | 1.82E-06 |
| GO:0010948 | negative regulation of cell cycle process | 18 | 1.82E-06 |
| GO:0007346 | regulation of mitotic cell cycle | 23 | 1.82E-06 |
| GO:1902099 | regulation of metaphase/anaphase transition of cell cycle | 11 | 1.82E-06 |
| GO:0045787 | positive regulation of cell cycle | 18 | 1.84E-06 |
| GO:0090068 | positive regulation of cell cycle process | 16 | 2.11E-06 |
| GO:0007091 | metaphase/anaphase transition of mitotic cell cycle | 11 | 2.23E-06 |
| GO:0044770 | cell cycle phase transition | 24 | 2.28E-06 |
| GO:0044784 | metaphase/anaphase transition of cell cycle | 11 | 2.28E-06 |
| GO:0045839 | negative regulation of mitotic nuclear division | 9 | 2.32E-06 |
| GO:1901987 | regulation of cell cycle phase transition | 21 | 2.40E-06 |
| GO:0010965 | regulation of mitotic sister chromatid separation | 9 | 3.54E-06 |
| GO:0051983 | regulation of chromosome segregation | 12 | 4.27E-06 |
| GO:0051784 | negative regulation of nuclear division | 9 | 4.50E-06 |
| GO:1901988 | negative regulation of cell cycle phase transition | 16 | 4.64E-06 |
| GO:0051306 | mitotic sister chromatid separation | 9 | 4.64E-06 |
| GO:0044772 | mitotic cell cycle phase transition | 21 | 4.64E-06 |
| GO:1901990 | regulation of mitotic cell cycle phase transition | 18 | 4.64E-06 |
| GO:0007094 | mitotic spindle assembly checkpoint signaling | 8 | 4.64E-06 |
| GO:0071173 | spindle assembly checkpoint signaling | 8 | 4.64E-06 |
| GO:0071174 | mitotic spindle checkpoint signaling | 8 | 4.64E-06 |
| GO:0045786 | negative regulation of cell cycle | 19 | 5.04E-06 |
| GO:0031577 | spindle checkpoint signaling | 8 | 5.29E-06 |
| GO:0033046 | negative regulation of sister chromatid segregation | 8 | 5.73E-06 |
| GO:0033048 | negative regulation of mitotic sister chromatid segregation | 8 | 5.73E-06 |
| GO:0045841 | negative regulation of mitotic metaphase/anaphase transition | 8 | 5.73E-06 |
| GO:2000816 | negative regulation of mitotic sister chromatid separation | 8 | 5.73E-06 |
| GO:0051985 | negative regulation of chromosome segregation | 8 | 6.38E-06 |
| GO:1902100 | negative regulation of metaphase/anaphase transition of cell cycle | 8 | 6.38E-06 |
| GO:1905819 | negative regulation of chromosome separation | 8 | 6.38E-06 |
| GO:1901991 | negative regulation of mitotic cell cycle phase transition | 13 | 1.01E-05 |
| GO:1905818 | regulation of chromosome separation | 9 | 1.27E-05 |
| GO:0033047 | regulation of mitotic sister chromatid segregation | 8 | 1.38E-05 |
| GO:0051304 | chromosome separation | 9 | 2.53E-05 |
| GO:0007052 | mitotic spindle organization | 11 | 2.69E-05 |
| GO:1902850 | microtubule cytoskeleton organization involved in mitosis | 12 | 3.78E-05 |
| GO:0140013 | meiotic nuclear division | 11 | 6.09E-05 |
| GO:2001251 | negative regulation of chromosome organization | 9 | 7.56E-05 |
| GO:0006261 | DNA-templated DNA replication | 11 | 1.56E-04 |
| GO:1903046 | meiotic cell cycle process | 11 | 1.74E-04 |
| GO:0007051 | spindle organization | 12 | 2.21E-04 |
| GO:0006260 | DNA replication | 14 | 2.55E-04 |
| GO:0006270 | DNA replication initiation | 6 | 2.62E-04 |
| GO:0033044 | regulation of chromosome organization | 13 | 2.97E-04 |
| GO:0007080 | mitotic metaphase chromosome alignment | 7 | 3.03E-04 |
| GO:0051321 | meiotic cell cycle | 12 | 6.34E-04 |
| GO:0051310 | metaphase chromosome alignment | 8 | 7.12E-04 |
| GO:0008608 | attachment of spindle microtubules to kinetochore | 6 | 9.13E-04 |
| GO:0061982 | meiosis I cell cycle process | 8 | 9.55E-04 |
| GO:0051303 | establishment of chromosome localization | 8 | 1.10E-03 |
| GO:0000076 | DNA replication checkpoint signaling | 4 | 1.66E-03 |
| GO:0050000 | chromosome localization | 8 | 1.93E-03 |
| GO:0044839 | cell cycle G2/M phase transition | 9 | 2.75E-03 |
| GO:0007127 | meiosis I | 7 | 4.71E-03 |
| GO:0045132 | meiotic chromosome segregation | 6 | 4.71E-03 |
| GO:0051315 | attachment of mitotic spindle microtubules to kinetochore | 4 | 4.80E-03 |
| GO:0044786 | cell cycle DNA replication | 5 | 5.40E-03 |
| GO:1901989 | positive regulation of cell cycle phase transition | 7 | 1.00E-02 |
| GO:0051785 | positive regulation of nuclear division | 5 | 1.03E-02 |
| GO:1902749 | regulation of cell cycle G2/M phase transition | 7 | 1.10E-02 |
| GO:0001525 | angiogenesis | 15 | 1.26E-02 |
| GO:0002040 | sprouting angiogenesis | 7 | 1.44E-02 |
| GO:0044774 | mitotic DNA integrity checkpoint signaling | 6 | 1.52E-02 |
| GO:2000105 | positive regulation of DNA-templated DNA replication | 3 | 1.75E-02 |
| GO:0010720 | positive regulation of cell development | 13 | 1.97E-02 |
| GO:0010639 | negative regulation of organelle organization | 12 | 2.61E-02 |
| GO:0000212 | meiotic spindle organization | 3 | 2.63E-02 |
| GO:0045840 | positive regulation of mitotic nuclear division | 4 | 3.11E-02 |
| GO:0010842 | retina layer formation | 3 | 3.12E-02 |
| GO:0031570 | DNA integrity checkpoint signaling | 7 | 3.12E-02 |
| GO:0061351 | neural precursor cell proliferation | 7 | 3.12E-02 |
| GO:0045740 | positive regulation of DNA replication | 4 | 3.30E-02 |
| GO:0010389 | regulation of G2/M transition of mitotic cell cycle | 6 | 3.44E-02 |
| GO:0000086 | G2/M transition of mitotic cell cycle | 7 | 3.46E-02 |
| GO:0033260 | nuclear DNA replication | 4 | 3.54E-02 |
| GO:0090231 | regulation of spindle checkpoint | 3 | 4.74E-02 |
| GO:0090266 | regulation of mitotic cell cycle spindle assembly checkpoint | 3 | 4.74E-02 |
| GO:1903504 | regulation of mitotic spindle checkpoint | 3 | 4.74E-02 |
| GO:2000177 | regulation of neural precursor cell proliferation | 5 | 4.87E-02 |
| GO:2000179 | positive regulation of neural precursor cell proliferation | 4 | 4.93E-02 |

Table S3. Significantly up- and downregulated GSEA terms with age in SHOCK primary dermal fibroblasts.

| Primary dermal fibroblasts | | | | | |
|---|---|---|---|---|---|
| GSEA Gene Set Database | Gene Set | Size | Normalized enrichment score (NES) | Nominal p-value (NOM p-val) | Adjusted p-value |
| c1: Hallmark | HALLMARK_TGF_BETA_SIGNALING | 52 | 2.18 | 5.53E-56 | 2.77E-54 |
| | HALLMARK_MYOGENESIS | 150 | 1.90 | 2.26E-47 | 5.65E-46 |
| | HALLMARK_COAGULATION | 96 | 1.89 | 2.31E-30 | 3.84E-29 |
| | HALLMARK_TNFA_SIGNALING_VIA_NFKB | 174 | 1.87 | 1.77E-08 | 2.21E-07 |
| | HALLMARK_HEME_METABOLISM | 159 | 1.79 | 3.00E-06 | 3.00E-05 |
| | HALLMARK_IL6_JAK_STAT3_SIGNALING | 62 | 1.75 | 3.74E-06 | 3.12E-05 |
| | HALLMARK_EPITHELIAL_MESENCHYMAL_TRANSITION | 194 | 1.71 | 4.40E-06 | 3.14E-05 |
| | HALLMARK_ANGIOGENESIS | 29 | 1.66 | 2.88E-05 | 1.60E-04 |
| | HALLMARK_BILE_ACID_METABOLISM | 80 | 1.65 | 2.78E-05 | 1.60E-04 |
| | HALLMARK_P53_PATHWAY | 179 | 1.59 | 5.97E-05 | 2.99E-04 |
| | HALLMARK_APOPTOSIS | 141 | 1.53 | 7.33E-05 | 3.33E-04 |
| | HALLMARK_HYPOXIA | 168 | 1.53 | 1.19E-04 | 4.97E-04 |
| | HALLMARK_COMPLEMENT | 149 | 1.49 | 1.78E-04 | 6.85E-04 |
| | HALLMARK_IL2_STAT5_SIGNALING | 157 | 1.43 | 1.41E-03 | 4.70E-03 |
| | HALLMARK_ADIPOGENESIS | 179 | 1.37 | 1.34E-03 | 4.70E-03 |
| | HALLMARK_MTORC1_SIGNALING | 189 | -1.35 | 2.65E-03 | 8.29E-03 |
| | HALLMARK_DNA_REPAIR | 146 | -1.80 | 3.05E-03 | 8.96E-03 |
| | HALLMARK_MYC_TARGETS_V2 | 56 | -1.91 | 3.86E-03 | 1.07E-02 |
| | HALLMARK_SPERMATOGENESIS | 76 | -1.93 | 4.90E-03 | 1.29E-02 |
| | HALLMARK_MITOTIC_SPINDLE | 197 | -2.02 | 6.49E-03 | 1.62E-02 |
| | HALLMARK_MYC_TARGETS_V1 | 193 | -3.00 | 8.39E-03 | 2.00E-02 |
| | HALLMARK_G2M_CHECKPOINT | 188 | -3.45 | 1.60E-02 | 3.65E-02 |
| | HALLMARK_E2F_TARGETS | 195 | -3.56 | 2.00E-02 | 4.35E-02 |
| c2: KEGG Medicus | KEGG_LEISHMANIA_INFECTION | 44 | 2.10 | 1.29E-05 | 2.70E-04 |
| | KEGG_STEROID_HORMONE_BIOSYNTHESIS | 17 | 1.93 | 5.95E-04 | 5.88E-03 |
| | KEGG_MEDICUS_REFERENCE_TLR1_2_4_NFKB_SIGNALING_PATHWAY | 18 | 1.81 | 3.39E-03 | 2.57E-02 |
| | KEGG_MEDICUS_REFERENCE_ITGA_B_FAK_CDC42_SIGNALING_PATHWAY | 24 | 1.78 | 3.39E-03 | 2.57E-02 |
| | KEGG_MEDICUS_REFERENCE_ITGA_B_RHOGAP_RHOA_SIGNALING_PATHWAY | 21 | 1.78 | 6.04E-03 | 3.80E-02 |
| | KEGG_MEDICUS_REFERENCE_RELN_VLDLR_PI3K_SIGNALING_PATHWAY | 15 | 1.76 | 7.59E-03 | 4.51E-02 |
| | KEGG_MEDICUS_REFERENCE_ITGA_B_FAK_RAC_SIGNALING_PATHWAY | 27 | 1.71 | 7.82E-03 | 4.56E-02 |
| | KEGG_HYPERTROPHIC_CARDIOMYOPATHY_HCM | 60 | 1.69 | 1.63E-03 | 1.47E-02 |
| | KEGG_LYSOSOME | 110 | 1.60 | 1.35E-03 | 1.24E-02 |
| | KEGG_TGF_BETA_SIGNALING_PATHWAY | 72 | 1.57 | 4.67E-03 | 3.31E-02 |
| | KEGG_FOCAL_ADHESION | 169 | 1.51 | 3.95E-03 | 2.91E-02 |
| | KEGG_CYTOKINE_CYTOKINE_RECEPTOR_INTERACTION | 120 | 1.51 | 5.58E-03 | 3.70E-02 |
| | KEGG_PROGESTERONE_MEDIATED_OOCYTE_MATURATION | 76 | -1.52 | 8.50E-03 | 4.89E-02 |
| | KEGG_PARKINSONS_DISEASE | 89 | -1.57 | 5.32E-03 | 3.58E-02 |
| | KEGG_MEDICUS_VARIANT_MUTATION_CAUSED_ABERRANT_SNCA_TO_ELECTRON_TRANSFER_IN_COMPLEX_I | 39 | -1.62 | 7.67E-03 | 4.51E-02 |
| | KEGG_MEDICUS_VARIANT_MUTATION_INACTIVATED_PINK1_TO_ELECTRON_TRANSFER_IN_COMPLEX_I | 39 | -1.64 | 5.67E-03 | 3.71E-02 |
| | KEGG_MEDICUS_VARIANT_MUTATION_CAUSED_ABERRANT_ABETA_TO_ELECTRON_TRANSFER_IN_COMPLEX_I | 44 | -1.68 | 7.08E-03 | 4.31E-02 |
| | KEGG_MEDICUS_REFERENCE_DNA_END_RESECTION_AND_RPA_LOADING | 14 | -1.69 | 7.25E-03 | 4.37E-02 |
| | KEGG_PURINE_METABOLISM | 127 | -1.70 | 5.45E-04 | 5.50E-03 |
| | KEGG_MEDICUS_REFERENCE_ELECTRON_TRANSFER_IN_COMPLEX_I | 38 | -1.71 | 5.33E-03 | 3.58E-02 |
| | KEGG_MEDICUS_REFERENCE_CORE_NER_REACTION | 28 | -1.74 | 5.94E-03 | 3.80E-02 |
| | KEGG_OOCYTE_MEIOSIS | 98 | -1.76 | 2.21E-04 | 2.57E-03 |
| | KEGG_RNA_DEGRADATION | 50 | -1.76 | 2.02E-03 | 1.73E-02 |
| | KEGG_MEDICUS_VARIANT_MUTATION_CAUSED_ABERRANT_HTT_TO_RETROGRADE_AXONAL_TRANSPORT | 29 | -1.76 | 3.81E-03 | 2.85E-02 |
| | KEGG_MEDICUS_REFERENCE_MITOCHONDRIAL_COMPLEX_UCP1_IN_THERMOGENESIS | 54 | -1.77 | 1.77E-03 | 1.57E-02 |
| | KEGG_MEDICUS_VARIANT_MUTATION_CAUSED_ABERRANT_TDP43_TO_ELECTRON_TRANSFER_IN_COMPLEX_I | 39 | -1.78 | 1.35E-03 | 1.24E-02 |
| | KEGG_MEDICUS_REFERENCE_CHOLESTEROL_BIOSYNTHESIS | 13 | -1.78 | 6.68E-03 | 4.12E-02 |
| | KEGG_MEDICUS_REFERENCE_RAD51_DSDNA_DESTABILIZATION | 10 | -1.78 | 5.25E-03 | 3.58E-02 |
| | KEGG_MEDICUS_VARIANT_MUTATION_CAUSED_ABERRANT_DCTN1_TO_RETROGRADE_AXONAL_TRANSPORT | 12 | -1.81 | 6.09E-03 | 3.80E-02 |
| | KEGG_MEDICUS_REFERENCE_GLYCOLYSIS | 19 | -1.81 | 4.46E-03 | 3.24E-02 |
| | KEGG_MEDICUS_REFERENCE_RETROGRADE_AXONAL_TRANSPORT | 28 | -1.81 | 2.73E-03 | 2.24E-02 |
| | KEGG_MEDICUS_REFERENCE_OKAZAKI_FRAGMENT_MATURATION | 11 | -1.82 | 5.94E-03 | 3.80E-02 |
| | KEGG_PYRIMIDINE_METABOLISM | 89 | -1.82 | 1.90E-04 | 2.49E-03 |
| | KEGG_MEDICUS_REFERENCE_DISASSEMBLY_OF_MCC | 17 | -1.83 | 3.33E-03 | 2.57E-02 |
| | KEGG_MEDICUS_VARIANT_AMPLIFIED_MYC_TO_CELL_CYCLE_G1_S | 11 | -1.83 | 5.11E-03 | 3.57E-02 |
| | KEGG_MEDICUS_REFERENCE_P16_CELL_CYCLE_G1_S | 10 | -1.84 | 3.18E-03 | 2.52E-02 |
| | KEGG_MEDICUS_ENV_FACTOR_ZN_TO_ANTEROGRADE_AXONAL_TRANSPORT | 13 | -1.86 | 2.25E-03 | 1.90E-02 |
| | KEGG_MEDICUS_PATHOGEN_SALMONELLA_SIFA_TO_MICROTUBULE_PLUS_END_DIRECTED_TRANSPORT | 18 | -1.87 | 4.51E-03 | 3.24E-02 |
| | KEGG_MEDICUS_VARIANT_MUTATION_CAUSED_ABERRANT_HTT_TO_ANTEROGRADE_AXONAL_TRANSPORT | 19 | -1.89 | 2.00E-03 | 1.73E-02 |
| | KEGG_MEDICUS_REFERENCE_WNT_SIGNALING_MODULATION_WNT_ACYLATION | 11 | -1.90 | 2.28E-03 | 1.90E-02 |
| | KEGG_MEDICUS_ENV_FACTOR_IRON_TO_ANTEROGRADE_AXONAL_TRANSPORT | 13 | -1.91 | 1.28E-03 | 1.22E-02 |
| | KEGG_MEDICUS_VARIANT_MUTATION_CAUSED_ABERRANT_SOD1_TO_26S_PROTEASOME_MEDIATED_PROTEIN_DEGRADATION | 39 | -1.91 | 2.88E-04 | 3.21E-03 |
| | KEGG_MEDICUS_VARIANT_MUTATION_CAUSED_ABERRANT_SNCA_TO_ANTEROGRADE_AXONAL_TRANSPORT | 18 | -1.92 | 2.95E-03 | 2.38E-02 |
| | KEGG_MEDICUS_VARIANT_MUTATION_INACTIVATED_UBQLN2_TO_26S_PROTEASOME_MEDIATED_PROTEIN_DEGRADATION | 38 | -1.94 | 2.99E-04 | 3.27E-03 |
| | KEGG_MEDICUS_REFERENCE_26S_PROTEASOME_MEDIATED_PROTEIN_DEGRADATION | 37 | -1.97 | 1.98E-04 | 2.50E-03 |
| | KEGG_MEDICUS_VARIANT_MUTATION_INACTIVATED_VCP_TO_26S_PROTEASOME_MEDIATED_PROTEIN_DEGRADATION | 38 | -1.97 | 2.01E-04 | 2.50E-03 |
| | KEGG_MEDICUS_REFERENCE_SPINDLE_ASSEMBLY_CHECKPOINT_SIGNALING | 20 | -1.98 | 1.15E-03 | 1.12E-02 |
| | KEGG_MEDICUS_REFERENCE_ANTEROGRADE_AXONAL_TRANSPORT | 17 | -2.00 | 3.41E-04 | 3.57E-03 |
| | KEGG_MEDICUS_REFERENCE_MICROTUBULE_RHOA_SIGNALING_PATHWAY | 15 | -2.01 | 3.29E-04 | 3.52E-03 |
| | KEGG_RIBOSOME | 85 | -2.02 | 1.74E-05 | 3.50E-04 |
| | KEGG_MEDICUS_REFERENCE_CONDENSIN_LOADING | 20 | -2.03 | 5.11E-04 | 5.25E-03 |
| | KEGG_MEDICUS_VARIANT_MUTATION_CAUSED_ABERRANT_ABETA_TO_ANTEROGRADE_AXONAL_TRANSPORT | 17 | -2.03 | 2.17E-04 | 2.57E-03 |
| | KEGG_MEDICUS_REFERENCE_ASSEMBLY_AND_TRAFFICKING_OF_TELOMERASE | 17 | -2.04 | 2.17E-04 | 2.57E-03 |
| | KEGG_MEDICUS_PATHOGEN_ESCHERICHIA_ESPG_TO_MICROTUBULE_RHOA_SIGNALING_PATHWAY | 11 | -2.04 | 2.59E-04 | 2.94E-03 |
| | KEGG_MEDICUS_VARIANT_SCRAPIE_CONFORMATION_PRPSC_TO_26S_PROTEASOME_MEDIATED_PROTEIN_DEGRADATION | 34 | -2.04 | 1.32E-04 | 1.87E-03 |
| | KEGG_NUCLEOTIDE_EXCISION_REPAIR | 43 | -2.06 | 5.63E-05 | 8.43E-04 |
| | KEGG_MEDICUS_REFERENCE_MICROTUBULE_DEPOLYMERIZATION | 17 | -2.07 | 1.39E-04 | 1.92E-03 |
| | KEGG_MEDICUS_VARIANT_MUTATION_CAUSED_ABERRANT_SNCA_TO_26S_PROTEASOME_MEDIATED_PROTEIN_DEGRADATION | 34 | -2.08 | 7.01E-05 | 1.02E-03 |
| | KEGG_MEDICUS_REFERENCE_TRANSLATION_INITIATION | 80 | -2.08 | 4.26E-06 | 9.71E-05 |
| | KEGG_BASE_EXCISION_REPAIR | 33 | -2.09 | 4.43E-05 | 7.03E-04 |
| | KEGG_MEDICUS_VARIANT_MUTATION_CAUSED_ABERRANT_HTT_TO_26S_PROTEASOME_MEDIATED_PROTEIN_DEGRADATION | 34 | -2.11 | 5.27E-05 | 8.12E-04 |
| | KEGG_MISMATCH_REPAIR | 22 | -2.11 | 1.51E-04 | 2.03E-03 |
| | KEGG_MEDICUS_VARIANT_MUTATION_CAUSED_ABERRANT_ABETA_TO_26S_PROTEASOME_MEDIATED_PROTEIN_DEGRADATION | 34 | -2.12 | 4.41E-05 | 7.03E-04 |
| | KEGG_MEDICUS_REFERENCE_HOMOLOGOUS_RECOMBINATION_IN_ICLR | 13 | -2.15 | 3.18E-05 | 5.56E-04 |
| | KEGG_MEDICUS_VARIANT_AMPLIFIED_MYC_TO_P27_CELL_CYCLE_G1_S | 15 | -2.15 | 3.39E-05 | 5.73E-04 |
| | KEGG_MEDICUS_REFERENCE_KINETOCHORE_FIBER_ORGANIZATION | 17 | -2.16 | 3.07E-05 | 5.55E-04 |
| | KEGG_MEDICUS_REFERENCE_P27_CELL_CYCLE_G1_S | 13 | -2.17 | 2.17E-05 | 4.22E-04 |
| | KEGG_HOMOLOGOUS_RECOMBINATION | 25 | -2.19 | 3.06E-05 | 5.55E-04 |
| | KEGG_MEDICUS_REFERENCE_MICROTUBULE_DEPOLYMERIZATION_AT_THE_MINUS_ENDS | 14 | -2.21 | 2.12E-06 | 5.83E-05 |
| | KEGG_MEDICUS_REFERENCE_KINETOCHORE_MICROTUBULE_ATTACHMENT | 14 | -2.22 | 1.70E-06 | 5.09E-05 |
| | KEGG_MEDICUS_PATHOGEN_EBV_EBNA3C_TO_P27_CELL_CYCLE_G1_S_N00264 | 10 | -2.23 | 1.75E-06 | 5.09E-05 |
| | KEGG_MEDICUS_REFERENCE_ORGANIZATION_OF_THE_INNER_KINETOCHORE | 15 | -2.25 | 3.57E-06 | 8.90E-05 |
| | KEGG_MEDICUS_REFERENCE_NUCLEAR_EXPORT_OF_MRNA | 28 | -2.27 | 3.05E-06 | 8.00E-05 |
| | KEGG_MEDICUS_REFERENCE_MICROTUBULE_NUCLEATION | 22 | -2.28 | 6.11E-06 | 1.33E-04 |
| | KEGG_MEDICUS_REFERENCE_FANCONI_ANEMIA_PATHWAY | 14 | -2.28 | 4.90E-07 | 1.71E-05 |
| | KEGG_PROTEASOME | 40 | -2.29 | 2.10E-07 | 7.85E-06 |
| | KEGG_MEDICUS_REFERENCE_MISMATCH_REPAIR | 26 | -2.29 | 3.83E-06 | 9.12E-05 |
| | KEGG_MEDICUS_REFERENCE_LONG_PATCH_BER | 17 | -2.34 | 6.04E-07 | 1.98E-05 |
| | KEGG_MEDICUS_REFERENCE_BRANCHING_MICROTUBULE_NUCLEATION | 27 | -2.43 | 5.77E-08 | 2.33E-06 |
| | KEGG_MEDICUS_REFERENCE_DEPHOSPHORYLATION_OF_KINETOCHORE | 12 | -2.48 | 1.24E-09 | 8.12E-08 |
| | KEGG_MEDICUS_REFERENCE_CENPE_INTERACTION_WITH_NDC80_COMPLEX | 12 | -2.48 | 1.19E-09 | 8.12E-08 |
| | KEGG_MEDICUS_REFERENCE_ORGANIZATION_OF_THE_OUTER_KINETOCHORE | 16 | -2.50 | 8.32E-09 | 3.96E-07 |
| | KEGG_MEDICUS_REFERENCE_HOMOLOGOUS_RECOMBINATION | 31 | -2.51 | 1.13E-08 | 4.95E-07 |
| | KEGG_MEDICUS_REFERENCE_TRAIP_DEPENDENT_REPLISOME_DISASSEMBLY | 15 | -2.51 | 3.76E-09 | 2.19E-07 |
| | KEGG_MEDICUS_REFERENCE_DNA_REPLICATION_TERMINATION | 18 | -2.54 | 7.10E-09 | 3.72E-07 |
| | KEGG_CELL_CYCLE | 119 | -2.57 | 8.63E-14 | 1.51E-11 |
| | KEGG_MEDICUS_REFERENCE_DNA_REPLICATION_LICENSING | 23 | -2.67 | 4.92E-11 | 4.30E-09 |
| | KEGG_DNA_REPLICATION | 35 | -2.72 | 2.45E-12 | 2.57E-10 |
| | KEGG_MEDICUS_REFERENCE_PRE_IC_FORMATION | 27 | -2.79 | 2.61E-13 | 3.42E-11 |
| | KEGG_SPLICEOSOME | 123 | -2.87 | 1.52E-19 | 7.95E-17 |
| | KEGG_MEDICUS_REFERENCE_ORIGIN_UNWINDING_AND_ELONGATION | 30 | -2.95 | 1.22E-16 | 3.21E-14 |
| c5: GO Biological Process | GOBP_ENDOSOME_ORGANIZATION | 91 | 2.36 | 2.34E-09 | 1.39E-07 |
| | GOBP_AMYLOID_BETA_CLEARANCE | 29 | 2.30 | 2.44E-06 | 7.76E-05 |
| | GOBP_AMYLOID_BETA_METABOLIC_PROCESS | 47 | 2.24 | 5.84E-07 | 2.32E-05 |

| | | | | |
|---|---|---|---|---|
| GOBP_AUTOPHAGOSOME_ORGANIZATION | 118 | 2.24 | 7.87E-09 | 4.43E-07 |
| GOBP_STEROID_HORMONE_SECRETION | 18 | 2.23 | 5.38E-06 | 1.55E-04 |
| GOBP_MACROAUTOPHAGY | 343 | 2.20 | 5.47E-14 | 5.93E-12 |
| GOBP_SMOOTH_MUSCLE_CELL_PROLIFERATION | 114 | 2.12 | 2.29E-07 | 1.00E-05 |
| GOBP_NEGATIVE_REGULATION_OF_AMYLOID_PRECURSOR_PROTEIN_CATABOLIC_PROCESS | 17 | 2.11 | 1.17E-04 | 2.08E-03 |
| GOBP_NUTRIENT_STORAGE | 69 | 2.11 | 2.39E-06 | 7.74E-05 |
| GOBP_REGULATION_OF_AUTOPHAGY | 324 | 2.11 | 1.97E-12 | 1.75E-10 |
| GOBP_CELLULAR_OXIDANT_DETOXIFICATION | 60 | 2.09 | 4.80E-06 | 1.41E-04 |
| GOBP_VACUOLE_ORGANIZATION | 217 | 2.09 | 1.83E-09 | 1.10E-07 |
| GOBP_DETOXIFICATION | 93 | 2.08 | 2.05E-06 | 6.74E-05 |
| GOBP_AMYLOID_BETA_FORMATION | 37 | 2.07 | 3.30E-05 | 7.18E-04 |
| GOBP_PHOSPHOLIPID_TRANSLOCATION | 42 | 2.07 | 2.07E-05 | 4.98E-04 |
| GOBP_PROTEIN_LOCALIZATION_TO_PHAGOPHORE_ASSEMBLY_SITE | 16 | 2.07 | 3.06E-04 | 4.43E-03 |
| GOBP_LIPID_LOCALIZATION | 295 | 2.07 | 4.07E-11 | 3.12E-09 |
| GOBP_REGULATION_OF_STEROID_HORMONE_SECRETION | 15 | 2.06 | 2.88E-04 | 4.21E-03 |
| GOBP_PROTEIN_LIPID_COMPLEX_ORGANIZATION | 30 | 2.05 | 7.51E-05 | 1.44E-03 |
| GOBP_POSITIVE_REGULATION_OF_SMOOTH_MUSCLE_CELL_PROLIFERATION | 64 | 2.04 | 1.19E-05 | 3.10E-04 |
| GOBP_HYDROGEN_PEROXIDE_BIOSYNTHETIC_PROCESS | 11 | 2.03 | 1.67E-04 | 2.78E-03 |
| GOBP_POSITIVE_REGULATION_OF_REACTIVE_OXYGEN_SPECIES_METABOLIC_PROCESS | 54 | 2.03 | 2.45E-05 | 5.64E-04 |
| GOBP_RESPONSE_TO_INTERLEUKIN_6 | 24 | 2.03 | 2.64E-04 | 3.92E-03 |
| GOBP_REGULATION_OF_RENAL_SYSTEM_PROCESS | 15 | 2.03 | 4.33E-04 | 5.71E-03 |
| GOBP_RENAL_SYSTEM_PROCESS_INVOLVED_IN_REGULATION_OF_SYSTEMIC_ARTERIAL_BLOOD_PRESSURE | 14 | 2.03 | 4.75E-04 | 6.17E-03 |
| GOBP_FRUCTOSE_6_PHOSPHATE_METABOLIC_PROCESS | 11 | 2.03 | 1.72E-04 | 2.83E-03 |
| GOBP_REACTIVE_OXYGEN_SPECIES_METABOLIC_PROCESS | 161 | 2.03 | 5.04E-08 | 2.41E-06 |
| GOBP_ENDOSOMAL_LUMEN_ACIDIFICATION | 13 | 2.03 | 3.57E-04 | 4.95E-03 |
| GOBP_NEGATIVE_REGULATION_OF_AMIDE_METABOLIC_PROCESS | 22 | 2.02 | 4.00E-04 | 5.34E-03 |
| GOBP_REGULATION_OF_LIPID_STORAGE | 36 | 2.02 | 8.60E-05 | 1.61E-03 |
| GOBP_NEUTRAL_LIPID_BIOSYNTHETIC_PROCESS | 36 | 2.02 | 8.72E-05 | 1.61E-03 |
| GOBP_MULTIVESICULAR_BODY_ORGANIZATION | 32 | 2.01 | 1.25E-04 | 2.19E-03 |
| GOBP_REGULATION_OF_PLASMA_LIPOPROTEIN_PARTICLE_LEVELS | 50 | 2.01 | 2.96E-05 | 6.60E-04 |
| GOBP_NEUTRAL_LIPID_METABOLIC_PROCESS | 86 | 2.01 | 5.08E-06 | 1.48E-04 |
| GOBP_POTASSIUM_ION_IMPORT_ACROSS_PLASMA_MEMBRANE | 21 | 2.00 | 3.44E-04 | 4.82E-03 |
| GOBP_REGULATION_OF_MEMBRANE_LIPID_DISTRIBUTION | 50 | 2.00 | 3.70E-05 | 7.95E-04 |
| GOBP_CORTICOSTEROID_HORMONE_SECRETION | 12 | 2.00 | 3.74E-04 | 5.06E-03 |
| GOBP_POSITIVE_REGULATION_OF_VASCULATURE_DEVELOPMENT | 127 | 1.99 | 1.59E-06 | 5.61E-05 |
| GOBP_MONOSACCHARIDE_TRANSMEMBRANE_TRANSPORT | 82 | 1.98 | 1.19E-05 | 3.10E-04 |
| GOBP_OSTEOCLAST_DIFFERENTIATION | 73 | 1.98 | 4.87E-05 | 1.02E-03 |
| GOBP_CELLULAR_RESPONSE_TO_TOXIC_SUBSTANCE | 86 | 1.98 | 8.61E-06 | 2.36E-04 |
| GOBP_EMBRYO_IMPLANTATION | 55 | 1.98 | 4.18E-05 | 8.86E-04 |
| GOBP_CELLULAR_RESPONSE_TO_GROWTH_HORMONE_STIMULUS | 21 | 1.98 | 4.81E-04 | 6.17E-03 |
| GOBP_POSITIVE_REGULATION_OF_NUCLEOTIDE_CATABOLIC_PROCESS | 18 | 1.97 | 5.58E-04 | 6.89E-03 |
| GOBP_REGULATION_OF_B_CELL_DIFFERENTIATION | 20 | 1.97 | 7.05E-04 | 8.37E-03 |
| GOBP_INORGANIC_ION_IMPORT_ACROSS_PLASMA_MEMBRANE | 63 | 1.97 | 4.79E-05 | 1.01E-03 |
| GOBP_ASTROCYTE_DEVELOPMENT | 28 | 1.97 | 3.08E-04 | 4.44E-03 |
| GOBP_HORMONE_METABOLIC_PROCESS | 133 | 1.96 | 2.67E-06 | 8.35E-05 |
| GOBP_RESPONSE_TO_OXYGEN_LEVELS | 264 | 1.96 | 2.14E-08 | 1.11E-06 |
| GOBP_RESPONSE_TO_PH | 23 | 1.95 | 8.52E-04 | 9.78E-03 |
| GOBP_NEGATIVE_REGULATION_OF_OSSIFICATION | 31 | 1.95 | 2.77E-04 | 4.09E-03 |
| GOBP_POSITIVE_REGULATION_OF_AUTOPHAGY | 144 | 1.95 | 1.22E-06 | 4.47E-05 |
| GOBP_RESPONSE_TO_ACTIVITY | 60 | 1.95 | 5.42E-05 | 1.12E-03 |
| GOBP_HETEROTYPIC_CELL_CELL_ADHESION | 34 | 1.95 | 2.05E-04 | 3.24E-03 |
| GOBP_MACROPHAGE_ACTIVATION | 69 | 1.95 | 6.36E-05 | 1.27E-03 |
| GOBP_RENAL_TUBULAR_SECRETION | 12 | 1.94 | 8.92E-04 | 1.00E-02 |
| GOBP_ACYLGLYCEROL_HOMEOSTASIS | 23 | 1.94 | 1.23E-03 | 1.27E-02 |
| GOBP_REGULATION_OF_LIPID_METABOLIC_PROCESS | 234 | 1.94 | 5.71E-08 | 2.70E-06 |
| GOBP_PEPTIDE_BIOSYNTHETIC_PROCESS | 23 | 1.94 | 1.23E-03 | 1.27E-02 |
| GOBP_BONE_CELL_DEVELOPMENT | 14 | 1.93 | 1.82E-03 | 1.69E-02 |
| GOBP_MIDDLE_EAR_MORPHOGENESIS | 20 | 1.93 | 1.13E-03 | 1.19E-02 |
| GOBP_SUPEROXIDE_METABOLIC_PROCESS | 49 | 1.92 | 1.24E-04 | 2.17E-03 |
| GOBP_RESPONSE_TO_TOXIC_SUBSTANCE | 163 | 1.92 | 1.54E-06 | 5.51E-05 |
| GOBP_REACTIVE_NITROGEN_SPECIES_METABOLIC_PROCESS | 67 | 1.91 | 1.49E-04 | 2.52E-03 |
| GOBP_REGULATION_OF_OSTEOCLAST_DIFFERENTIATION | 41 | 1.91 | 4.81E-04 | 6.17E-03 |
| GOBP_GOLGI_TO_ENDOSOME_TRANSPORT | 15 | 1.91 | 1.77E-03 | 1.66E-02 |
| GOBP_POSITIVE_REGULATION_OF_VACUOLE_ORGANIZATION | 23 | 1.91 | 1.65E-03 | 1.59E-02 |
| GOBP_NEUTRAL_AMINO_ACID_TRANSPORT | 40 | 1.90 | 1.28E-04 | 2.21E-03 |
| GOBP_REGULATION_OF_D_GLUCOSE_TRANSMEMBRANE_TRANSPORT | 57 | 1.90 | 8.27E-05 | 1.55E-03 |
| GOBP_REGULATION_OF_PH | 74 | 1.90 | 6.00E-05 | 1.22E-03 |
| GOBP_MULTICELLULAR_ORGANISMAL_LEVEL_IRON_ION_HOMEOSTASIS | 21 | 1.90 | 1.22E-03 | 1.26E-02 |
| GOBP_CIRCULATORY_SYSTEM_PROCESS | 402 | 1.90 | 5.53E-10 | 3.62E-08 |
| GOBP_CONNECTIVE_TISSUE_DEVELOPMENT | 229 | 1.89 | 2.31E-07 | 1.00E-05 |
| GOBP_ORGANIC_CATION_TRANSPORT | 73 | 1.89 | 2.03E-04 | 3.23E-03 |
| GOBP_VACUOLAR_ACIDIFICATION | 33 | 1.89 | 5.08E-04 | 6.44E-03 |
| GOBP_NEGATIVE_REGULATION_OF_LIPID_STORAGE | 16 | 1.89 | 2.90E-03 | 2.39E-02 |
| GOBP_PHOSPHOLIPID_TRANSPORT | 74 | 1.89 | 7.14E-05 | 1.40E-03 |
| GOBP_LATE_ENDOSOME_TO_LYSOSOME_TRANSPORT | 20 | 1.89 | 1.82E-03 | 1.69E-02 |
| GOBP_T_HELPER_17_CELL_DIFFERENTIATION | 27 | 1.89 | 9.17E-04 | 1.02E-02 |
| GOBP_MEMBRANE_FISSION | 44 | 1.89 | 3.31E-04 | 4.73E-03 |
| GOBP_POSITIVE_REGULATION_OF_SMOOTH_MUSCLE_CELL_MIGRATION | 31 | 1.89 | 9.04E-04 | 1.01E-02 |
| GOBP_TRANSMISSION_OF_NERVE_IMPULSE | 38 | 1.88 | 1.91E-04 | 3.09E-03 |
| GOBP_VESICLE_ORGANIZATION | 323 | 1.88 | 1.28E-08 | 6.85E-07 |
| GOBP_VASCULAR_TRANSPORT | 63 | 1.88 | 1.84E-04 | 2.99E-03 |
| GOBP_REGULATION_OF_HEART_CONTRACTION | 127 | 1.88 | 1.40E-05 | 3.57E-04 |
| GOBP_LATE_ENDOSOME_TO_VACUOLE_TRANSPORT_VIA_MULTIVESICULAR_BODY_SORTING_PATHWAY | 26 | 1.88 | 8.80E-04 | 9.95E-03 |
| GOBP_POSITIVE_REGULATION_OF_GLYCOPROTEIN_METABOLIC_PROCESS | 15 | 1.88 | 2.49E-03 | 2.16E-02 |
| GOBP_REGULATION_OF_MACROAUTOPHAGY | 177 | 1.88 | 2.03E-06 | 6.73E-05 |
| GOBP_LYTIC_VACUOLE_ORGANIZATION | 96 | 1.87 | 8.13E-05 | 1.54E-03 |
| GOBP_SYNAPTIC_VESICLE_MATURATION | 25 | 1.87 | 1.38E-03 | 1.38E-02 |
| GOBP_COLLAGEN_CATABOLIC_PROCESS | 30 | 1.87 | 7.21E-04 | 8.46E-03 |
| GOBP_INFLAMMATORY_RESPONSE_TO_WOUNDING | 20 | 1.87 | 2.16E-03 | 1.93E-02 |
| GOBP_BONE_DEVELOPMENT | 165 | 1.87 | 3.13E-06 | 9.57E-05 |
| GOBP_NUCLEOPHAGY | 22 | 1.86 | 2.59E-03 | 2.21E-02 |
| GOBP_CARBOHYDRATE_TRANSPORT | 92 | 1.86 | 9.17E-05 | 1.67E-03 |
| GOBP_ASTROCYTE_DIFFERENTIATION | 58 | 1.86 | 2.57E-04 | 3.83E-03 |
| GOBP_AUTOPHAGY_OF_MITOCHONDRION | 103 | 1.86 | 7.73E-05 | 1.48E-03 |
| GOBP_EXTRACELLULAR_MATRIX_DISASSEMBLY | 49 | 1.86 | 2.53E-04 | 3.81E-03 |
| GOBP_INTRACELLULAR_PH_REDUCTION | 46 | 1.86 | 4.81E-04 | 6.17E-03 |
| GOBP_REGULATION_OF_EXTRACELLULAR_MATRIX_ORGANIZATION | 54 | 1.86 | 4.78E-04 | 6.17E-03 |
| GOBP_SMAD_PROTEIN_SIGNAL_TRANSDUCTION | 73 | 1.85 | 3.13E-04 | 4.50E-03 |
| GOBP_ASTROCYTE_ACTIVATION | 14 | 1.85 | 4.75E-03 | 3.44E-02 |
| GOBP_REGULATION_OF_BLOOD_CIRCULATION | 160 | 1.85 | 9.04E-06 | 2.46E-04 |
| GOBP_GLYCOPHAGY | 12 | 1.85 | 2.65E-03 | 2.23E-02 |
| GOBP_AMINO_SUGAR_CATABOLIC_PROCESS | 12 | 1.85 | 2.69E-03 | 2.26E-02 |
| GOBP_GLIAL_CELL_ACTIVATION | 32 | 1.85 | 9.41E-04 | 1.04E-02 |
| GOBP_SYNAPTIC_VESICLE_LUMEN_ACIDIFICATION | 16 | 1.84 | 4.70E-03 | 3.43E-02 |
| GOBP_RESPONSE_TO_POTASSIUM_ION | 12 | 1.84 | 2.77E-03 | 2.31E-02 |
| GOBP_POSITIVE_REGULATION_OF_INTERLEUKIN_4_PRODUCTION | 13 | 1.84 | 2.86E-03 | 2.37E-02 |
| GOBP_VASODILATION | 37 | 1.84 | 7.19E-04 | 8.45E-03 |
| GOBP_POSITIVE_REGULATION_OF_D_GLUCOSE_TRANSMEMBRANE_TRANSPORT | 34 | 1.84 | 7.06E-04 | 8.37E-03 |
| GOBP_FOAM_CELL_DIFFERENTIATION | 22 | 1.84 | 3.44E-03 | 2.73E-02 |
| GOBP_VESICLE_TETHERING | 31 | 1.84 | 1.48E-03 | 1.46E-02 |
| GOBP_REGULATION_OF_LYSOSOMAL_LUMEN_PH | 29 | 1.83 | 2.26E-03 | 2.00E-02 |
| GOBP_HEART_PROCESS | 163 | 1.83 | 1.06E-05 | 2.88E-04 |
| GOBP_POSITIVE_REGULATION_OF_HORMONE_METABOLIC_PROCESS | 10 | 1.83 | 5.87E-03 | 3.93E-02 |
| GOBP_MIRNA_TRANSCRIPTION | 62 | 1.82 | 4.29E-04 | 5.69E-03 |
| GOBP_DIACYLGLYCEROL_METABOLIC_PROCESS | 19 | 1.82 | 2.64E-03 | 2.23E-02 |
| GOBP_BODY_MORPHOGENESIS | 44 | 1.82 | 6.07E-04 | 7.35E-03 |
| GOBP_MUSCLE_CELL_PROLIFERATION | 171 | 1.82 | 3.95E-06 | 1.19E-04 |
| GOBP_COLLAGEN_METABOLIC_PROCESS | 78 | 1.82 | 2.43E-04 | 3.70E-03 |
| GOBP_INTERLEUKIN_6_MEDIATED_SIGNALING_PATHWAY | 14 | 1.82 | 6.60E-03 | 4.26E-02 |
| GOBP_GOLGI_LUMEN_ACIDIFICATION | 13 | 1.82 | 3.92E-03 | 3.00E-02 |
| GOBP_LIPID_EXPORT_FROM_CELL | 32 | 1.82 | 1.31E-03 | 1.33E-02 |
| GOBP_PROTEIN_TRANSPORT_TO_VACUOLE_INVOLVED_IN_UBIQUITIN_DEPENDENT_PROTEIN_CATABOLIC_PROCESS_VIA_THE_MULTIVESICULAR_BODY_SORTING_PATHWAY | 16 | 1.82 | 5.82E-03 | 3.91E-02 |
| GOBP_REGULATION_OF_BLOOD_PRESSURE | 111 | 1.81 | 3.80E-05 | 8.11E-04 |

| | | | | |
|---|---|---|---|---|
| GOBP_REGULATION_OF_AMYLOID_BETA_CLEARANCE | 13 | 1.81 | 4.15E-03 | 3.12E-02 |
| GOBP_ENDOSOMAL_TRANSPORT | 248 | 1.81 | 1.67E-06 | 5.76E-05 |
| GOBP_REVERSE_CHOLESTEROL_TRANSPORT | 11 | 1.81 | 5.02E-03 | 3.59E-02 |
| GOBP_LYSOSOMAL_LUMEN_ACIDIFICATION | 24 | 1.81 | 2.60E-03 | 2.21E-02 |
| GOBP_REGULATION_OF_MIRNA_METABOLIC_PROCESS | 75 | 1.81 | 2.21E-04 | 3.42E-03 |
| GOBP_NEGATIVE_REGULATION_OF_MIRNA_METABOLIC_PROCESS | 25 | 1.81 | 3.01E-03 | 2.46E-02 |
| GOBP_CELLULAR_RESPONSE_TO_OXYGEN_RADICAL | 16 | 1.81 | 6.50E-03 | 4.22E-02 |
| GOBP_CELL_PROLIFERATION_INVOLVED_IN_KIDNEY_DEVELOPMENT | 17 | 1.81 | 4.84E-03 | 3.50E-02 |
| GOBP_REGULATION_OF_LEUKOCYTE_ADHESION_TO_VASCULAR_ENDOTHELIAL_CELL | 19 | 1.80 | 3.54E-03 | 2.79E-02 |
| GOBP_TRANSITION_METAL_ION_TRANSPORT | 73 | 1.80 | 5.55E-04 | 6.87E-03 |
| GOBP_PLASMA_LIPOPROTEIN_PARTICLE_CLEARANCE | 29 | 1.80 | 2.74E-03 | 2.29E-02 |
| GOBP_CELL_DIFFERENTIATION_INVOLVED_IN_METANEPHROS_DEVELOPMENT | 16 | 1.80 | 6.57E-03 | 4.25E-02 |
| GOBP_MITOPHAGY | 84 | 1.80 | 2.36E-04 | 3.62E-03 |
| GOBP_REGULATION_OF_EPITHELIAL_TO_MESENCHYMAL_TRANSITION | 85 | 1.80 | 2.07E-04 | 3.26E-03 |
| GOBP_IMPORT_ACROSS_PLASMA_MEMBRANE | 120 | 1.80 | 1.82E-04 | 2.96E-03 |
| GOBP_AUTOPHAGOSOME_MATURATION | 62 | 1.80 | 5.80E-04 | 7.08E-03 |
| GOBP_POSITIVE_REGULATION_OF_ENDOTHELIAL_CELL_PROLIFERATION | 77 | 1.80 | 3.88E-04 | 5.22E-03 |
| GOBP_INTRACELLULAR_GLUCOSE_HOMEOSTASIS | 112 | 1.80 | 1.68E-04 | 2.79E-03 |
| GOBP_VACUOLAR_TRANSPORT | 153 | 1.80 | 2.13E-05 | 5.04E-04 |
| GOBP_REGULATION_OF_AMIDE_METABOLIC_PROCESS | 58 | 1.79 | 5.63E-04 | 6.93E-03 |
| GOBP_TRIGLYCERIDE_METABOLIC_PROCESS | 65 | 1.79 | 3.43E-04 | 4.82E-03 |
| GOBP_SEX_DETERMINATION | 13 | 1.79 | 4.99E-03 | 3.58E-02 |
| GOBP_NEURON_CELLULAR_HOMEOSTASIS | 43 | 1.79 | 1.11E-03 | 1.18E-02 |
| GOBP_RESPONSE_TO_MUSCLE_ACTIVITY | 21 | 1.79 | 4.21E-03 | 3.16E-02 |
| GOBP_POSITIVE_REGULATION_OF_NUCLEOTIDE_METABOLIC_PROCESS | 28 | 1.79 | 2.42E-03 | 2.12E-02 |
| GOBP_VENTRAL_SPINAL_CORD_DEVELOPMENT | 25 | 1.78 | 3.61E-03 | 2.83E-02 |
| GOBP_OLEFINIC_COMPOUND_METABOLIC_PROCESS | 93 | 1.78 | 4.87E-04 | 6.18E-03 |
| GOBP_LIPID_HOMEOSTASIS | 119 | 1.78 | 1.77E-04 | 2.90E-03 |
| GOBP_RETINOL_METABOLIC_PROCESS | 27 | 1.78 | 2.52E-03 | 2.18E-02 |
| GOBP_NON_LYTIC_VIRAL_RELEASE | 13 | 1.78 | 5.75E-03 | 3.89E-02 |
| GOBP_VASCULAR_PROCESS_IN_CIRCULATORY_SYSTEM | 186 | 1.78 | 2.64E-05 | 6.01E-04 |
| GOBP_TRANSCYTOSIS | 13 | 1.78 | 5.75E-03 | 3.89E-02 |
| GOBP_ENDOPLASMIC_RETICULUM_CALCIUM_ION_HOMEOSTASIS | 23 | 1.78 | 5.90E-03 | 3.94E-02 |
| GOBP_NEGATIVE_REGULATION_OF_EPITHELIAL_TO_MESENCHYMAL_TRANSITION | 31 | 1.78 | 2.40E-03 | 2.11E-02 |
| GOBP_CELLULAR_RESPONSE_TO_CARBOHYDRATE_STIMULUS | 105 | 1.78 | 2.29E-04 | 3.53E-03 |
| GOBP_REGULATION_OF_GLYCOLYTIC_PROCESS | 42 | 1.78 | 1.01E-03 | 1.09E-02 |
| GOBP_RESPONSE_TO_BMP | 126 | 1.78 | 1.43E-04 | 2.43E-03 |
| GOBP_CARTILAGE_DEVELOPMENT | 159 | 1.78 | 3.69E-05 | 7.95E-04 |
| GOBP_REGULATION_OF_REACTIVE_OXYGEN_SPECIES_METABOLIC_PROCESS | 108 | 1.77 | 1.94E-04 | 3.12E-03 |
| GOBP_POSITIVE_REGULATION_OF_VASCULAR_ASSOCIATED_SMOOTH_MUSCLE_CELL_PROLIFERATION | 32 | 1.77 | 1.97E-03 | 1.79E-02 |
| GOBP_REGULATION_OF_GLYCOPROTEIN_METABOLIC_PROCESS | 31 | 1.77 | 2.55E-03 | 2.20E-02 |
| GOBP_SKELETAL_SYSTEM_MORPHOGENESIS | 181 | 1.77 | 1.89E-05 | 4.62E-04 |
| GOBP_LATE_ENDOSOME_TO_VACUOLE_TRANSPORT | 35 | 1.77 | 1.64E-03 | 1.58E-02 |
| GOBP_NEGATIVE_REGULATION_OF_CHONDROCYTE_DIFFERENTIATION | 19 | 1.77 | 4.44E-03 | 3.30E-02 |
| GOBP_REGULATION_OF_CALCIUM_ION_DEPENDENT_EXOCYTOSIS | 22 | 1.77 | 5.20E-03 | 3.67E-02 |
| GOBP_POSITIVE_REGULATION_OF_MACROAUTOPHAGY | 85 | 1.77 | 3.69E-04 | 5.03E-03 |
| GOBP_SKELETAL_SYSTEM_DEVELOPMENT | 410 | 1.77 | 4.36E-08 | 2.13E-06 |
| GOBP_RESPONSE_TO_PLATELET_DERIVED_GROWTH_FACTOR | 22 | 1.76 | 5.59E-03 | 3.85E-02 |
| GOBP_REGULATION_OF_WNT_SIGNALING_PATHWAY_PLANAR_CELL_POLARITY_PATHWAY | 13 | 1.76 | 7.20E-03 | 4.56E-02 |
| GOBP_PEROXISOME_ORGANIZATION | 35 | 1.76 | 1.81E-03 | 1.69E-02 |
| GOBP_EXTRACELLULAR_VESICLE_BIOGENESIS | 22 | 1.76 | 5.59E-03 | 3.85E-02 |
| GOBP_NEGATIVE_REGULATION_OF_OSTEOCLAST_DIFFERENTIATION | 20 | 1.76 | 6.57E-03 | 4.25E-02 |
| GOBP_DIGESTIVE_SYSTEM_PROCESS | 53 | 1.76 | 8.59E-04 | 9.84E-03 |
| GOBP_POSITIVE_REGULATION_OF_AUTOPHAGOSOME_ASSEMBLY | 19 | 1.75 | 5.15E-03 | 3.64E-02 |
| GOBP_ENDOMEMBRANE_SYSTEM_ORGANIZATION | 489 | 1.75 | 7.43E-09 | 4.29E-07 |
| GOBP_HEAD_MORPHOGENESIS | 36 | 1.75 | 2.55E-03 | 2.20E-02 |
| GOBP_PEROXISOMAL_TRANSPORT | 21 | 1.75 | 5.76E-03 | 3.89E-02 |
| GOBP_VIRION_ASSEMBLY | 33 | 1.75 | 2.85E-03 | 2.36E-02 |
| GOBP_REGULATION_OF_CHONDROCYTE_DIFFERENTIATION | 42 | 1.75 | 1.42E-03 | 1.42E-02 |
| GOBP_D_GLUCOSE_IMPORT | 57 | 1.74 | 6.71E-04 | 8.04E-03 |
| GOBP_MONOAMINE_TRANSPORT | 33 | 1.74 | 3.07E-03 | 2.49E-02 |
| GOBP_RENAL_FILTRATION | 22 | 1.74 | 6.76E-03 | 4.35E-02 |
| GOBP_STEROL_HOMEOSTASIS | 60 | 1.74 | 9.42E-04 | 1.04E-02 |
| GOBP_L_AMINO_ACID_TRANSPORT | 61 | 1.74 | 1.12E-03 | 1.18E-02 |
| GOBP_REGULATION_OF_D_GLUCOSE_IMPORT | 41 | 1.74 | 2.44E-03 | 2.13E-02 |
| GOBP_ENDODERM_FORMATION | 47 | 1.74 | 2.27E-03 | 2.01E-02 |
| GOBP_LUNG_CELL_DIFFERENTIATION | 21 | 1.74 | 6.46E-03 | 4.20E-02 |
| GOBP_REGULATION_OF_CARBOHYDRATE_CATABOLIC_PROCESS | 51 | 1.74 | 8.68E-04 | 9.89E-03 |
| GOBP_REGULATION_OF_VASCULATURE_DEVELOPMENT | 218 | 1.73 | 2.74E-05 | 6.21E-04 |
| GOBP_REGULATION_OF_TRIGLYCERIDE_METABOLIC_PROCESS | 29 | 1.73 | 5.96E-03 | 3.95E-02 |
| GOBP_MULTICELLULAR_ORGANISMAL_LEVEL_CHEMICAL_HOMEOSTASIS | 51 | 1.73 | 9.15E-04 | 1.02E-02 |
| GOBP_METANEPHRIC_NEPHRON_DEVELOPMENT | 29 | 1.73 | 6.04E-03 | 3.99E-02 |
| GOBP_POSITIVE_REGULATION_OF_MIRNA_METABOLIC_PROCESS | 53 | 1.73 | 1.29E-03 | 1.31E-02 |
| GOBP_NEGATIVE_REGULATION_OF_SMAD_PROTEIN_SIGNAL_TRANSDUCTION | 22 | 1.73 | 7.54E-03 | 4.71E-02 |
| GOBP_MULTI_MULTICELLULAR_ORGANISM_PROCESS | 152 | 1.73 | 1.00E-04 | 1.80E-03 |
| GOBP_POSITIVE_REGULATION_OF_INFLAMMATORY_RESPONSE | 96 | 1.73 | 7.36E-04 | 8.61E-03 |
| GOBP_REGULATION_OF_NITRIC_OXIDE_METABOLIC_PROCESS | 50 | 1.72 | 1.01E-03 | 1.09E-02 |
| GOBP_MYOBLAST_PROLIFERATION | 21 | 1.72 | 7.62E-03 | 4.74E-02 |
| GOBP_POSITIVE_REGULATION_OF_NITRIC_OXIDE_METABOLIC_PROCESS | 33 | 1.72 | 3.71E-03 | 2.86E-02 |
| GOBP_SMOOTH_MUSCLE_CELL_MIGRATION | 63 | 1.72 | 1.38E-03 | 1.38E-02 |
| GOBP_REGULATION_OF_HORMONE_METABOLIC_PROCESS | 28 | 1.72 | 4.20E-03 | 3.15E-02 |
| GOBP_POSITIVE_REGULATION_OF_EPITHELIAL_CELL_PROLIFERATION | 156 | 1.72 | 6.36E-05 | 1.27E-03 |
| GOBP_CARDIAC_CONDUCTION_SYSTEM_DEVELOPMENT | 26 | 1.72 | 5.63E-03 | 3.86E-02 |
| GOBP_REGULATION_OF_OSTEOBLAST_DIFFERENTIATION | 108 | 1.72 | 4.49E-04 | 5.87E-03 |
| GOBP_NEPHRON_DEVELOPMENT | 126 | 1.72 | 3.41E-04 | 4.82E-03 |
| GOBP_CELLULAR_RESPONSE_TO_AMYLOID_BETA | 33 | 1.71 | 3.98E-03 | 3.04E-02 |
| GOBP_REGULATION_OF_SYSTEM_PROCESS | 345 | 1.71 | 3.09E-06 | 9.54E-05 |
| GOBP_BONE_MORPHOGENESIS | 78 | 1.71 | 1.07E-03 | 1.15E-02 |
| GOBP_NEGATIVE_REGULATION_OF_NON_CANONICAL_NF_KAPPAB_SIGNAL_TRANSDUCTION | 18 | 1.71 | 7.92E-03 | 4.87E-02 |
| GOBP_PROXIMAL_DISTAL_PATTERN_FORMATION | 28 | 1.70 | 5.02E-03 | 3.59E-02 |
| GOBP_REGULATION_OF_MYELOID_LEUKOCYTE_DIFFERENTIATION | 84 | 1.70 | 1.11E-03 | 1.18E-02 |
| GOBP_RESPONSE_TO_AMYLOID_BETA | 38 | 1.70 | 1.82E-03 | 1.69E-02 |
| GOBP_SYNAPSE_MATURATION | 31 | 1.70 | 5.67E-03 | 3.86E-02 |
| GOBP_GLOMERULUS_DEVELOPMENT | 60 | 1.69 | 1.49E-03 | 1.46E-02 |
| GOBP_VIRAL_BUDDING | 24 | 1.69 | 7.48E-03 | 4.68E-02 |
| GOBP_REGULATION_OF_INFLAMMATORY_RESPONSE | 251 | 1.69 | 2.11E-05 | 5.01E-04 |
| GOBP_TERPENOID_METABOLIC_PROCESS | 53 | 1.69 | 1.88E-03 | 1.72E-02 |
| GOBP_RESPONSE_TO_WOUNDING | 412 | 1.69 | 5.36E-07 | 2.19E-05 |
| GOBP_NEUROMUSCULAR_PROCESS | 107 | 1.69 | 5.19E-04 | 6.54E-03 |
| GOBP_SUPEROXIDE_ANION_GENERATION | 26 | 1.69 | 7.14E-03 | 4.53E-02 |
| GOBP_LEUKOCYTE_ACTIVATION_INVOLVED_IN_INFLAMMATORY_RESPONSE | 30 | 1.69 | 5.03E-03 | 3.59E-02 |
| GOBP_NEGATIVE_REGULATION_OF_FAT_CELL_DIFFERENTIATION | 43 | 1.69 | 3.02E-03 | 2.46E-02 |
| GOBP_CYTOSOLIC_TRANSPORT | 120 | 1.68 | 1.07E-03 | 1.15E-02 |
| GOBP_ORGANIC_HYDROXY_COMPOUND_TRANSPORT | 163 | 1.68 | 1.94E-04 | 3.12E-03 |
| GOBP_NEGATIVE_REGULATION_OF_AUTOPHAGY | 83 | 1.68 | 8.85E-04 | 9.98E-03 |
| GOBP_REGULATION_OF_STEROID_METABOLIC_PROCESS | 77 | 1.68 | 1.64E-03 | 1.58E-02 |
| GOBP_FAT_CELL_DIFFERENTIATION | 197 | 1.68 | 5.44E-05 | 1.12E-03 |
| GOBP_LIPID_OXIDATION | 88 | 1.68 | 1.64E-03 | 1.58E-02 |
| GOBP_MULTIVESICULAR_BODY_SORTING_PATHWAY | 45 | 1.68 | 2.62E-03 | 2.22E-02 |
| GOBP_LIPID_MODIFICATION | 135 | 1.68 | 2.11E-04 | 3.30E-03 |
| GOBP_T_CELL_ACTIVATION_INVOLVED_IN_IMMUNE_RESPONSE | 79 | 1.68 | 1.72E-03 | 1.64E-02 |
| GOBP_POSITIVE_REGULATION_OF_SMALL_MOLECULE_METABOLIC_PROCESS | 108 | 1.68 | 7.16E-04 | 8.44E-03 |
| GOBP_WNT_SIGNALING_PATHWAY | 384 | 1.67 | 2.37E-06 | 7.74E-05 |
| GOBP_PLASMA_MEMBRANE_ORGANIZATION | 117 | 1.67 | 7.39E-04 | 8.62E-03 |
| GOBP_ENDODERM_DEVELOPMENT | 68 | 1.67 | 1.80E-03 | 1.68E-02 |
| GOBP_REGULATION_OF_B_CELL_ACTIVATION | 87 | 1.67 | 2.06E-03 | 1.86E-02 |
| GOBP_REGULATION_OF_HEART_RATE | 61 | 1.67 | 2.28E-03 | 2.01E-02 |
| GOBP_REGULATION_OF_AMYLOID_PRECURSOR_PROTEIN_CATABOLIC_PROCESS | 37 | 1.67 | 4.99E-03 | 3.58E-02 |
| GOBP_BODY_FLUID_SECRETION | 61 | 1.67 | 2.28E-03 | 2.01E-02 |
| GOBP_NEGATIVE_REGULATION_OF_CELL_DIFFERENTIATION | 462 | 1.67 | 5.06E-07 | 2.08E-05 |
| GOBP_POSITIVE_REGULATION_OF_SMALL_GTPASE_MEDIATED_SIGNAL_TRANSDUCTION | 57 | 1.67 | 1.77E-03 | 1.66E-02 |
| GOBP_POSITIVE_REGULATION_OF_ANIMAL_ORGAN_MORPHOGENESIS | 27 | 1.67 | 8.11E-03 | 4.97E-02 |
| GOBP_RENAL_SYSTEM_DEVELOPMENT | 269 | 1.67 | 2.47E-05 | 5.67E-04 |

| | | | | |
|---|---|---|---|---|
| GOBP_ACTIN_FILAMENT_BASED_MOVEMENT | 84 | 1.66 | 1.95E-03 | 1.78E-02 |
| GOBP_VESICLE_LOCALIZATION | 191 | 1.66 | 7.02E-05 | 1.38E-03 |
| GOBP_NEGATIVE_REGULATION_OF_MUSCLE_CELL_APOPTOTIC_PROCESS | 37 | 1.66 | 5.53E-03 | 3.84E-02 |
| GOBP_VASCULAR_ASSOCIATED_SMOOTH_MUSCLE_CELL_PROLIFERATION | 55 | 1.66 | 2.14E-03 | 1.91E-02 |
| GOBP_POSITIVE_REGULATION_OF_PHAGOCYTOSIS | 42 | 1.66 | 3.47E-03 | 2.74E-02 |
| GOBP_RESPONSE_TO_OXIDATIVE_STRESS | 320 | 1.66 | 1.09E-05 | 2.93E-04 |
| GOBP_MIRNA_METABOLIC_PROCESS | 87 | 1.66 | 2.45E-03 | 2.13E-02 |
| GOBP_EPITHELIAL_CELL_DIFFERENTIATION_INVOLVED_IN_KIDNEY_DEVELOPMENT | 38 | 1.66 | 2.65E-03 | 2.23E-02 |
| GOBP_CELL_CELL_ADHESION_VIA_PLASMA_MEMBRANE_ADHESION_MOLECULES | 122 | 1.66 | 6.01E-04 | 7.29E-03 |
| GOBP_METANEPHROS_DEVELOPMENT | 67 | 1.66 | 4.68E-03 | 3.42E-02 |
| GOBP_REGULATION_OF_LIPID_LOCALIZATION | 109 | 1.66 | 9.81E-04 | 1.07E-02 |
| GOBP_ENDOTHELIAL_CELL_PROLIFERATION | 124 | 1.66 | 5.80E-04 | 7.08E-03 |
| GOBP_POSITIVE_REGULATION_OF_LOCOMOTION | 441 | 1.66 | 1.36E-06 | 4.95E-05 |
| GOBP_RENAL_SYSTEM_PROCESS | 73 | 1.66 | 2.61E-03 | 2.22E-02 |
| GOBP_PLATELET_DERIVED_GROWTH_FACTOR_RECEPTOR_SIGNALING_PATHWAY | 54 | 1.65 | 5.77E-03 | 3.89E-02 |
| GOBP_MEMORY | 91 | 1.65 | 1.84E-03 | 1.70E-02 |
| GOBP_REGULATION_OF_LIPID_BIOSYNTHETIC_PROCESS | 136 | 1.65 | 8.52E-04 | 9.78E-03 |
| GOBP_REGULATION_OF_SMALL_MOLECULE_METABOLIC_PROCESS | 241 | 1.65 | 5.44E-05 | 1.12E-03 |
| GOBP_NEGATIVE_REGULATION_OF_SMOOTH_MUSCLE_CELL_PROLIFERATION | 43 | 1.65 | 4.09E-03 | 3.09E-02 |
| GOBP_PROTEIN_QUALITY_CONTROL_FOR_MISFOLDED_OR_INCOMPLETELY_SYNTHESIZED_PROTEINS | 28 | 1.65 | 8.15E-03 | 4.97E-02 |
| GOBP_REGULATION_OF_VACUOLE_ORGANIZATION | 58 | 1.65 | 4.11E-03 | 3.10E-02 |
| GOBP_SYNAPTIC_TRANSMISSION_GLUTAMATERGIC | 62 | 1.65 | 3.48E-03 | 2.75E-02 |
| GOBP_POSITIVE_REGULATION_OF_MIRNA_TRANSCRIPTION | 44 | 1.65 | 5.24E-03 | 3.69E-02 |
| GOBP_CELLULAR_RESPONSE_TO_OXIDATIVE_STRESS | 211 | 1.64 | 7.89E-05 | 1.50E-03 |
| GOBP_POST_EMBRYONIC_DEVELOPMENT | 72 | 1.64 | 2.94E-03 | 2.42E-02 |
| GOBP_POSITIVE_REGULATION_OF_B_CELL_ACTIVATION | 56 | 1.64 | 2.97E-03 | 2.44E-02 |
| GOBP_ACUTE_INFLAMMATORY_RESPONSE | 50 | 1.64 | 2.59E-03 | 2.21E-02 |
| GOBP_MONOATOMIC_ION_HOMEOSTASIS | 402 | 1.64 | 3.58E-06 | 1.08E-04 |
| GOBP_REGULATION_OF_SYSTEMIC_ARTERIAL_BLOOD_PRESSURE | 57 | 1.64 | 2.37E-03 | 2.08E-02 |
| GOBP_NEGATIVE_REGULATION_OF_CANONICAL_NF_KAPPAB_SIGNAL_TRANSDUCTION | 63 | 1.64 | 2.93E-03 | 2.42E-02 |
| GOBP_CELLULAR_RESPONSE_TO_OXYGEN_LEVELS | 143 | 1.64 | 4.78E-04 | 6.17E-03 |
| GOBP_ADULT_BEHAVIOR | 98 | 1.64 | 2.70E-03 | 2.26E-02 |
| GOBP_RESPONSE_TO_ENDOPLASMIC_RETICULUM_STRESS | 237 | 1.64 | 1.63E-04 | 2.74E-03 |
| GOBP_HEART_MORPHOGENESIS | 204 | 1.64 | 1.26E-04 | 2.19E-03 |
| GOBP_CELL_SURFACE_RECEPTOR_PROTEIN_SERINE_THREONINE_KINASE_SIGNALING_PATHWAY | 301 | 1.64 | 1.89E-05 | 4.62E-04 |
| GOBP_BONE_MINERALIZATION | 94 | 1.63 | 1.75E-03 | 1.65E-02 |
| GOBP_REGULATION_OF_RESPONSE_TO_ENDOPLASMIC_RETICULUM_STRESS | 82 | 1.63 | 1.96E-03 | 1.79E-02 |
| GOBP_REGULATION_OF_OSSIFICATION | 92 | 1.63 | 2.58E-03 | 2.21E-02 |
| GOBP_FACE_MORPHOGENESIS | 30 | 1.63 | 7.29E-03 | 4.59E-02 |
| GOBP_POSITIVE_REGULATION_OF_CATABOLIC_PROCESS | 462 | 1.63 | 1.58E-06 | 5.61E-05 |
| GOBP_HORMONE_BIOSYNTHETIC_PROCESS | 39 | 1.63 | 5.63E-03 | 3.86E-02 |
| GOBP_REGULATION_OF_SMAD_PROTEIN_SIGNAL_TRANSDUCTION | 56 | 1.63 | 3.35E-03 | 2.69E-02 |
| GOBP_PRIMARY_ALCOHOL_METABOLIC_PROCESS | 60 | 1.63 | 3.26E-03 | 2.63E-02 |
| GOBP_NEGATIVE_REGULATION_OF_INFLAMMATORY_RESPONSE | 103 | 1.63 | 2.06E-03 | 1.86E-02 |
| GOBP_GLIAL_CELL_DEVELOPMENT | 97 | 1.63 | 1.67E-03 | 1.60E-02 |
| GOBP_WOUND_HEALING | 307 | 1.63 | 1.47E-05 | 3.72E-04 |
| GOBP_PROTEIN_LOCALIZATION_TO_VACUOLE | 76 | 1.63 | 4.07E-03 | 3.09E-02 |
| GOBP_NEUROINFLAMMATORY_RESPONSE | 52 | 1.63 | 5.88E-03 | 3.93E-02 |
| GOBP_REGULATION_OF_TISSUE_REMODELING | 52 | 1.62 | 5.95E-03 | 3.95E-02 |
| GOBP_ACTION_POTENTIAL | 98 | 1.62 | 3.24E-03 | 2.61E-02 |
| GOBP_NEGATIVE_REGULATION_OF_LIPID_METABOLIC_PROCESS | 66 | 1.62 | 3.66E-03 | 2.84E-02 |
| GOBP_DIGESTION | 60 | 1.62 | 3.65E-03 | 2.84E-02 |
| GOBP_REGULATION_OF_STEROID_BIOSYNTHETIC_PROCESS | 56 | 1.62 | 3.62E-03 | 2.83E-02 |
| GOBP_OSSIFICATION | 325 | 1.62 | 3.18E-05 | 6.99E-04 |
| GOBP_VESICLE_BUDDING_FROM_MEMBRANE | 81 | 1.62 | 3.03E-03 | 2.46E-02 |
| GOBP_LYMPHOCYTE_ACTIVATION_INVOLVED_IN_IMMUNE_RESPONSE | 139 | 1.62 | 9.95E-04 | 1.08E-02 |
| GOBP_POSITIVE_REGULATION_OF_EPITHELIAL_CELL_MIGRATION | 43 | 1.62 | 5.23E-03 | 3.68E-02 |
| GOBP_LEUKOCYTE_MIGRATION | 234 | 1.62 | 7.36E-05 | 1.42E-03 |
| GOBP_VESICLE_TARGETING | 60 | 1.62 | 3.88E-03 | 2.98E-02 |
| GOBP_MYELOID_LEUKOCYTE_ACTIVATION | 147 | 1.62 | 8.26E-04 | 9.54E-03 |
| GOBP_RETROGRADE_TRANSPORT_ENDOSOME_TO_GOLGI | 86 | 1.62 | 2.56E-03 | 2.20E-02 |
| GOBP_AMYLOID_PRECURSOR_PROTEIN_CATABOLIC_PROCESS | 50 | 1.62 | 3.58E-03 | 2.81E-02 |
| GOBP_LOCOMOTORY_BEHAVIOR | 142 | 1.61 | 7.11E-04 | 8.40E-03 |
| GOBP_ORGANOPHOSPHATE_ESTER_TRANSPORT | 125 | 1.61 | 1.41E-03 | 1.41E-02 |
| GOBP_POSITIVE_REGULATION_OF_LIPID_METABOLIC_PROCESS | 92 | 1.61 | 3.24E-03 | 2.61E-02 |
| GOBP_REGULATION_OF_CARTILAGE_DEVELOPMENT | 56 | 1.61 | 4.27E-03 | 3.20E-02 |
| GOBP_METAL_ION_TRANSPORT | 496 | 1.61 | 3.14E-06 | 9.57E-05 |
| GOBP_REACTIVE_OXYGEN_SPECIES_BIOSYNTHETIC_PROCESS | 39 | 1.61 | 8.19E-03 | 4.99E-02 |
| GOBP_MYELOID_LEUKOCYTE_DIFFERENTIATION | 174 | 1.60 | 5.50E-04 | 6.82E-03 |
| GOBP_RELEASE_OF_CYTOCHROME_C_FROM_MITOCHONDRIA | 49 | 1.60 | 5.73E-03 | 3.89E-02 |
| GOBP_RESPONSE_TO_REACTIVE_OXYGEN_SPECIES | 159 | 1.60 | 6.62E-04 | 7.96E-03 |
| GOBP_BIOLOGICAL_PROCESS_INVOLVED_IN_INTERACTION_WITH_HOST | 126 | 1.60 | 1.56E-03 | 1.52E-02 |
| GOBP_MESENCHYMAL_CELL_PROLIFERATION | 40 | 1.60 | 6.07E-03 | 4.00E-02 |
| GOBP_INTRACELLULAR_IRON_ION_HOMEOSTASIS | 61 | 1.60 | 4.74E-03 | 3.44E-02 |
| GOBP_ADULT_LOCOMOTORY_BEHAVIOR | 63 | 1.60 | 4.58E-03 | 3.38E-02 |
| GOBP_AMINO_SUGAR_METABOLIC_PROCESS | 34 | 1.60 | 7.46E-03 | 4.68E-02 |
| GOBP_CELLULAR_RESPONSE_TO_CHEMICAL_STRESS | 271 | 1.60 | 8.67E-05 | 1.61E-03 |
| GOBP_NEUROTRANSMITTER_TRANSPORT | 138 | 1.60 | 1.59E-03 | 1.54E-02 |
| GOBP_ENDOCHONDRAL_BONE_MORPHOGENESIS | 50 | 1.60 | 4.14E-03 | 3.12E-02 |
| GOBP_REGULATION_OF_SYNAPTIC_TRANSMISSION_GLUTAMATERGIC | 40 | 1.60 | 6.22E-03 | 4.07E-02 |
| GOBP_KIDNEY_MORPHOGENESIS | 76 | 1.59 | 6.38E-03 | 4.16E-02 |
| GOBP_MESONEPHROS_DEVELOPMENT | 80 | 1.59 | 7.10E-03 | 4.52E-02 |
| GOBP_RECEPTOR_MEDIATED_ENDOCYTOSIS | 204 | 1.59 | 3.53E-04 | 4.92E-03 |
| GOBP_ACTIN_MEDIATED_CELL_CONTRACTION | 68 | 1.59 | 5.16E-03 | 3.64E-02 |
| GOBP_INORGANIC_ION_HOMEOSTASIS | 342 | 1.59 | 6.44E-05 | 1.28E-03 |
| GOBP_CELLULAR_RESPONSE_TO_STARVATION | 157 | 1.58 | 9.77E-04 | 1.07E-02 |
| GOBP_NEGATIVE_REGULATION_OF_CELL_ADHESION | 202 | 1.58 | 2.03E-04 | 3.23E-03 |
| GOBP_RESPONSE_TO_PEPTIDE_HORMONE | 332 | 1.58 | 8.00E-05 | 1.52E-03 |
| GOBP_REGULATION_OF_CARBOHYDRATE_METABOLIC_PROCESS | 135 | 1.57 | 1.08E-03 | 1.16E-02 |
| GOBP_VESICLE_MEDIATED_TRANSPORT_IN_SYNAPSE | 202 | 1.57 | 2.46E-04 | 3.73E-03 |
| GOBP_CELL_ACTIVATION_INVOLVED_IN_IMMUNE_RESPONSE | 190 | 1.57 | 4.26E-04 | 5.66E-03 |
| GOBP_CELLULAR_RESPONSE_TO_NUTRIENT_LEVELS | 212 | 1.57 | 3.93E-04 | 5.28E-03 |
| GOBP_REGULATION_OF_TUBE_SIZE | 98 | 1.57 | 6.67E-03 | 4.30E-02 |
| GOBP_REGULATION_OF_HORMONE_LEVELS | 325 | 1.57 | 1.28E-04 | 2.21E-03 |
| GOBP_POTASSIUM_ION_TRANSPORT | 120 | 1.57 | 4.94E-03 | 3.55E-02 |
| GOBP_POSITIVE_REGULATION_OF_LIPID_LOCALIZATION | 68 | 1.57 | 6.36E-03 | 4.15E-02 |
| GOBP_AMINO_ACID_TRANSPORT | 106 | 1.57 | 4.05E-03 | 3.08E-02 |
| GOBP_NEPHRON_EPITHELIUM_DEVELOPMENT | 92 | 1.57 | 5.42E-03 | 3.79E-02 |
| GOBP_T_CELL_DIFFERENTIATION_INVOLVED_IN_IMMUNE_RESPONSE | 62 | 1.56 | 6.92E-03 | 4.43E-02 |
| GOBP_CELLULAR_RESPONSE_TO_PEPTIDE_HORMONE_STIMULUS | 252 | 1.56 | 2.70E-04 | 4.00E-03 |
| GOBP_CELLULAR_RESPONSE_TO_HORMONE_STIMULUS | 496 | 1.56 | 1.17E-05 | 3.08E-04 |
| GOBP_BIOLOGICAL_PROCESS_INVOLVED_IN_SYMBIOTIC_INTERACTION | 144 | 1.56 | 1.75E-03 | 1.65E-02 |
| GOBP_MUSCLE_CELL_MIGRATION | 77 | 1.56 | 7.54E-03 | 4.71E-02 |
| GOBP_AMYLOID_PRECURSOR_PROTEIN_METABOLIC_PROCESS | 63 | 1.56 | 7.82E-03 | 4.82E-02 |
| GOBP_VASCULOGENESIS | 68 | 1.55 | 7.34E-03 | 4.61E-02 |
| GOBP_RESPONSE_TO_HYDROGEN_PEROXIDE | 84 | 1.55 | 7.66E-03 | 4.75E-02 |
| GOBP_RESPONSE_TO_STARVATION | 183 | 1.55 | 1.00E-03 | 1.09E-02 |
| GOBP_CHONDROCYTE_DIFFERENTIATION | 96 | 1.55 | 7.37E-03 | 4.63E-02 |
| GOBP_STEROID_METABOLIC_PROCESS | 210 | 1.55 | 9.34E-04 | 1.04E-02 |
| GOBP_INTRACELLULAR_MONOATOMIC_ION_HOMEOSTASIS | 347 | 1.55 | 6.29E-05 | 1.26E-03 |
| GOBP_FORMATION_OF_PRIMARY_GERM_LAYER | 104 | 1.55 | 3.81E-03 | 2.93E-02 |
| GOBP_MONOATOMIC_CATION_TRANSMEMBRANE_TRANSPORT | 498 | 1.55 | 1.13E-05 | 3.02E-04 |
| GOBP_MESENCHYME_DEVELOPMENT | 249 | 1.55 | 2.21E-04 | 3.42E-03 |
| GOBP_CARBOHYDRATE_HOMEOSTASIS | 176 | 1.54 | 1.34E-03 | 1.35E-02 |
| GOBP_INORGANIC_ANION_TRANSPORT | 91 | 1.54 | 7.29E-03 | 4.59E-02 |
| GOBP_NEGATIVE_REGULATION_OF_PROTEIN_CONTAINING_COMPLEX_ASSEMBLY | 115 | 1.54 | 4.63E-03 | 3.40E-02 |
| GOBP_CANONICAL_WNT_SIGNALING_PATHWAY | 255 | 1.54 | 2.43E-04 | 3.70E-03 |
| GOBP_REGULATION_OF_FAT_CELL_DIFFERENTIATION | 107 | 1.54 | 2.82E-03 | 2.34E-02 |
| GOBP_PROTEIN_LOCALIZATION_TO_EXTRACELLULAR_REGION | 252 | 1.54 | 4.59E-04 | 5.98E-03 |
| GOBP_RESPONSE_TO_INTERLEUKIN_1 | 89 | 1.54 | 3.64E-03 | 2.84E-02 |
| GOBP_RESPONSE_TO_MECHANICAL_STIMULUS | 161 | 1.54 | 1.98E-03 | 1.80E-02 |
| GOBP_TRANSFORMING_GROWTH_FACTOR_BETA_RECEPTOR_SIGNALING_PATHWAY | 184 | 1.53 | 8.72E-04 | 9.91E-03 |
| GOBP_REGULATION_OF_CANONICAL_WNT_SIGNALING_PATHWAY | 210 | 1.53 | 1.28E-03 | 1.30E-02 |

| | | | | |
|---|---|---|---|---|
| GOBP_CELLULAR_RESPONSE_TO_INTERLEUKIN_1 | 71 | 1.53 | 6.75E-03 | 4.35E-02 |
| GOBP_ESTABLISHMENT_OF_PROTEIN_LOCALIZATION_TO_VACUOLE | 57 | 1.53 | 7.23E-03 | 4.57E-02 |
| GOBP_FIBROBLAST_PROLIFERATION | 99 | 1.53 | 6.19E-03 | 4.06E-02 |
| GOBP_CELL_SURFACE_RECEPTOR_PROTEIN_TYROSINE_KINASE_SIGNALING_PATHWAY | 491 | 1.53 | 2.45E-05 | 5.64E-04 |
| GOBP_BIOMINERAL_TISSUE_DEVELOPMENT | 120 | 1.52 | 7.07E-03 | 4.51E-02 |
| GOBP_HEART_DEVELOPMENT | 469 | 1.52 | 2.87E-05 | 6.44E-04 |
| GOBP_NEGATIVE_REGULATION_OF_WNT_SIGNALING_PATHWAY | 136 | 1.52 | 4.65E-03 | 3.41E-02 |
| GOBP_RESPONSE_TO_NUTRIENT_LEVELS | 383 | 1.52 | 1.36E-04 | 2.32E-03 |
| GOBP_RESPONSE_TO_FIBROBLAST_GROWTH_FACTOR | 86 | 1.52 | 8.02E-03 | 4.93E-02 |
| GOBP_SENSORY_ORGAN_DEVELOPMENT | 416 | 1.51 | 8.73E-05 | 1.61E-03 |
| GOBP_APOPTOTIC_MITOCHONDRIAL_CHANGES | 97 | 1.51 | 8.12E-03 | 4.97E-02 |
| GOBP_MYELOID_LEUKOCYTE_MIGRATION | 136 | 1.51 | 5.62E-03 | 3.86E-02 |
| GOBP_CELLULAR_RESPONSE_TO_METAL_ION | 126 | 1.51 | 4.77E-03 | 3.45E-02 |
| GOBP_KIDNEY_EPITHELIUM_DEVELOPMENT | 117 | 1.50 | 6.87E-03 | 4.41E-02 |
| GOBP_REGULATION_OF_CELL_MORPHOGENESIS | 196 | 1.50 | 1.81E-03 | 1.69E-02 |
| GOBP_INTRACELLULAR_RECEPTOR_SIGNALING_PATHWAY | 317 | 1.50 | 5.48E-04 | 6.82E-03 |
| GOBP_REGULATION_OF_WNT_SIGNALING_PATHWAY | 270 | 1.50 | 9.30E-04 | 1.03E-02 |
| GOBP_SENSORY_PERCEPTION_OF_MECHANICAL_STIMULUS | 107 | 1.50 | 4.54E-03 | 3.36E-02 |
| GOBP_POSITIVE_REGULATION_OF_SECRETION | 212 | 1.50 | 1.21E-03 | 1.26E-02 |
| GOBP_FATTY_ACID_METABOLIC_PROCESS | 277 | 1.50 | 9.58E-04 | 1.05E-02 |
| GOBP_EPITHELIAL_TO_MESENCHYMAL_TRANSITION | 144 | 1.49 | 3.89E-03 | 2.98E-02 |
| GOBP_GLYCEROLIPID_METABOLIC_PROCESS | 286 | 1.49 | 2.99E-04 | 4.34E-03 |
| GOBP_EMBRYONIC_SKELETAL_SYSTEM_DEVELOPMENT | 111 | 1.49 | 4.71E-03 | 3.43E-02 |
| GOBP_RESPONSE_TO_XENOBIOTIC_STIMULUS | 272 | 1.49 | 1.31E-03 | 1.33E-02 |
| GOBP_NEGATIVE_REGULATION_OF_CELL_DEVELOPMENT | 187 | 1.49 | 1.72E-03 | 1.64E-02 |
| GOBP_RESPONSE_TO_TRANSFORMING_GROWTH_FACTOR_BETA | 233 | 1.49 | 1.18E-03 | 1.23E-02 |
| GOBP_LYSOSOMAL_TRANSPORT | 118 | 1.48 | 5.65E-03 | 3.86E-02 |
| GOBP_MONONUCLEAR_CELL_MIGRATION | 134 | 1.48 | 7.24E-03 | 4.57E-02 |
| GOBP_ORGANIC_ANION_TRANSPORT | 265 | 1.48 | 8.68E-04 | 9.89E-03 |
| GOBP_EPITHELIAL_CELL_PROLIFERATION | 339 | 1.48 | 3.87E-04 | 5.22E-03 |
| GOBP_RESPONSE_TO_CARBOHYDRATE | 153 | 1.47 | 4.68E-03 | 3.42E-02 |
| GOBP_EMBRYONIC_MORPHOGENESIS | 473 | 1.47 | 2.90E-04 | 4.23E-03 |
| GOBP_MUSCLE_CELL_DIFFERENTIATION | 313 | 1.47 | 9.92E-04 | 1.08E-02 |
| GOBP_REGULATION_OF_CELLULAR_RESPONSE_TO_GROWTH_FACTOR_STIMULUS | 256 | 1.46 | 2.04E-03 | 1.85E-02 |
| GOBP_SODIUM_ION_TRANSPORT | 131 | 1.46 | 7.56E-03 | 4.71E-02 |
| GOBP_EAR_DEVELOPMENT | 153 | 1.46 | 6.01E-03 | 3.98E-02 |
| GOBP_NEGATIVE_REGULATION_OF_CELL_GROWTH | 131 | 1.46 | 7.64E-03 | 4.74E-02 |
| GOBP_PROTEIN_LOCALIZATION_TO_PLASMA_MEMBRANE | 239 | 1.46 | 1.81E-03 | 1.69E-02 |
| GOBP_GOLGI_VESICLE_TRANSPORT | 281 | 1.46 | 1.88E-03 | 1.73E-02 |
| GOBP_CANONICAL_NF_KAPPAB_SIGNAL_TRANSDUCTION | 269 | 1.45 | 1.57E-03 | 1.53E-02 |
| GOBP_CELL_SUBSTRATE_ADHESION | 280 | 1.45 | 2.07E-03 | 1.86E-02 |
| GOBP_OSTEOBLAST_DIFFERENTIATION | 189 | 1.45 | 4.56E-03 | 3.36E-02 |
| GOBP_REGULATION_OF_BODY_FLUID_LEVELS | 247 | 1.45 | 1.46E-03 | 1.45E-02 |
| GOBP_TISSUE_HOMEOSTASIS | 183 | 1.45 | 4.56E-03 | 3.36E-02 |
| GOBP_ERK1_AND_ERK2_CASCADE | 212 | 1.45 | 3.40E-03 | 2.71E-02 |
| GOBP_POSITIVE_REGULATION_OF_CELL_ADHESION | 344 | 1.45 | 6.09E-04 | 7.35E-03 |
| GOBP_GLYCEROLIPID_BIOSYNTHETIC_PROCESS | 198 | 1.45 | 5.66E-03 | 3.86E-02 |
| GOBP_REGULATION_OF_TRANSMEMBRANE_RECEPTOR_PROTEIN_SERINE_THREONINE_KINASE_SIGNALING_PATHWAY | 220 | 1.44 | 4.33E-03 | 3.23E-02 |
| GOBP_MONOCARBOXYLIC_ACID_METABOLIC_PROCESS | 435 | 1.44 | 4.85E-04 | 6.17E-03 |
| GOBP_POSITIVE_REGULATION_OF_CELL_PROJECTION_ORGANIZATION | 285 | 1.44 | 2.60E-03 | 2.21E-02 |
| GOBP_REGULATION_OF_SECRETION | 397 | 1.44 | 5.33E-04 | 6.69E-03 |
| GOBP_CELL_MATRIX_ADHESION | 189 | 1.44 | 5.46E-03 | 3.81E-02 |
| GOBP_BLOOD_VESSEL_MORPHOGENESIS | 490 | 1.44 | 2.82E-04 | 4.14E-03 |
| GOBP_CARDIAC_MUSCLE_TISSUE_DEVELOPMENT | 178 | 1.43 | 5.15E-03 | 3.64E-02 |
| GOBP_REGULATION_OF_EPITHELIAL_CELL_PROLIFERATION | 280 | 1.43 | 3.37E-03 | 2.70E-02 |
| GOBP_REGULATION_OF_CELL_SUBSTRATE_ADHESION | 172 | 1.42 | 8.09E-03 | 4.96E-02 |
| GOBP_LIPID_CATABOLIC_PROCESS | 220 | 1.42 | 6.27E-03 | 4.10E-02 |
| GOBP_REGULATION_OF_IMMUNE_EFFECTOR_PROCESS | 240 | 1.42 | 4.63E-03 | 3.40E-02 |
| GOBP_ORGANIC_ACID_TRANSPORT | 204 | 1.41 | 5.52E-03 | 3.84E-02 |
| GOBP_REGULATION_OF_DEVELOPMENTAL_GROWTH | 237 | 1.41 | 7.81E-03 | 4.82E-02 |
| GOBP_ADAPTIVE_IMMUNE_RESPONSE_BASED_ON_SOMATIC_RECOMBINATION_OF_IMMUNE_RECEPTORS_BUILT_FROM_IMMUNOGLOBULIN_SUPERFAMILY_DOMAINS | 204 | 1.41 | 5.76E-03 | 3.89E-02 |
| GOBP_POSITIVE_REGULATION_OF_PROGRAMMED_CELL_DEATH | 429 | 1.40 | 5.48E-04 | 6.82E-03 |
| GOBP_SENSORY_SYSTEM_DEVELOPMENT | 296 | 1.40 | 4.92E-03 | 3.55E-02 |
| GOBP_SIGNAL_RELEASE | 321 | 1.40 | 3.39E-03 | 2.71E-02 |
| GOBP_POSITIVE_REGULATION_OF_PHOSPHORUS_METABOLIC_PROCESS | 252 | 1.40 | 6.12E-03 | 4.02E-02 |
| GOBP_MUSCLE_TISSUE_DEVELOPMENT | 322 | 1.40 | 3.43E-03 | 2.73E-02 |
| GOBP_CELL_JUNCTION_ASSEMBLY | 361 | 1.39 | 2.42E-03 | 2.12E-02 |
| GOBP_REGULATION_OF_VESICLE_MEDIATED_TRANSPORT | 396 | 1.39 | 2.01E-03 | 1.82E-02 |
| GOBP_REGULATION_OF_NEURON_PROJECTION_DEVELOPMENT | 344 | 1.38 | 2.24E-03 | 1.99E-02 |
| GOBP_IMMUNE_EFFECTOR_PROCESS | 389 | 1.37 | 5.04E-03 | 3.59E-02 |
| GOBP_AMIDE_METABOLIC_PROCESS | 341 | 1.37 | 5.72E-03 | 3.89E-02 |
| GOBP_EPITHELIAL_CELL_DIFFERENTIATION | 465 | 1.37 | 1.32E-03 | 1.33E-02 |
| GOBP_GLYCOPROTEIN_METABOLIC_PROCESS | 301 | 1.37 | 5.67E-03 | 3.86E-02 |
| GOBP_CELLULAR_RESPONSE_TO_NITROGEN_COMPOUND | 472 | 1.36 | 3.29E-03 | 2.65E-02 |
| GOBP_REGULATION_OF_GROWTH | 460 | 1.35 | 4.33E-03 | 3.23E-02 |
| GOBP_POSITIVE_REGULATION_OF_MAPK_CASCADE | 316 | 1.35 | 6.08E-03 | 4.01E-02 |
| GOBP_CELLULAR_RESPONSE_TO_LIPID | 452 | 1.35 | 2.79E-03 | 2.32E-02 |
| GOBP_CELL_MORPHOGENESIS_INVOLVED_IN_NEURON_DIFFERENTIATION | 445 | 1.34 | 4.86E-03 | 3.51E-02 |
| GOBP_MUSCLE_SYSTEM_PROCESS | 290 | 1.33 | 7.62E-03 | 4.74E-02 |
| GOBP_BEHAVIOR | 440 | 1.32 | 3.66E-03 | 2.84E-02 |
| GOBP_NEGATIVE_REGULATION_OF_CELL_POPULATION_PROLIFERATION | 494 | 1.32 | 2.59E-03 | 2.21E-02 |
| GOBP_TISSUE_MORPHOGENESIS | 476 | 1.32 | 6.97E-03 | 4.45E-02 |
| GOBP_POSITIVE_REGULATION_OF_RESPONSE_TO_EXTERNAL_STIMULUS | 453 | 1.31 | 5.26E-03 | 3.69E-02 |
| GOBP_GAMETE_GENERATION | 484 | -1.33 | 5.45E-03 | 3.81E-02 |
| GOBP_FOREBRAIN_DEVELOPMENT | 295 | -1.44 | 1.77E-03 | 1.66E-02 |
| GOBP_REGULATION_OF_TRANSFERASE_ACTIVITY | 261 | -1.46 | 3.54E-03 | 2.79E-02 |
| GOBP_NUCLEIC_ACID_CATABOLIC_PROCESS | 300 | -1.46 | 1.27E-03 | 1.30E-02 |
| GOBP_REGULATION_OF_CELL_CYCLE_G1_S_PHASE_TRANSITION | 159 | -1.47 | 8.04E-03 | 4.93E-02 |
| GOBP_POSITIVE_REGULATION_OF_ORGANELLE_ORGANIZATION | 413 | -1.51 | 7.19E-05 | 1.40E-03 |
| GOBP_NUCLEOBASE_CONTAINING_COMPOUND_TRANSPORT | 195 | -1.52 | 1.89E-03 | 1.73E-02 |
| GOBP_CELLULAR_RESPIRATION | 178 | -1.54 | 1.66E-03 | 1.59E-02 |
| GOBP_NUCLEAR_TRANSPORT | 302 | -1.56 | 3.65E-04 | 5.00E-03 |
| GOBP_REGULATION_OF_DNA_BIOSYNTHETIC_PROCESS | 80 | -1.57 | 7.17E-03 | 4.54E-02 |
| GOBP_FEMALE_GAMETE_GENERATION | 111 | -1.57 | 3.01E-03 | 2.46E-02 |
| GOBP_ESTABLISHMENT_OF_ORGANELLE_LOCALIZATION | 407 | -1.59 | 3.16E-05 | 6.99E-04 |
| GOBP_NEGATIVE_REGULATION_OF_ORGANELLE_ORGANIZATION | 304 | -1.60 | 7.22E-05 | 1.40E-03 |
| GOBP_EPIGENETIC_REGULATION_OF_GENE_EXPRESSION | 214 | -1.61 | 3.67E-04 | 5.02E-03 |
| GOBP_MITOCHONDRIAL_RNA_METABOLIC_PROCESS | 46 | -1.64 | 5.86E-03 | 3.93E-02 |
| GOBP_MITOTIC_DNA_INTEGRITY_CHECKPOINT_SIGNALING | 68 | -1.64 | 4.54E-03 | 3.36E-02 |
| GOBP_POSITIVE_REGULATION_OF_DNA_BIOSYNTHETIC_PROCESS | 44 | -1.64 | 5.54E-03 | 3.84E-02 |
| GOBP_REGULATION_OF_STEM_CELL_POPULATION_MAINTENANCE | 60 | -1.65 | 3.84E-03 | 2.96E-02 |
| GOBP_MICROTUBULE_DEPOLYMERIZATION | 43 | -1.65 | 5.55E-03 | 3.84E-02 |
| GOBP_POSITIVE_REGULATION_OF_DNA_REPAIR | 121 | -1.65 | 1.25E-03 | 1.29E-02 |
| GOBP_POSTREPLICATION_REPAIR | 34 | -1.65 | 5.87E-03 | 3.93E-02 |
| GOBP_NEGATIVE_REGULATION_OF_CELL_CYCLE_G2_M_PHASE_TRANSITION | 62 | -1.66 | 3.22E-03 | 2.61E-02 |
| GOBP_TRNA_METABOLIC_PROCESS | 173 | -1.67 | 3.53E-04 | 4.92E-03 |
| GOBP_REGULATION_OF_MICROTUBULE_CYTOSKELETON_ORGANIZATION | 148 | -1.67 | 3.71E-04 | 5.05E-03 |
| GOBP_RESPONSE_TO_IONIZING_RADIATION | 126 | -1.68 | 5.20E-04 | 6.54E-03 |
| GOBP_REGULATION_OF_SPINDLE_ORGANIZATION | 45 | -1.68 | 4.74E-03 | 3.44E-02 |
| GOBP_REGULATION_OF_CENTROSOME_CYCLE | 52 | -1.68 | 5.09E-03 | 3.62E-02 |
| GOBP_CYTOSKELETON_DEPENDENT_CYTOKINESIS | 101 | -1.68 | 1.16E-03 | 1.22E-02 |
| GOBP_NEGATIVE_REGULATION_OF_GENE_EXPRESSION_EPIGENETIC | 141 | -1.68 | 5.35E-04 | 6.70E-03 |
| GOBP_MALE_MEIOTIC_NUCLEAR_DIVISION | 28 | -1.68 | 7.90E-03 | 4.87E-02 |
| GOBP_POSITIVE_REGULATION_OF_RNA_SPLICING | 35 | -1.69 | 5.94E-03 | 3.95E-02 |
| GOBP_MICROTUBULE_NUCLEATION | 46 | -1.69 | 3.68E-03 | 2.85E-02 |
| GOBP_NEGATIVE_REGULATION_OF_DNA_METABOLIC_PROCESS | 102 | -1.70 | 1.16E-03 | 1.22E-02 |
| GOBP_POSITIVE_REGULATION_OF_NUCLEAR_DIVISION | 43 | -1.71 | 3.69E-03 | 2.85E-02 |
| GOBP_REGULATION_OF_MEIOTIC_NUCLEAR_DIVISION | 23 | -1.71 | 6.21E-03 | 4.07E-02 |
| GOBP_RIBOSOMAL_SMALL_SUBUNIT_BIOGENESIS | 100 | -1.71 | 7.88E-04 | 9.14E-03 |
| GOBP_REGULATION_OF_CENTROSOME_DUPLICATION | 43 | -1.71 | 3.58E-03 | 2.81E-02 |
| GOBP_NEGATIVE_REGULATION_OF_RNA_CATABOLIC_PROCESS | 71 | -1.71 | 1.20E-03 | 1.25E-02 |
| GOBP_RNA_5_END_PROCESSING | 34 | -1.71 | 4.05E-03 | 3.08E-02 |
| GOBP_POSITIVE_REGULATION_OF_DOUBLE_STRAND_BREAK_REPAIR | 83 | -1.71 | 8.90E-04 | 1.00E-02 |

| | | | | |
|---|---|---|---|---|
| GOBP_MITOCHONDRIAL_RESPIRATORY_CHAIN_COMPLEX_ASSEMBLY | 78 | -1.71 | 1.86E-03 | 1.71E-02 |
| GOBP_MITOTIC_CYTOKINESIS | 89 | -1.71 | 8.06E-04 | 9.33E-03 |
| GOBP_POSITIVE_REGULATION_OF_MITOTIC_NUCLEAR_DIVISION | 33 | -1.72 | 5.59E-03 | 3.85E-02 |
| GOBP_REGULATION_OF_NUCLEOTIDE_EXCISION_REPAIR | 28 | -1.72 | 5.25E-03 | 3.69E-02 |
| GOBP_ESTABLISHMENT_OF_RNA_LOCALIZATION | 143 | -1.72 | 3.34E-04 | 4.76E-03 |
| GOBP_TRANSLESION_SYNTHESIS | 24 | -1.73 | 7.10E-03 | 4.52E-02 |
| GOBP_NEGATIVE_REGULATION_OF_MRNA_CATABOLIC_PROCESS | 59 | -1.73 | 1.80E-03 | 1.68E-02 |
| GOBP_RIBOSOMAL_LARGE_SUBUNIT_BIOGENESIS | 59 | -1.73 | 1.67E-03 | 1.60E-02 |
| GOBP_SIGNAL_TRANSDUCTION_IN_RESPONSE_TO_DNA_DAMAGE | 177 | -1.73 | 1.28E-04 | 2.21E-03 |
| GOBP_REGULATION_OF_PINOCYTOSIS | 12 | -1.74 | 6.53E-03 | 4.24E-02 |
| GOBP_MICROTUBULE_POLYMERIZATION | 93 | -1.74 | 1.36E-03 | 1.37E-02 |
| GOBP_POSITIVE_REGULATION_OF_TELOMERE_MAINTENANCE | 70 | -1.75 | 1.11E-03 | 1.18E-02 |
| GOBP_NEGATIVE_REGULATION_OF_DNA_REPAIR | 31 | -1.75 | 5.79E-03 | 3.90E-02 |
| GOBP_POSITIVE_REGULATION_OF_DOUBLE_STRAND_BREAK_REPAIR_VIA_HOMOLOGOUS_RECOMBINATION | 38 | -1.76 | 1.51E-03 | 1.48E-02 |
| GOBP_FOREBRAIN_GENERATION_OF_NEURONS | 59 | -1.76 | 1.18E-03 | 1.23E-02 |
| GOBP_CELL_DIFFERENTIATION_INVOLVED_IN_EMBRYONIC_PLACENTA_DEVELOPMENT | 17 | -1.76 | 6.85E-03 | 4.40E-02 |
| GOBP_OXIDATIVE_PHOSPHORYLATION | 105 | -1.76 | 6.92E-04 | 8.24E-03 |
| GOBP_TELOMERE_MAINTENANCE_IN_RESPONSE_TO_DNA_DAMAGE | 30 | -1.76 | 4.09E-03 | 3.09E-02 |
| GOBP_REGULATION_OF_UBIQUITIN_PROTEIN_TRANSFERASE_ACTIVITY | 25 | -1.76 | 7.08E-03 | 4.52E-02 |
| GOBP_HOMOLOGOUS_CHROMOSOME_PAIRING_AT_MEIOSIS | 28 | -1.76 | 3.45E-03 | 2.74E-02 |
| GOBP_ELECTRON_TRANSPORT_CHAIN | 102 | -1.77 | 3.64E-04 | 5.00E-03 |
| GOBP_REGULATION_OF_ALTERNATIVE_MRNA_SPLICING_VIA_SPLICEOSOME | 46 | -1.77 | 1.59E-03 | 1.54E-02 |
| GOBP_ESTABLISHMENT_OF_SPINDLE_ORIENTATION | 42 | -1.78 | 1.29E-03 | 1.31E-02 |
| GOBP_MACROMOLECULE_METHYLATION | 100 | -1.78 | 2.53E-04 | 3.81E-03 |
| GOBP_POSITIVE_REGULATION_OF_DNA_METABOLIC_PROCESS | 245 | -1.78 | 5.12E-06 | 1.49E-04 |
| GOBP_POSITIVE_REGULATION_OF_CARDIOCYTE_DIFFERENTIATION | 11 | -1.78 | 7.67E-03 | 4.75E-02 |
| GOBP_CELLULAR_SENESCENCE | 91 | -1.78 | 2.54E-04 | 3.81E-03 |
| GOBP_MRNA_EXPORT_FROM_NUCLEUS | 62 | -1.78 | 7.71E-04 | 8.96E-03 |
| GOBP_POSITIVE_REGULATION_OF_MEIOTIC_CELL_CYCLE | 16 | -1.79 | 5.49E-03 | 3.83E-02 |
| GOBP_OOCYTE_MATURATION | 22 | -1.79 | 5.09E-03 | 3.62E-02 |
| GOBP_REGULATION_OF_DNA_RECOMBINATION | 106 | -1.79 | 2.84E-04 | 4.16E-03 |
| GOBP_MISMATCH_REPAIR | 32 | -1.79 | 2.76E-03 | 2.31E-02 |
| GOBP_REGULATION_OF_MRNA_METABOLIC_PROCESS | 252 | -1.80 | 1.76E-06 | 5.93E-05 |
| GOBP_REGULATION_OF_HAIR_FOLLICLE_DEVELOPMENT | 15 | -1.80 | 4.41E-03 | 3.28E-02 |
| GOBP_CYTOPLASMIC_TRANSLATION | 160 | -1.80 | 3.00E-05 | 6.67E-04 |
| GOBP_SPINDLE_LOCALIZATION | 58 | -1.80 | 1.47E-03 | 1.45E-02 |
| GOBP_NUCLEOSOME_ORGANIZATION | 64 | -1.80 | 5.88E-04 | 7.15E-03 |
| GOBP_SINGLE_STRANDED_VIRAL_RNA_REPLICATION_VIA_DOUBLE_STRANDED_DNA_INTERMEDIATE | 11 | -1.80 | 6.03E-03 | 3.99E-02 |
| GOBP_MRNA_TRANSPORT | 117 | -1.82 | 9.28E-05 | 1.68E-03 |
| GOBP_NUCLEAR_EXPORT | 153 | -1.82 | 3.20E-05 | 7.00E-04 |
| GOBP_NEGATIVE_REGULATION_OF_INTRINSIC_APOPTOTIC_SIGNALING_PATHWAY_IN_RESPONSE_TO_DNA_DAMAGE_BY_P53_CLASS_MEDIATOR | 15 | -1.83 | 3.44E-03 | 2.73E-02 |
| GOBP_RRNA_TRANSCRIPTION | 35 | -1.83 | 1.13E-03 | 1.19E-02 |
| GOBP_DNA_SYNTHESIS_INVOLVED_IN_DNA_REPAIR | 36 | -1.83 | 1.40E-03 | 1.39E-02 |
| GOBP_SPINDLE_MIDZONE_ASSEMBLY | 13 | -1.83 | 2.98E-03 | 2.44E-02 |
| GOBP_PROTEIN_LOCALIZATION_TO_CELL_CORTEX | 11 | -1.83 | 4.03E-03 | 3.07E-02 |
| GOBP_MITOCHONDRIAL_ELECTRON_TRANSPORT_CYTOCHROME_C_TO_OXYGEN | 14 | -1.84 | 5.92E-03 | 3.95E-02 |
| GOBP_AEROBIC_RESPIRATION | 144 | -1.84 | 4.19E-05 | 8.86E-04 |
| GOBP_REGULATION_OF_MICROTUBULE_BASED_PROCESS | 213 | -1.84 | 1.63E-06 | 5.71E-05 |
| GOBP_DNA_BIOSYNTHETIC_PROCESS | 142 | -1.85 | 2.39E-05 | 5.59E-04 |
| GOBP_RNA_EXPORT_FROM_NUCLEUS | 80 | -1.85 | 2.30E-04 | 3.53E-03 |
| GOBP_RNA_TEMPLATED_DNA_BIOSYNTHETIC_PROCESS | 56 | -1.85 | 2.55E-04 | 3.81E-03 |
| GOBP_POSITIVE_REGULATION_OF_CENTRIOLE_REPLICATION | 10 | -1.85 | 3.04E-03 | 2.47E-02 |
| GOBP_REGULATION_OF_UBIQUITIN_PROTEIN_LIGASE_ACTIVITY | 16 | -1.85 | 2.48E-03 | 2.16E-02 |
| GOBP_CYTOKINESIS | 153 | -1.85 | 1.59E-05 | 3.97E-04 |
| GOBP_RNA_CAPPING | 13 | -1.86 | 2.12E-03 | 1.90E-02 |
| GOBP_REGULATION_OF_TELOMERE_MAINTENANCE | 102 | -1.86 | 6.80E-05 | 1.34E-03 |
| GOBP_REGULATION_OF_CYCLIN_DEPENDENT_PROTEIN_SERINE_THREONINE_KINASE_ACTIVITY | 32 | -1.86 | 1.47E-03 | 1.45E-02 |
| GOBP_REGULATION_OF_TELOMERE_MAINTENANCE_VIA_TELOMERASE | 43 | -1.86 | 4.46E-04 | 5.86E-03 |
| GOBP_REGULATION_OF_TELOMERASE_RNA_LOCALIZATION_TO_CAJAL_BODY | 12 | -1.87 | 1.55E-03 | 1.51E-02 |
| GOBP_POSITIVE_REGULATION_OF_PROTEIN_LOCALIZATION_TO_CAJAL_BODY | 10 | -1.87 | 2.50E-03 | 2.16E-02 |
| GOBP_POSITIVE_REGULATION_OF_CELL_CYCLE_G1_S_PHASE_TRANSITION | 50 | -1.87 | 3.94E-04 | 5.28E-03 |
| GOBP_MRNA_CIS_SPLICING_VIA_SPLICEOSOME | 29 | -1.88 | 8.75E-04 | 9.93E-03 |
| GOBP_PROTEIN_LOCALIZATION_TO_NUCLEOPLASM | 13 | -1.88 | 1.75E-03 | 1.65E-02 |
| GOBP_CELL_CYCLE_G1_S_PHASE_TRANSITION | 214 | -1.89 | 8.00E-07 | 3.12E-05 |
| GOBP_RNA_3_END_PROCESSING | 79 | -1.89 | 8.98E-05 | 1.64E-03 |
| GOBP_MICROTUBULE_POLYMERIZATION_OR_DEPOLYMERIZATION | 127 | -1.90 | 2.44E-05 | 5.64E-04 |
| GOBP_RNA_MODIFICATION | 147 | -1.90 | 5.40E-06 | 1.55E-04 |
| GOBP_PROTEIN_DNA_COMPLEX_DISASSEMBLY | 19 | -1.91 | 2.20E-03 | 1.96E-02 |
| GOBP_ATP_SYNTHESIS_COUPLED_ELECTRON_TRANSPORT | 81 | -1.92 | 4.39E-05 | 9.26E-04 |
| GOBP_RRNA_MODIFICATION | 30 | -1.92 | 6.82E-04 | 8.14E-03 |
| GOBP_REGULATION_OF_DNA_REPAIR | 200 | -1.94 | 4.49E-07 | 1.86E-05 |
| GOBP_REGULATION_OF_INTRINSIC_APOPTOTIC_SIGNALING_PATHWAY_IN_RESPONSE_TO_DNA_DAMAGE_BY_P53_CLASS_MEDIATOR | 17 | -1.94 | 1.25E-03 | 1.29E-02 |
| GOBP_RRNA_PROCESSING | 203 | -1.94 | 1.88E-07 | 8.45E-06 |
| GOBP_DNA_TOPOLOGICAL_CHANGE | 10 | -1.94 | 1.10E-03 | 1.17E-02 |
| GOBP_REGULATION_OF_CELL_DIVISION | 141 | -1.94 | 4.72E-06 | 1.40E-04 |
| GOBP_SPLICEOSOMAL_SNRNP_ASSEMBLY | 40 | -1.95 | 3.41E-04 | 4.82E-03 |
| GOBP_RIBOSOME_BIOGENESIS | 287 | -1.96 | 8.38E-09 | 4.60E-07 |
| GOBP_RNA_LOCALIZATION | 181 | -1.97 | 5.61E-07 | 2.24E-05 |
| GOBP_NUCLEOTIDE_EXCISION_REPAIR | 83 | -1.97 | 2.07E-05 | 4.98E-04 |
| GOBP_MICROTUBULE_ORGANIZING_CENTER_LOCALIZATION | 27 | -1.97 | 3.23E-04 | 4.63E-03 |
| GOBP_REGULATION_OF_TELOMERE_MAINTENANCE_VIA_TELOMERE_LENGTHENING | 52 | -1.97 | 1.36E-04 | 2.32E-03 |
| GOBP_RNA_METHYLATION | 73 | -1.97 | 3.63E-05 | 7.86E-04 |
| GOBP_POSITIVE_REGULATION_OF_CYTOKINESIS | 36 | -1.99 | 2.09E-04 | 3.27E-03 |
| GOBP_POSITIVE_REGULATION_OF_ATTACHMENT_OF_SPINDLE_MICROTUBULES_TO_KINETOCHORE | 11 | -1.99 | 5.67E-04 | 6.96E-03 |
| GOBP_REGULATION_OF_DOUBLE_STRAND_BREAK_REPAIR | 123 | -2.00 | 2.60E-06 | 8.18E-05 |
| GOBP_NEGATIVE_REGULATION_OF_MRNA_PROCESSING | 21 | -2.00 | 4.49E-04 | 5.87E-03 |
| GOBP_TELOMERE_MAINTENANCE_VIA_TELOMERE_LENGTHENING | 66 | -2.00 | 6.26E-05 | 1.26E-03 |
| GOBP_REGULATION_OF_SISTER_CHROMATID_COHESION | 14 | -2.00 | 1.06E-03 | 1.15E-02 |
| GOBP_ESTABLISHMENT_OF_SISTER_CHROMATID_COHESION | 10 | -2.00 | 4.85E-04 | 6.17E-03 |
| GOBP_NEGATIVE_REGULATION_OF_RNA_SPLICING | 25 | -2.01 | 3.64E-04 | 5.00E-03 |
| GOBP_POSITIVE_REGULATION_OF_METAPHASE_ANAPHASE_TRANSITION_OF_CELL_CYCLE | 16 | -2.01 | 4.30E-04 | 5.69E-03 |
| GOBP_RRNA_METABOLIC_PROCESS | 238 | -2.01 | 1.37E-08 | 7.23E-07 |
| GOBP_NEGATIVE_REGULATION_OF_SPROUTING_ANGIOGENESIS | 12 | -2.01 | 1.66E-04 | 2.78E-03 |
| GOBP_POSITIVE_REGULATION_OF_DNA_REPLICATION | 37 | -2.01 | 1.72E-04 | 2.83E-03 |
| GOBP_PROTEIN_RNA_COMPLEX_ORGANIZATION | 193 | -2.02 | 1.83E-08 | 9.54E-07 |
| GOBP_POSITIVE_REGULATION_OF_CELL_CYCLE_G2_M_PHASE_TRANSITION | 28 | -2.02 | 1.29E-04 | 2.21E-03 |
| GOBP_NEGATIVE_REGULATION_OF_MRNA_METABOLIC_PROCESS | 80 | -2.02 | 1.15E-05 | 3.04E-04 |
| GOBP_REGULATION_OF_METAPHASE_PLATE_CONGRESSION | 15 | -2.02 | 3.58E-04 | 4.95E-03 |
| GOBP_POSITIVE_REGULATION_OF_DNA_TEMPLATED_DNA_REPLICATION | 15 | -2.03 | 3.40E-04 | 4.82E-03 |
| GOBP_PROTEIN_LOCALIZATION_TO_CONDENSED_CHROMOSOME | 14 | -2.03 | 6.34E-04 | 7.63E-03 |
| GOBP_DOUBLE_STRAND_BREAK_REPAIR_VIA_NONHOMOLOGOUS_END_JOINING | 64 | -2.03 | 2.00E-05 | 4.87E-04 |
| GOBP_REGULATION_OF_DOUBLE_STRAND_BREAK_REPAIR_VIA_HOMOLOGOUS_RECOMBINATION | 69 | -2.03 | 1.24E-05 | 3.20E-04 |
| GOBP_ALTERNATIVE_MRNA_SPLICING_VIA_SPLICEOSOME | 64 | -2.04 | 1.87E-05 | 4.62E-04 |
| GOBP_HOMOLOGOUS_CHROMOSOME_SEGREGATION | 37 | -2.04 | 1.13E-04 | 2.01E-03 |
| GOBP_REGULATION_OF_RNA_SPLICING | 161 | -2.04 | 4.79E-08 | 2.31E-06 |
| GOBP_MEIOTIC_SPINDLE_ORGANIZATION | 18 | -2.04 | 2.09E-04 | 3.27E-03 |
| GOBP_REGULATION_OF_MEIOTIC_CELL_CYCLE | 51 | -2.04 | 2.84E-05 | 6.42E-04 |
| GOBP_MITOTIC_SISTER_CHROMATID_COHESION | 26 | -2.04 | 1.07E-04 | 1.91E-03 |
| GOBP_REGULATION_OF_DNA_METABOLIC_PROCESS | 416 | -2.04 | 8.22E-13 | 8.12E-11 |
| GOBP_MITOTIC_RECOMBINATION | 26 | -2.05 | 9.93E-05 | 1.79E-03 |
| GOBP_NEGATIVE_REGULATION_OF_CELL_DIVISION | 12 | -2.05 | 9.80E-05 | 1.77E-03 |
| GOBP_RIBONUCLEOPROTEIN_COMPLEX_BIOGENESIS | 424 | -2.06 | 3.36E-13 | 3.55E-11 |
| GOBP_POSITIVE_REGULATION_OF_TELOMERE_MAINTENANCE_VIA_TELOMERE_LENGTHENING | 26 | -2.06 | 8.17E-05 | 1.54E-03 |
| GOBP_SPINDLE_ASSEMBLY_INVOLVED_IN_FEMALE_MEIOSIS | 10 | -2.06 | 1.79E-04 | 2.93E-03 |
| GOBP_DNA_INTEGRITY_CHECKPOINT_SIGNALING | 114 | -2.06 | 9.62E-07 | 3.62E-05 |
| GOBP_POSITIVE_REGULATION_OF_MITOTIC_CELL_CYCLE | 114 | -2.06 | 9.62E-07 | 3.62E-05 |
| GOBP_REGULATION_OF_ATTACHMENT_OF_MITOTIC_SPINDLE_MICROTUBULES_TO_KINETOCHORE | 10 | -2.07 | 1.64E-04 | 2.75E-03 |
| GOBP_REGULATION_OF_DNA_DAMAGE_CHECKPOINT | 21 | -2.07 | 1.95E-04 | 3.12E-03 |
| GOBP_REGULATION_OF_CELL_CYCLE_G2_M_PHASE_TRANSITION | 103 | -2.08 | 1.50E-06 | 5.40E-05 |
| GOBP_DNA_CONFORMATION_CHANGE | 16 | -2.09 | 1.23E-04 | 2.17E-03 |
| GOBP_MEMBRANELESS_ORGANELLE_ASSEMBLY | 363 | -2.11 | 9.01E-13 | 8.71E-11 |
| GOBP_REGULATION_OF_CYTOKINESIS | 77 | -2.11 | 2.42E-06 | 7.74E-05 |
| GOBP_MITOTIC_DNA_REPLICATION_CHECKPOINT_SIGNALING | 10 | -2.11 | 5.86E-05 | 1.20E-03 |

| | | | | | |
|---|---|---|---|---|---|
| | GOBP_REGULATION_OF_MRNA_PROCESSING | 113 | -2.12 | 4.01E-07 | 1.68E-05 |
| | GOBP_CELL_CYCLE_G2_M_PHASE_TRANSITION | 139 | -2.13 | 8.70E-08 | 4.03E-06 |
| | GOBP_SISTER_CHROMATID_COHESION | 49 | -2.13 | 7.20E-06 | 2.01E-04 |
| | GOBP_TELOMERASE_RNA_LOCALIZATION | 19 | -2.13 | 8.83E-05 | 1.62E-03 |
| | GOBP_POSITIVE_REGULATION_OF_SPINDLE_CHECKPOINT | 11 | -2.14 | 6.22E-05 | 1.26E-03 |
| | GOBP_REGULATION_OF_MITOTIC_CELL_CYCLE | 434 | -2.14 | 1.24E-15 | 1.72E-13 |
| | GOBP_MEIOTIC_SPINDLE_ASSEMBLY | 12 | -2.15 | 1.46E-05 | 3.71E-04 |
| | GOBP_SPLICEOSOMAL_COMPLEX_ASSEMBLY | 71 | -2.15 | 8.27E-07 | 3.17E-05 |
| | GOBP_MITOTIC_METAPHASE_CHROMOSOME_ALIGNMENT | 64 | -2.16 | 2.40E-06 | 7.74E-05 |
| | GOBP_POSITIVE_REGULATION_OF_MRNA_PROCESSING | 26 | -2.16 | 1.78E-05 | 4.42E-04 |
| | GOBP_PROTEIN_LOCALIZATION_TO_CHROMATIN | 35 | -2.17 | 1.08E-05 | 2.92E-04 |
| | GOBP_SPINDLE_ELONGATION | 13 | -2.17 | 1.33E-05 | 3.43E-04 |
| | GOBP_MEIOTIC_CELL_CYCLE_PHASE_TRANSITION | 12 | -2.18 | 8.33E-06 | 2.30E-04 |
| | GOBP_MICROTUBULE_ORGANIZING_CENTER_ORGANIZATION | 140 | -2.18 | 7.67E-09 | 4.37E-07 |
| | GOBP_PROTEIN_DNA_COMPLEX_ORGANIZATION | 169 | -2.18 | 1.17E-09 | 7.22E-08 |
| | GOBP_POSITIVE_REGULATION_OF_CELL_CYCLE_CHECKPOINT | 16 | -2.20 | 2.20E-05 | 5.17E-04 |
| | GOBP_REGULATION_OF_CENTRIOLE_REPLICATION | 23 | -2.20 | 5.64E-06 | 1.61E-04 |
| | GOBP_KINETOCHORE_ASSEMBLY | 16 | -2.20 | 2.09E-05 | 4.99E-04 |
| | GOBP_CHROMATIN_REMODELING_AT_CENTROMERE | 10 | -2.20 | 5.81E-06 | 1.64E-04 |
| | GOBP_MITOTIC_SPINDLE_ASSEMBLY | 71 | -2.20 | 2.81E-07 | 1.20E-05 |
| | GOBP_BASE_EXCISION_REPAIR | 47 | -2.21 | 2.82E-06 | 8.77E-05 |
| | GOBP_MITOTIC_DNA_REPLICATION | 16 | -2.21 | 1.54E-05 | 3.88E-04 |
| | GOBP_TELOMERE_ORGANIZATION | 147 | -2.22 | 1.83E-09 | 1.10E-07 |
| | GOBP_NEGATIVE_REGULATION_OF_MITOTIC_CELL_CYCLE | 198 | -2.22 | 4.95E-11 | 3.67E-09 |
| | GOBP_DOUBLE_STRAND_BREAK_REPAIR_VIA_BREAK_INDUCED_REPLICATION | 12 | -2.25 | 1.96E-06 | 6.56E-05 |
| | GOBP_NEGATIVE_REGULATION_OF_CELL_CYCLE | 322 | -2.25 | 5.45E-15 | 6.55E-13 |
| | GOBP_CHROMOSOME_ORGANIZATION_INVOLVED_IN_MEIOTIC_CELL_CYCLE | 39 | -2.26 | 1.13E-06 | 4.21E-05 |
| | GOBP_POSITIVE_REGULATION_OF_MITOTIC_SISTER_CHROMATID_SEPARATION | 20 | -2.27 | 7.25E-06 | 2.01E-04 |
| | GOBP_REGULATION_OF_CHROMOSOME_CONDENSATION | 12 | -2.27 | 1.14E-06 | 4.23E-05 |
| | GOBP_U2_TYPE_PRESPLICEOSOME_ASSEMBLY | 24 | -2.27 | 4.24E-06 | 1.26E-04 |
| | GOBP_REGULATION_OF_DNA_REPLICATION | 103 | -2.28 | 2.29E-08 | 1.17E-06 |
| | GOBP_MITOTIC_SPINDLE_ELONGATION | 11 | -2.29 | 1.65E-06 | 5.73E-05 |
| | GOBP_ATTACHMENT_OF_MITOTIC_SPINDLE_MICROTUBULES_TO_KINETOCHORE | 26 | -2.29 | 1.76E-06 | 5.93E-05 |
| | GOBP_MITOTIC_CELL_CYCLE_PHASE_TRANSITION | 382 | -2.29 | 3.34E-18 | 6.74E-16 |
| | GOBP_POSITIVE_REGULATION_OF_MITOTIC_CELL_CYCLE_PHASE_TRANSITION | 83 | -2.30 | 2.72E-08 | 1.38E-06 |
| | GOBP_CENTROSOME_SEPARATION | 15 | -2.30 | 1.74E-06 | 5.93E-05 |
| | GOBP_NEGATIVE_REGULATION_OF_MITOTIC_CELL_CYCLE_PHASE_TRANSITION | 157 | -2.31 | 1.06E-10 | 7.61E-09 |
| | GOBP_PROTEIN_LOCALIZATION_TO_CHROMOSOME_CENTROMERIC_REGION | 22 | -2.31 | 6.42E-06 | 1.81E-04 |
| | GOBP_REGULATION_OF_MRNA_SPLICING_VIA_SPLICEOSOME | 91 | -2.33 | 5.92E-09 | 3.46E-07 |
| | GOBP_CENTROSOME_DUPLICATION | 71 | -2.34 | 1.06E-08 | 5.75E-07 |
| | GOBP_CELL_CYCLE_PHASE_TRANSITION | 473 | -2.34 | 3.40E-22 | 1.26E-19 |
| | GOBP_MITOTIC_CHROMOSOME_CONDENSATION | 15 | -2.35 | 5.54E-07 | 2.24E-05 |
| | GOBP_DNA_REPLICATION_CHECKPOINT_SIGNALING | 18 | -2.35 | 8.25E-07 | 3.17E-05 |
| | GOBP_REGULATION_OF_DNA_TEMPLATED_DNA_REPLICATION_INITIATION | 14 | -2.36 | 7.03E-07 | 2.76E-05 |
| | GOBP_MITOCHONDRIAL_GENE_EXPRESSION | 156 | -2.36 | 2.54E-11 | 2.02E-09 |
| | GOBP_REGULATION_OF_MITOTIC_CELL_CYCLE_PHASE_TRANSITION | 295 | -2.36 | 1.19E-16 | 1.89E-14 |
| | GOBP_HOMOLOGOUS_RECOMBINATION | 46 | -2.36 | 7.37E-08 | 3.45E-06 |
| | GOBP_DNA_STRAND_ELONGATION | 35 | -2.36 | 1.56E-07 | 7.08E-06 |
| | GOBP_CHROMOSOME_CONDENSATION | 26 | -2.37 | 2.70E-07 | 1.16E-05 |
| | GOBP_DNA_STRAND_ELONGATION_INVOLVED_IN_DNA_REPLICATION | 15 | -2.38 | 2.96E-07 | 1.25E-05 |
| | GOBP_MRNA_PROCESSING | 435 | -2.38 | 7.97E-22 | 2.72E-19 |
| | GOBP_FEMALE_MEIOTIC_NUCLEAR_DIVISION | 30 | -2.38 | 1.94E-07 | 8.61E-06 |
| | GOBP_POSITIVE_REGULATION_OF_CHROMOSOME_ORGANIZATION | 95 | -2.41 | 8.96E-10 | 5.61E-08 |
| | GOBP_NEGATIVE_REGULATION_OF_CELL_CYCLE_PROCESS | 261 | -2.41 | 4.27E-16 | 6.55E-14 |
| | GOBP_REGULATION_OF_CELL_CYCLE_PHASE_TRANSITION | 383 | -2.41 | 1.87E-21 | 5.55E-19 |
| | GOBP_POSITIVE_REGULATION_OF_CELL_CYCLE | 268 | -2.41 | 7.63E-17 | 1.26E-14 |
| | GOBP_POSITIVE_REGULATION_OF_CELL_CYCLE_PHASE_TRANSITION | 100 | -2.42 | 1.24E-10 | 8.74E-09 |
| | GOBP_MITOCHONDRIAL_TRANSLATION | 125 | -2.43 | 1.38E-11 | 1.14E-09 |
| | GOBP_INTERSTRAND_CROSS_LINK_REPAIR | 35 | -2.44 | 3.11E-08 | 1.55E-06 |
| | GOBP_PROTEIN_LOCALIZATION_TO_CHROMOSOME | 91 | -2.44 | 3.25E-10 | 2.22E-08 |
| | GOBP_RNA_SPLICING | 408 | -2.44 | 1.91E-22 | 7.70E-20 |
| | GOBP_REGULATION_OF_SPINDLE_CHECKPOINT | 20 | -2.45 | 9.49E-08 | 4.35E-06 |
| | GOBP_MITOTIC_CELL_CYCLE_CHECKPOINT_SIGNALING | 121 | -2.45 | 4.50E-12 | 3.92E-10 |
| | GOBP_SPINDLE_ASSEMBLY | 126 | -2.47 | 1.91E-12 | 1.73E-10 |
| | GOBP_KINETOCHORE_ORGANIZATION | 20 | -2.48 | 3.77E-08 | 1.86E-06 |
| | GOBP_MICROTUBULE_CYTOSKELETON_ORGANIZATION_INVOLVED_IN_MITOSIS | 159 | -2.49 | 5.45E-14 | 5.93E-12 |
| | GOBP_REGULATION_OF_DNA_TEMPLATED_DNA_REPLICATION | 36 | -2.49 | 8.20E-09 | 4.55E-07 |
| | GOBP_REGULATION_OF_CELL_CYCLE_CHECKPOINT | 42 | -2.52 | 5.84E-10 | 3.76E-08 |
| | GOBP_DNA_RECOMBINATION | 267 | -2.52 | 4.43E-20 | 1.04E-17 |
| | GOBP_MITOTIC_SPINDLE_ORGANIZATION | 127 | -2.54 | 3.94E-13 | 4.07E-11 |
| | GOBP_CELL_CYCLE_CHECKPOINT_SIGNALING | 167 | -2.55 | 7.56E-16 | 1.08E-13 |
| | GOBP_ORGANELLE_FISSION | 397 | -2.55 | 2.09E-26 | 1.55E-23 |
| | GOBP_NEGATIVE_REGULATION_OF_CHROMOSOME_ORGANIZATION | 86 | -2.55 | 5.14E-12 | 4.39E-10 |
| | GOBP_DNA_TEMPLATED_DNA_REPLICATION_MAINTENANCE_OF_FIDELITY | 52 | -2.56 | 3.98E-10 | 2.68E-08 |
| | GOBP_SPINDLE_ORGANIZATION | 188 | -2.56 | 4.45E-17 | 7.90E-15 |
| | GOBP_CHROMOSOME_LOCALIZATION | 111 | -2.57 | 6.25E-13 | 6.32E-11 |
| | GOBP_METAPHASE_CHROMOSOME_ALIGNMENT | 97 | -2.57 | 1.61E-12 | 1.49E-10 |
| | GOBP_POSITIVE_REGULATION_OF_CHROMOSOME_SEPARATION | 30 | -2.58 | 8.90E-10 | 5.61E-08 |
| | GOBP_POSITIVE_REGULATION_OF_CHROMOSOME_SEGREGATION | 29 | -2.58 | 4.07E-10 | 2.70E-08 |
| | GOBP_REGULATION_OF_ATTACHMENT_OF_SPINDLE_MICROTUBULES_TO_KINETOCHORE | 23 | -2.59 | 1.62E-10 | 1.12E-08 |
| | GOBP_MEIOTIC_CELL_CYCLE | 195 | -2.60 | 8.69E-18 | 1.68E-15 |
| | GOBP_MEIOSIS_I_CELL_CYCLE_PROCESS | 88 | -2.60 | 1.48E-12 | 1.40E-10 |
| | GOBP_MEIOTIC_CHROMOSOME_SEGREGATION | 60 | -2.60 | 7.54E-11 | 5.49E-09 |
| | GOBP_REGULATION_OF_NUCLEAR_DIVISION | 119 | -2.62 | 2.69E-14 | 3.06E-12 |
| | GOBP_CENTRIOLE_ASSEMBLY | 46 | -2.64 | 2.08E-11 | 1.68E-09 |
| | GOBP_DOUBLE_STRAND_BREAK_REPAIR | 275 | -2.64 | 1.54E-21 | 4.88E-19 |
| | GOBP_CENTROMERE_COMPLEX_ASSEMBLY | 27 | -2.64 | 4.60E-11 | 3.46E-09 |
| | GOBP_CELL_CYCLE_DNA_REPLICATION | 41 | -2.67 | 2.70E-11 | 2.11E-09 |
| | GOBP_DNA_REPLICATION_INITIATION | 35 | -2.68 | 1.01E-11 | 8.44E-10 |
| | GOBP_RNA_SPLICING_VIA_TRANSESTERIFICATION_REACTIONS | 277 | -2.68 | 6.24E-24 | 2.77E-21 |
| | GOBP_REGULATION_OF_CHROMOSOME_ORGANIZATION | 219 | -2.68 | 5.76E-21 | 1.51E-18 |
| | GOBP_MEIOTIC_CELL_CYCLE_PROCESS | 145 | -2.69 | 5.05E-17 | 8.63E-15 |
| | GOBP_POSITIVE_REGULATION_OF_CELL_CYCLE_PROCESS | 212 | -2.70 | 2.17E-21 | 6.04E-19 |
| | GOBP_REGULATION_OF_MITOTIC_NUCLEAR_DIVISION | 101 | -2.71 | 4.44E-15 | 5.64E-13 |
| | GOBP_RECOMBINATIONAL_REPAIR | 166 | -2.73 | 2.31E-19 | 5.12E-17 |
| | GOBP_MITOTIC_NUCLEAR_DIVISION | 251 | -2.75 | 2.98E-24 | 1.47E-21 |
| | GOBP_DNA_REPLICATION | 249 | -2.77 | 4.24E-25 | 2.35E-22 |
| | GOBP_ATTACHMENT_OF_SPINDLE_MICROTUBULES_TO_KINETOCHORE | 50 | -2.82 | 4.65E-15 | 5.73E-13 |
| | GOBP_NEGATIVE_REGULATION_OF_CHROMOSOME_SEGREGATION | 46 | -2.84 | 6.90E-15 | 8.07E-13 |
| | GOBP_NEGATIVE_REGULATION_OF_NUCLEAR_DIVISION | 56 | -2.85 | 1.60E-15 | 2.15E-13 |
| | GOBP_CHROMOSOME_SEGREGATION | 367 | -2.87 | 8.40E-37 | 1.87E-33 |
| | GOBP_MITOTIC_SISTER_CHROMATID_SEPARATION | 58 | -2.88 | 2.00E-15 | 2.61E-13 |
| | GOBP_CHROMOSOME_ORGANIZATION | 462 | -2.89 | 5.06E-43 | 2.25E-39 |
| | GOBP_REGULATION_OF_SISTER_CHROMATID_SEGREGATION | 97 | -2.89 | 2.15E-18 | 4.55E-16 |
| | GOBP_CHROMOSOME_SEPARATION | 74 | -2.90 | 1.27E-17 | 2.36E-15 |
| | GOBP_REGULATION_OF_MITOTIC_SISTER_CHROMATID_SEGREGATION | 51 | -2.90 | 4.52E-16 | 6.70E-14 |
| | GOBP_REGULATION_OF_CHROMOSOME_SEGREGATION | 120 | -2.93 | 8.43E-21 | 2.08E-18 |
| | GOBP_NUCLEAR_CHROMOSOME_SEGREGATION | 270 | -2.96 | 3.70E-33 | 5.48E-30 |
| | GOBP_SISTER_CHROMATID_SEGREGATION | 215 | -2.99 | 2.89E-29 | 3.21E-26 |
| | GOBP_MITOTIC_SISTER_CHROMATID_SEGREGATION | 178 | -3.02 | 8.15E-29 | 7.25E-26 |
| | GOBP_DNA_TEMPLATED_DNA_REPLICATION | 145 | -3.04 | 2.85E-25 | 1.81E-22 |

Table S4. Significantly up- and downregulated GSEA terms with age in SHOCK induced neurons.

| Induced neurons | | | | | |
|---|---|---|---|---|---|
| GSEA Gene Set Database | Gene Set | Size | Normalized enrichment score (NES) | Nominal p-value (NOM p-val) | Adjusted p-value |
| c1: Hallmark | HALLMARK_TNFA_SIGNALING_VIA_NFKB | 183 | 2.77 | 8.70E-21 | 2.17E-19 |
| | HALLMARK_MYC_TARGETS_V1 | 200 | 2.76 | 4.35E-21 | 2.17E-19 |
| | HALLMARK_EPITHELIAL_MESENCHYMAL_TRANSITION | 195 | 2.63 | 8.16E-18 | 1.36E-16 |
| | HALLMARK_INFLAMMATORY_RESPONSE | 148 | 2.50 | 5.75E-14 | 5.75E-13 |
| | HALLMARK_MTORC1_SIGNALING | 199 | 2.48 | 3.87E-15 | 4.84E-14 |
| | HALLMARK_UNFOLDED_PROTEIN_RESPONSE | 112 | 2.45 | 1.03E-10 | 7.39E-10 |
| | HALLMARK_COMPLEMENT | 159 | 2.43 | 1.37E-12 | 1.14E-11 |
| | HALLMARK_IL6_JAK_STAT3_SIGNALING | 66 | 2.42 | 2.44E-08 | 1.02E-07 |
| | HALLMARK_COAGULATION | 98 | 2.36 | 3.95E-09 | 1.80E-08 |
| | HALLMARK_XENOBIOTIC_METABOLISM | 165 | 2.22 | 6.36E-10 | 3.53E-09 |
| | HALLMARK_GLYCOLYSIS | 188 | 2.19 | 4.64E-10 | 2.90E-09 |
| | HALLMARK_HYPOXIA | 191 | 2.15 | 1.59E-09 | 7.97E-09 |
| | HALLMARK_ALLOGRAFT_REJECTION | 131 | 2.13 | 4.56E-08 | 1.76E-07 |
| | HALLMARK_MYC_TARGETS_V2 | 58 | 2.08 | 2.46E-05 | 5.58E-05 |
| | HALLMARK_APOPTOSIS | 144 | 2.07 | 1.34E-07 | 4.18E-07 |
| | HALLMARK_INTERFERON_GAMMA_RESPONSE | 177 | 1.99 | 7.63E-08 | 2.73E-07 |
| | HALLMARK_P53_PATHWAY | 192 | 1.98 | 1.02E-07 | 3.40E-07 |
| | HALLMARK_ANDROGEN_RESPONSE | 93 | 1.98 | 1.44E-05 | 3.43E-05 |
| | HALLMARK_TGF_BETA_SIGNALING | 52 | 1.95 | 2.12E-04 | 4.07E-04 |
| | HALLMARK_UV_RESPONSE_DN | 144 | 1.94 | 1.89E-06 | 5.24E-06 |
| | HALLMARK_PROTEIN_SECRETION | 95 | 1.93 | 2.67E-05 | 5.80E-05 |
| | HALLMARK_INTERFERON_ALPHA_RESPONSE | 91 | 1.91 | 6.10E-05 | 1.22E-04 |
| | HALLMARK_UV_RESPONSE_UP | 152 | 1.91 | 1.74E-06 | 5.12E-06 |
| | HALLMARK_FATTY_ACID_METABOLISM | 140 | 1.90 | 7.76E-06 | 1.94E-05 |
| | HALLMARK_MYOGENESIS | 171 | 1.86 | 3.69E-06 | 9.72E-06 |
| | HALLMARK_KRAS_SIGNALING_UP | 165 | 1.76 | 4.58E-05 | 9.54E-05 |
| | HALLMARK_CHOLESTEROL_HOMEOSTASIS | 72 | 1.75 | 1.16E-03 | 1.94E-03 |
| | HALLMARK_ANGIOGENESIS | 30 | 1.68 | 1.15E-02 | 1.69E-02 |
| | HALLMARK_REACTIVE_OXYGEN_SPECIES_PATHWAY | 47 | 1.65 | 6.30E-03 | 9.84E-03 |
| | HALLMARK_IL2_STAT5_SIGNALING | 166 | 1.65 | 2.67E-04 | 4.95E-04 |
| | HALLMARK_ESTROGEN_RESPONSE_EARLY | 178 | 1.62 | 3.54E-04 | 6.32E-04 |
| | HALLMARK_ADIPOGENESIS | 192 | 1.59 | 4.60E-04 | 7.93E-04 |
| | HALLMARK_APICAL_JUNCTION | 170 | 1.57 | 1.26E-03 | 2.03E-03 |
| | HALLMARK_BILE_ACID_METABOLISM | 91 | 1.52 | 1.40E-02 | 2.00E-02 |
| | HALLMARK_HEME_METABOLISM | 169 | 1.37 | 1.51E-02 | 2.10E-02 |
| | HALLMARK_SPERMATOGENESIS | 89 | -1.41 | 2.98E-02 | 4.02E-02 |
| | HALLMARK_MITOTIC_SPINDLE | 198 | -1.44 | 7.02E-03 | 1.06E-02 |
| c2: KEGG Medicus | KEGG_CYTOKINE_CYTOKINE_RECEPTOR_INTERACTION | 142 | 2.56 | 7.31E-14 | 3.99E-11 |
| | KEGG_LEISHMANIA_INFECTION | 51 | 2.40 | 5.88E-08 | 1.61E-05 |
| | KEGG_PRION_DISEASES | 26 | 2.28 | 8.02E-06 | 6.25E-04 |
| | KEGG_HEMATOPOIETIC_CELL_LINEAGE | 41 | 2.22 | 3.75E-06 | 4.09E-04 |
| | KEGG_COMPLEMENT_AND_COAGULATION_CASCADES | 42 | 2.20 | 4.71E-06 | 4.28E-04 |
| | KEGG_PROPANOATE_METABOLISM | 30 | 2.18 | 2.98E-05 | 1.48E-03 |
| | KEGG_AMINOACYL_TRNA_BIOSYNTHESIS | 41 | 2.16 | 1.21E-05 | 7.32E-04 |
| | KEGG_MEDICUS_PATHOGEN_SARS_COV_2_S_TO_ANGII_AT1R_NOX2_SIGNALING_PATHWAY | 10 | 2.11 | 3.29E-05 | 1.50E-03 |
| | KEGG_MEDICUS_VARIANT_SCRAPIE_CONFORMATION_PRPSC_TO_26S_PROTEASOME_MEDIATED_PROTEIN_DEGRADATION | 35 | 2.10 | 6.48E-05 | 2.53E-03 |
| | KEGG_MEDICUS_VARIANT_MUTATION_CAUSED_ABERRANT_ABETA_TO_26S_PROTEASOME_MEDIATED_PROTEIN_DEGRADATION | 35 | 2.09 | 7.18E-05 | 2.53E-03 |
| | KEGG_MEDICUS_VARIANT_MUTATION_INACTIVATED_VCP_TO_26S_PROTEASOME_MEDIATED_PROTEIN_DEGRADATION | 39 | 2.07 | 7.40E-05 | 2.53E-03 |
| | KEGG_RIBOSOME | 85 | 2.06 | 2.69E-06 | 3.68E-04 |
| | KEGG_MEDICUS_VARIANT_MUTATION_CAUSED_ABERRANT_HTT_TO_26S_PROTEASOME_MEDIATED_PROTEIN_DEGRADATION | 35 | 2.06 | 1.23E-04 | 3.39E-03 |
| | KEGG_MEDICUS_REFERENCE_CROSSTALK_BETWEEN_EXTRINSIC_AND_INTRINSIC_APOPTOTIC_PATHWAYS | 14 | 2.05 | 3.07E-04 | 6.20E-03 |
| | KEGG_MEDICUS_VARIANT_MUTATION_INACTIVATED_UBQLN2_TO_26S_PROTEASOME_MEDIATED_PROTEIN_DEGRADATION | 39 | 2.05 | 1.01E-04 | 3.06E-03 |
| | KEGG_MEDICUS_REFERENCE_TRANSLATION_INITIATION | 80 | 2.04 | 1.08E-05 | 7.32E-04 |
| | KEGG_MEDICUS_REFERENCE_26S_PROTEASOME_MEDIATED_PROTEIN_DEGRADATION | 38 | 2.01 | 2.53E-04 | 5.52E-03 |
| | KEGG_MEDICUS_VARIANT_MUTATION_CAUSED_ABERRANT_SNCA_TO_26S_PROTEASOME_MEDIATED_PROTEIN_DEGRADATION | 35 | 2.01 | 2.08E-04 | 4.93E-03 |
| | KEGG_MEDICUS_REFERENCE_ASSEMBLY_AND_TRAFFICKING_OF_TELOMERASE | 17 | 2.00 | 9.62E-04 | 1.42E-02 |
| | KEGG_MEDICUS_VARIANT_MUTATION_CAUSED_ABERRANT_SOD1_TO_26S_PROTEASOME_MEDIATED_PROTEIN_DEGRADATION | 40 | 1.99 | 2.07E-04 | 4.93E-03 |
| | KEGG_METABOLISM_OF_XENOBIOTICS_BY_CYTOCHROME_P450 | 29 | 1.99 | 8.19E-04 | 1.24E-02 |
| | KEGG_ECM_RECEPTOR_INTERACTION | 72 | 1.98 | 4.24E-05 | 1.78E-03 |
| | KEGG_GLUTATHIONE_METABOLISM | 41 | 1.98 | 1.83E-04 | 4.75E-03 |
| | KEGG_MEDICUS_VARIANT_MUTATION_CAUSED_ABERRANT_ABETA_TO_CROSSTALK_BETWEEN_EXTRINSIC_AND_INTRINSIC_APOPTOTIC_PATHWAYS | 11 | 1.98 | 5.62E-04 | 9.30E-03 |
| | KEGG_MEDICUS_REFERENCE_COPI_VESICLE_FORMATION | 22 | 1.97 | 4.74E-04 | 8.36E-03 |
| | KEGG_ARGININE_AND_PROLINE_METABOLISM | 46 | 1.97 | 2.31E-04 | 5.25E-03 |
| | KEGG_MEDICUS_REFERENCE_PRNP_PI3K_NOX2_SIGNALING_PATHWAY | 25 | 1.95 | 1.04E-03 | 1.46E-02 |
| | KEGG_NOD_LIKE_RECEPTOR_SIGNALING_PATHWAY | 52 | 1.94 | 1.24E-04 | 3.39E-03 |
| | KEGG_GLYCOSAMINOGLYCAN_BIOSYNTHESIS_HEPARAN_SULFATE | 26 | 1.93 | 1.22E-03 | 1.62E-02 |
| | KEGG_CITRATE_CYCLE_TCA_CYCLE | 29 | 1.93 | 1.65E-03 | 1.95E-02 |
| | KEGG_PROTEASOME | 41 | 1.92 | 4.44E-04 | 8.36E-03 |
| | KEGG_GLYCOLYSIS_GLUCONEOGENESIS | 47 | 1.90 | 3.82E-04 | 7.45E-03 |
| | KEGG_N_GLYCAN_BIOSYNTHESIS | 46 | 1.90 | 5.32E-04 | 9.08E-03 |
| | KEGG_ANTIGEN_PROCESSING_AND_PRESENTATION | 50 | 1.90 | 2.99E-04 | 6.20E-03 |
| | KEGG_MEDICUS_REFERENCE_N_GLYCAN_PRECURSOR_BIOSYNTHESIS_ALG6_TO_OST | 11 | 1.89 | 1.59E-03 | 1.93E-02 |
| | KEGG_PATHWAYS_IN_CANCER | 287 | 1.86 | 9.79E-08 | 1.78E-05 |
| | KEGG_CHEMOKINE_SIGNALING_PATHWAY | 134 | 1.86 | 2.78E-05 | 1.48E-03 |
| | KEGG_PYRUVATE_METABOLISM | 35 | 1.84 | 1.57E-03 | 1.93E-02 |
| | KEGG_VALINE_LEUCINE_AND_ISOLEUCINE_DEGRADATION | 42 | 1.82 | 1.52E-03 | 1.93E-02 |
| | KEGG_MEDICUS_REFERENCE_ITGA_B_TALIN_VINCULIN_SIGNALING_PATHWAY | 28 | 1.81 | 3.70E-03 | 4.04E-02 |
| | KEGG_CYSTEINE_AND_METHIONINE_METABOLISM | 28 | 1.80 | 4.52E-03 | 4.65E-02 |
| | KEGG_GRAFT_VERSUS_HOST_DISEASE | 20 | 1.80 | 2.59E-03 | 2.95E-02 |
| | KEGG_CHRONIC_MYELOID_LEUKEMIA | 70 | 1.79 | 4.73E-04 | 8.36E-03 |
| | KEGG_ADHERENS_JUNCTION | 70 | 1.74 | 9.91E-04 | 1.42E-02 |
| | KEGG_APOPTOSIS | 77 | 1.71 | 7.74E-04 | 1.23E-02 |
| | KEGG_TGF_BETA_SIGNALING_PATHWAY | 78 | 1.70 | 1.56E-03 | 1.93E-02 |
| | KEGG_FOCAL_ADHESION | 180 | 1.69 | 8.09E-05 | 2.60E-03 |
| | KEGG_COLORECTAL_CANCER | 60 | 1.67 | 4.12E-03 | 4.33E-02 |
| | KEGG_TIGHT_JUNCTION | 105 | 1.65 | 1.14E-03 | 1.56E-02 |
| | KEGG_VIRAL_MYOCARDITIS | 53 | 1.64 | 4.83E-03 | 4.74E-02 |
| | KEGG_PURINE_METABOLISM | 139 | 1.59 | 7.90E-04 | 1.23E-02 |
| | KEGG_RENAL_CELL_CARCINOMA | 67 | 1.58 | 4.86E-03 | 4.74E-02 |
| | KEGG_LYSOSOME | 113 | 1.50 | 5.20E-03 | 4.99E-02 |
| | KEGG_SPLICEOSOME | 126 | 1.50 | 3.97E-03 | 4.25E-02 |
| | KEGG_LONG_TERM_POTENTIATION | 62 | -1.67 | 2.94E-03 | 3.27E-02 |
| | KEGG_MEDICUS_REFERENCE_ORGANIZATION_OF_THE_OUTER_KINETOCHORE | 19 | -1.86 | 4.79E-03 | 4.74E-02 |
| | KEGG_MEDICUS_REFERENCE_DEPHOSPHORYLATION_OF_KINETOCHORE | 12 | -1.92 | 1.81E-03 | 2.10E-02 |
| c5: GO Biological Process | GOBP_HUMORAL_IMMUNE_RESPONSE | 121 | 2.53 | 4.60E-13 | 1.57E-10 |
| | GOBP_NEGATIVE_REGULATION_OF_INTRINSIC_APOPTOTIC_SIGNALING_PATHWAY | 101 | 2.47 | 1.74E-11 | 3.47E-09 |
| | GOBP_MYELOID_LEUKOCYTE_MIGRATION | 160 | 2.46 | 1.19E-12 | 3.79E-10 |
| | GOBP_REGULATION_OF_NITRIC_OXIDE_METABOLIC_PROCESS | 54 | 2.45 | 2.29E-08 | 1.79E-06 |
| | GOBP_CELL_CHEMOTAXIS | 212 | 2.44 | 2.11E-15 | 2.01E-12 |
| | GOBP_NEGATIVE_REGULATION_OF_APOPTOTIC_SIGNALING_PATHWAY | 207 | 2.44 | 1.05E-14 | 8.21E-12 |

| | | | | |
|---|---|---|---|---|
| GOBP_GRANULOCYTE_CHEMOTAXIS | 77 | 2.42 | 1.78E-09 | 1.98E-07 |
| GOBP_POSITIVE_REGULATION_OF_EPITHELIAL_TO_MESENCHYMAL_TRANSITION | 53 | 2.42 | 2.05E-08 | 1.63E-06 |
| GOBP_GRANULOCYTE_MIGRATION | 96 | 2.42 | 9.20E-11 | 1.46E-08 |
| GOBP_COMPLEMENT_ACTIVATION | 27 | 2.39 | 1.71E-06 | 6.36E-05 |
| GOBP_ANTIMICROBIAL_HUMORAL_IMMUNE_RESPONSE_MEDIATED_BY_ANTIMICROBIAL_PEPTIDE | 50 | 2.38 | 4.75E-08 | 3.24E-06 |
| GOBP_RIBOSOME_BIOGENESIS | 315 | 2.37 | 2.92E-17 | 1.03E-13 |
| GOBP_REACTIVE_NITROGEN_SPECIES_METABOLIC_PROCESS | 73 | 2.35 | 6.61E-09 | 5.95E-07 |
| GOBP_CYTOPLASMIC_TRANSLATIONAL_INITIATION | 46 | 2.35 | 2.12E-07 | 1.15E-05 |
| GOBP_LEUKOCYTE_CHEMOTAXIS | 155 | 2.35 | 8.77E-12 | 2.46E-09 |
| GOBP_POSITIVE_REGULATION_OF_SMOOTH_MUSCLE_CELL_PROLIFERATION | 68 | 2.34 | 3.49E-08 | 2.52E-06 |
| GOBP_POSITIVE_REGULATION_OF_NITRIC_OXIDE_METABOLIC_PROCESS | 34 | 2.32 | 1.17E-06 | 4.66E-05 |
| GOBP_LEUKOCYTE_MIGRATION | 272 | 2.32 | 1.20E-14 | 8.21E-12 |
| GOBP_REGULATION_OF_INFLAMMATORY_RESPONSE_TO_ANTIGENIC_STIMULUS | 26 | 2.31 | 3.23E-06 | 1.03E-04 |
| GOBP_POSITIVE_REGULATION_OF_VASCULATURE_DEVELOPMENT | 137 | 2.31 | 8.36E-11 | 1.38E-08 |
| GOBP_RRNA_PROCESSING | 220 | 2.31 | 4.18E-13 | 1.54E-10 |
| GOBP_REGULATION_OF_INTRINSIC_APOPTOTIC_SIGNALING_PATHWAY | 170 | 2.30 | 1.21E-11 | 2.83E-09 |
| GOBP_CYTOPLASMIC_TRANSLATION | 167 | 2.30 | 3.62E-11 | 6.40E-09 |
| GOBP_REGULATION_OF_LEUKOCYTE_MIGRATION | 170 | 2.30 | 1.29E-11 | 2.83E-09 |
| GOBP_GLYCOSAMINOGLYCAN_BIOSYNTHETIC_PROCESS | 44 | 2.30 | 5.18E-07 | 2.33E-05 |
| GOBP_REGULATION_OF_APOPTOTIC_SIGNALING_PATHWAY | 342 | 2.29 | 6.48E-17 | 1.03E-13 |
| GOBP_RRNA_METABOLIC_PROCESS | 263 | 2.29 | 3.77E-14 | 2.25E-11 |
| GOBP_ANTIMICROBIAL_HUMORAL_RESPONSE | 62 | 2.28 | 1.38E-07 | 8.34E-06 |
| GOBP_REGULATION_OF_EXTRACELLULAR_MATRIX_ORGANIZATION | 60 | 2.27 | 3.10E-07 | 1.57E-05 |
| GOBP_VIRAL_PROCESS | 358 | 2.26 | 8.89E-17 | 1.06E-13 |
| GOBP_BIOLOGICAL_PROCESS_INVOLVED_IN_SYMBIOTIC_INTERACTION | 165 | 2.26 | 3.91E-10 | 4.85E-08 |
| GOBP_RIBOSOMAL_SMALL_SUBUNIT_BIOGENESIS | 108 | 2.26 | 1.09E-08 | 9.42E-07 |
| GOBP_INTRINSIC_APOPTOTIC_SIGNALING_PATHWAY | 300 | 2.26 | 4.87E-14 | 2.58E-11 |
| GOBP_POSITIVE_REGULATION_OF_MIRNA_TRANSCRIPTION | 51 | 2.24 | 1.23E-06 | 4.87E-05 |
| GOBP_MACROPHAGE_CHEMOTAXIS | 31 | 2.24 | 1.28E-05 | 3.31E-04 |
| GOBP_OVULATION_CYCLE_PROCESS | 37 | 2.24 | 3.21E-06 | 1.03E-04 |
| GOBP_CHAPERONE_MEDIATED_PROTEIN_FOLDING | 66 | 2.24 | 1.93E-07 | 1.07E-05 |
| GOBP_AMINOGLYCAN_BIOSYNTHETIC_PROCESS | 51 | 2.23 | 1.43E-06 | 5.47E-05 |
| GOBP_AMINO_ACID_ACTIVATION | 45 | 2.23 | 4.52E-06 | 1.39E-04 |
| GOBP_EXTRINSIC_APOPTOTIC_SIGNALING_PATHWAY_VIA_DEATH_DOMAIN_RECEPTORS | 75 | 2.22 | 2.04E-07 | 1.12E-05 |
| GOBP_POSITIVE_REGULATION_OF_REACTIVE_OXYGEN_SPECIES_METABOLIC_PROCESS | 52 | 2.21 | 2.72E-06 | 9.28E-05 |
| GOBP_REGULATION_OF_NEUROINFLAMMATORY_RESPONSE | 22 | 2.21 | 2.25E-05 | 5.15E-04 |
| GOBP_RESPONSE_TO_ENDOPLASMIC_RETICULUM_STRESS | 260 | 2.21 | 3.82E-13 | 1.54E-10 |
| GOBP_REGULATION_OF_VIRAL_PROCESS | 123 | 2.21 | 1.78E-09 | 1.98E-07 |
| GOBP_REGULATION_OF_LEUKOCYTE_CHEMOTAXIS | 91 | 2.21 | 2.25E-07 | 1.21E-05 |
| GOBP_MATURATION_OF_SSU_RRNA | 54 | 2.20 | 6.31E-06 | 1.78E-04 |
| GOBP_COMPLEMENT_ACTIVATION_CLASSICAL_PATHWAY | 16 | 2.20 | 2.80E-05 | 6.05E-04 |
| GOBP_RIBONUCLEOPROTEIN_COMPLEX_BIOGENESIS | 462 | 2.19 | 4.98E-17 | 1.03E-13 |
| GOBP_MIRNA_TRANSCRIPTION | 70 | 2.18 | 3.44E-07 | 1.68E-05 |
| GOBP_MATURATION_OF_5_8S_RRNA | 34 | 2.18 | 1.34E-05 | 3.41E-04 |
| GOBP_REGULATION_OF_VASCULATURE_DEVELOPMENT | 243 | 2.18 | 1.30E-11 | 2.83E-09 |
| GOBP_OSTEOBLAST_DIFFERENTIATION | 211 | 2.18 | 2.73E-10 | 3.72E-08 |
| GOBP_NEGATIVE_REGULATION_OF_INFLAMMATORY_RESPONSE_TO_ANTIGENIC_STIMULUS | 19 | 2.18 | 7.76E-05 | 1.42E-03 |
| GOBP_RESPONSE_TO_GAMMA_RADIATION | 49 | 2.18 | 5.89E-06 | 1.68E-04 |
| GOBP_RNA_TEMPLATED_DNA_BIOSYNTHETIC_PROCESS | 59 | 2.17 | 2.45E-06 | 8.61E-05 |
| GOBP_NEUTROPHIL_CHEMOTAXIS | 52 | 2.17 | 5.62E-06 | 1.63E-04 |
| GOBP_AROMATIC_AMINO_ACID_METABOLIC_PROCESS | 25 | 2.17 | 7.41E-05 | 1.37E-03 |
| GOBP_PROTEIN_FOLDING | 202 | 2.16 | 2.55E-10 | 3.57E-08 |
| GOBP_PROTEIN_N_LINKED_GLYCOSYLATION | 67 | 2.16 | 7.45E-07 | 3.20E-05 |
| GOBP_POSITIVE_REGULATION_OF_EXTRACELLULAR_MATRIX_ORGANIZATION | 27 | 2.16 | 8.51E-05 | 1.52E-03 |
| GOBP_REGULATION_OF_GRANULOCYTE_CHEMOTAXIS | 40 | 2.15 | 8.72E-06 | 2.37E-04 |
| GOBP_EXTRINSIC_APOPTOTIC_SIGNALING_PATHWAY | 200 | 2.15 | 3.96E-10 | 4.85E-08 |
| GOBP_VIRAL_LIFE_CYCLE | 254 | 2.15 | 1.21E-11 | 2.83E-09 |
| GOBP_OVULATION_CYCLE | 53 | 2.15 | 4.88E-06 | 1.49E-04 |
| GOBP_DISRUPTION_OF_ANATOMICAL_STRUCTURE_IN_ANOTHER_ORGANISM | 47 | 2.15 | 9.17E-06 | 2.47E-04 |
| GOBP_MACROPHAGE_MIGRATION | 43 | 2.15 | 1.03E-05 | 2.70E-04 |
| GOBP_REGULATION_OF_EXECUTION_PHASE_OF_APOPTOSIS | 21 | 2.15 | 4.90E-05 | 1.00E-03 |
| GOBP_PROTEIN_N_LINKED_GLYCOSYLATION_VIA_ASPARAGINE | 41 | 2.15 | 2.22E-05 | 5.09E-04 |
| GOBP_CHRONIC_INFLAMMATORY_RESPONSE | 16 | 2.14 | 7.81E-05 | 1.42E-03 |
| GOBP_HUMORAL_IMMUNE_RESPONSE_MEDIATED_BY_CIRCULATING_IMMUNOGLOBULIN | 23 | 2.14 | 1.04E-04 | 1.79E-03 |
| GOBP_POSITIVE_REGULATION_OF_LEUKOCYTE_MIGRATION | 112 | 2.13 | 7.34E-08 | 4.73E-06 |
| GOBP_DE_NOVO_PROTEIN_FOLDING | 36 | 2.12 | 1.90E-05 | 4.51E-04 |
| GOBP_SYMBIONT_ENTRY_INTO_HOST | 114 | 2.12 | 1.33E-07 | 8.11E-06 |
| GOBP_BIOLOGICAL_PROCESS_INVOLVED_IN_INTERACTION_WITH_HOST | 144 | 2.12 | 2.35E-08 | 1.81E-06 |
| GOBP_INTRINSIC_APOPTOTIC_SIGNALING_PATHWAY_IN_RESPONSE_TO_DNA_DAMAGE | 105 | 2.12 | 1.55E-07 | 9.04E-06 |
| GOBP_PROTEOGLYCAN_METABOLIC_PROCESS | 102 | 2.11 | 2.56E-07 | 1.34E-05 |
| GOBP_GLYCOPROTEIN_METABOLIC_PROCESS | 346 | 2.11 | 4.11E-13 | 1.54E-10 |
| GOBP_REGULATION_OF_OSTEOBLAST_DIFFERENTIATION | 120 | 2.11 | 1.58E-07 | 9.11E-06 |
| GOBP_POSITIVE_REGULATION_OF_CELL_MATRIX_ADHESION | 47 | 2.11 | 1.79E-05 | 4.37E-04 |
| GOBP_REACTIVE_OXYGEN_SPECIES_METABOLIC_PROCESS | 187 | 2.11 | 5.40E-09 | 4.95E-07 |
| GOBP_NEUTROPHIL_MIGRATION | 69 | 2.10 | 2.58E-06 | 8.97E-05 |
| GOBP_HETEROTYPIC_CELL_CELL_ADHESION | 40 | 2.10 | 2.28E-05 | 5.19E-04 |
| GOBP_REGULATION_OF_TRANSLATION_IN_RESPONSE_TO_STRESS | 21 | 2.10 | 1.11E-04 | 1.89E-03 |
| GOBP_REGULATION_OF_MONONUCLEAR_CELL_MIGRATION | 106 | 2.10 | 1.75E-07 | 9.94E-06 |
| GOBP_CELLULAR_RESPONSE_TO_INTERLEUKIN_1 | 76 | 2.09 | 2.85E-06 | 9.46E-05 |
| GOBP_INTRINSIC_APOPTOTIC_SIGNALING_PATHWAY_BY_P53_CLASS_MEDIATOR | 84 | 2.09 | 2.59E-06 | 8.97E-05 |
| GOBP_TELOMERASE_RNA_LOCALIZATION | 19 | 2.09 | 2.66E-04 | 3.80E-03 |
| GOBP_CELLULAR_RESPONSE_TO_FATTY_ACID | 33 | 2.09 | 1.40E-04 | 2.22E-03 |
| GOBP_RESPONSE_TO_INTERLEUKIN_7 | 19 | 2.08 | 2.90E-04 | 4.07E-03 |
| GOBP_MONONUCLEAR_CELL_MIGRATION | 157 | 2.08 | 4.50E-08 | 3.12E-06 |
| GOBP_EPIBOLY | 37 | 2.08 | 5.11E-05 | 1.03E-03 |
| GOBP_DE_NOVO_POST_TRANSLATIONAL_PROTEIN_FOLDING | 31 | 2.08 | 1.26E-04 | 2.07E-03 |
| GOBP_ACUTE_PHASE_RESPONSE | 26 | 2.08 | 1.55E-04 | 2.43E-03 |
| GOBP_NEGATIVE_REGULATION_OF_OXIDATIVE_STRESS_INDUCED_NEURON_INTRINSIC_APOPTOTIC_SIGNALING_PATHWAY | 13 | 2.08 | 1.80E-04 | 2.78E-03 |
| GOBP_RESPONSE_TO_TOPOLOGICALLY_INCORRECT_PROTEIN | 157 | 2.08 | 5.55E-08 | 3.73E-06 |
| GOBP_CELLULAR_RESPONSE_TO_GAMMA_RADIATION | 27 | 2.07 | 2.96E-04 | 4.13E-03 |
| GOBP_TAXIS | 331 | 2.07 | 2.78E-12 | 8.30E-10 |
| GOBP_INFLAMMATORY_RESPONSE_TO_ANTIGENIC_STIMULUS | 42 | 2.07 | 6.49E-05 | 1.22E-03 |
| GOBP_REGULATION_OF_BIOLOGICAL_PROCESS_INVOLVED_IN_SYMBIOTIC_INTERACTION | 33 | 2.07 | 1.88E-04 | 2.89E-03 |
| GOBP_REGULATION_OF_MYELOID_CELL_DIFFERENTIATION | 174 | 2.06 | 3.14E-08 | 2.34E-06 |
| GOBP_POSITIVE_REGULATION_OF_PHAGOCYTOSIS | 41 | 2.06 | 8.36E-05 | 1.49E-03 |
| GOBP_NEGATIVE_REGULATION_OF_OXIDATIVE_STRESS_INDUCED_INTRINSIC_APOPTOTIC_SIGNALING_PATHWAY | 28 | 2.06 | 2.09E-04 | 3.11E-03 |
| GOBP_GLYCOSAMINOGLYCAN_METABOLIC_PROCESS | 69 | 2.06 | 5.35E-06 | 1.57E-04 |
| GOBP_TRANSLATIONAL_INITIATION | 119 | 2.06 | 3.44E-07 | 1.68E-05 |
| GOBP_MATURATION_OF_LSU_RRNA | 26 | 2.06 | 2.01E-04 | 3.04E-03 |
| GOBP_SMOOTH_MUSCLE_CELL_PROLIFERATION | 119 | 2.06 | 3.62E-07 | 1.75E-05 |
| GOBP_NEGATIVE_REGULATION_OF_RESPONSE_TO_ENDOPLASMIC_RETICULUM_STRESS | 48 | 2.05 | 5.74E-05 | 1.13E-03 |
| GOBP_NEGATIVE_REGULATION_OF_VIRAL_PROCESS | 64 | 2.05 | 7.33E-06 | 2.02E-04 |
| GOBP_CRANIAL_SKELETAL_SYSTEM_DEVELOPMENT | 62 | 2.05 | 1.38E-05 | 3.48E-04 |
| GOBP_GLYCOPROTEIN_BIOSYNTHETIC_PROCESS | 286 | 2.05 | 2.16E-10 | 3.13E-08 |
| GOBP_REGULATION_OF_EPITHELIAL_TO_MESENCHYMAL_TRANSITION | 95 | 2.05 | 2.02E-06 | 7.31E-05 |
| GOBP_NEURON_INTRINSIC_APOPTOTIC_SIGNALING_PATHWAY_IN_RESPONSE_TO_OXIDATIVE_STRESS | 20 | 2.04 | 6.18E-04 | 7.28E-03 |

| | | | | |
|---|---|---|---|---|
| GOBP_AMINOGLYCAN_METABOLIC_PROCESS | 79 | 2.04 | 5.63E-06 | 1.63E-04 |
| GOBP_AMINO_ACID_METABOLIC_PROCESS | 237 | 2.04 | 2.55E-09 | 2.53E-07 |
| GOBP_ENDOTHELIAL_CELL_APOPTOTIC_PROCESS | 51 | 2.04 | 5.05E-05 | 1.03E-03 |
| GOBP_REGULATION_OF_VIRAL_LIFE_CYCLE | 100 | 2.04 | 2.76E-06 | 9.35E-05 |
| GOBP_FORMATION_OF_CYTOPLASMIC_TRANSLATION_INITIATION_COMPLEX | 15 | 2.04 | 2.02E-04 | 3.05E-03 |
| GOBP_POSITIVE_REGULATION_OF_MONONUCLEAR_CELL_MIGRATION | 70 | 2.03 | 9.94E-06 | 2.65E-04 |
| GOBP_WOUND_HEALING_SPREADING_OF_EPIDERMAL_CELLS | 22 | 2.03 | 2.72E-04 | 3.86E-03 |
| GOBP_REGULATION_OF_CALCIUM_ION_IMPORT | 30 | 2.03 | 2.03E-04 | 3.06E-03 |
| GOBP_REGULATION_OF_MACROPHAGE_CHEMOTAXIS | 25 | 2.03 | 4.38E-04 | 5.61E-03 |
| GOBP_POSITIVE_REGULATION_OF_CANONICAL_NF_KAPPAB_SIGNAL_TRANSDUCTION | 196 | 2.03 | 1.42E-08 | 1.19E-06 |
| GOBP_NEGATIVE_REGULATION_OF_SIGNAL_TRANSDUCTION_BY_P53_CLASS_MEDIATOR | 39 | 2.02 | 1.24E-04 | 2.04E-03 |
| GOBP_POSITIVE_REGULATION_OF_CHEMOTAXIS | 110 | 2.02 | 1.67E-06 | 6.29E-05 |
| GOBP_REGULATION_OF_REACTIVE_OXYGEN_SPECIES_METABOLIC_PROCESS | 118 | 2.02 | 1.87E-06 | 6.86E-05 |
| GOBP_CELL_ADHESION_MEDIATED_BY_INTEGRIN | 71 | 2.02 | 5.11E-06 | 1.54E-04 |
| GOBP_POSITIVE_REGULATION_OF_MYELOID_CELL_DIFFERENTIATION | 87 | 2.02 | 7.42E-06 | 2.04E-04 |
| GOBP_VASCULAR_ENDOTHELIAL_GROWTH_FACTOR_PRODUCTION | 33 | 2.02 | 3.20E-04 | 4.37E-03 |
| GOBP_POSITIVE_REGULATION_OF_LOCOMOTION | 492 | 2.02 | 9.24E-14 | 4.41E-11 |
| GOBP_POSITIVE_REGULATION_OF_MIRNA_METABOLIC_PROCESS | 60 | 2.02 | 1.87E-05 | 4.50E-04 |
| GOBP_REGULATION_OF_CELL_MATRIX_ADHESION | 109 | 2.02 | 9.22E-07 | 3.76E-05 |
| GOBP_CELLULAR_RESPONSE_TO_CHEMICAL_STRESS | 296 | 2.02 | 1.72E-10 | 2.56E-08 |
| GOBP_REGULATION_OF_CELL_ADHESION_MEDIATED_BY_INTEGRIN | 40 | 2.01 | 7.92E-05 | 1.43E-03 |
| GOBP_BRANCH_ELONGATION_OF_AN_EPITHELIUM | 17 | 2.01 | 3.60E-04 | 4.79E-03 |
| GOBP_TRANSFORMING_GROWTH_FACTOR_BETA_RECEPTOR_SIGNALING_PATHWAY | 207 | 2.01 | 4.05E-08 | 2.84E-06 |
| GOBP_REGULATION_OF_MYELOID_LEUKOCYTE_DIFFERENTIATION | 96 | 2.00 | 2.83E-06 | 9.46E-05 |
| GOBP_REGULATION_OF_PROTEIN_LOCALIZATION_TO_NUCLEUS | 134 | 2.00 | 4.50E-07 | 2.08E-05 |
| GOBP_GLAND_MORPHOGENESIS | 115 | 2.00 | 2.18E-06 | 7.76E-05 |
| GOBP_REGULATION_OF_EXTRINSIC_APOPTOTIC_SIGNALING_PATHWAY | 137 | 2.00 | 3.19E-07 | 1.60E-05 |
| GOBP_REGULATION_OF_SMAD_PROTEIN_SIGNAL_TRANSDUCTION | 60 | 2.00 | 2.59E-05 | 5.73E-04 |
| GOBP_ANTRAL_OVARIAN_FOLLICLE_GROWTH | 10 | 2.00 | 3.10E-04 | 4.27E-03 |
| GOBP_RIBOSOMAL_LARGE_SUBUNIT_BIOGENESIS | 66 | 2.00 | 2.10E-05 | 4.83E-04 |
| GOBP_REGULATION_OF_MACROPHAGE_MIGRATION | 34 | 1.99 | 2.05E-04 | 3.07E-03 |
| GOBP_CHONDROITIN_SULFATE_PROTEOGLYCAN_BIOSYNTHETIC_PROCESS | 22 | 1.99 | 4.98E-04 | 6.25E-03 |
| GOBP_NEGATIVE_REGULATION_OF_VIRAL_TRANSCRIPTION | 15 | 1.99 | 4.10E-04 | 5.35E-03 |
| GOBP_CHONDROITIN_SULFATE_PROTEOGLYCAN_METABOLIC_PROCESS | 27 | 1.99 | 7.39E-04 | 8.20E-03 |
| GOBP_MODULATION_BY_SYMBIONT_OF_ENTRY_INTO_HOST | 27 | 1.99 | 7.97E-04 | 8.63E-03 |
| GOBP_NEGATIVE_REGULATION_OF_PROTEIN_MATURATION | 18 | 1.99 | 5.59E-04 | 6.70E-03 |
| GOBP_MAMMARY_GLAND_DUCT_MORPHOGENESIS | 31 | 1.99 | 4.65E-04 | 5.89E-03 |
| GOBP_CHEMOKINE_PRODUCTION | 69 | 1.99 | 1.95E-05 | 4.55E-04 |
| GOBP_POSITIVE_REGULATION_OF_VASCULAR_ENDOTHELIAL_GROWTH_FACTOR_PRODUCTION | 29 | 1.98 | 6.44E-04 | 7.49E-03 |
| GOBP_INTRINSIC_APOPTOTIC_SIGNALING_PATHWAY_IN_RESPONSE_TO_DNA_DAMAGE_BY_P53_CLASS_MEDIATOR | 47 | 1.98 | 1.18E-04 | 1.98E-03 |
| GOBP_MATURATION_OF_SSU_RRNA_FROM_TRICISTRONIC_RRNA_TRANSCRIPT_SSU_RRNA_5_8S_RRNA_LSU_RRNA | 35 | 1.98 | 5.31E-04 | 6.51E-03 |
| GOBP_POSITIVE_REGULATION_OF_PROTEIN_LOCALIZATION_TO_CAJAL_BODY | 10 | 1.98 | 4.11E-04 | 5.35E-03 |
| GOBP_RESPONSE_TO_OXYGEN_LEVELS | 306 | 1.98 | 3.38E-10 | 4.36E-08 |
| GOBP_REGULATION_OF_PHAGOCYTOSIS | 62 | 1.98 | 5.50E-05 | 1.09E-03 |
| GOBP_NEGATIVE_REGULATION_OF_BLOOD_PRESSURE | 28 | 1.97 | 6.86E-04 | 7.79E-03 |
| GOBP_BIOLOGICAL_PROCESS_INVOLVED_IN_INTERACTION_WITH_SYMBIONT | 18 | 1.97 | 6.58E-04 | 7.57E-03 |
| GOBP_PLASMINOGEN_ACTIVATION | 20 | 1.97 | 1.88E-03 | 1.66E-02 |
| GOBP_REGULATION_OF_TRANSLATIONAL_INITIATION_IN_RESPONSE_TO_STRESS | 11 | 1.97 | 5.86E-04 | 6.98E-03 |
| GOBP_REGULATION_OF_VASCULAR_ENDOTHELIAL_GROWTH_FACTOR_RECEPTOR_SIGNALING_PATHWAY | 29 | 1.97 | 7.75E-04 | 8.48E-03 |
| GOBP_POSITIVE_REGULATION_OF_EXECUTION_PHASE_OF_APOPTOSIS | 18 | 1.97 | 6.88E-04 | 7.79E-03 |
| GOBP_NEGATIVE_REGULATION_OF_PROTEOLYSIS | 120 | 1.96 | 3.98E-06 | 1.25E-04 |
| GOBP_POSITIVE_REGULATION_OF_LEUKOCYTE_CHEMOTAXIS | 70 | 1.96 | 4.32E-05 | 9.00E-04 |
| GOBP_POSITIVE_REGULATION_OF_MYELOID_LEUKOCYTE_DIFFERENTIATION | 48 | 1.96 | 2.10E-04 | 3.12E-03 |
| GOBP_HEPARAN_SULFATE_PROTEOGLYCAN_BIOSYNTHETIC_PROCESS | 34 | 1.96 | 3.46E-04 | 4.65E-03 |
| GOBP_INTRINSIC_APOPTOTIC_SIGNALING_PATHWAY_IN_RESPONSE_TO_OXIDATIVE_STRESS | 56 | 1.96 | 1.27E-04 | 2.07E-03 |
| GOBP_PROTEOGLYCAN_BIOSYNTHETIC_PROCESS | 78 | 1.96 | 2.41E-05 | 5.37E-04 |
| GOBP_REGULATION_OF_EXTRACELLULAR_MATRIX_ASSEMBLY | 14 | 1.96 | 5.27E-04 | 6.48E-03 |
| GOBP_ESTABLISHMENT_OF_PROTEIN_LOCALIZATION_TO_PEROXISOME | 20 | 1.96 | 2.10E-03 | 1.81E-02 |
| GOBP_ENDODERM_DEVELOPMENT | 76 | 1.95 | 3.65E-05 | 7.81E-04 |
| GOBP_BLOOD_VESSEL_ENDOTHELIAL_CELL_PROLIFERATION_INVOLVED_IN_SPROUTING_ANGIOGENESIS | 22 | 1.95 | 8.07E-04 | 8.71E-03 |
| GOBP_CELLULAR_RESPONSE_TO_UNFOLDED_PROTEIN | 96 | 1.95 | 6.76E-06 | 1.89E-04 |
| GOBP_NADPH_REGENERATION | 23 | 1.95 | 1.43E-03 | 1.35E-02 |
| GOBP_PROTEIN_PROCESSING | 191 | 1.95 | 3.21E-07 | 1.60E-05 |
| GOBP_ENDODERM_FORMATION | 50 | 1.95 | 1.16E-04 | 1.95E-03 |
| GOBP_ENDOTHELIAL_CELL_PROLIFERATION | 142 | 1.95 | 1.93E-06 | 7.01E-05 |
| GOBP_POSITIVE_REGULATION_OF_FIBROBLAST_PROLIFERATION | 48 | 1.95 | 2.70E-04 | 3.85E-03 |
| GOBP_RESPONSE_TO_INTERLEUKIN_17 | 16 | 1.95 | 1.25E-03 | 1.22E-02 |
| GOBP_POSITIVE_REGULATION_OF_PROTEIN_LOCALIZATION_TO_NUCLEUS | 86 | 1.94 | 1.91E-05 | 4.51E-04 |
| GOBP_POSITIVE_REGULATION_OF_VASCULAR_ASSOCIATED_SMOOTH_MUSCLE_CELL_PROLIFERATION | 31 | 1.94 | 6.18E-04 | 7.28E-03 |
| GOBP_RESPONSE_TO_WOUNDING | 459 | 1.94 | 1.40E-11 | 2.90E-09 |
| GOBP_REGULATION_OF_CELL_SUBSTRATE_ADHESION | 191 | 1.94 | 3.75E-07 | 1.79E-05 |
| GOBP_GASTRULATION | 166 | 1.94 | 5.35E-07 | 2.39E-05 |
| GOBP_CELL_MATRIX_ADHESION | 200 | 1.94 | 2.62E-07 | 1.36E-05 |
| GOBP_EPITHELIAL_CELL_PROLIFERATION_INVOLVED_IN_PROSTATE_GLAND_DEVELOPMENT | 10 | 1.94 | 8.46E-04 | 9.03E-03 |
| GOBP_ODONTOBLAST_DIFFERENTIATION | 17 | 1.94 | 9.17E-04 | 9.70E-03 |
| GOBP_MYELOID_CELL_DIFFERENTIATION | 371 | 1.94 | 2.85E-10 | 3.78E-08 |
| GOBP_PROTEIN_LOCALIZATION_TO_NUCLEUS | 304 | 1.94 | 4.64E-09 | 4.34E-07 |
| GOBP_RESPONSE_TO_MOLECULE_OF_BACTERIAL_ORIGIN | 259 | 1.93 | 1.59E-08 | 1.31E-06 |
| GOBP_REGULATION_OF_MONOCYTE_CHEMOTAXIS | 19 | 1.93 | 1.79E-03 | 1.60E-02 |
| GOBP_NEGATIVE_REGULATION_OF_ENDOPLASMIC_RETICULUM_STRESS_INDUCED_INTRINSIC_APOPTOTIC_SIGNALING_PATHWAY | 20 | 1.93 | 2.70E-03 | 2.17E-02 |
| GOBP_NEGATIVE_REGULATION_OF_NITRIC_OXIDE_METABOLIC_PROCESS | 12 | 1.93 | 1.16E-03 | 1.15E-02 |
| GOBP_IRE1_MEDIATED_UNFOLDED_PROTEIN_RESPONSE | 20 | 1.93 | 2.70E-03 | 2.17E-02 |
| GOBP_CANONICAL_NF_KAPPAB_SIGNAL_TRANSDUCTION | 285 | 1.93 | 2.51E-09 | 2.53E-07 |
| GOBP_REGULATION_OF_RELEASE_OF_CYTOCHROME_C_FROM_MITOCHONDRIA | 39 | 1.93 | 4.50E-04 | 5.73E-03 |
| GOBP_POSITIVE_REGULATION_OF_OSTEOBLAST_DIFFERENTIATION | 66 | 1.93 | 5.77E-05 | 1.13E-03 |
| GOBP_REGULATION_OF_INFLAMMATORY_RESPONSE | 285 | 1.93 | 2.51E-09 | 2.53E-07 |
| GOBP_RESPONSE_TO_INTERLEUKIN_1 | 96 | 1.93 | 1.04E-05 | 2.70E-04 |
| GOBP_FORMATION_OF_TRANSLATION_PREINITIATION_COMPLEX | 10 | 1.93 | 9.62E-04 | 1.00E-02 |
| GOBP_MYELOID_CELL_HOMEOSTASIS | 162 | 1.93 | 7.85E-07 | 3.28E-05 |
| GOBP_ZYMOGEN_ACTIVATION | 34 | 1.93 | 5.22E-04 | 6.46E-03 |
| GOBP_POSITIVE_REGULATION_OF_MACROPHAGE_CHEMOTAXIS | 18 | 1.93 | 1.26E-03 | 1.23E-02 |
| GOBP_REGULATION_OF_TELOMERASE_RNA_LOCALIZATION_TO_CAJAL_BODY | 12 | 1.93 | 1.30E-03 | 1.26E-02 |
| GOBP_REGULATION_OF_PLASMINOGEN_ACTIVATION | 17 | 1.93 | 1.12E-03 | 1.13E-02 |
| GOBP_AMP_METABOLIC_PROCESS | 20 | 1.93 | 2.90E-03 | 2.29E-02 |
| GOBP_NEGATIVE_REGULATION_OF_EXTRINSIC_APOPTOTIC_SIGNALING_PATHWAY | 84 | 1.92 | 5.38E-05 | 1.08E-03 |
| GOBP_PROTEIN_MATURATION | 491 | 1.92 | 1.02E-11 | 2.69E-09 |
| GOBP_OUTFLOW_TRACT_SEPTUM_MORPHOGENESIS | 27 | 1.92 | 1.51E-03 | 1.40E-02 |
| GOBP_FOLIC_ACID_CONTAINING_COMPOUND_METABOLIC_PROCESS | 25 | 1.92 | 1.75E-03 | 1.57E-02 |
| GOBP_RESPONSE_TO_MANGANESE_ION | 15 | 1.92 | 1.06E-03 | 1.09E-02 |
| GOBP_CELLULAR_RESPONSE_TO_MANGANESE_ION | 10 | 1.92 | 1.10E-03 | 1.12E-02 |
| GOBP_REGULATION_OF_VIRAL_ENTRY_INTO_HOST_CELL | 21 | 1.92 | 1.13E-03 | 1.14E-02 |
| GOBP_HEPARIN_PROTEOGLYCAN_BIOSYNTHETIC_PROCESS | 11 | 1.92 | 1.48E-03 | 1.38E-02 |
| GOBP_CELLULAR_RESPONSE_TO_BIOTIC_STIMULUS | 183 | 1.91 | 1.86E-06 | 6.86E-05 |
| GOBP_REGULATION_OF_CARDIAC_EPITHELIAL_TO_MESENCHYMAL_TRANSITION | 12 | 1.91 | 1.42E-03 | 1.35E-02 |
| GOBP_MORPHOGENESIS_OF_AN_EPITHELIAL_SHEET | 59 | 1.91 | 1.11E-04 | 1.89E-03 |
| GOBP_REGULATION_OF_CHEMOTAXIS | 160 | 1.91 | 3.54E-06 | 1.13E-04 |

| | | | | |
|---|---|---|---|---|
| GOBP_AROMATIC_AMINO_ACID_FAMILY_CATABOLIC_PROCESS | 15 | 1.91 | 1.14E-03 | 1.14E-02 |
| GOBP_CELL_CELL_ADHESION_MEDIATED_BY_INTEGRIN | 16 | 1.91 | 1.63E-03 | 1.49E-02 |
| GOBP_SYNCYTIUM_FORMATION | 45 | 1.91 | 3.60E-04 | 4.79E-03 |
| GOBP_SMAD_PROTEIN_SIGNAL_TRANSDUCTION | 78 | 1.91 | 4.77E-05 | 9.82E-04 |
| GOBP_RESPONSE_TO_LIPOPROTEIN_PARTICLE | 27 | 1.91 | 1.78E-03 | 1.59E-02 |
| GOBP_REGULATION_OF_MIRNA_METABOLIC_PROCESS | 84 | 1.91 | 6.59E-05 | 1.23E-03 |
| GOBP_WOUND_HEALING | 336 | 1.91 | 1.71E-09 | 1.98E-07 |
| GOBP_REGULATION_OF_VIRAL_GENOME_REPLICATION | 68 | 1.91 | 2.29E-04 | 3.36E-03 |
| GOBP_POSITIVE_REGULATION_OF_PROGRAMMED_CELL_DEATH | 468 | 1.90 | 3.40E-11 | 6.36E-09 |
| GOBP_ALPHA_AMINO_ACID_BIOSYNTHETIC_PROCESS | 49 | 1.90 | 3.84E-04 | 5.06E-03 |
| GOBP_REGULATION_OF_MORPHOGENESIS_OF_AN_EPITHELIUM | 60 | 1.90 | 1.11E-04 | 1.89E-03 |
| GOBP_REGULATION_OF_COMPLEMENT_ACTIVATION | 13 | 1.90 | 1.42E-03 | 1.35E-02 |
| GOBP_NEGATIVE_REGULATION_OF_LEUKOCYTE_MIGRATION | 38 | 1.90 | 7.98E-04 | 8.63E-03 |
| GOBP_REGULATION_OF_SULFUR_METABOLIC_PROCESS | 16 | 1.90 | 1.75E-03 | 1.57E-02 |
| GOBP_REGULATION_OF_TELOMERE_MAINTENANCE_VIA_TELOMERASE | 45 | 1.90 | 3.93E-04 | 5.16E-03 |
| GOBP_POSITIVE_REGULATION_OF_ENDOTHELIAL_CELL_MIGRATION | 98 | 1.90 | 1.89E-05 | 4.51E-04 |
| GOBP_LEUKOCYTE_APOPTOTIC_PROCESS | 98 | 1.90 | 1.92E-05 | 4.51E-04 |
| GOBP_VIRAL_GENOME_REPLICATION | 107 | 1.90 | 1.48E-05 | 3.68E-04 |
| GOBP_LEUKOCYTE_ACTIVATION_INVOLVED_IN_INFLAMMATORY_RESPONSE | 30 | 1.90 | 9.19E-04 | 9.71E-03 |
| GOBP_REGULATION_OF_HUMORAL_IMMUNE_RESPONSE | 21 | 1.90 | 1.45E-03 | 1.36E-02 |
| GOBP_CELL_SURFACE_RECEPTOR_PROTEIN_SERINE_THREONINE_KINASE_SIGNALING_PATHWAY | 347 | 1.90 | 2.59E-09 | 2.53E-07 |
| GOBP_RENAL_SYSTEM_DEVELOPMENT | 289 | 1.90 | 3.45E-08 | 2.52E-06 |
| GOBP_REGULATION_OF_INTRINSIC_APOPTOTIC_SIGNALING_PATHWAY_BY_P53_CLASS_MEDIATOR | 34 | 1.89 | 7.43E-04 | 8.22E-03 |
| GOBP_REGULATION_OF_TRANSLATION_IN_RESPONSE_TO_ENDOPLASMIC_RETICULUM_STRESS | 10 | 1.89 | 1.56E-03 | 1.45E-02 |
| GOBP_BRANCHING_MORPHOGENESIS_OF_AN_EPITHELIAL_TUBE | 145 | 1.89 | 5.18E-06 | 1.56E-04 |
| GOBP_ERAD_PATHWAY | 105 | 1.89 | 2.79E-05 | 6.05E-04 |
| GOBP_NEGATIVE_REGULATION_OF_PROTEOLYSIS_INVOLVED_IN_PROTEIN_CATABOLIC_PROCESS | 72 | 1.89 | 1.31E-04 | 2.11E-03 |
| GOBP_IMMUNE_EFFECTOR_PROCESS | 442 | 1.89 | 1.63E-10 | 2.50E-08 |
| GOBP_CELL_SUBSTRATE_ADHESION | 307 | 1.89 | 2.81E-08 | 2.13E-06 |
| GOBP_AMYLOID_BETA_CLEARANCE | 26 | 1.89 | 1.77E-03 | 1.58E-02 |
| GOBP_POSITIVE_REGULATION_OF_ASTROCYTE_DIFFERENTIATION | 10 | 1.89 | 1.70E-03 | 1.54E-02 |
| GOBP_EXTERNAL_ENCAPSULATING_STRUCTURE_ORGANIZATION | 285 | 1.89 | 1.01E-08 | 8.96E-07 |
| GOBP_POSITIVE_REGULATION_OF_CHEMOKINE_PRODUCTION | 50 | 1.88 | 3.29E-04 | 4.46E-03 |
| GOBP_MORPHOGENESIS_OF_A_BRANCHING_STRUCTURE | 186 | 1.88 | 1.30E-06 | 5.04E-05 |
| GOBP_ANTIGEN_PROCESSING_AND_PRESENTATION_OF_PEPTIDE_OR_POLYSACCHARIDE_ANTIGEN_VIA_MHC_CLASS_II | 26 | 1.88 | 1.89E-03 | 1.67E-02 |
| GOBP_EPITHELIAL_CELL_PROLIFERATION | 371 | 1.88 | 2.48E-09 | 2.53E-07 |
| GOBP_MAMMARY_GLAND_MORPHOGENESIS | 44 | 1.88 | 3.91E-04 | 5.14E-03 |
| GOBP_POSITIVE_CHEMOTAXIS | 48 | 1.88 | 6.58E-04 | 7.57E-03 |
| GOBP_NEGATIVE_REGULATION_OF_STRIATED_MUSCLE_CELL_DIFFERENTIATION | 33 | 1.88 | 1.71E-03 | 1.55E-02 |
| GOBP_REGULATION_OF_DEFENSE_RESPONSE_TO_VIRUS_BY_HOST | 37 | 1.88 | 8.73E-04 | 9.26E-03 |
| GOBP_CELL_SUBSTRATE_JUNCTION_ORGANIZATION | 97 | 1.88 | 5.85E-05 | 1.14E-03 |
| GOBP_REGULATION_OF_SUPEROXIDE_METABOLIC_PROCESS | 20 | 1.88 | 4.62E-03 | 3.14E-02 |
| GOBP_RESPONSE_TO_VIRUS | 346 | 1.88 | 3.22E-09 | 3.07E-07 |
| GOBP_POSITIVE_REGULATION_OF_PEPTIDYL_SERINE_PHOSPHORYLATION | 19 | 1.88 | 2.66E-03 | 2.16E-02 |
| GOBP_LOW_DENSITY_LIPOPROTEIN_PARTICLE_CLEARANCE | 23 | 1.88 | 3.00E-03 | 2.34E-02 |
| GOBP_NADP_METABOLIC_PROCESS | 38 | 1.88 | 1.07E-03 | 1.10E-02 |
| GOBP_SULFUR_COMPOUND_BIOSYNTHETIC_PROCESS | 150 | 1.88 | 4.22E-06 | 1.32E-04 |
| GOBP_MATURATION_OF_5_8S_RRNA_FROM_TRICISTRONIC_RRNA_TRANSCRIPT_SSU_RRNA_5_8S_RRNA_LSU_RRNA | 23 | 1.88 | 3.12E-03 | 2.40E-02 |
| GOBP_SUPPRESSION_OF_VIRAL_RELEASE_BY_HOST | 16 | 1.88 | 2.56E-03 | 2.09E-02 |
| GOBP_VIRAL_GENE_EXPRESSION | 102 | 1.88 | 2.38E-05 | 5.33E-04 |
| GOBP_CELLULAR_RESPONSE_TO_OXIDATIVE_STRESS | 232 | 1.88 | 1.90E-07 | 1.06E-05 |
| GOBP_MACROPHAGE_ACTIVATION | 73 | 1.87 | 1.07E-04 | 1.84E-03 |
| GOBP_MYELOID_LEUKOCYTE_ACTIVATION | 163 | 1.87 | 5.38E-06 | 1.57E-04 |
| GOBP_RESPONSE_TO_TRANSFORMING_GROWTH_FACTOR_BETA | 260 | 1.87 | 7.19E-08 | 4.70E-06 |
| GOBP_RESPONSE_TO_EPIDERMAL_GROWTH_FACTOR | 50 | 1.87 | 3.81E-04 | 5.04E-03 |
| GOBP_MUSCLE_CELL_PROLIFERATION | 181 | 1.87 | 2.14E-06 | 7.66E-05 |
| GOBP_MYELOID_LEUKOCYTE_DIFFERENTIATION | 196 | 1.87 | 1.09E-06 | 4.36E-05 |
| GOBP_ICOSANOID_METABOLIC_PROCESS | 69 | 1.87 | 1.33E-04 | 2.13E-03 |
| GOBP_SIGNAL_TRANSDUCTION_BY_P53_CLASS_MEDIATOR | 182 | 1.87 | 3.78E-06 | 1.20E-04 |
| GOBP_ACUTE_INFLAMMATORY_RESPONSE | 62 | 1.87 | 2.56E-04 | 3.71E-03 |
| GOBP_TRAIL_ACTIVATED_APOPTOTIC_SIGNALING_PATHWAY | 11 | 1.87 | 2.41E-03 | 2.00E-02 |
| GOBP_ALDEHYDE_METABOLIC_PROCESS | 65 | 1.87 | 1.65E-04 | 2.58E-03 |
| GOBP_MESENCHYME_MORPHOGENESIS | 56 | 1.87 | 6.29E-04 | 7.37E-03 |
| GOBP_GLYCOSYLATION | 222 | 1.87 | 1.24E-06 | 4.87E-05 |
| GOBP_MIRNA_METABOLIC_PROCESS | 101 | 1.87 | 2.77E-05 | 6.05E-04 |
| GOBP_POSITIVE_REGULATION_OF_DEFENSE_RESPONSE_TO_VIRUS_BY_HOST | 28 | 1.87 | 2.49E-03 | 2.05E-02 |
| GOBP_FOAM_CELL_DIFFERENTIATION | 25 | 1.87 | 3.47E-03 | 2.61E-02 |
| GOBP_HEPARIN_PROTEOGLYCAN_METABOLIC_PROCESS | 17 | 1.86 | 2.69E-03 | 2.17E-02 |
| GOBP_REGULATION_OF_PROTEIN_IMPORT_INTO_NUCLEUS | 56 | 1.86 | 6.87E-04 | 7.79E-03 |
| GOBP_MODIFIED_AMINO_ACID_BIOSYNTHETIC_PROCESS | 42 | 1.86 | 8.65E-04 | 9.20E-03 |
| GOBP_POSITIVE_REGULATION_OF_SYSTEMIC_ARTERIAL_BLOOD_PRESSURE | 12 | 1.86 | 2.85E-03 | 2.26E-02 |
| GOBP_POSITIVE_REGULATION_OF_SMOOTH_MUSCLE_CELL_MIGRATION | 33 | 1.86 | 1.86E-03 | 1.65E-02 |
| GOBP_RESPONSE_TO_CHEMOKINE | 50 | 1.86 | 4.25E-04 | 5.48E-03 |
| GOBP_POSITIVE_REGULATION_OF_BLOOD_PRESSURE | 20 | 1.86 | 5.10E-03 | 3.41E-02 |
| GOBP_RESPONSE_TO_BACTERIUM | 436 | 1.86 | 1.31E-09 | 1.56E-07 |
| GOBP_AMINO_ACID_BIOSYNTHETIC_PROCESS | 60 | 1.85 | 2.48E-04 | 3.61E-03 |
| GOBP_PRODUCTION_OF_MOLECULAR_MEDIATOR_OF_IMMUNE_RESPONSE | 171 | 1.85 | 5.79E-06 | 1.67E-04 |
| GOBP_EOSINOPHIL_CHEMOTAXIS | 10 | 1.85 | 2.79E-03 | 2.23E-02 |
| GOBP_REGULATION_OF_PEPTIDYL_SERINE_PHOSPHORYLATION | 27 | 1.85 | 2.91E-03 | 2.29E-02 |
| GOBP_REGULATION_OF_VIRAL_TRANSCRIPTION | 19 | 1.85 | 3.67E-03 | 2.70E-02 |
| GOBP_KERATINOCYTE_PROLIFERATION | 46 | 1.85 | 5.12E-04 | 6.37E-03 |
| GOBP_ODONTOGENESIS | 102 | 1.85 | 4.24E-05 | 8.87E-04 |
| GOBP_NEGATIVE_REGULATION_OF_PROTEASOMAL_PROTEIN_CATABOLIC_PROCESS | 57 | 1.85 | 4.76E-04 | 6.01E-03 |
| GOBP_NEGATIVE_REGULATION_OF_RELEASE_OF_CYTOCHROME_C_FROM_MITOCHONDRIA | 20 | 1.85 | 5.42E-03 | 3.57E-02 |
| GOBP_REGULATION_OF_MITOCHONDRIAL_FUSION | 15 | 1.85 | 2.73E-03 | 2.19E-02 |
| GOBP_NEGATIVE_REGULATION_OF_ENDOTHELIAL_CELL_PROLIFERATION | 34 | 1.85 | 1.11E-03 | 1.12E-02 |
| GOBP_MUSCLE_CELL_CELLULAR_HOMEOSTASIS | 21 | 1.85 | 2.15E-03 | 1.83E-02 |
| GOBP_NEGATIVE_REGULATION_OF_PROTEIN_METABOLIC_PROCESS | 500 | 1.85 | 5.15E-11 | 8.78E-09 |
| GOBP_MONOCYTE_CHEMOTAXIS | 31 | 1.85 | 1.60E-03 | 1.48E-02 |
| GOBP_ASPARTATE_FAMILY_AMINO_ACID_BIOSYNTHETIC_PROCESS | 14 | 1.85 | 2.33E-03 | 1.95E-02 |
| GOBP_RESPONSE_TO_FIBROBLAST_GROWTH_FACTOR | 102 | 1.85 | 4.42E-05 | 9.17E-04 |
| GOBP_POSITIVE_REGULATION_OF_APOPTOTIC_SIGNALING_PATHWAY | 130 | 1.85 | 2.78E-05 | 6.05E-04 |
| GOBP_NEGATIVE_REGULATION_BY_HOST_OF_VIRAL_PROCESS | 12 | 1.85 | 3.49E-03 | 2.61E-02 |
| GOBP_POSITIVE_REGULATION_OF_RECEPTOR_CLUSTERING | 11 | 1.85 | 3.68E-03 | 2.70E-02 |
| GOBP_REGULATION_OF_EPITHELIAL_CELL_PROLIFERATION | 302 | 1.84 | 1.24E-07 | 7.81E-06 |
| GOBP_REGULATION_OF_GLYCOPROTEIN_METABOLIC_PROCESS | 37 | 1.84 | 1.44E-03 | 1.36E-02 |
| GOBP_POSITIVE_REGULATION_OF_TRANSLATION | 122 | 1.84 | 1.81E-05 | 4.42E-04 |
| GOBP_OSSIFICATION | 370 | 1.84 | 1.32E-08 | 1.12E-06 |
| GOBP_MATURATION_OF_LSU_RRNA_FROM_TRICISTRONIC_RRNA_TRANSCRIPT_SSU_RRNA_5_8S_RRNA_LSU_RRNA | 17 | 1.84 | 3.47E-03 | 2.61E-02 |
| GOBP_GLYCOLIPID_CATABOLIC_PROCESS | 17 | 1.84 | 3.51E-03 | 2.62E-02 |
| GOBP_POSITIVE_REGULATION_OF_CELL_ADHESION_MEDIATED_BY_INTEGRIN | 15 | 1.84 | 2.97E-03 | 2.33E-02 |
| GOBP_REGULATION_OF_GLIAL_CELL_MIGRATION | 12 | 1.84 | 3.89E-03 | 2.79E-02 |
| GOBP_REGULATION_OF_MACROPHAGE_ACTIVATION | 42 | 1.84 | 1.19E-03 | 1.18E-02 |
| GOBP_ADAPTIVE_IMMUNE_RESPONSE_BASED_ON_SOMATIC_RECOMBINATION_OF_IMMUNE_RECEPTORS_BUILT_FROM_IMMUNOGLOBULIN_SUPERFAN | 227 | 1.84 | 5.70E-07 | 2.49E-05 |
| GOBP_PROTEIN_FOLDING_IN_ENDOPLASMIC_RETICULUM | 12 | 1.84 | 4.01E-03 | 2.86E-02 |
| GOBP_NUCLEAR_PORE_ORGANIZATION | 15 | 1.84 | 3.01E-03 | 2.34E-02 |

| | | | | |
|---|---|---|---|---|
| GOBP_B_CELL_MEDIATED_IMMUNITY | 85 | 1.84 | 1.21E-04 | 2.02E-03 |
| GOBP_FORMATION_OF_PRIMARY_GERM_LAYER | 111 | 1.84 | 9.51E-05 | 1.67E-03 |
| GOBP_NEGATIVE_REGULATION_OF_MYELOID_LEUKOCYTE_DIFFERENTIATION | 41 | 1.84 | 1.54E-03 | 1.43E-02 |
| GOBP_PROSTANOID_METABOLIC_PROCESS | 36 | 1.84 | 1.33E-03 | 1.28E-02 |
| GOBP_REGULATION_OF_NEUTROPHIL_MIGRATION | 36 | 1.83 | 1.33E-03 | 1.28E-02 |
| GOBP_NEPHRON_DEVELOPMENT | 135 | 1.83 | 1.54E-05 | 3.82E-04 |
| GOBP_REGULATION_OF_DNA_DAMAGE_RESPONSE_SIGNAL_TRANSDUCTION_BY_P53_CLASS_MEDIATOR | 37 | 1.83 | 1.48E-03 | 1.38E-02 |
| GOBP_REGULATION_OF_ENDOTHELIAL_CELL_MIGRATION | 149 | 1.83 | 1.43E-05 | 3.59E-04 |
| GOBP_NEUROINFLAMMATORY_RESPONSE | 55 | 1.83 | 7.83E-04 | 8.53E-03 |
| GOBP_REGULATION_OF_HORMONE_BIOSYNTHETIC_PROCESS | 18 | 1.83 | 4.62E-03 | 3.14E-02 |
| GOBP_POSITIVE_REGULATION_OF_MACROPHAGE_MIGRATION | 21 | 1.83 | 2.51E-03 | 2.06E-02 |
| GOBP_REGULATION_OF_ODONTOBLAST_DIFFERENTIATION | 10 | 1.83 | 3.42E-03 | 2.58E-02 |
| GOBP_POSITIVE_REGULATION_OF_RESPONSE_TO_EXTERNAL_STIMULUS | 499 | 1.83 | 3.47E-11 | 6.36E-09 |
| GOBP_ENDOCARDIAL_CUSHION_DEVELOPMENT | 50 | 1.83 | 7.28E-04 | 8.15E-03 |
| GOBP_PYRIDINE_CONTAINING_COMPOUND_METABOLIC_PROCESS | 155 | 1.83 | 1.03E-05 | 2.70E-04 |
| GOBP_HEPARAN_SULFATE_PROTEOGLYCAN_METABOLIC_PROCESS | 46 | 1.83 | 6.93E-04 | 7.83E-03 |
| GOBP_REGULATION_OF_PLASMA_MEMBRANE_ORGANIZATION | 16 | 1.83 | 3.71E-03 | 2.71E-02 |
| GOBP_RESPONSE_TO_ISCHEMIA | 51 | 1.83 | 7.96E-04 | 8.63E-03 |
| GOBP_MONOATOMIC_ANION_HOMEOSTASIS | 15 | 1.82 | 3.21E-03 | 2.46E-02 |
| GOBP_RESPONSE_TO_SALT | 20 | 1.82 | 6.39E-03 | 3.99E-02 |
| GOBP_NEGATIVE_REGULATION_OF_MYELOID_CELL_DIFFERENTIATION | 75 | 1.82 | 2.98E-04 | 4.13E-03 |
| GOBP_TYROSINE_METABOLIC_PROCESS | 10 | 1.82 | 3.69E-03 | 2.70E-02 |
| GOBP_NEGATIVE_REGULATION_OF_TRANSFORMING_GROWTH_FACTOR_BETA_RECEPTOR_SIGNALING_PATHWAY | 94 | 1.82 | 1.98E-04 | 3.02E-03 |
| GOBP_EXTRACELLULAR_MATRIX_DISASSEMBLY | 52 | 1.82 | 1.19E-03 | 1.17E-02 |
| GOBP_NEUTROPHIL_HOMEOSTASIS | 17 | 1.82 | 4.14E-03 | 2.92E-02 |
| GOBP_POSITIVE_REGULATION_OF_REPRODUCTIVE_PROCESS | 53 | 1.82 | 8.32E-04 | 8.92E-03 |
| GOBP_POSITIVE_REGULATION_OF_CELL_SUBSTRATE_ADHESION | 108 | 1.82 | 1.13E-04 | 1.91E-03 |
| GOBP_REGULATION_OF_SIGNAL_TRANSDUCTION_BY_P53_CLASS_MEDIATOR | 114 | 1.81 | 6.49E-05 | 1.22E-03 |
| GOBP_NEGATIVE_REGULATION_OF_ENDOPLASMIC_RETICULUM_UNFOLDED_PROTEIN_RESPONSE | 21 | 1.81 | 2.83E-03 | 2.25E-02 |
| GOBP_DEFENSE_RESPONSE_TO_VIRUS | 261 | 1.81 | 4.32E-07 | 2.02E-05 |
| GOBP_CELL_KILLING | 128 | 1.81 | 5.64E-05 | 1.11E-03 |
| GOBP_REGULATION_OF_EXTRINSIC_APOPTOTIC_SIGNALING_PATHWAY_VIA_DEATH_DOMAIN_RECEPTORS | 40 | 1.81 | 1.02E-03 | 1.06E-02 |
| GOBP_NEGATIVE_REGULATION_OF_STEROID_METABOLIC_PROCESS | 24 | 1.81 | 6.07E-03 | 3.87E-02 |
| GOBP_CYTOKINE_MEDIATED_SIGNALING_PATHWAY | 372 | 1.81 | 1.84E-08 | 1.49E-06 |
| GOBP_EMBRYONIC_CRANIAL_SKELETON_MORPHOGENESIS | 40 | 1.81 | 1.04E-03 | 1.07E-02 |
| GOBP_VASODILATION | 42 | 1.81 | 1.60E-03 | 1.48E-02 |
| GOBP_RESPONSE_TO_ACTIVITY | 69 | 1.81 | 3.97E-04 | 5.20E-03 |
| GOBP_MATURE_B_CELL_DIFFERENTIATION | 30 | 1.81 | 2.27E-03 | 1.92E-02 |
| GOBP_LEUKOTRIENE_METABOLIC_PROCESS | 16 | 1.81 | 4.43E-03 | 3.07E-02 |
| GOBP_CELLULAR_RESPONSE_TO_MOLECULE_OF_BACTERIAL_ORIGIN | 160 | 1.81 | 2.88E-05 | 6.18E-04 |
| GOBP_CELLULAR_RESPONSE_TO_TOPOLOGICALLY_INCORRECT_PROTEIN | 111 | 1.81 | 1.54E-04 | 2.43E-03 |
| GOBP_RENAL_SYSTEM_VASCULATURE_DEVELOPMENT | 26 | 1.81 | 3.63E-03 | 2.68E-02 |
| GOBP_RESPONSE_TO_MECHANICAL_STIMULUS | 190 | 1.81 | 6.40E-06 | 1.80E-04 |
| GOBP_PROTEIN_TARGETING_TO_PEROXISOME | 10 | 1.81 | 4.35E-03 | 3.04E-02 |
| GOBP_POSITIVE_REGULATION_OF_VIRAL_GENOME_REPLICATION | 27 | 1.81 | 4.43E-03 | 3.07E-02 |
| GOBP_REGULATION_OF_BLOOD_PRESSURE | 133 | 1.81 | 4.87E-05 | 9.98E-04 |
| GOBP_HOMOTYPIC_CELL_CELL_ADHESION | 76 | 1.81 | 4.43E-04 | 5.67E-03 |
| GOBP_ODONTOGENESIS_OF_DENTIN_CONTAINING_TOOTH | 65 | 1.80 | 4.47E-04 | 5.71E-03 |
| GOBP_BRANCHING_INVOLVED_IN_MAMMARY_GLAND_DUCT_MORPHOGENESIS | 23 | 1.80 | 6.19E-03 | 3.90E-02 |
| GOBP_POSITIVE_REGULATION_OF_MACROPHAGE_DERIVED_FOAM_CELL_DIFFERENTIATION | 10 | 1.80 | 4.43E-03 | 3.07E-02 |
| GOBP_FOCAL_ADHESION_ASSEMBLY | 81 | 1.80 | 2.41E-04 | 3.52E-03 |
| GOBP_MATING_BEHAVIOR | 22 | 1.80 | 4.04E-03 | 2.87E-02 |
| GOBP_SUPEROXIDE_ANION_GENERATION | 26 | 1.80 | 3.79E-03 | 2.74E-02 |
| GOBP_PURINE_NUCLEOSIDE_MONOPHOSPHATE_METABOLIC_PROCESS | 40 | 1.80 | 1.14E-03 | 1.14E-02 |
| GOBP_NEGATIVE_REGULATION_OF_MUSCLE_HYPERTROPHY | 28 | 1.80 | 3.77E-03 | 2.73E-02 |
| GOBP_POSITIVE_REGULATION_OF_ENDOTHELIAL_CELL_APOPTOTIC_PROCESS | 18 | 1.80 | 5.99E-03 | 3.85E-02 |
| GOBP_RESPONSE_TO_OXIDATIVE_STRESS | 363 | 1.80 | 4.00E-08 | 2.84E-06 |
| GOBP_NEGATIVE_REGULATION_OF_PRODUCTION_OF_MOLECULAR_MEDIATOR_OF_IMMUNE_RESPONSE | 26 | 1.80 | 3.83E-03 | 2.77E-02 |
| GOBP_ANTIGEN_PROCESSING_AND_PRESENTATION_OF_EXOGENOUS_PEPTIDE_ANTIGEN_VIA_MHC_CLASS_II | 22 | 1.80 | 4.04E-03 | 2.87E-02 |
| GOBP_CARDIAC_CHAMBER_MORPHOGENESIS | 115 | 1.80 | 9.95E-05 | 1.73E-03 |
| GOBP_NEGATIVE_REGULATION_OF_PROTEIN_CATABOLIC_PROCESS | 112 | 1.80 | 7.15E-05 | 1.33E-03 |
| GOBP_REGULATION_OF_NEUTROPHIL_CHEMOTAXIS | 25 | 1.80 | 6.09E-03 | 3.87E-02 |
| GOBP_ENDOPLASMIC_RETICULUM_TO_CYTOSOL_TRANSPORT | 28 | 1.80 | 4.13E-03 | 2.92E-02 |
| GOBP_EMBRYONIC_SKELETAL_SYSTEM_DEVELOPMENT | 121 | 1.80 | 7.82E-05 | 1.42E-03 |
| GOBP_LUTEINIZATION | 10 | 1.79 | 4.67E-03 | 3.17E-02 |
| GOBP_REGULATION_OF_CELLULAR_RESPONSE_TO_TRANSFORMING_GROWTH_FACTOR_BETA_STIMULUS | 142 | 1.79 | 4.01E-05 | 8.42E-04 |
| GOBP_FRUCTOSE_6_PHOSPHATE_METABOLIC_PROCESS | 11 | 1.79 | 6.04E-03 | 3.86E-02 |
| GOBP_SMOOTH_MUSCLE_CELL_MIGRATION | 64 | 1.79 | 4.14E-04 | 5.37E-03 |
| GOBP_REGULATION_OF_CELLULAR_RESPONSE_TO_GROWTH_FACTOR_STIMULUS | 296 | 1.79 | 2.54E-07 | 1.34E-05 |
| GOBP_ZINC_ION_TRANSPORT | 26 | 1.79 | 3.99E-03 | 2.85E-02 |
| GOBP_POSITIVE_REGULATION_OF_VIRAL_PROCESS | 58 | 1.79 | 7.55E-04 | 8.32E-03 |
| GOBP_NEGATIVE_REGULATION_OF_ENDOTHELIAL_CELL_APOPTOTIC_PROCESS | 28 | 1.79 | 4.61E-03 | 3.14E-02 |
| GOBP_DEFENSE_RESPONSE_TO_GRAM_POSITIVE_BACTERIUM | 59 | 1.79 | 6.54E-04 | 7.55E-03 |
| GOBP_LEUKOCYTE_DIFFERENTIATION | 482 | 1.79 | 2.19E-09 | 2.37E-07 |
| GOBP_LYMPHOCYTE_APOPTOTIC_PROCESS | 70 | 1.79 | 5.47E-04 | 6.66E-03 |
| GOBP_REGULATION_OF_MONOCYTE_DIFFERENTIATION | 17 | 1.79 | 5.08E-03 | 3.41E-02 |
| GOBP_MYOBLAST_FUSION | 35 | 1.79 | 4.54E-03 | 3.13E-02 |
| GOBP_OSTEOCLAST_DIFFERENTIATION | 87 | 1.79 | 5.34E-04 | 6.52E-03 |
| GOBP_RESPONSE_TO_FATTY_ACID | 51 | 1.79 | 1.11E-03 | 1.13E-02 |
| GOBP_KETONE_METABOLIC_PROCESS | 174 | 1.79 | 2.43E-05 | 5.40E-04 |
| GOBP_ADAPTIVE_IMMUNE_RESPONSE | 311 | 1.79 | 1.28E-07 | 7.95E-06 |
| GOBP_NEGATIVE_REGULATION_OF_INFLAMMATORY_RESPONSE | 116 | 1.79 | 1.38E-04 | 2.21E-03 |
| GOBP_POSITIVE_REGULATION_OF_TRANSMEMBRANE_RECEPTOR_PROTEIN_SERINE_THREONINE_KINASE_SIGNALING_PATHWAY | 99 | 1.79 | 2.00E-04 | 3.04E-03 |
| GOBP_RESPONSE_TO_UV_A | 11 | 1.79 | 6.36E-03 | 3.98E-02 |
| GOBP_INTRINSIC_APOPTOTIC_SIGNALING_PATHWAY_IN_RESPONSE_TO_ENDOPLASMIC_RETICULUM_STRESS | 61 | 1.78 | 1.63E-03 | 1.49E-02 |
| GOBP_REGULATION_OF_RECEPTOR_LOCALIZATION_TO_SYNAPSE | 28 | 1.78 | 5.41E-03 | 3.57E-02 |
| GOBP_EXTRACELLULAR_MATRIX_CONSTITUENT_SECRETION | 11 | 1.78 | 6.68E-03 | 4.11E-02 |
| GOBP_T_CELL_EXTRAVASATION | 11 | 1.78 | 6.68E-03 | 4.11E-02 |
| GOBP_POSITIVE_REGULATION_OF_STEM_CELL_PROLIFERATION | 37 | 1.78 | 2.75E-03 | 2.20E-02 |
| GOBP_REGULATION_OF_OXIDATIVE_STRESS_INDUCED_INTRINSIC_APOPTOTIC_SIGNALING_PATHWAY | 38 | 1.78 | 3.06E-03 | 2.37E-02 |
| GOBP_GLIAL_CELL_PROLIFERATION | 47 | 1.78 | 1.69E-03 | 1.54E-02 |
| GOBP_PYROPTOTIC_INFLAMMATORY_RESPONSE | 30 | 1.78 | 3.35E-03 | 2.55E-02 |
| GOBP_REGULATION_OF_PHOSPHOLIPID_BIOSYNTHETIC_PROCESS | 13 | 1.78 | 5.91E-03 | 3.82E-02 |
| GOBP_NUCLEOSIDE_SALVAGE | 12 | 1.78 | 6.53E-03 | 4.05E-02 |
| GOBP_REGULATION_OF_DEFENSE_RESPONSE_TO_VIRUS | 54 | 1.78 | 2.40E-03 | 2.00E-02 |
| GOBP_VITAMIN_METABOLIC_PROCESS | 92 | 1.78 | 5.35E-04 | 6.52E-03 |
| GOBP_REGULATION_OF_RESPONSE_TO_WOUNDING | 132 | 1.78 | 3.69E-05 | 7.86E-04 |
| GOBP_REGULATION_OF_WOUND_HEALING | 97 | 1.78 | 3.31E-04 | 4.46E-03 |
| GOBP_POSITIVE_REGULATION_OF_SMAD_PROTEIN_SIGNAL_TRANSDUCTION | 37 | 1.77 | 2.94E-03 | 2.31E-02 |
| GOBP_CELL_CELL_ADHESION_VIA_PLASMA_MEMBRANE_ADHESION_MOLECULES | 215 | 1.77 | 7.03E-06 | 1.95E-04 |
| GOBP_REGULATION_OF_SUPEROXIDE_ANION_GENERATION | 12 | 1.77 | 6.85E-03 | 4.20E-02 |
| GOBP_HORMONE_METABOLIC_PROCESS | 166 | 1.77 | 1.92E-05 | 4.51E-04 |
| GOBP_LEUKOCYTE_PROLIFERATION | 239 | 1.77 | 2.42E-06 | 8.56E-05 |
| GOBP_RELEASE_OF_CYTOCHROME_C_FROM_MITOCHONDRIA | 50 | 1.77 | 1.15E-03 | 1.15E-02 |
| GOBP_GLANDULAR_EPITHELIAL_CELL_DIFFERENTIATION | 31 | 1.77 | 3.02E-03 | 2.34E-02 |
| GOBP_PROTEIN_TRANSMEMBRANE_IMPORT_INTO_INTRACELLULAR_ORGANELLE | 42 | 1.77 | 2.60E-03 | 2.11E-02 |

| | | | | |
|---|---|---|---|---|
| GOBP_POSITIVE_REGULATION_OF_ANIMAL_ORGAN_MORPHOGENESIS | 32 | 1.77 | 7.92E-03 | 4.66E-02 |
| GOBP_DEVELOPMENT_OF_PRIMARY_FEMALE_SEXUAL_CHARACTERISTICS | 91 | 1.77 | 5.88E-04 | 6.98E-03 |
| GOBP_NEGATIVE_REGULATION_OF_SMAD_PROTEIN_SIGNAL_TRANSDUCTION | 23 | 1.77 | 8.58E-03 | 4.87E-02 |
| GOBP_SUPEROXIDE_METABOLIC_PROCESS | 54 | 1.77 | 2.56E-03 | 2.09E-02 |
| GOBP_CELLULAR_RESPONSE_TO_COLD | 12 | 1.77 | 7.25E-03 | 4.38E-02 |
| GOBP_REGULATION_OF_IRE1_MEDIATED_UNFOLDED_PROTEIN_RESPONSE | 15 | 1.77 | 5.73E-03 | 3.73E-02 |
| GOBP_POSITIVE_REGULATION_OF_INFLAMMATORY_RESPONSE | 109 | 1.77 | 1.05E-04 | 1.81E-03 |
| GOBP_HOST_MEDIATED_SUPPRESSION_OF_SYMBIONT_INVASION | 18 | 1.77 | 7.67E-03 | 4.55E-02 |
| GOBP_ENDOPLASMIC_RETICULUM_TO_GOLGI_VESICLE_MEDIATED_TRANSPORT | 131 | 1.76 | 5.89E-05 | 1.14E-03 |
| GOBP_IMPORT_INTO_NUCLEUS | 164 | 1.76 | 7.67E-05 | 1.41E-03 |
| GOBP_RIBONUCLEOSIDE_MONOPHOSPHATE_METABOLIC_PROCESS | 52 | 1.76 | 2.00E-03 | 1.75E-02 |
| GOBP_POSITIVE_REGULATION_OF_PROTEIN_IMPORT_INTO_NUCLEUS | 35 | 1.76 | 5.94E-03 | 3.83E-02 |
| GOBP_REGULATION_OF_ANIMAL_ORGAN_MORPHOGENESIS | 75 | 1.76 | 7.31E-04 | 8.15E-03 |
| GOBP_REGULATION_OF_SYSTEMIC_ARTERIAL_BLOOD_PRESSURE_BY_RENIN_ANGIOTENSIN | 22 | 1.76 | 6.08E-03 | 3.87E-02 |
| GOBP_ERYTHROSE_4_PHOSPHATE_PHOSPHOENOLPYRUVATE_FAMILY_AMINO_ACID_CATABOLIC_PROCESS | 10 | 1.76 | 6.47E-03 | 4.02E-02 |
| GOBP_AUTOPHAGY_OF_PEROXISOME | 16 | 1.76 | 7.46E-03 | 4.47E-02 |
| GOBP_POSITIVE_REGULATION_OF_CYTOPLASMIC_TRANSLATION | 15 | 1.76 | 5.73E-03 | 3.73E-02 |
| GOBP_REGULATION_OF_RESPONSE_TO_ENDOPLASMIC_RETICULUM_STRESS | 88 | 1.76 | 3.31E-04 | 4.46E-03 |
| GOBP_POSITIVE_REGULATION_OF_CELLULAR_RESPONSE_TO_TRANSFORMING_GROWTH_FACTOR_BETA_STIMULUS | 31 | 1.76 | 3.38E-03 | 2.56E-02 |
| GOBP_CELLULAR_RESPONSE_TO_REACTIVE_OXYGEN_SPECIES | 132 | 1.76 | 5.65E-05 | 1.11E-03 |
| GOBP_TYPE_2_IMMUNE_RESPONSE | 26 | 1.76 | 5.31E-03 | 3.53E-02 |
| GOBP_PROTEINOGENIC_AMINO_ACID_BIOSYNTHETIC_PROCESS | 38 | 1.76 | 3.70E-03 | 2.71E-02 |
| GOBP_REGULATION_OF_CELL_SUBSTRATE_JUNCTION_ORGANIZATION | 66 | 1.76 | 7.69E-04 | 8.43E-03 |
| GOBP_RESPONSE_TO_HYDROGEN_PEROXIDE | 91 | 1.76 | 6.80E-04 | 7.78E-03 |
| GOBP_RESPONSE_TO_IONIZING_RADIATION | 136 | 1.76 | 9.06E-05 | 1.61E-03 |
| GOBP_CHAPERONE_MEDIATED_AUTOPHAGY | 15 | 1.75 | 6.13E-03 | 3.89E-02 |
| GOBP_RESPONSE_TO_AMYLOID_BETA | 40 | 1.75 | 1.89E-03 | 1.67E-02 |
| GOBP_NEGATIVE_REGULATION_OF_CELL_GROWTH_INVOLVED_IN_CARDIAC_MUSCLE_CELL_DEVELOPMENT | 11 | 1.75 | 8.60E-03 | 4.87E-02 |
| GOBP_NEURON_APOPTOTIC_PROCESS | 275 | 1.75 | 2.85E-06 | 9.46E-05 |
| GOBP_INFLAMMATORY_RESPONSE_TO_WOUNDING | 22 | 1.75 | 6.39E-03 | 3.99E-02 |
| GOBP_VESICLE_CARGO_LOADING | 32 | 1.75 | 8.47E-03 | 4.83E-02 |
| GOBP_REGULATION_OF_EARLY_ENDOSOME_TO_LATE_ENDOSOME_TRANSPORT | 19 | 1.75 | 8.40E-03 | 4.81E-02 |
| GOBP_NEGATIVE_REGULATION_BY_HOST_OF_VIRAL_GENOME_REPLICATION | 10 | 1.75 | 6.87E-03 | 4.20E-02 |
| GOBP_POSITIVE_REGULATION_OF_TUMOR_NECROSIS_FACTOR_SUPERFAMILY_CYTOKINE_PRODUCTION | 80 | 1.75 | 6.35E-04 | 7.43E-03 |
| GOBP_POSITIVE_REGULATION_OF_ERK1_AND_ERK2_CASCADE | 140 | 1.75 | 1.22E-04 | 2.02E-03 |
| GOBP_CELL_PROLIFERATION_INVOLVED_IN_KIDNEY_DEVELOPMENT | 18 | 1.75 | 8.48E-03 | 4.83E-02 |
| GOBP_NEPHRON_EPITHELIUM_DEVELOPMENT | 101 | 1.75 | 1.83E-04 | 2.82E-03 |
| GOBP_REGULATION_OF_SYNAPTIC_VESICLE_CYCLE | 15 | 1.75 | 6.61E-03 | 4.09E-02 |
| GOBP_TELOMERE_MAINTENANCE_VIA_TELOMERE_LENGTHENING | 70 | 1.75 | 9.41E-04 | 9.87E-03 |
| GOBP_PROGRAMMED_CELL_DEATH_INVOLVED_IN_CELL_DEVELOPMENT | 17 | 1.75 | 6.89E-03 | 4.21E-02 |
| GOBP_REGULATION_OF_SYSTEMIC_ARTERIAL_BLOOD_PRESSURE_MEDIATED_BY_A_CHEMICAL_SIGNAL | 31 | 1.74 | 3.78E-03 | 2.74E-02 |
| GOBP_GTP_METABOLIC_PROCESS | 22 | 1.74 | 6.94E-03 | 4.21E-02 |
| GOBP_NEGATIVE_REGULATION_OF_TRANSMEMBRANE_RECEPTOR_PROTEIN_SERINE_THREONINE_KINASE_SIGNALING_PATHWAY | 142 | 1.74 | 9.21E-05 | 1.62E-03 |
| GOBP_ENDOCARDIAL_CUSHION_MORPHOGENESIS | 40 | 1.74 | 2.13E-03 | 1.82E-02 |
| GOBP_GRANULOCYTE_ACTIVATION | 29 | 1.74 | 7.37E-03 | 4.43E-02 |
| GOBP_MUSCLE_CELL_MIGRATION | 78 | 1.74 | 5.55E-04 | 6.69E-03 |
| GOBP_NEGATIVE_REGULATION_OF_CHONDROCYTE_DIFFERENTIATION | 22 | 1.74 | 6.94E-03 | 4.21E-02 |
| GOBP_UNSATURATED_FATTY_ACID_BIOSYNTHETIC_PROCESS | 46 | 1.74 | 1.85E-03 | 1.65E-02 |
| GOBP_C21_STEROID_HORMONE_METABOLIC_PROCESS | 26 | 1.74 | 6.11E-03 | 3.87E-02 |
| GOBP_NEGATIVE_REGULATION_OF_DNA_DAMAGE_RESPONSE_SIGNAL_TRANSDUCTION_BY_P53_CLASS_MEDIATOR | 15 | 1.74 | 6.93E-03 | 4.21E-02 |
| GOBP_REGULATION_OF_TRANSMEMBRANE_RECEPTOR_PROTEIN_SERINE_THREONINE_KINASE_SIGNALING_PATHWAY | 254 | 1.74 | 5.02E-06 | 1.53E-04 |
| GOBP_TRNA_METABOLIC_PROCESS | 200 | 1.74 | 2.78E-05 | 6.05E-04 |
| GOBP_CELLULAR_EXTRAVASATION | 51 | 1.74 | 2.20E-03 | 1.87E-02 |
| GOBP_VASCULAR_PROCESS_IN_CIRCULATORY_SYSTEM | 210 | 1.74 | 2.35E-05 | 5.32E-04 |
| GOBP_REGULATION_OF_TUBE_SIZE | 109 | 1.74 | 1.70E-04 | 2.63E-03 |
| GOBP_NEGATIVE_REGULATION_OF_CELL_ADHESION | 225 | 1.73 | 1.30E-05 | 3.34E-04 |
| GOBP_REGULATION_OF_TRANSLATIONAL_INITIATION | 79 | 1.73 | 6.54E-04 | 7.55E-03 |
| GOBP_PROTEIN_TARGETING | 319 | 1.73 | 7.64E-07 | 3.23E-05 |
| GOBP_REGULATION_OF_ERYTHROCYTE_DIFFERENTIATION | 46 | 1.73 | 1.99E-03 | 1.74E-02 |
| GOBP_MULTI_MULTICELLULAR_ORGANISM_PROCESS | 165 | 1.73 | 1.52E-04 | 2.41E-03 |
| GOBP_SULFUR_AMINO_ACID_METABOLIC_PROCESS | 29 | 1.73 | 8.19E-03 | 4.72E-02 |
| GOBP_REGULATION_OF_TRANSLATION | 339 | 1.73 | 5.74E-07 | 2.49E-05 |
| GOBP_FEMALE_SEX_DIFFERENTIATION | 102 | 1.73 | 3.20E-04 | 4.37E-03 |
| GOBP_POSITIVE_REGULATION_OF_ENDOCYTOSIS | 117 | 1.73 | 2.81E-04 | 3.97E-03 |
| GOBP_GLIAL_CELL_ACTIVATION | 35 | 1.73 | 8.19E-03 | 4.72E-02 |
| GOBP_REGULATION_OF_HORMONE_METABOLIC_PROCESS | 30 | 1.73 | 5.45E-03 | 3.58E-02 |
| GOBP_PROTEIN_LOCALIZATION_TO_NUCLEOPLASM | 14 | 1.73 | 8.60E-03 | 4.87E-02 |
| GOBP_POSITIVE_REGULATION_OF_NUCLEOCYTOPLASMIC_TRANSPORT | 57 | 1.72 | 1.98E-03 | 1.74E-02 |
| GOBP_NEGATIVE_REGULATION_OF_IMMUNE_SYSTEM_PROCESS | 388 | 1.72 | 4.73E-07 | 2.15E-05 |
| GOBP_NEGATIVE_REGULATION_OF_LEUKOCYTE_CHEMOTAXIS | 15 | 1.72 | 8.14E-03 | 4.72E-02 |
| GOBP_REGULATION_OF_NUCLEOCYTOPLASMIC_TRANSPORT | 112 | 1.72 | 2.84E-04 | 3.99E-03 |
| GOBP_IMPORT_ACROSS_PLASMA_MEMBRANE | 156 | 1.72 | 1.23E-04 | 2.03E-03 |
| GOBP_REGULATION_OF_ENDOPLASMIC_RETICULUM_STRESS_INDUCED_INTRINSIC_APOPTOTIC_SIGNALING_PATHWAY | 33 | 1.72 | 6.09E-03 | 3.87E-02 |
| GOBP_POSITIVE_REGULATION_OF_FIBROBLAST_MIGRATION | 15 | 1.72 | 8.30E-03 | 4.76E-02 |
| GOBP_MACROPHAGE_CYTOKINE_PRODUCTION | 33 | 1.72 | 6.33E-03 | 3.97E-02 |
| GOBP_DEFENSE_RESPONSE_TO_BACTERIUM | 156 | 1.72 | 1.28E-04 | 2.08E-03 |
| GOBP_UNSATURATED_FATTY_ACID_METABOLIC_PROCESS | 78 | 1.72 | 7.69E-04 | 8.43E-03 |
| GOBP_COLUMNAR_CUBOIDAL_EPITHELIAL_CELL_DIFFERENTIATION | 90 | 1.71 | 7.77E-04 | 8.48E-03 |
| GOBP_REGULATION_OF_HEMOPOIESIS | 331 | 1.71 | 8.51E-07 | 3.53E-05 |
| GOBP_ESTABLISHMENT_OF_PROTEIN_LOCALIZATION_TO_ORGANELLE | 434 | 1.71 | 1.44E-07 | 8.59E-06 |
| GOBP_NEGATIVE_REGULATION_OF_STEROID_BIOSYNTHETIC_PROCESS | 22 | 1.71 | 7.96E-03 | 4.67E-02 |
| GOBP_NEGATIVE_REGULATION_OF_UBIQUITIN_DEPENDENT_PROTEIN_CATABOLIC_PROCESS | 54 | 1.71 | 4.35E-03 | 3.04E-02 |
| GOBP_COPII_COATED_VESICLE_BUDDING | 42 | 1.71 | 5.22E-03 | 3.48E-02 |
| GOBP_INTERLEUKIN_1_BETA_PRODUCTION | 63 | 1.71 | 3.53E-03 | 2.62E-02 |
| GOBP_REGULATION_OF_INTEGRIN_MEDIATED_SIGNALING_PATHWAY | 22 | 1.71 | 8.12E-03 | 4.72E-02 |
| GOBP_OLEFINIC_COMPOUND_METABOLIC_PROCESS | 110 | 1.71 | 4.36E-04 | 5.60E-03 |
| GOBP_NUCLEAR_TRANSPORT | 317 | 1.71 | 2.93E-06 | 9.57E-05 |
| GOBP_MODIFIED_AMINO_ACID_METABOLIC_PROCESS | 152 | 1.71 | 1.94E-04 | 2.97E-03 |
| GOBP_SULFUR_COMPOUND_METABOLIC_PROCESS | 280 | 1.71 | 4.42E-06 | 1.37E-04 |
| GOBP_ICOSANOID_BIOSYNTHETIC_PROCESS | 36 | 1.71 | 4.83E-03 | 3.26E-02 |
| GOBP_TERPENOID_METABOLIC_PROCESS | 64 | 1.71 | 1.27E-03 | 1.24E-02 |
| GOBP_POSITIVE_REGULATION_OF_CELL_ADHESION | 388 | 1.71 | 8.60E-07 | 3.54E-05 |
| GOBP_ENDOTHELIAL_CELL_MIGRATION | 204 | 1.70 | 6.37E-05 | 1.22E-03 |
| GOBP_ASPARTATE_FAMILY_AMINO_ACID_METABOLIC_PROCESS | 34 | 1.70 | 4.86E-03 | 3.27E-02 |
| GOBP_CARDIAC_EPITHELIAL_TO_MESENCHYMAL_TRANSITION | 37 | 1.70 | 6.15E-03 | 3.89E-02 |
| GOBP_T_CELL_ACTIVATION | 419 | 1.70 | 2.88E-07 | 1.48E-05 |
| GOBP_ER_NUCLEUS_SIGNALING_PATHWAY | 52 | 1.70 | 3.47E-03 | 2.61E-02 |
| GOBP_REGULATION_OF_LIPID_STORAGE | 39 | 1.70 | 4.94E-03 | 3.32E-02 |
| GOBP_LIPID_EXPORT_FROM_CELL | 41 | 1.70 | 5.99E-03 | 3.85E-02 |
| GOBP_RESPONSE_TO_REACTIVE_OXYGEN_SPECIES | 177 | 1.70 | 7.26E-05 | 1.35E-03 |
| GOBP_PROTEIN_LOCALIZATION_TO_CELL_CELL_JUNCTION | 22 | 1.70 | 8.75E-03 | 4.95E-02 |
| GOBP_CELLULAR_RESPONSE_TO_OSMOTIC_STRESS | 53 | 1.70 | 3.52E-03 | 2.62E-02 |
| GOBP_TUMOR_NECROSIS_FACTOR_SUPERFAMILY_CYTOKINE_PRODUCTION | 127 | 1.70 | 2.60E-04 | 3.76E-03 |
| GOBP_MUSCLE_CELL_DIFFERENTIATION | 348 | 1.70 | 5.28E-06 | 1.57E-04 |
| GOBP_SUBSTRATE_ADHESION_DEPENDENT_CELL_SPREADING | 91 | 1.69 | 1.43E-03 | 1.35E-02 |
| GOBP_POSITIVE_REGULATION_OF_BLOOD_VESSEL_ENDOTHELIAL_CELL_MIGRATION | 52 | 1.69 | 3.90E-03 | 2.80E-02 |

| | | | | | |
|---|---|---|---|---|---|
| | GOBP_ORGANIC_ACID_BIOSYNTHETIC_PROCESS | 245 | 1.69 | 1.83E-05 | 4.42E-04 |
| | GOBP_REGULATION_OF_STRESS_ACTIVATED_PROTEIN_KINASE_SIGNALING_CASCADE | 37 | 1.69 | 6.62E-03 | 4.09E-02 |
| | GOBP_CIRCULATORY_SYSTEM_PROCESS | 483 | 1.69 | 7.11E-08 | 4.70E-06 |
| | GOBP_REGULATION_OF_IMMUNE_EFFECTOR_PROCESS | 270 | 1.69 | 2.05E-05 | 4.76E-04 |
| | GOBP_POSITIVE_REGULATION_OF_EXTRINSIC_APOPTOTIC_SIGNALING_PATHWAY | 48 | 1.69 | 4.17E-03 | 2.94E-02 |
| | GOBP_AMINO_ACID_IMPORT_ACROSS_PLASMA_MEMBRANE | 39 | 1.69 | 5.66E-03 | 3.70E-02 |
| | GOBP_PROTEIN_O_LINKED_GLYCOSYLATION | 91 | 1.69 | 1.49E-03 | 1.39E-02 |
| | GOBP_HORMONE_BIOSYNTHETIC_PROCESS | 44 | 1.69 | 4.14E-03 | 2.92E-02 |
| | GOBP_RESPONSE_TO_INTERLEUKIN_4 | 31 | 1.69 | 5.58E-03 | 3.65E-02 |
| | GOBP_VASCULAR_ASSOCIATED_SMOOTH_MUSCLE_CELL_PROLIFERATION | 55 | 1.69 | 3.91E-03 | 2.80E-02 |
| | GOBP_DEVELOPMENT_OF_PRIMARY_SEXUAL_CHARACTERISTICS | 193 | 1.69 | 9.90E-05 | 1.72E-03 |
| | GOBP_LYMPHOCYTE_MEDIATED_IMMUNITY | 199 | 1.69 | 1.60E-04 | 2.51E-03 |
| | GOBP_POSITIVE_REGULATION_OF_INTRINSIC_APOPTOTIC_SIGNALING_PATHWAY | 60 | 1.69 | 2.02E-03 | 1.76E-02 |
| | GOBP_PLATELET_AGGREGATION | 56 | 1.69 | 4.23E-03 | 2.98E-02 |
| | GOBP_MUSCLE_ORGAN_MORPHOGENESIS | 66 | 1.69 | 1.77E-03 | 1.58E-02 |
| | GOBP_AMIDE_METABOLIC_PROCESS | 393 | 1.69 | 1.05E-06 | 4.25E-05 |
| | GOBP_POSITIVE_REGULATION_OF_G_PROTEIN_COUPLED_RECEPTOR_SIGNALING_PATHWAY | 30 | 1.68 | 8.19E-03 | 4.72E-02 |
| | GOBP_POSITIVE_REGULATION_OF_DEFENSE_RESPONSE | 399 | 1.68 | 4.57E-07 | 2.10E-05 |
| | GOBP_REGULATION_OF_KERATINOCYTE_PROLIFERATION | 36 | 1.68 | 6.24E-03 | 3.93E-02 |
| | GOBP_INTEGRIN_MEDIATED_SIGNALING_PATHWAY | 96 | 1.68 | 7.54E-04 | 8.32E-03 |
| | GOBP_MONONUCLEAR_CELL_DIFFERENTIATION | 400 | 1.68 | 1.62E-06 | 6.12E-05 |
| | GOBP_COLLAGEN_METABOLIC_PROCESS | 90 | 1.68 | 1.24E-03 | 1.21E-02 |
| | GOBP_POSITIVE_REGULATION_OF_PRODUCTION_OF_MOLECULAR_MEDIATOR_OF_IMMUNE_RESPONSE | 106 | 1.68 | 6.51E-04 | 7.55E-03 |
| | GOBP_RIBOSOME_ASSEMBLY | 60 | 1.68 | 2.17E-03 | 1.84E-02 |
| | GOBP_REGULATION_OF_REPRODUCTIVE_PROCESS | 135 | 1.68 | 2.18E-04 | 3.21E-03 |
| | GOBP_MEMBRANE_LIPID_CATABOLIC_PROCESS | 39 | 1.68 | 6.47E-03 | 4.02E-02 |
| | GOBP_RETROGRADE_VESICLE_MEDIATED_TRANSPORT_GOLGI_TO_ENDOPLASMIC_RETICULUM | 49 | 1.68 | 7.43E-03 | 4.46E-02 |
| | GOBP_CARDIAC_CHAMBER_DEVELOPMENT | 154 | 1.68 | 2.05E-04 | 3.07E-03 |
| | GOBP_T_CELL_MIGRATION | 53 | 1.68 | 4.07E-03 | 2.88E-02 |
| | GOBP_GLYCOSPHINGOLIPID_METABOLIC_PROCESS | 56 | 1.68 | 4.47E-03 | 3.09E-02 |
| | GOBP_DNA_DAMAGE_RESPONSE_SIGNAL_TRANSDUCTION_BY_P53_CLASS_MEDIATOR | 73 | 1.68 | 1.75E-03 | 1.57E-02 |
| | GOBP_FIBROBLAST_MIGRATION | 55 | 1.68 | 4.78E-03 | 3.24E-02 |
| | GOBP_O_GLYCAN_PROCESSING | 34 | 1.68 | 6.03E-03 | 3.86E-02 |
| | GOBP_POSITIVE_REGULATION_OF_CYTOKINE_PRODUCTION | 354 | 1.67 | 3.18E-06 | 1.03E-04 |
| | GOBP_NEGATIVE_REGULATION_OF_EPITHELIAL_CELL_PROLIFERATION | 115 | 1.67 | 6.20E-04 | 7.28E-03 |
| | GOBP_NEGATIVE_REGULATION_OF_OSTEOBLAST_DIFFERENTIATION | 44 | 1.67 | 4.60E-03 | 3.14E-02 |
| | GOBP_ENDOTHELIUM_DEVELOPMENT | 123 | 1.67 | 2.61E-04 | 3.76E-03 |
| | GOBP_PROTEIN_RNA_COMPLEX_ORGANIZATION | 209 | 1.67 | 6.36E-05 | 1.22E-03 |
| | GOBP_RESPONSE_TO_TOXIC_SUBSTANCE | 209 | 1.67 | 6.42E-05 | 1.22E-03 |
| | GOBP_REGULATION_OF_MORPHOGENESIS_OF_A_BRANCHING_STRUCTURE | 51 | 1.67 | 4.42E-03 | 3.07E-02 |
| | GOBP_EMBRYO_IMPLANTATION | 60 | 1.67 | 2.36E-03 | 1.97E-02 |
| | GOBP_PROTEIN_METHYLATION | 40 | 1.67 | 4.75E-03 | 3.22E-02 |
| | GOBP_IMMUNOGLOBULIN_PRODUCTION | 85 | 1.67 | 1.28E-03 | 1.24E-02 |
| | GOBP_DEFENSE_RESPONSE_TO_GRAM_NEGATIVE_BACTERIUM | 45 | 1.67 | 5.25E-03 | 3.50E-02 |
| | GOBP_REGULATION_OF_RESPONSE_TO_BIOTIC_STIMULUS | 438 | 1.67 | 3.90E-07 | 1.84E-05 |
| | GOBP_CELLULAR_RESPONSE_TO_LIPID | 500 | 1.67 | 8.63E-08 | 5.49E-06 |
| | GOBP_NEGATIVE_REGULATION_OF_LOCOMOTION | 279 | 1.67 | 2.37E-05 | 5.33E-04 |
| | GOBP_WATER_SOLUBLE_VITAMIN_METABOLIC_PROCESS | 60 | 1.67 | 2.40E-03 | 2.00E-02 |
| | GOBP_MORPHOGENESIS_OF_AN_EPITHELIUM | 442 | 1.66 | 7.64E-07 | 3.23E-05 |
| | GOBP_MODULATION_OF_PROCESS_OF_ANOTHER_ORGANISM | 91 | 1.66 | 2.13E-03 | 1.82E-02 |
| | GOBP_REGULATION_OF_CELL_MIGRATION_INVOLVED_IN_SPROUTING_ANGIOGENESIS | 36 | 1.66 | 7.57E-03 | 4.50E-02 |
| | GOBP_HOMEOSTASIS_OF_NUMBER_OF_CELLS | 294 | 1.66 | 1.62E-05 | 3.97E-04 |
| | GOBP_KIDNEY_EPITHELIUM_DEVELOPMENT | 129 | 1.66 | 5.16E-04 | 6.42E-03 |
| | GOBP_NEGATIVE_REGULATION_OF_CATABOLIC_PROCESS | 329 | 1.66 | 1.02E-05 | 2.70E-04 |
| | GOBP_LYMPHOCYTE_MIGRATION | 77 | 1.66 | 2.49E-03 | 2.05E-02 |
| | GOBP_DETECTION_OF_MECHANICAL_STIMULUS | 46 | 1.66 | 5.09E-03 | 3.41E-02 |
| | GOBP_RESPONSE_TO_TYPE_II_INTERFERON | 102 | 1.66 | 8.39E-04 | 8.97E-03 |
| | GOBP_PLACENTA_DEVELOPMENT | 132 | 1.66 | 2.95E-04 | 4.12E-03 |
| | GOBP_PROTEIN_TARGETING_TO_MEMBRANE | 113 | 1.66 | 1.39E-03 | 1.32E-02 |
| | GOBP_PRODUCTION_OF_MOLECULAR_MEDIATOR_INVOLVED_IN_INFLAMMATORY_RESPONSE | 72 | 1.66 | 2.46E-03 | 2.03E-02 |
| | GOBP_IN_UTERO_EMBRYONIC_DEVELOPMENT | 378 | 1.66 | 2.88E-06 | 9.49E-05 |
| | GOBP_METANEPHROS_DEVELOPMENT | 73 | 1.66 | 2.42E-03 | 2.01E-02 |
| | GOBP_NEGATIVE_REGULATION_OF_INTRINSIC_APOPTOTIC_SIGNALING_PATHWAY_IN_RESPONSE_TO_DNA_DAMAGE | 31 | 1.66 | 8.15E-03 | 4.72E-02 |
| | GOBP_CELL_DIFFERENTIATION_INVOLVED_IN_KIDNEY_DEVELOPMENT | 48 | 1.66 | 5.81E-03 | 3.76E-02 |
| | GOBP_POSITIVE_REGULATION_OF_MAPK_CASCADE | 360 | 1.66 | 2.68E-06 | 9.19E-05 |
| | GOBP_NEGATIVE_REGULATION_OF_RESPONSE_TO_WOUNDING | 69 | 1.65 | 3.31E-03 | 2.53E-02 |
| | GOBP_ERK1_AND_ERK2_CASCADE | 244 | 1.65 | 6.26E-05 | 1.21E-03 |
| | GOBP_RESPIRATORY_SYSTEM_DEVELOPMENT | 196 | 1.65 | 1.57E-04 | 2.47E-03 |
| | GOBP_HEART_MORPHOGENESIS | 233 | 1.65 | 5.41E-05 | 1.08E-03 |
| | GOBP_MONOCARBOXYLIC_ACID_METABOLIC_PROCESS | 488 | 1.65 | 5.63E-07 | 2.49E-05 |
| | GOBP_RESPONSE_TO_TUMOR_NECROSIS_FACTOR | 195 | 1.65 | 1.23E-04 | 2.03E-03 |
| | GOBP_OSTEOBLAST_PROLIFERATION | 31 | 1.65 | 8.32E-03 | 4.76E-02 |
| | GOBP_ACTIVATION_OF_IMMUNE_RESPONSE | 433 | 1.65 | 1.32E-06 | 5.10E-05 |
| | GOBP_EMBRYONIC_SKELETAL_SYSTEM_MORPHOGENESIS | 85 | 1.65 | 1.71E-03 | 1.55E-02 |
| | GOBP_CELLULAR_RESPONSE_TO_ABIOTIC_STIMULUS | 304 | 1.65 | 1.23E-05 | 3.19E-04 |
| | GOBP_LEUKOCYTE_MEDIATED_IMMUNITY | 266 | 1.65 | 3.86E-05 | 8.19E-04 |
| | GOBP_REGULATION_OF_OSSIFICATION | 108 | 1.65 | 1.05E-03 | 1.08E-02 |
| | GOBP_REGULATION_OF_PROTEOLYSIS | 350 | 1.65 | 5.99E-06 | 1.70E-04 |
| | GOBP_CARDIAC_MUSCLE_TISSUE_DEVELOPMENT | 201 | 1.65 | 9.16E-05 | 1.62E-03 |
| | GOBP_SMALL_MOLECULE_BIOSYNTHETIC_PROCESS | 485 | 1.65 | 1.50E-07 | 8.81E-06 |
| | GOBP_EPITHELIAL_TO_MESENCHYMAL_TRANSITION | 158 | 1.65 | 5.73E-04 | 6.85E-03 |
| | GOBP_T_CELL_PROLIFERATION | 152 | 1.65 | 5.49E-04 | 6.66E-03 |
| | GOBP_FATTY_ACID_METABOLIC_PROCESS | 309 | 1.64 | 1.34E-05 | 3.41E-04 |
| | GOBP_AMEBOIDAL_TYPE_CELL_MIGRATION | 223 | 1.64 | 1.19E-04 | 1.99E-03 |
| | GOBP_REPRODUCTIVE_SYSTEM_DEVELOPMENT | 252 | 1.64 | 8.13E-05 | 1.46E-03 |
| | GOBP_REGULATION_OF_SYSTEMIC_ARTERIAL_BLOOD_PRESSURE | 71 | 1.64 | 2.57E-03 | 2.10E-02 |
| | GOBP_LEUKOCYTE_HOMEOSTASIS | 95 | 1.64 | 1.93E-03 | 1.70E-02 |
| | GOBP_LYMPHOCYTE_DIFFERENTIATION | 323 | 1.64 | 1.47E-05 | 3.67E-04 |
| | GOBP_ANTIGEN_PROCESSING_AND_PRESENTATION_OF_EXOGENOUS_ANTIGEN | 34 | 1.64 | 7.74E-03 | 4.57E-02 |
| | GOBP_ENDOTHELIAL_CELL_DIFFERENTIATION | 106 | 1.64 | 1.25E-03 | 1.22E-02 |
| | GOBP_T_CELL_APOPTOTIC_PROCESS | 50 | 1.64 | 5.13E-03 | 3.43E-02 |
| | GOBP_NEGATIVE_REGULATION_OF_CELL_SUBSTRATE_ADHESION | 52 | 1.64 | 6.35E-03 | 3.98E-02 |
| | GOBP_CELLULAR_RESPONSE_TO_OXYGEN_LEVELS | 157 | 1.64 | 4.84E-04 | 6.10E-03 |
| | GOBP_ALPHA_AMINO_ACID_METABOLIC_PROCESS | 160 | 1.63 | 7.39E-04 | 8.20E-03 |
| | GOBP_PROSTATE_GLAND_DEVELOPMENT | 46 | 1.63 | 6.79E-03 | 4.17E-02 |
| | GOBP_ENDOCRINE_PROCESS | 72 | 1.63 | 3.67E-03 | 2.70E-02 |
| | GOBP_REGULATION_OF_KETONE_METABOLIC_PROCESS | 105 | 1.63 | 2.08E-03 | 1.80E-02 |
| | GOBP_GLOMERULUS_DEVELOPMENT | 63 | 1.62 | 8.10E-03 | 4.72E-02 |
| | GOBP_REGULATION_OF_CELL_CELL_ADHESION | 371 | 1.62 | 9.87E-06 | 2.65E-04 |
| | GOBP_VASCULAR_ENDOTHELIAL_GROWTH_FACTOR_RECEPTOR_SIGNALING_PATHWAY | 53 | 1.62 | 7.62E-03 | 4.52E-02 |
| | GOBP_INTERLEUKIN_1_PRODUCTION | 74 | 1.62 | 3.58E-03 | 2.66E-02 |
| | GOBP_PROTEIN_TRANSMEMBRANE_TRANSPORT | 74 | 1.62 | 3.71E-03 | 2.71E-02 |
| | GOBP_DICARBOXYLIC_ACID_METABOLIC_PROCESS | 89 | 1.62 | 2.42E-03 | 2.01E-02 |
| | GOBP_POSITIVE_REGULATION_OF_HEMOPOIESIS | 155 | 1.62 | 5.03E-04 | 6.31E-03 |
| | GOBP_REGULATION_OF_LEUKOCYTE_PROLIFERATION | 186 | 1.62 | 3.25E-04 | 4.43E-03 |
| | GOBP_REGULATION_OF_LEUKOCYTE_DIFFERENTIATION | 249 | 1.62 | 5.18E-05 | 1.04E-03 |

| | | | | |
|---|---|---|---|---|
| GOBP_SKELETAL_SYSTEM_MORPHOGENESIS | 206 | 1.62 | 4.22E-04 | 5.45E-03 |
| GOBP_REGULATION_OF_CHONDROCYTE_DIFFERENTIATION | 50 | 1.61 | 6.22E-03 | 3.92E-02 |
| GOBP_FIBROBLAST_PROLIFERATION | 98 | 1.61 | 1.37E-03 | 1.31E-02 |
| GOBP_SEMI_LUNAR_VALVE_DEVELOPMENT | 44 | 1.61 | 7.54E-03 | 4.50E-02 |
| GOBP_CYTOKINE_PRODUCTION_INVOLVED_IN_IMMUNE_RESPONSE | 93 | 1.61 | 2.30E-03 | 1.94E-02 |
| GOBP_BLOOD_VESSEL_ENDOTHELIAL_CELL_MIGRATION | 109 | 1.61 | 1.16E-03 | 1.15E-02 |
| GOBP_CELLULAR_RESPONSE_TO_VIRUS | 51 | 1.61 | 7.56E-03 | 4.50E-02 |
| GOBP_REGULATION_OF_FIBROBLAST_PROLIFERATION | 80 | 1.61 | 3.02E-03 | 2.34E-02 |
| GOBP_NEGATIVE_REGULATION_OF_PROTEASOMAL_UBIQUITIN_DEPENDENT_PROTEIN_CATABOLIC_PROCESS | 40 | 1.61 | 7.49E-03 | 4.49E-02 |
| GOBP_POSITIVE_REGULATION_OF_NEURON_APOPTOTIC_PROCESS | 55 | 1.61 | 8.49E-03 | 4.84E-02 |
| GOBP_ERYTHROCYTE_HOMEOSTASIS | 135 | 1.61 | 6.98E-04 | 7.86E-03 |
| GOBP_REGULATION_OF_NEURON_APOPTOTIC_PROCESS | 227 | 1.61 | 1.68E-04 | 2.61E-03 |
| GOBP_REGULATION_OF_IMMUNOGLOBULIN_PRODUCTION | 57 | 1.61 | 8.09E-03 | 4.71E-02 |
| GOBP_CELLULAR_RESPONSE_TO_UV | 88 | 1.61 | 2.93E-03 | 2.31E-02 |
| GOBP_REGULATION_OF_CELL_SHAPE | 121 | 1.61 | 1.17E-03 | 1.16E-02 |
| GOBP_RESPONSE_TO_NICOTINE | 40 | 1.60 | 7.96E-03 | 4.67E-02 |
| GOBP_HOMOPHILIC_CELL_ADHESION_VIA_PLASMA_MEMBRANE_ADHESION_MOLECULES | 143 | 1.60 | 9.49E-04 | 9.91E-03 |
| GOBP_CELLULAR_RESPONSE_TO_RADIATION | 162 | 1.60 | 5.19E-04 | 6.43E-03 |
| GOBP_MYELOID_CELL_DEVELOPMENT | 67 | 1.60 | 3.73E-03 | 2.71E-02 |
| GOBP_POSITIVE_REGULATION_OF_PHOSPHORUS_METABOLIC_PROCESS | 281 | 1.60 | 6.60E-05 | 1.23E-03 |
| GOBP_SEX_DIFFERENTIATION | 224 | 1.60 | 3.07E-04 | 4.25E-03 |
| GOBP_RESPONSE_TO_LEUKEMIA_INHIBITORY_FACTOR | 96 | 1.59 | 2.06E-03 | 1.79E-02 |
| GOBP_NEGATIVE_REGULATION_OF_LEUKOCYTE_PROLIFERATION | 65 | 1.59 | 4.85E-03 | 3.27E-02 |
| GOBP_EPITHELIAL_TUBE_MORPHOGENESIS | 299 | 1.59 | 1.31E-04 | 2.11E-03 |
| GOBP_RHYTHMIC_PROCESS | 252 | 1.59 | 2.13E-04 | 3.15E-03 |
| GOBP_GLIAL_CELL_MIGRATION | 50 | 1.59 | 7.87E-03 | 4.64E-02 |
| GOBP_EMBRYONIC_ORGAN_DEVELOPMENT | 399 | 1.59 | 8.27E-06 | 2.25E-04 |
| GOBP_REGULATION_OF_LIPID_BIOSYNTHETIC_PROCESS | 150 | 1.59 | 7.30E-04 | 8.15E-03 |
| GOBP_POSITIVE_REGULATION_OF_ENDOTHELIAL_CELL_PROLIFERATION | 83 | 1.59 | 3.61E-03 | 2.67E-02 |
| GOBP_POSITIVE_REGULATION_OF_EPITHELIAL_CELL_PROLIFERATION | 167 | 1.59 | 1.03E-03 | 1.07E-02 |
| GOBP_NEGATIVE_REGULATION_OF_PROTEIN_LOCALIZATION | 198 | 1.58 | 3.76E-04 | 4.98E-03 |
| GOBP_POSITIVE_REGULATION_OF_INTERLEUKIN_6_PRODUCTION | 76 | 1.58 | 5.58E-03 | 3.65E-02 |
| GOBP_POSITIVE_REGULATION_OF_AUTOPHAGY | 153 | 1.58 | 1.59E-03 | 1.47E-02 |
| GOBP_GLIOGENESIS | 291 | 1.58 | 1.28E-04 | 2.08E-03 |
| GOBP_REGULATION_OF_INTRACELLULAR_TRANSPORT | 247 | 1.58 | 3.27E-04 | 4.45E-03 |
| GOBP_RESPONSE_TO_NUTRIENT_LEVELS | 429 | 1.58 | 5.35E-06 | 1.57E-04 |
| GOBP_ANTIGEN_PROCESSING_AND_PRESENTATION | 86 | 1.57 | 4.61E-03 | 3.14E-02 |
| GOBP_COLLAGEN_FIBRIL_ORGANIZATION | 66 | 1.57 | 6.41E-03 | 3.99E-02 |
| GOBP_REGULATION_OF_INTRACELLULAR_PROTEIN_TRANSPORT | 140 | 1.57 | 1.98E-03 | 1.74E-02 |
| GOBP_NEGATIVE_REGULATION_OF_IMMUNE_RESPONSE | 144 | 1.57 | 1.22E-03 | 1.20E-02 |
| GOBP_POSITIVE_REGULATION_OF_LIPID_LOCALIZATION | 82 | 1.57 | 6.01E-03 | 3.85E-02 |
| GOBP_CELLULAR_RESPONSE_TO_TYPE_II_INTERFERON | 81 | 1.57 | 3.85E-03 | 2.77E-02 |
| GOBP_CELLULAR_RESPONSE_TO_VASCULAR_ENDOTHELIAL_GROWTH_FACTOR_STIMULUS | 68 | 1.57 | 8.17E-03 | 4.72E-02 |
| GOBP_PROTEIN_STABILIZATION | 216 | 1.57 | 5.57E-04 | 6.69E-03 |
| GOBP_RESPONSE_TO_XENOBIOTIC_STIMULUS | 324 | 1.57 | 9.55E-05 | 1.67E-03 |
| GOBP_T_CELL_ACTIVATION_INVOLVED_IN_IMMUNE_RESPONSE | 87 | 1.57 | 6.60E-03 | 4.09E-02 |
| GOBP_RESPONSE_TO_METAL_ION | 280 | 1.56 | 1.31E-04 | 2.11E-03 |
| GOBP_CHONDROCYTE_DIFFERENTIATION | 110 | 1.56 | 2.86E-03 | 2.27E-02 |
| GOBP_ISOPRENOID_METABOLIC_PROCESS | 83 | 1.56 | 4.41E-03 | 3.07E-02 |
| GOBP_CELLULAR_RESPONSE_TO_XENOBIOTIC_STIMULUS | 134 | 1.56 | 1.45E-03 | 1.36E-02 |
| GOBP_AMIDE_BIOSYNTHETIC_PROCESS | 151 | 1.56 | 1.29E-03 | 1.25E-02 |
| GOBP_REGULATION_OF_LEUKOCYTE_APOPTOTIC_PROCESS | 73 | 1.56 | 6.09E-03 | 3.87E-02 |
| GOBP_TISSUE_REMODELING | 149 | 1.56 | 2.11E-03 | 1.81E-02 |
| GOBP_REGULATION_OF_STEM_CELL_PROLIFERATION | 66 | 1.56 | 7.73E-03 | 4.57E-02 |
| GOBP_POSITIVE_REGULATION_OF_IMMUNE_EFFECTOR_PROCESS | 177 | 1.56 | 9.46E-04 | 9.90E-03 |
| GOBP_PLASMA_MEMBRANE_ORGANIZATION | 140 | 1.55 | 2.62E-03 | 2.13E-02 |
| GOBP_NEGATIVE_REGULATION_OF_HEMOPOIESIS | 87 | 1.55 | 7.74E-03 | 4.57E-02 |
| GOBP_MYOTUBE_DIFFERENTIATION | 104 | 1.55 | 3.32E-03 | 2.53E-02 |
| GOBP_CELL_ACTIVATION_INVOLVED_IN_IMMUNE_RESPONSE | 214 | 1.55 | 8.66E-04 | 9.20E-03 |
| GOBP_XENOBIOTIC_METABOLIC_PROCESS | 79 | 1.55 | 5.48E-03 | 3.59E-02 |
| GOBP_PROTEIN_LOCALIZATION_TO_MITOCHONDRION | 133 | 1.55 | 3.25E-03 | 2.49E-02 |
| GOBP_NEGATIVE_REGULATION_OF_CELL_CELL_ADHESION | 145 | 1.55 | 2.10E-03 | 1.81E-02 |
| GOBP_REGULATION_OF_POST_TRANSLATIONAL_PROTEIN_MODIFICATION | 236 | 1.55 | 5.77E-04 | 6.89E-03 |
| GOBP_STEM_CELL_PROLIFERATION | 98 | 1.55 | 3.15E-03 | 2.42E-02 |
| GOBP_PHAGOCYTOSIS | 170 | 1.55 | 9.74E-04 | 1.01E-02 |
| GOBP_VESICLE_BUDDING_FROM_MEMBRANE | 84 | 1.55 | 8.23E-03 | 4.73E-02 |
| GOBP_POSITIVE_REGULATION_OF_MACROAUTOPHAGY | 90 | 1.55 | 4.60E-03 | 3.14E-02 |
| GOBP_RESPONSE_TO_STARVATION | 203 | 1.54 | 1.10E-03 | 1.12E-02 |
| GOBP_MUSCLE_TISSUE_DEVELOPMENT | 364 | 1.54 | 7.87E-05 | 1.42E-03 |
| GOBP_CELL_CELL_JUNCTION_ORGANIZATION | 175 | 1.54 | 1.74E-03 | 1.57E-02 |
| GOBP_TYPE_I_INTERFERON_PRODUCTION | 113 | 1.54 | 5.52E-03 | 3.62E-02 |
| GOBP_REGULATION_OF_GLIOGENESIS | 82 | 1.54 | 7.91E-03 | 4.66E-02 |
| GOBP_NEGATIVE_REGULATION_OF_CELL_CYCLE_G1_S_PHASE_TRANSITION | 60 | 1.54 | 8.30E-03 | 4.76E-02 |
| GOBP_DIGESTIVE_SYSTEM_DEVELOPMENT | 119 | 1.54 | 3.12E-03 | 2.41E-02 |
| GOBP_MONOATOMIC_ION_HOMEOSTASIS | 486 | 1.54 | 1.97E-05 | 4.59E-04 |
| GOBP_CELLULAR_RESPONSE_TO_NUTRIENT_LEVELS | 233 | 1.54 | 6.83E-04 | 7.79E-03 |
| GOBP_NEGATIVE_REGULATION_OF_RESPONSE_TO_EXTERNAL_STIMULUS | 292 | 1.54 | 2.84E-04 | 3.99E-03 |
| GOBP_STEROID_METABOLIC_PROCESS | 243 | 1.54 | 6.10E-04 | 7.22E-03 |
| GOBP_HEART_VALVE_DEVELOPMENT | 71 | 1.54 | 7.37E-03 | 4.43E-02 |
| GOBP_MUSCLE_ADAPTATION | 90 | 1.54 | 5.25E-03 | 3.50E-02 |
| GOBP_T_CELL_DIFFERENTIATION | 238 | 1.53 | 5.56E-04 | 6.69E-03 |
| GOBP_POSITIVE_REGULATION_OF_PHOSPHORYLATION | 237 | 1.53 | 9.29E-04 | 9.78E-03 |
| GOBP_REGULATION_OF_LIPID_LOCALIZATION | 132 | 1.53 | 2.85E-03 | 2.26E-02 |
| GOBP_PROTEIN_TARGETING_TO_MITOCHONDRION | 107 | 1.53 | 3.55E-03 | 2.64E-02 |
| GOBP_NEGATIVE_REGULATION_OF_DEFENSE_RESPONSE | 193 | 1.53 | 1.64E-03 | 1.49E-02 |
| GOBP_POSITIVE_REGULATION_OF_RESPONSE_TO_BIOTIC_STIMULUS | 326 | 1.53 | 2.97E-04 | 4.13E-03 |
| GOBP_REGULATION_OF_BLOOD_CIRCULATION | 206 | 1.53 | 1.42E-03 | 1.35E-02 |
| GOBP_LYMPHOCYTE_ACTIVATION_INVOLVED_IN_IMMUNE_RESPONSE | 157 | 1.53 | 2.98E-03 | 2.33E-02 |
| GOBP_MAMMARY_GLAND_EPITHELIUM_DEVELOPMENT | 67 | 1.53 | 8.58E-03 | 4.87E-02 |
| GOBP_RNA_LOCALIZATION | 192 | 1.53 | 1.63E-03 | 1.49E-02 |
| GOBP_INORGANIC_ION_HOMEOSTASIS | 418 | 1.53 | 4.58E-05 | 9.47E-04 |
| GOBP_LEUKOCYTE_CELL_CELL_ADHESION | 304 | 1.52 | 2.63E-04 | 3.78E-03 |
| GOBP_HEMOSTASIS | 181 | 1.52 | 1.61E-03 | 1.48E-02 |
| GOBP_NERVE_DEVELOPMENT | 83 | 1.52 | 6.17E-03 | 3.90E-02 |
| GOBP_SKELETAL_SYSTEM_DEVELOPMENT | 468 | 1.52 | 3.89E-05 | 8.22E-04 |
| GOBP_ORGANIC_ACID_TRANSMEMBRANE_TRANSPORT | 120 | 1.52 | 3.39E-03 | 2.56E-02 |
| GOBP_MESENCHYME_DEVELOPMENT | 281 | 1.52 | 3.52E-04 | 4.72E-03 |
| GOBP_INTRACELLULAR_MONOATOMIC_ION_HOMEOSTASIS | 422 | 1.52 | 6.33E-05 | 1.22E-03 |
| GOBP_RESPONSE_TO_STEROID_HORMONE | 279 | 1.52 | 7.21E-04 | 8.10E-03 |
| GOBP_STRIATED_MUSCLE_CELL_DIFFERENTIATION | 248 | 1.51 | 8.28E-04 | 8.90E-03 |
| GOBP_REGULATION_OF_ACTIN_FILAMENT_BUNDLE_ASSEMBLY | 94 | 1.51 | 8.19E-03 | 4.72E-02 |
| GOBP_MONOSACCHARIDE_METABOLIC_PROCESS | 231 | 1.51 | 9.98E-04 | 1.04E-02 |
| GOBP_BONE_MINERALIZATION | 108 | 1.51 | 6.89E-03 | 4.21E-02 |
| GOBP_NEGATIVE_REGULATION_OF_IMMUNE_EFFECTOR_PROCESS | 83 | 1.51 | 7.01E-03 | 4.24E-02 |
| GOBP_POSITIVE_REGULATION_OF_MRNA_METABOLIC_PROCESS | 128 | 1.51 | 5.36E-03 | 3.55E-02 |
| GOBP_CELLULAR_RESPONSE_TO_LIGHT_STIMULUS | 103 | 1.51 | 5.43E-03 | 3.58E-02 |

| | | | | | |
|---|---|---|---|---|---|
| | GOBP_B_CELL_ACTIVATION | 211 | 1.50 | 2.11E-03 | 1.81E-02 |
| | GOBP_CELLULAR_RESPONSE_TO_STARVATION | 174 | 1.50 | 3.35E-03 | 2.55E-02 |
| | GOBP_POSITIVE_REGULATION_OF_INTRACELLULAR_TRANSPORT | 126 | 1.50 | 3.97E-03 | 2.84E-02 |
| | GOBP_REGULATION_OF_PROTEASOMAL_PROTEIN_CATABOLIC_PROCESS | 193 | 1.50 | 2.36E-03 | 1.97E-02 |
| | GOBP_SKIN_DEVELOPMENT | 208 | 1.50 | 2.05E-03 | 1.78E-02 |
| | GOBP_SPROUTING_ANGIOGENESIS | 124 | 1.50 | 6.99E-03 | 4.23E-02 |
| | GOBP_REGULATION_OF_PROTEIN_STABILITY | 342 | 1.49 | 3.08E-04 | 4.25E-03 |
| | GOBP_REGULATION_OF_ACTIN_FILAMENT_BASED_PROCESS | 341 | 1.49 | 2.08E-04 | 3.11E-03 |
| | GOBP_BIOMINERAL_TISSUE_DEVELOPMENT | 138 | 1.49 | 4.57E-03 | 3.14E-02 |
| | GOBP_POSITIVE_REGULATION_OF_CELL_CELL_ADHESION | 245 | 1.49 | 9.40E-04 | 9.87E-03 |
| | GOBP_GLIAL_CELL_DIFFERENTIATION | 224 | 1.49 | 2.12E-03 | 1.82E-02 |
| | GOBP_NEGATIVE_REGULATION_OF_CATALYTIC_ACTIVITY | 150 | 1.49 | 3.21E-03 | 2.46E-02 |
| | GOBP_CELL_CELL_JUNCTION_ASSEMBLY | 123 | 1.48 | 3.38E-03 | 2.56E-02 |
| | GOBP_PIGMENTATION | 103 | 1.48 | 8.01E-03 | 4.69E-02 |
| | GOBP_TRNA_PROCESSING | 135 | 1.48 | 4.46E-03 | 3.08E-02 |
| | GOBP_REGULATION_OF_INNATE_IMMUNE_RESPONSE | 368 | 1.48 | 2.64E-04 | 3.78E-03 |
| | GOBP_MUSCLE_CELL_DEVELOPMENT | 164 | 1.48 | 5.39E-03 | 3.57E-02 |
| | GOBP_REGULATION_OF_PROTEIN_CATABOLIC_PROCESS | 351 | 1.48 | 5.53E-04 | 6.69E-03 |
| | GOBP_NUCLEAR_EXPORT | 161 | 1.48 | 5.76E-03 | 3.73E-02 |
| | GOBP_B_CELL_DIFFERENTIATION | 114 | 1.48 | 8.04E-03 | 4.70E-02 |
| | GOBP_RESPONSE_TO_KETONE | 181 | 1.48 | 3.03E-03 | 2.35E-02 |
| | GOBP_DNA_BIOSYNTHETIC_PROCESS | 154 | 1.48 | 5.36E-03 | 3.55E-02 |
| | GOBP_CARBOHYDRATE_METABOLIC_PROCESS | 471 | 1.47 | 7.78E-05 | 1.42E-03 |
| | GOBP_FATTY_ACID_BIOSYNTHETIC_PROCESS | 123 | 1.47 | 3.76E-03 | 2.73E-02 |
| | GOBP_PROTEINOGENIC_AMINO_ACID_METABOLIC_PROCESS | 127 | 1.47 | 6.91E-03 | 4.21E-02 |
| | GOBP_BONE_DEVELOPMENT | 188 | 1.47 | 3.72E-03 | 2.71E-02 |
| | GOBP_REGULATION_OF_PHOSPHORYLATION | 408 | 1.47 | 2.32E-04 | 3.39E-03 |
| | GOBP_ESTABLISHMENT_OF_PROTEIN_LOCALIZATION_TO_MEMBRANE | 259 | 1.47 | 1.05E-03 | 1.08E-02 |
| | GOBP_REGULATION_OF_CELL_ACTIVATION | 461 | 1.46 | 3.56E-04 | 4.76E-03 |
| | GOBP_TISSUE_HOMEOSTASIS | 211 | 1.46 | 4.80E-03 | 3.25E-02 |
| | GOBP_RESPONSE_TO_TEMPERATURE_STIMULUS | 146 | 1.46 | 7.55E-03 | 4.50E-02 |
| | GOBP_NUCLEOSIDE_DIPHOSPHATE_METABOLIC_PROCESS | 120 | 1.46 | 5.75E-03 | 3.73E-02 |
| | GOBP_POSITIVE_REGULATION_OF_CELL_ACTIVATION | 290 | 1.46 | 1.16E-03 | 1.15E-02 |
| | GOBP_SMALL_MOLECULE_CATABOLIC_PROCESS | 307 | 1.46 | 1.18E-03 | 1.17E-02 |
| | GOBP_NEGATIVE_REGULATION_OF_TRANSPORT | 336 | 1.46 | 1.10E-03 | 1.12E-02 |
| | GOBP_MESENCHYMAL_CELL_DIFFERENTIATION | 228 | 1.45 | 3.49E-03 | 2.61E-02 |
| | GOBP_NUCLEOBASE_CONTAINING_COMPOUND_TRANSPORT | 213 | 1.45 | 4.57E-03 | 3.14E-02 |
| | GOBP_GLAND_DEVELOPMENT | 375 | 1.45 | 4.56E-04 | 5.78E-03 |
| | GOBP_RESPONSE_TO_NUTRIENT | 131 | 1.45 | 5.95E-03 | 3.83E-02 |
| | GOBP_POSITIVE_REGULATION_OF_LEUKOCYTE_CELL_CELL_ADHESION | 206 | 1.45 | 4.11E-03 | 2.91E-02 |
| | GOBP_REGULATION_OF_CELL_CYCLE_G1_S_PHASE_TRANSITION | 164 | 1.45 | 8.03E-03 | 4.69E-02 |
| | GOBP_CONNECTIVE_TISSUE_DEVELOPMENT | 262 | 1.45 | 2.03E-03 | 1.76E-02 |
| | GOBP_HEART_PROCESS | 210 | 1.45 | 6.26E-03 | 3.94E-02 |
| | GOBP_MACROAUTOPHAGY | 365 | 1.44 | 5.27E-04 | 6.48E-03 |
| | GOBP_REGULATION_OF_LIPID_METABOLIC_PROCESS | 262 | 1.44 | 2.12E-03 | 1.82E-02 |
| | GOBP_REGULATION_OF_AUTOPHAGY | 347 | 1.44 | 1.23E-03 | 1.21E-02 |
| | GOBP_ACTIVATION_OF_INNATE_IMMUNE_RESPONSE | 261 | 1.44 | 2.87E-03 | 2.27E-02 |
| | GOBP_STEM_CELL_DIFFERENTIATION | 225 | 1.43 | 4.43E-03 | 3.07E-02 |
| | GOBP_RESPONSE_TO_ALCOHOL | 209 | 1.43 | 5.69E-03 | 3.71E-02 |
| | GOBP_ALCOHOL_METABOLIC_PROCESS | 286 | 1.43 | 2.78E-03 | 2.22E-02 |
| | GOBP_REGULATION_OF_PROTEOLYSIS_INVOLVED_IN_PROTEIN_CATABOLIC_PROCESS | 230 | 1.43 | 4.06E-03 | 2.88E-02 |
| | GOBP_REGULATION_OF_ADAPTIVE_IMMUNE_RESPONSE | 155 | 1.43 | 8.74E-03 | 4.95E-02 |
| | GOBP_NUCLEOTIDE_METABOLIC_PROCESS | 440 | 1.43 | 6.94E-04 | 7.83E-03 |
| | GOBP_NEGATIVE_REGULATION_OF_PROTEIN_MODIFICATION_PROCESS | 203 | 1.43 | 6.67E-03 | 4.11E-02 |
| | GOBP_REGULATION_OF_ACTIN_FILAMENT_ORGANIZATION | 239 | 1.43 | 3.00E-03 | 2.34E-02 |
| | GOBP_REGULATION_OF_BODY_FLUID_LEVELS | 277 | 1.43 | 2.74E-03 | 2.20E-02 |
| | GOBP_POSITIVE_REGULATION_OF_CELL_DEVELOPMENT | 371 | 1.43 | 1.33E-03 | 1.28E-02 |
| | GOBP_CELLULAR_COMPONENT_DISASSEMBLY | 412 | 1.43 | 6.37E-04 | 7.43E-03 |
| | GOBP_ACTIN_FILAMENT_ORGANIZATION | 408 | 1.42 | 6.66E-04 | 7.65E-03 |
| | GOBP_REGULATION_OF_PROTEIN_LOCALIZATION_TO_MEMBRANE | 180 | 1.42 | 6.27E-03 | 3.94E-02 |
| | GOBP_IMMUNE_SYSTEM_DEVELOPMENT | 171 | 1.42 | 7.29E-03 | 4.40E-02 |
| | GOBP_REGULATION_OF_T_CELL_ACTIVATION | 279 | 1.42 | 4.31E-03 | 3.03E-02 |
| | GOBP_LIPID_LOCALIZATION | 355 | 1.41 | 1.40E-03 | 1.33E-02 |
| | GOBP_REGULATION_OF_CELL_JUNCTION_ASSEMBLY | 245 | 1.41 | 3.09E-03 | 2.38E-02 |
| | GOBP_WNT_SIGNALING_PATHWAY | 430 | 1.41 | 8.24E-04 | 8.88E-03 |
| | GOBP_EPIDERMIS_DEVELOPMENT | 253 | 1.41 | 3.37E-03 | 2.56E-02 |
| | GOBP_REGULATION_OF_ENDOCYTOSIS | 250 | 1.40 | 4.55E-03 | 3.13E-02 |
| | GOBP_REGULATION_OF_HORMONE_LEVELS | 407 | 1.39 | 1.34E-03 | 1.28E-02 |
| | GOBP_MUSCLE_ORGAN_DEVELOPMENT | 308 | 1.39 | 2.97E-03 | 2.33E-02 |
| | GOBP_POSITIVE_REGULATION_OF_PROTEIN_LOCALIZATION | 443 | 1.38 | 2.30E-03 | 1.94E-02 |
| | GOBP_RESPONSE_TO_PEPTIDE_HORMONE | 363 | 1.38 | 2.55E-03 | 2.09E-02 |
| | GOBP_CELL_JUNCTION_ASSEMBLY | 452 | 1.37 | 2.46E-03 | 2.03E-02 |
| | GOBP_RESPONSE_TO_RADIATION | 393 | 1.37 | 4.33E-03 | 3.03E-02 |
| | GOBP_POSITIVE_REGULATION_OF_PROTEIN_MODIFICATION_PROCESS | 355 | 1.37 | 2.62E-03 | 2.13E-02 |
| | GOBP_REGULATION_OF_SMALL_MOLECULE_METABOLIC_PROCESS | 279 | 1.37 | 7.94E-03 | 4.67E-02 |
| | GOBP_SENSORY_ORGAN_MORPHOGENESIS | 244 | 1.37 | 8.20E-03 | 4.72E-02 |
| | GOBP_MRNA_CATABOLIC_PROCESS | 245 | 1.37 | 7.54E-03 | 4.50E-02 |
| | GOBP_REGULATION_OF_LYMPHOCYTE_ACTIVATION | 374 | 1.36 | 5.74E-03 | 3.73E-02 |
| | GOBP_PURINE_CONTAINING_COMPOUND_METABOLIC_PROCESS | 498 | 1.35 | 2.33E-03 | 1.95E-02 |
| | GOBP_ORGANIC_ANION_TRANSPORT | 323 | 1.34 | 8.06E-03 | 4.70E-02 |
| | GOBP_MUSCLE_SYSTEM_PROCESS | 355 | 1.34 | 4.78E-03 | 3.24E-02 |
| | GOBP_IMMUNE_RESPONSE_REGULATING_SIGNALING_PATHWAY | 408 | 1.31 | 7.30E-03 | 4.40E-02 |
| | GOBP_PEPTIDYL_AMINO_ACID_MODIFICATION | 391 | 1.31 | 6.85E-03 | 4.20E-02 |
| | GOBP_POSITIVE_REGULATION_OF_MOLECULAR_FUNCTION | 499 | 1.29 | 3.64E-03 | 2.68E-02 |
| | GOBP_REGULATION_OF_PROTEIN_TRANSPORT | 377 | 1.29 | 6.91E-03 | 4.21E-02 |
| | GOBP_NUCLEAR_CHROMOSOME_SEGREGATION | 288 | -1.39 | 6.77E-03 | 4.16E-02 |
| | GOBP_CHROMOSOME_SEGREGATION | 394 | -1.40 | 2.23E-03 | 1.89E-02 |
| | GOBP_MEIOTIC_CHROMOSOME_SEGREGATION | 65 | -1.60 | 7.54E-03 | 4.50E-02 |
| | GOBP_POSTSYNAPTIC_DENSITY_ASSEMBLY | 26 | -1.77 | 3.61E-03 | 2.67E-02 |
| | GOBP_MAINTENANCE_OF_SISTER_CHROMATID_COHESION | 13 | -1.78 | 8.42E-03 | 4.81E-02 |
| | GOBP_FOREBRAIN_REGIONALIZATION | 18 | -1.80 | 5.37E-03 | 3.56E-02 |
| | GOBP_LIGAND_GATED_ION_CHANNEL_SIGNALING_PATHWAY | 34 | -1.80 | 2.68E-03 | 2.17E-02 |
| | GOBP_KINETOCHORE_ORGANIZATION | 21 | -1.85 | 3.85E-03 | 2.77E-02 |
| | GOBP_MITOTIC_SISTER_CHROMATID_COHESION | 31 | -1.85 | 1.86E-03 | 1.65E-02 |
| | GOBP_IONOTROPIC_GLUTAMATE_RECEPTOR_SIGNALING_PATHWAY | 27 | -1.90 | 2.17E-03 | 1.85E-02 |
| | GOBP_REGULATION_OF_NEUROTRANSMITTER_RECEPTOR_ACTIVITY | 12 | -1.97 | 5.11E-04 | 6.37E-03 |

**Table S5. Summary of autophagy-related gene modules defined by the PANTHER Classification System.**

| Category | List ID | Full list name | Abbreviated list name | Total genes | Genes detected in SHOCK fibroblasts | Older vs young t-test p-value in fibroblasts | Genes detected in SHOCK iNs | Older vs young t-test p-value in iNs |
|---|---|---|---|---|---|---|---|---|
| GO Cellular Component | GO:0005776 | autophagosome | autophagosome | 29 | 26 | 0.05 | 28 | 0.59 |
| | GO:0005764 | lysosome | lysosome | 107 | 89 | 0.11 | 98 | 0.34 |
| GO Biological Process | GO:0006914 | autophagy | autophagy | 86 | 73 | 0.04 | 80 | 0.09 |
| | GO:0016236 | macroautophagy | macroatg | 76 | 64 | 0.03 | 70 | 0.13 |
| | GO:0000045 | autophagosome assembly | AP assembly | 58 | 50 | 0.04 | 53 | 0.10 |
| | GO:0097352 | autophagosome maturation | AP maturation | 16 | 12 | 0.03 | 14 | 0.56 |
| | GO:0016239 | positive regulation of macroautophagy | pos reg macroatg | 9 | 8 | 0.09 | 9 | 0.82 |
| | GO:0016242 | negative regulation of macroautophagy | neg reg macroatg | 5 | 4 | 0.23 | 5 | 0.52 |
| Manually curated lists | - | selective autophagy receptors | selective atg | 20 | 10 | 0.21 | 11 | 0.11 |
| | - | chaperone-mediated autophagy | CMA | 21 | 8 | 0.13 | 9 | 0.72 |

Manually curated lists obtained from Khawaja et al., 2025[18].

Table S6. Assignment of autophagy-related genes to GO biological Process modules and regulatory categories.

| Gene name | Detected in SHOCK fibroblasts | Detected in SHOCK iNs | Gene module | | | | | | | | | |
|---|---|---|---|---|---|---|---|---|---|---|---|---|
| | | | autophagosome | lysosome | autophagy | macroautophagy | autophagosome assembly | autophagosome maturation | positive regulation of macroautophagy | negative regulation of macroautophagy | selective autophagy receptors | chaperone-mediated autophagy |
| ACP2 | X | X | | X | | | | | | | | |
| ACP4 | | | | X | | | | | | | | |
| AFG2B | | X | | | X | X | | X | | | | |
| AKT1 | X | | | | | | | | | | | X |
| AKT2 | X | | | | | | | | | | | X |
| AMBRA1 | X | X | | | X | X | X | | | | | |
| AP5M1 | X | X | | X | | | | | | | | |
| AP5S1 | X | X | | X | | | | | | | | |
| ARL8A | X | X | | X | | | | | | | | |
| ARL8B | X | X | | X | | | | | | | | |
| ATG10 | X | X | | | X | X | X | | | | | |
| ATG101 | | X | | | X | X | X | | | | | |
| ATG12 | X | X | X | | X | X | X | X | | | | |
| ATG13 | X | X | | | X | X | X | | | | | |
| ATG14 | X | X | X | | X | X | X | X | | | | |
| ATG16L1 | X | X | X | | X | X | X | | | | | |
| ATG16L2 | X | X | X | | X | X | X | | | | | |
| ATG2A | X | X | | | X | X | X | | | | | |
| ATG2B | X | X | | | X | X | X | | | | | |
| ATG3 | X | X | | | X | X | X | | | | | |
| ATG4A | X | X | | | X | X | X | | | | | |
| ATG4B | X | X | | | X | X | X | | | | | |
| ATG4C | X | X | | | X | X | X | | | | | |
| ATG4D | X | X | | | X | X | X | | | | | |
| ATG5 | X | X | X | | X | X | X | | | | | |
| ATG7 | X | X | | | X | X | X | | | | | |
| ATG9A | X | X | X | | X | X | X | | | | | |
| ATG9B | X | X | X | | X | X | X | | | | | |
| ATP13A2 | X | X | | | X | X | | | | | | |
| ATP2A2 | X | X | | | X | X | X | | | | | |
| ATP6V0D1 | X | X | | X | | | | | | | | |
| ATP6V0D2 | | X | | X | | | | | | | | |
| ATP6V1A | X | X | | X | | | | | | | | |
| ATP6V1C1 | X | X | | X | | | | | | | | |
| ATP6V1C2 | | X | | X | | | | | | | | |
| BECN1 | X | X | | | X | X | X | | | | | |
| BECN2 | | | | | X | X | X | | | | | |
| BMAL1 | | | | | | | | | | | X | |
| BMF | X | X | | | | | | | | | | |
| BNI3L | X | | | | | | | | | | X | |
| BNIP3 | X | X | | | | | | | | | X | |
| BORCS5 | | X | | X | | | | | | | | |
| CACO1 | | | | | | | | | | | X | |
| CALCOCO2 | X | X | X | | X | X | | | X | | | |
| CD68 | X | X | | X | | | | | | | | |
| CDIP1 | X | X | | X | | | | | | | | |
| CLEC16A | X | X | | | | | | | | | | |
| CLN5 | X | X | | X | | | | | | | | |
| CST7 | | | | X | | | | | | | | |
| CTSB | X | X | | X | | | | | | | | |
| CTSC | X | X | | X | | | | | | | | |
| CTSF | X | X | | X | | | | | | | | |
| CTSH | X | X | | X | | | | | | | | |
| CTSK | X | X | | X | | | | | | | | |
| CTSL | X | X | | X | | | | | | | | |
| CTSO | X | X | | X | | | | | | | | |
| CTSS | X | X | | X | | | | | | | | |
| CTSV | X | X | | X | | | | | | | | |
| CTSW | X | X | | X | | | | | | | | |
| CTSZ | X | X | | X | | | | | | | | |
| CYB561 | X | X | | X | | | | | | | | |
| CYB561A3 | X | X | | X | | | | | | | | |
| CYBRD1 | X | X | | X | | | | | | | | |
| DAP | X | X | | | | | | | | | | |
| DAPL1 | | X | | | | | | | | | | |
| DEPDC5 | X | X | | | | | | | | | | |
| DEPP1 | | X | | | | | | | | | | |
| DNJB1 | | | | | | | | | | | | X |
| DRAM1 | X | X | | X | | | | | | | | |
| DRAM2 | X | X | | X | | | | | | | | |
| EF1A1 | | | | | | | | | | | | X |
| EI24 | X | X | | | X | X | | | | | | |
| ELAPOR1 | | X | | X | X | X | X | | | | | |
| EMC6 | X | X | | | X | X | X | | | | | |
| EPDR1 | X | X | | X | | | | | | | | |
| EPG5 | X | X | | | X | X | | X | | | | |
| FIS1 | X | X | | | X | | | | | | | |
| FUCA1 | X | X | | X | | | | | | | | |
| FUCA2 | X | X | | X | | | | | | | | |
| FUNDC1 | X | X | | | X | | | | | | | |
| FUNDC2 | X | X | | | X | | | | | | | |
| FYCO1 | X | X | X | X | | | | | X | | | |
| GAA | X | X | | X | | | | | | | | |
| GABARAP | X | X | X | | X | X | X | X | | | | |
| GABARAPL1 | X | X | X | | X | X | X | X | | | | |
| GABARAPL2 | X | X | X | | X | X | X | X | | | | |
| GABARAPL3 | | | X | | X | X | X | X | | | | |
| GALC | X | X | | X | | | | | | | | |
| GFAP | X | X | | | | | | | | | | X |
| GLMP | | X | | X | | | | | | | | |
| GPR137 | X | X | | X | | | | | | | | |

| | | | | | | | | | | | | |
|---|---|---|---|---|---|---|---|---|---|---|---|---|
| GPR137B | X | X | | X | | | | | | | | |
| GPR137C | X | X | | X | | | | | | | | |
| HEXA | X | X | | X | | | | | | | | |
| HEXB | X | X | | X | | | | | | | | |
| HPS6 | X | X | | X | | | | | | | | |
| HS90A | | | | | | | | | | | | X |
| HS90B | | | | | | | | | | | | X |
| HSP7C | | | | | | | | | | | | X |
| IRGM | | | | | X | X | X | | | | | |
| KAT6A | X | | | | | | | | | | | X |
| LAMP1 | X | X | | X | | | | | | | | |
| LAMP2 | X | X | | X | | | | | | | | X |
| LAMP3 | X | X | | X | | | | | | | | |
| LAMP5 | X | X | | X | | | | | | | | |
| LAMTOR4 | X | X | | X | | | | | | | | |
| LAMTOR5 | X | X | | X | | | | | | | | |
| LAPTM4A | X | X | | X | | | | | | | | |
| LAPTM4B | X | X | | X | | | | | | | | |
| LAPTM5 | X | X | | X | | | | | | | | |
| LITAF | X | X | | X | | | | | | | | |
| LITAFD | | | | X | | | | | | | | |
| LIX1 | | | | | X | X | | X | | | | |
| LIX1L | X | X | | | X | X | | X | | | | |
| MAN2B2 | X | X | | X | | | | | | | | |
| MAP1LC3A | X | X | X | | X | X | X | X | | | | |
| MAP1LC3B | X | X | X | | X | X | X | X | | | | |
| MAP1LC3B2 | X | X | X | | X | X | X | X | | | | |
| MAP1LC3C | X | X | X | | X | X | X | X | | | | |
| MARCHF1 | | X | | X | | | | | | | | |
| MARCHF8 | | X | | X | | | | | | | | |
| MCOLN1 | X | X | | X | | | | | | | | |
| MCOLN2 | X | X | | X | | | | | | | | |
| MCOLN3 | X | X | | X | | | | | | | | |
| MTM1 | X | X | | | | | | | | X | | |
| MTMR3 | X | X | | | | | | | | | | |
| MTMR4 | X | X | | | | | | | | | | |
| MTMR9 | X | X | | | | | | | | | | |
| MTOR | X | X | | | | | | | | X | | |
| NAPSA | X | X | | X | | | | | | | | |
| NBR1 | X | X | | | X | X | | | | | X | |
| NCOA4 | X | X | | | | | | | | | X | |
| NCOR1 | X | | | | | | | | | | | X |
| NF2L2 | | | | | | | | | | | | X |
| NFAC1 | | | | | | | | | | | | X |
| NPRL2 | X | X | | | | | | | | | | |
| NPRL3 | X | X | | | | | | | | | | |
| NRBF2 | X | X | | | X | | | | | | | |
| NSFL1C | X | X | | | X | X | X | | | | | |
| NUFP1 | | | | | | | | | | | X | |
| OPTN | X | X | | | | | | | | | X | |
| P53 | | | | | | | | | | | | X |
| PHB2 | X | X | | | | | | | | | X | |
| PHF23 | X | X | | | | | | | | X | | |
| PHLP1 | | | | | | | | | | | | X |
| PI4K2A | X | X | | X | | | | | | | | |
| PI4K2B | X | X | | X | | | | | | | | |
| PIK3C3 | X | X | | | X | X | X | | | | | |
| PIK3R4 | X | X | | | X | X | | | | | | |
| PINK1 | X | X | | | X | | | | | | X | |
| PIP4P1 | | X | | X | | | | | | | | |
| PIP4P2 | | X | | X | | | | | | | | |
| PLA2G15 | X | X | | X | | | | | | | | |
| PPGB | | | | | | | | | | | | X |
| PPT1 | X | X | | X | | | | | | | | |
| PPT2 | X | X | | X | | | | | | | | |
| PRKN | | X | | | X | | | | | | X | |
| RAB19 | | | | | X | X | X | | | | | |
| RAB1A | X | X | | | X | X | X | | | | | |
| RAB1B | X | X | | | X | X | X | | | | | |
| RAB1C | | | | | X | X | X | | | | | |
| RAB23 | X | X | X | | X | X | X | | | | | |
| RAB24 | X | X | X | | | | | | | | X | |
| RAB33A | X | X | | | X | X | X | | | | | |
| RAB33B | X | X | | | X | X | X | | | | | |
| RAB43 | X | X | | | X | X | X | | | | | |
| RAB7A | X | X | | X | | | | | | | | |
| RAB7B | | | | X | | | | | | | | |
| RAB9A | X | X | | X | | | | | | | | |
| RAB9B | X | X | | X | | | | | | | | |
| RAC1 | X | | | | | | | | | | | X |
| RARA | X | | | | | | | | | | | X |
| RB11A | | | | | | | | | | | | X |
| RB1CC1 | X | X | | | X | X | X | | | | | |
| RETR1 | | | | | | | | | | | X | |
| RETREG1 | | X | | | X | X | | | | | | |
| RICTR | | | | | | | | | | | | X |
| RMC1 | | X | | X | | | | | | | | |
| RNF19B | X | X | | X | | | | | | | | |
| RPTOR | X | X | | | | | | | | | | |
| RRAGA | X | X | | X | | | | | | | | |
| RRAGB | X | X | | X | | | | | | | | |
| RRAGC | X | X | | X | | | | | | | | |
| RRAGD | X | X | | X | | | | | | | | |
| RTN3 | X | X | | | | | | | | | X | |
| RUBCN | | X | | | | | | | | X | | |
| RUBCNL | | X | X | | X | X | | X | | | | |

| | | | | | | | | | | | |
|---|---|---|---|---|---|---|---|---|---|---|---|
| SESN1 | X | X | | | | | | | X | | |
| SESN2 | X | X | | | | | | | X | | |
| SESN3 | X | X | | | | | | | X | | |
| SIDT1 | | X | | X | | | | | | | |
| SIDT2 | X | X | | X | | | | | | | |
| SLC11A1 | X | | | X | | | | | | | |
| SLC11A2 | X | X | | X | | | | | | | |
| SLC15A3 | X | X | | X | | | | | | | |
| SLC38A9 | X | X | | X | | | | | | | |
| SLC46A3 | X | X | | X | | | | | | | |
| SLC48A1 | X | X | | X | | | | | | | |
| SLC66A1L | | | | X | | | | | | | |
| SNX30 | X | X | | | X | X | | | | | |
| SNX4 | X | X | | | | | | | X | | |
| SNX7 | X | X | | | X | X | | | | | |
| SPATA18 | X | X | | | X | | | | | | |
| SPPL2A | X | X | | X | | | | | | | |
| SPPL2B | X | X | | X | | | | | | | |
| SPPL2C | | X | | X | | | | | | | |
| SQSTM1 | X | X | X | | X | X | | | | | |
| STARD3 | X | X | | X | | | | | | | |
| STARD3NL | X | X | | X | | | | | | | |
| STBD1 | X | X | | | | | | | | | X |
| STING1 | | X | X | | X | X | X | | X | | |
| STX12 | X | X | | | X | X | X | | | | |
| SVIP | X | X | | | | | | | | | |
| TBC1D12 | X | X | X | | | | | | | | |
| TBC1D14 | X | X | X | | | | | | | | |
| TGFBRAP1 | X | X | | | X | | | | | | |
| TINAG | | | | X | | | | | | | |
| TINAGL1 | X | | | X | | | | | | | |
| TM6SF1 | X | | | X | | | | | | | |
| TMEM192 | X | X | | X | | | | | | | |
| TMEM39A | X | X | | | | | | | | X | |
| TMEM59 | X | X | | X | | | | | | | |
| TMEM79 | X | X | | X | | | | | | | |
| TOLIP | | | | | | | | | | | X |
| TP53INP1 | X | X | X | | X | X | X | | | | |
| TP53INP2 | X | X | X | | X | X | X | | | | |
| TPCN1 | X | X | | X | | | | | | | |
| TPCN2 | X | X | | X | | | | | | | |
| TRI16 | | | | | | | | | | | X |
| TRIM13 | X | X | | | | | | | X | | |
| TRIM23 | X | X | | X | | | | | | | |
| UBQLN2 | X | X | | | X | X | X | | | | |
| UBXN2A | X | X | | | X | X | X | | | | |
| UBXN2B | X | X | | | X | X | X | | | | |
| ULK1 | X | X | X | | X | X | X | | | | |
| ULK2 | X | X | X | | X | X | X | | | | |
| ULK3 | X | X | X | | X | X | X | | | | |
| UNC93B1 | X | X | | X | | | | | | | |
| VCP | X | X | | | X | X | | X | | | |
| VPS13A | X | X | | | X | | | | | | |
| VPS16 | X | X | | X | | | | | | | |
| VPS33A | X | X | | X | | | | | | | |
| VPS39 | X | X | | | X | | | | | | |
| VPS41 | X | X | | | X | X | | | | | |
| VPS4A | X | X | | | | | | | | | X |
| VTI1B | X | X | | | X | X | | | | | |
| WAC | X | X | | | | | | | | | |
| WDFY3 | X | X | | | X | X | | | | | |
| WDR24 | X | X | | | | | | | X | | |
| WDR45 | X | X | | | X | X | X | | | | |
| WDR45B | X | X | | | X | X | X | | | | |
| WIPI1 | X | X | | | X | X | X | | | | |
| WIPI2 | X | X | | | X | X | X | | | | |

X - gene detected or included in module list

**Table S7. Demographics and health measures of the STRONG cohort before and after a 12-week exercise intervention.**

| STRONG donor ID | Number of classes attended out of 12 | Demographics | | | | Health and fitness readouts | | | | | | | | | | | | | | | | | |
|---|---|---|---|---|---|---|---|---|---|---|---|---|---|---|---|---|---|---|---|---|---|---|---|
| | | Age (years) | Sex | Race | Hispanic/ Latino? | SPPB score | | BMI (kg/m$^2$) | | Systolic blood pressure (mmHg) | | Diastolic blood pressure (mmHg) | | Resting heart rate (bpm) | | Maximal grip strength (kg) | | Chair stand time (s) | | Occipital wall distance, normal posture (cm) | | Occipital wall distance, tall posture (cm) | |
| | | | | | | Week 0 | Week 12 | Week 0 | Week 12 | Week 0 | Week 12 | Week 0 | Week 12 | Week 0 | Week 12 | Week 0 | Week 12 | Week 0 | Week 12 | Week 0 | Week 12 | Week 0 | Week 12 |
| 7 | 11 | 84 | F | White | | 6 | 8 | 23.9 | 24.6 | 135 | 144 | 73 | 68 | 54.5 | 58.5 | 14 | 10 | - | - | 10.7 | 8.3 | 6.7 | 7.9 |
| 8 | 10 | 83 | F | White | | 10 | 12 | 26.5 | 27.0 | 127 | 113 | 84 | 73 | 79.0 | 92.0 | 16 | 30 | 14.38 | 8.84 | 0.0 | 2.0 | 0.0 | 0.5 |
| 9 | 9 | 88 | M | White | | 9 | 10 | 26.4 | 26.4 | 125 | 134 | 51 | 75 | 55.0 | 61.5 | 18 | 20 | 25.32 | 16.59 | 8.6 | 7.3 | 6.4 | 4.5 |
| 11 | 7 | 87 | M | White | | 7 | 8 | 31.6 | 32.0 | 137 | 131 | 62 | 55 | 58.5 | 67.5 | 23 | 23 | 42.28 | 25.03 | 12.3 | 8.5 | 7.2 | 7.9 |
| 13 | 12 | 77 | M | White | | 11 | 11 | 29.2 | 30.2 | 142 | 128 | 69 | 66 | 60.0 | 73.5 | 32 | 30 | 10.16 | 10.97 | 7.7 | 9.7 | 6.5 | 6.0 |

M, male; F, female; SPPB, Short Physical Performance Battery; BMI, body mass index, chair stand time - in SPPB test.

**Table S8. Demographics and health measures of the SHOCK PBMC cohort and STRONG cohort before and after a 12-week exercise intervention.**

| Demographics, health and fitness readouts | SHOCK PBMC cohort | | STRONG cohort | | | | Paired t-test p-value | Welch's t-test p-value | |
|---|---|---|---|---|---|---|---|---|---|
| | | | Week 0 | | Week12 | | STRONG Week 0 vs Week 12 | SHOCK vs STRONG Week 0 | SHOCK vs STRONG Week 12 |
| | Mean | Standard deviation | Mean | Standard deviation | Mean | Standard deviation | | | |
| Number of participants | 23 | | 5 | | 5 | | | | |
| Age (years) | 60.8 | 16.7 | 83.8 | 4.3 | 83.8 | 4.3 | | <0.0001 | |
| Sex (% male) | 47.8 | | 60 | | 60 | | | | |
| Race (% White) | 95.7 | | 100 | | 100 | | | | |
| Hispanic/Latino (%) | 21.7 | | 0 | | 0 | | | | |
| SPPB score | 11.8 | 0.9 | 8.6 | 2.1 | 9.8 | 1.8 | 0.03 | 0.03 | 0.07 |
| BMI ($kg/m^2$) | 24.5 | 2.8 | 27.5 | 2.9 | 29.0 | 4.7 | 0.20 | 0.08 | 0.10 |
| Systolic blood pressure (mmHg) | 123 | 16 | 133 | 7 | 130 | 11 | 0.55 | 0.05 | 0.30 |
| Diastolic blood pressure (mmHg) | 76 | 6 | 68 | 12 | 67 | 8 | 0.95 | 0.20 | 0.06 |
| Resting heart rate (bpm) | 60 | 9 | 61 | 10 | 71 | 13 | 0.007 | 0.71 | 0.12 |
| Maximal grip strength (kg) | 39 | 12 | 21 | 7 | 23 | 8 | 0.56 | 0.001 | 0.007 |
| Chair stand time (s) | 1.9 | 0.2 | 23.0 | 14.3 | 15.4 | 7.2 | 0.13 | 0.06 | 0.03 |
| Occipital wall distance, normal posture (cm) | N/A | | 7.9 | 4.7 | 7.2 | 3.0 | 0.58 | | |
| Occipital wall distance, tall posture (cm) | N/A | | 5.4 | 3.0 | 3.0 | 3.1 | >0.99 | | |

SPPB, Short Physical Performance Battery; BMI, body mass index; walking speed - in 6-minute walk test; chair stand time - in SPPB test
